# A non-genetic meiotic repair program inferred from spore survival values in fission yeast wild isolates: a clue for an epigenetic ratchet-like model of ageing?

**DOI:** 10.1101/223685

**Authors:** Xavi Marsellach

## Abstract

What is the nature of the ageing process? What is the spore survival, that one would expect upon analysing a self-cross, in a wild fission yeast strain? Could this two research questions be, somehow, related? In this manuscript, I am describing some interesting observations obtained while studying fission yeast spore survival values upon genetic crosses. Early findings brought my attention into mainly studying self-crosses (intra-strain crosses in which any cell can be involved in by mating with a sibling cell). This study, yield some interesting findings. As a summary: 1) most fission yeast self-crosses do show low spore survival values; 2) clonally related strains show a high phenotypic variability in self-cross spore survival values; 3) differences in self-cross spore survival values can be detected when comparing zygotic and azygotic matings; 4) self-cross spore survival values are highly affected by environmental factors, mainly producing a reduction in the spore survival values; 5) self-cross spore survival values are “recovered” when cells are subjected to several rounds of meiotic divisions; 6) signs of correlation between spore survival and vegetative cell survival (prior to the entry into meiosis) have been observed in this study. All those observations, among others, are discussed as part of an epigenetic variability that exist in fission yeast populations. A cyclical behaviour, of this epigenetic variability it is proposed, defining an underlying ratchet-like epigenetic mechanisms acting in all cells. In this manuscript, I propose that this mechanism, is, indeed, the main cause of the ageing process.

## 1. Introduction

Fission yeast cells, like most of the other unicellular eukaryotic cells^1^, can give rise to progeny by means of two types of cell divisions: 1) mitotic divisions, which gives rise to two genetically identical daughter cells (clonal cells), or 2) meiotic divisions, which give rise to four daughter cells. Meiotically produced daughter cells can either be genetically different daughter cells, i.e. when they arise from a cross involving two genetically different parental cells (inter-strain cross), or almost genetically identical daughter cells, i.e. when they arise from a cross of two almost genetically identical parental cells (intra-strain cross or self-cross). In pluricellular organisms, this two types of cell divisions happen in differentiated group of cells: somatic cells divide by means of mitotic divisions^1^, while germline cells ultimately give rise to gametes by means of a meiotic cell division. Gametes are responsible to give rise to the next generation of individuals.

The ageing process is the set of phenomena that occur from the birth of a new individual, generated from the fusion of two gametes in sexually dividing cells, up to their death. If a given individual does not die from an accidental cause, the loss of proper cell function, caused by the ageing process, will ultimately produce the death of this individual. The most accepted consensus in the ageing community is that ageing is a complex phenomenon with a multifactorial origin. Nine candidate hallmarks have been proposed as the major contributors to the ageing phenomena^2^, but despite this, the nature of the ageing process, and which pathways, or combination of pathways, generate the ageing phenomena, and why, are still an open debate. Several active controversies are still open inside the ageing research community. Just to name a few: ageing as a programmed *vs* a non-programed process^3–7^, the role of sirtuins in ageing^8–10^, the role of reactive oxygen species (ROS) in the ageing process^11–15^, or whether ageing is a disease or not^16–20^. In summary, there is no consensus agreement on what it is ageing *per se*, and why ageing exist at all. Many theories to explain the ageing process has been proposed to date (see some of the reviews available in the literature: as an example^6,16,21–29^). Most of the research that is being carried out nowadays by the leading scientist in the ageing research community is part of what have been called "the new biology of ageing"^30,31^. This denomination is mainly referring to the unexpected discovery that the rate of ageing can be modulated, at least to some extent, by genetic pathways, biochemical processes, and environmental modifications, like dietary restriction, that are conserved during evolution. Strikingly, in the ageing research community, even if this approach implies a new paradigm, or is just the continuation of the "old biology of ageing" is a controversial issue^32^. Nevertheless, the founding papers of this so-called "new biology of ageing" all share as common denominator that they all focus in lifespan modifications^33–48^.

A surprising observation, when it comes to the research that it is been carried out in the ageing field, is the scarce importance that is given to the different behaviour shown between mortal somatic cells (affected by the ageing process) and amortal germline cells (able to restart the ageing process, and therefore immune to the inevitable end of the ageing process: death). This is an observation that has been present in the ageing research field from the very beginning^49,50^, but despite this, just a few studies focus on this different behaviour^51–56^ as a central aspect to study ageing. In this manuscript, using fission yeast and spore survival data, I show an interesting phenomenon related to spore survival and vegetative cell survival, that, focusing on the different behaviour between mitotic and meiotic cells, might provide a simple and coherent explanation of the ageing process (see later).

Unlike in the budding yeast *Saccharomyces cerevisiae*, nearly all the studies that have been done in the fission yeast *Schizosaccharomyces pombe* are isogenic^57,58^, and derive from isolates characterized by Urs Leupold, the founder of *S. pombe* genetics^59,60^. Leupold began working with *S. pombe* in 1946 when he joined Øjvind Winge’s laboratory, the founder of budding yeast genetics. Winge suggested Leupold to study the basis of *S. pombe* homothalism as his *PhD* topic^60^. From the work carried out by Leupold in Winge’s laboratory, and Leupold’s further studies, three main strains became stablished as the reference strains in the growing community of researchers that had started to work with fission yeast as a model organism^61^: Leupold’s 968 (*h*^90^), 972 (*h^-s^*), and 975 (*h^+N^*). Those strains will be thereafter referred as L968, L972 and L975. All those three isolates come from a single culture obtained by Leupold from the *Centraalbureau voor Schimmelcultures* in Delft, The Netherlands. This culture was originally isolated by A. Osterwalder in Switzerland in 1924^59,60^. The two heterothallic strains (L972 and L975) derived from the homothallic strain (L968), which, spontaneously, give rise to heterothallic derivatives^59,62^. More details about the strains isolated by Leupold and colleagues, and their relationships, can be obtained from the literature^59,60,62–65^. Leupold’s strains are therefore considered as isogenic strains by the fission yeast community, and, indeed, in a recent study where two of those isolates were fully sequenced (L968 and L972), only 14 SNPs differences were found between them^58^.

Determination of the spore survival values after a meiotic division in yeast can be precisely measured by means of tetrad analysis (see below). Despite that, the spore survival values of the wild type fission yeast strains (all derived from Leupold’s isolates), have been measured, using tetrad analysis, in several previous studies showing no consistent results between them: some studies show wild type survivals close to 95%^66–68^, while some others show survivals close to 80%^69–71^. Most studies use the determination of wild type spore survival just as a value to compare with the mutant of interest. Given that all the *S. pombe* strains used in the community are nearly isogeneic, it remains to be answered why this variability in wild type spore survival determination is observed. In other words, there is no clear answer on which is the survival value that one would expect form a tetrad analysis carried out in a Leupold’s derived wild strain.

In recent studies, a growing interest for non-Leupold’s isolates is seen in the fission yeast community^58,67,72–77^. In one of those studies I collaborated determining the spore variability for 43 inter-strain crosses between different fission yeast wild isolates^58^. From this analysis, it was concluded that these strains have started to accumulate genetic differences that contribute to the reproductive isolation between them. A surprising observation was, as well, obtained in this analysis: the spore survival values obtained in self-crosses from those fission yeast wild isolates gave reduced spore survival values (see Supplementary Figure 1 in Jeffares et al 2015^58^, and bellow). The lowest value observed in a self-cross of a fission yeast wild isolate is as low as 9,78% survival (see below for detailed data). An accurate review of the literature showed that, indeed, this phenomenon was already described even in pioneering studies using fission yeast wild isolates other than Leupold’s ones^63,78^. In fact, even the proper foundation of the *S. pombe* genetic research showed this phenomenon: the very first isolate that Leupold was given in Winge’s laboratory showed fewer than 50% spore survival in a self-cross, and therefore was discarded by Leupold as a non-useful isolate for genetic research^60^.

In this study, I analyse in more detail the phenomenon of the observed lower spore survival values obtained from self-crosses in fission yeast wild type strains. This analysis gave several unexpected observations, the most interesting of which are: 1) high phenotypic differences are seen in clonally related strains when analysing the spore survival obtained from self-crosses; 2) environmental factors induces variation of the self-cross spores survival values in a given strain; 3) there are preliminary observations that suggest that, it might be a correlation of the low spore survival values with the lower cell survival of the vegetative cells (prior to the entry into meiosis); and 4) a non-genetic recovery (of the low spore survival values and of the vegetative cell fitness) is seen when several meiotic cycles are carried out. All those observations, together with more data, are discussed as part of a cyclical variation of non-genetic factors, likely epigenetic marks. This leads to the proposal of epigenetic ratchet-like mechanisms in analogy with the well-known genetic Muller’s ratchet.

In this manuscript, I propose several key follow-up experiments, to test this hypothesis. My current situation as an independent researcher, doesn’t allow me to carry out them just by myself. With this manuscript, I pretend to expose all the data that I have obtained so far to the scientific community, and to reason that, if some of the observations, obtained in this study, are proven to be the rule rather than the exception, this leads to a simple and coherent model to describe the ageing process as an epigenetically-based ratchet-like mechanisms, and to conceive ageing itself as a loss of epigenetic information phenomenon. The ultimate purpose of this manuscript is to gather supports, from fellow scientists, to consider aging as a process of epigenetic loss of information, and to contribute to a change on the focus on ageing research. In my opinion, ageing and anti-ageing research should be focussed into the mechanisms which leads to the loss of epigenetic information (ageing research), and the meiotic events that reverse ageing, through the recovery of lost epigenetic information (anti-ageing research). This is not the actual consensus in the ageing research community, which doesn’t yet agree in a common definition of what is ageing *per se*, and spend most of the research efforts from the ageing field in lifespan modifying factors (i.e. genetic pathways, biochemical processes, and environmental modifications). Finally, this manuscript introduces the definition the epigenetic degeneration phenomena, as a possible cause of many diseases, not initially related to ageing. These phenomena are proposed to occur, as well as ageing, due to the loss of epigenetic information. The epigenetic degeneration processes are described as the loss of epigenetic information due to non-full repair of epigenetic defects in-between two generations.

## 2. Materials and methods

### Yeast strain, media, vectors and general methods

Basic cell growth and media conditions were used according to standard fission yeast techniques^79,80^. To generate diploid hybrids or diploid homozygotic strains a customized protocol were used to ensured that mating took place between the two desired strains^73^. In short: strains were tagged by deleting the *ade6 locus* with the insertion of a dominant marker (Kanamycin, Nourseothricin or Hygromycin^81^). This allow to perform directed crosses by crossing differentially tagged strains, and select for the desired diploids by platting the mating mixture in a double selection plates accordingly. This strategy was used both, to obtain diploid hybrid strains (from inter-strain crosses), and diploid homozygotic strains (from self-crosses). Selected double resistant clones were tested for its ability to produce *asci* by plating them on Malt Extract plates (MEA) and inspect them under the microscope. This allow to discard the rare *ade6* intra-genic recombinants with double markers at this *locus*. Two independently isolated diploid strains were kept in each cross for further analysis. Three vectors were generated to routinely delete the *ade6 locus* in the desired strains, and replace it by a dominant selectable marker: pΔade6-K (Kanamycin), pΔade6-N (Nourseothiricin), and pΔade6-H (Hygromycin). To generate those vectors the methods described by Gregan and collaborators were used^82^. The vectors pΔade6-N and pΔade6-H were generated straight from the pCloneNat1 and pCloneHyg1 vectors available from the Gregan paper. To build the vector pΔade6-K, a modified version of the Gregan vectors, containing Kanamycin as selectable marker (the pCloneKan1 vector) was constructed by replacing the BglII-EcoRI region of the pCloneNat1 vector with the BglII-EcoRI region of the pFA6a-KanMX6 vector^83^. This strategy gave three vectors, with three different dominant selectable markers, that easily generate a deletion cassette with 735bp homology region in the 5’ region of the *ade6 locus* and 350bp homology region in the 3’ region of the same locus, just by digesting the desired vector with the XbaI restriction enzyme^82^. The PCR products to generate those plasmids were done using the DNA from the reference genome (the L972 strain^84^), but thanks to the long homology regions present on those vectors one can easily transform distantly related fission yeast strains successfully.

The strain JB32 is a Leupold’s derived strain, as in the Heim’s 1990 paper^85^, it is stated that this *h*^+s^ strains were constructed using Leupold’s derived strain. In this report, though, a unique clonal background (L968, L972 or L975), it is not indicated as the source for the creation of this strains. It is therefore a wrong assumption, as has recently been done^86^, to deduce that the JB32 strain (*h*^+s^: *h*^+^ stable wild type Leupold’s derived, obtained on purpose by Heim in 1990^85^) it is derived from the L975 strain (*h*^+N^: *h*^+^ unstable wild type Leupold’s derived, isolated by Leupold in 1950^59^.

Two more vectors were generated in this study: pMTS-h+ and pMTS-h- (MTS= Mating Type Switching). Those vectors target the *mat1 locus*, and allow to specifically switch the mating type of any given strain with a simple transformation protocol. Those vectors are built in a way that once inserted in any given strain, this strain is transformed into a non-switchable strain (stable heterothallic) of the chosen mating type (*h*^+^ for the pMTS-H+ vector and *h*^-^ for the pMTS-H- vector). To build those vectors, the following oligonucleotides: 5’-TTTGGAATTGGGAGGTTGAG-3’ and 5’-GAGTGGTTGAGCGAGGAAAG- 3’ were used to specifically amplify by PCR the *mat1 locus*. Two PCR reactions were carried out: one using as a template the DNA obtained from the strains JB32 (*h*^+^ non-switchable strain) and the other with DNA from the strain L972 (*h*^-^ non-switchable strain). Two clear DNA bands of approximately 2400bp were obtained with those PCRs. Those bands were cloned into the pCR2.1-TOPO TA vector (Invitrogen, life technologies). Subsequently, the Kanamycin cassette was added to those vectors by digesting the Kanamycin cassette from the pFA6a-KanMX6 vector^83^ with the SmaI and EcoRV restriction enzymes (both generate blunt ends) and by cloning the 1436bp band obtained into the previously obtained vectors digested with the SanDI restriction enzyme. The SanDI restriction enzyme generates 5’- overhang ends, that were converted to blunt ends with the NEBNext end Repair Module (New England Biolabs). The following vectors, though, were not yet useful to build a stable non-switchable strain (although they were successfully targeted, if the given strain has a functional *mat2-mat3 locus* the transformed strain would still be able to switch back the mating type). To avoid this problem, the H1 region in the *mat1 locus*, indispensable for the mating type switching reaction, were eliminated from the cloned vectors by doing a circular PCR in the vectors already obtained with the oligonucleotides: 5’- TTTCCAATTATGCTGTTCGTG-3’ in both vectors, and the oligonucleotide 5’- TTTAATGGGCCAATTCTACGA-3’ in the vector containing the *mat1P locus* or the oligonucleotide 5’-TTATTGAAAATAAATAAAAACG-3’ in the vector containing the *mat1M locus*. The PCR products obtained with this PCRs were treated with NEBNext end Repair Module (New England Biolabs) to repair ends to achieve self-ligation of those PCR products (PCR products normally do not self-ligate because commercially bought oligonucleotides do not have the 5’ phosphate needed for the ligation reaction). All this protocol generated the vectors named pMTS-h+ and pMTS-h- which were used to intentionally modify the *mat1 locus* of any given strain to produce non-switchable stable heterothallic strains (see main text). This author wants to point out that, when using those vectors, the user must carefully check that the H1 region is missing in the transformed strain, otherwise although the dominant selectable marker it is correctly inserted in the *mat1 locus* of the transformed strain, this strain might not be non-switchable if it contains a working *mat2-mat3 locus*. This author would like to encourage the reader to rebuild those vectors by putting the dominant selectable marker exactly in the region were the H1 region is deleted to avoid this problem.

Some spontaneous mating type revertants were isolated thanks to the double selection procedure: the double selection procedure described above allowed to isolate mating type variants in some of the naturally isolated heterothallic strains. Differential tagging in *ade6 locus* were carried out in different clones of a given strain. Mass mating and replica plating in double selection plates should give no clones, unless a spontaneous event of mating type switching occurred randomly. In some cases, no colonies were observed in repeated experiments, while in some other examples colonies do appear. Those colonies were tested for *asci* production (to check that they are diploid cells), and those *asci* were dissected to isolate a haploid strain with the spontaneously isolated mating type revertant.

Each strain is coloured in a different colour through the paper. To highlight strain similarity (both in clonally related strains or in cluster grouped strains), similar colours are used to plot related strains, although there is no linear relationship between colour similarity and strain similarity (i.e. in clonally related strains, strain differences are much lower than in cluster grouped strains). All inter-strain crosses are plotted in grey.

A detailed list of the strains generated with those methods can be found in the supplemental Table S1.

### Spore survival determination

To determine the spore survival of diploid strains (both diploid hybrids and diploid homozygotes), the desired strains were grown from -80°C stock tubes into YES plates containing the two dominant markers that were used to generate the given strain (see above). This was done to ensure that cells do not revert to haploid cells by spontaneously going through meiosis, as seems to be the norm in fission yeast^87^. A loop of the cells grown in the YES plates was subsequently patched into Malt Extract plates (MEA) or Minimal media plates without Nitrogen (MM-N; Formedium, UK) plates, and incubated at 25°C for 2 to 5 days. Visual inspection was carried out using a microscope, and, when the *asci* were ready, the tetrads were dissected using an MSM400 micromanipulator (Singer Instruments, UK).

As described in the main text, in this paper, not only azygotic meiosis were analysed (azygotic meiosis = meiosis from a stable diploid strain). Zygotic meiosis, those than arise straight from a mating mixture of haploid cells, were, as well, analysed (see bellow). This procedure is named, as well, as mass mating along the paper. To determine zygotic meiosis two scenarios were contemplated: 1) homothallic strains, which can switch the mating type, and therefore ensure the presence of both, *h*^+^ and *h*^-^ cells in the mating mixture, were simply patched on Malt Extract plates (or any other media that induces meiosis), and the *asci* obtained were dissected as before; and 2) heterothallic strains, in which spontaneously occurring mating type revertants (to either *h*^90^ or to the opposite mating type) were isolated (see above), or from which specific *mat1* modified strains from the opposite mating type were produced thanks to the vectors pMTS-h+ or pMTS-h-, as needed (see above). In this case, the mating mixture was made by mixing both strains (original and modified strain), or just patching the *h*^90^ revertant isolated, and plating them in the meiosis inducer media. The obtained *asci* were, as well, dissected using the MSM400 micromanipulator (Singer Instruments, UK).

To help with the tetrad dissection, and due that some of the non-Leupold’s isolates showed a harder cell wall, and therefore were more difficult to dissect, a specific protocol was developed to ensure that dissection could be done in all cases: 50 ml of 10 mg/ml of Zymolyase 100-T (Seikagaku, Japan) were spread in the dissection part of the plate to help cell breakage. After the first round of *asci* dissection (picking asci from the *inoculum* part of the plate) plates were incubated 2-4 hours at 37°C to help cell breakage. This protocol does not affect spore survival as indicated that, with this protocol survivals close to 100% were obtained in several different strains, and data obtained with or without Zymolyase treatment showed no difference (data not shown).

### Statistical analysis

All statistical analyses were performed with the freely available R software (https://www.r-project.org).

Spore survival values were compared using Fisher’s exact test function *fisher.test()* from the *stats* R package. ANOVA test, to detect differences in the spore survival values, were performed by fitting a Generalized Linear Model with a binomial error distribution using the *glm()* function from the *stats* R package. To test the number of strain frequencies showing a statistically significant spore survival value higher than a certain value were performed with a chi-square based test, with the *prop.test()* function from the *stats* R package.

To get an idea of the reproducibility of the values obtained, in the spore survival determination with tetrad analysis using the MSM400 micromanipulator, in some cases, biological replicates were analysed. In those cases, a high reproducibility was observed (see Supplemental Figure S4A), event by using different diploids isolated from crosses coming from different clones of the same original strain (see Table S1). In the cases where no biological replicates were analysed, to get, not just a single value for the spore survival (by summarizing all the different dissection plates carried out), but the spread of the data, every dissection plate was considered as a single measurement of the spore survival value. This was as well done in the cases where biological replicates are available, to compare both ways to show the data (i.e. Compare Figure 1A, Figure 1B, and figure 1C with Supplemental Figures 1A, 1B and 1C). To avoid deviations in the boxplot generation, due to the different number of successfully dissected *asci* in each dissection plates, the R function *wtd.boxplot()*, from the *ENmisc* R package, was used to produce boxplots with weighted values, in each graph where single plates were used as units of repetition.

**Figure 1:**
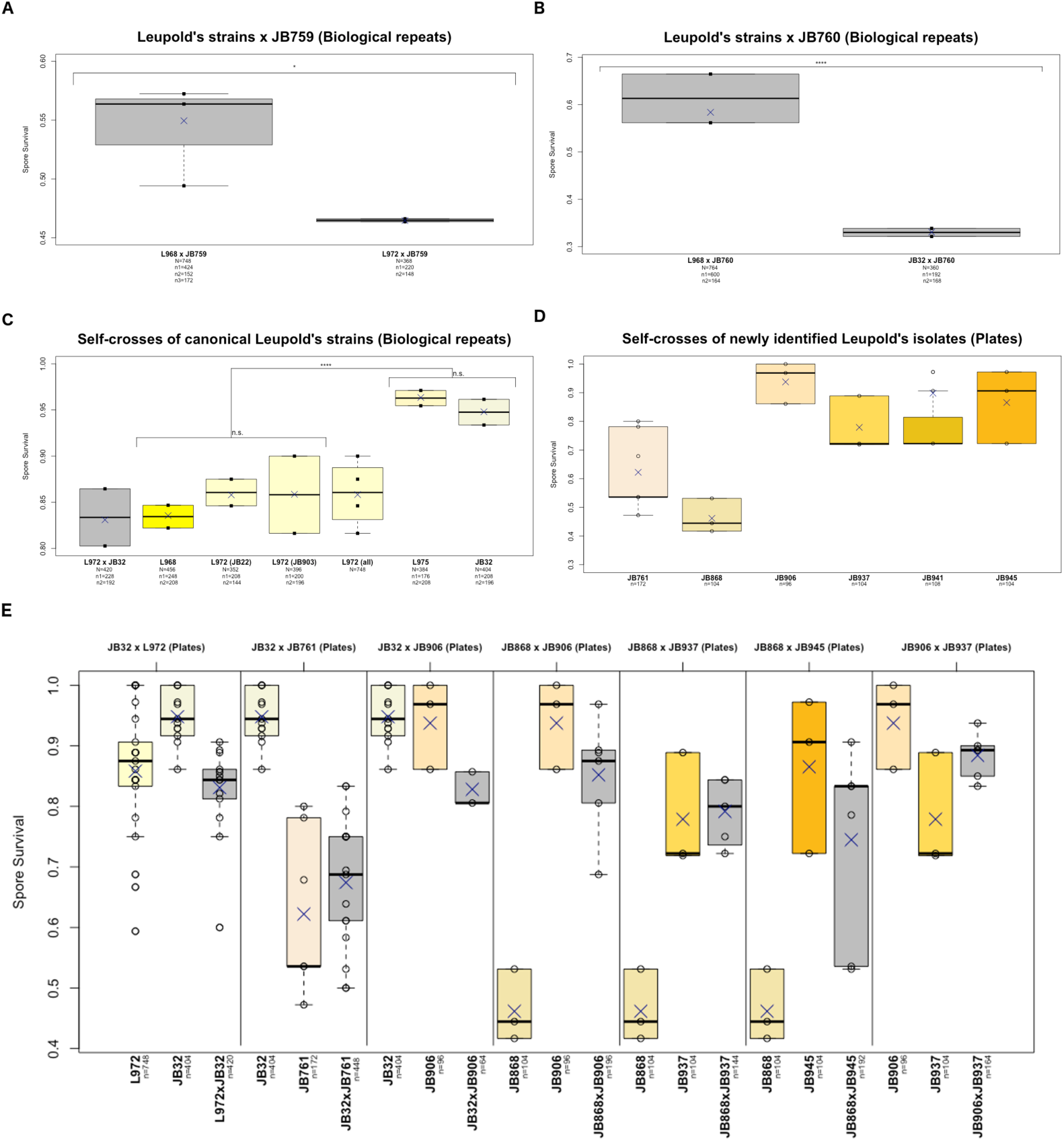
Leupold’s strains clonally related inter-strain crosses and Leupold’s strains self-crosses. Inter-strain crosses **(A)** and **(B)**, of Leupold’s strains L968 (h^90^), L972 (h^-s^) and JB32 (h^+s^), with non-Leopold’s wild isolates JB759 (h^+^) **(A)** or JB760 (h^-^) **(B)**, accordingly. Self-crosses **(C)** and **(D)**, of canonical **(C),** or newly identified **(D)** Leupold’s strains{Jeffares:2015hi}. Two individual diploid isolates (biological replicates) of the respective crosses were analysed by tetrad dissection in **(A)**, **(B)** and **(C). (E)** Inter-strain crosses involving Leupold’s derived strains. Related self-crosses of the performed inter-strain crosses are shown for comparison. In **(A)**, **(B)** and **(C)**, where biological replicates were analysed, the spore survival values obtained in each of the individual isolates are represented with a closed square. In **(D)** and **(E)**, where no biological replicates were analysed, individually dissected plates were used as units of repetition (see Materials and Methods). The spore survival values obtained in each of the individually dissected plates are plotted as open circles. In all panels, the arithmetic mean of all crosses is plotted with a blue X. Boxplots are coloured as indicated: inter-strain crosses boxplots are coloured in grey, self-crosses of individual strains boxplots are coloured with different, but closely related colours, to highlight that different strains correspond to different, but clonally related strains (see Materials and methods). Total number of cells analysed in each individual strain is indicated below each strain description. Numbers of cells analysed in each individual biological replicate are indicate where is relevant. Significance level are plotted as described in Material and Methods (see Materials and Methods, Table S2 and Table S4).

Significance levels, calculated with the Fisher’s exact test function *fisher.test()* or with the chi-square based *prop.test()* R function were transformed, to correct for the multiple testing problem. P-values obtained in each individual comparison were corrected with the Bonferroni transformation, to generate an adjusted p-value in each test (See table S2). Bonferroni correction is a highly conservative correction for the multiple testing problem. This ensures that detected differences are highly confident differences. In this paper’s figures, adjusted p-values are plotted as indicated: n.s., p>0.05; *, p<0.05; **, p<0.01; ***, p<0.001; ****, p<0.0001.

### JB32 SNPs determination, comparison of JB32 SNPs with the *Jeffares et al* 2015 results, and determination of shared SNPs in clonally related strains

SNPs from JB32 were determined as described in Fawcet et al 2014 study^74^, using data obtained from the study Prevorovsky et al 2015^88^. JB32 data was downloaded from ENA (https://www.ebi.ac.uk/ena) under the accession number: E-MTAB-2725. Data from Jeffares et al 2015 study^58^, were obtained from Figshare (https://doi.org/10.6084/m9.figshare.3978303.v1).

To compare the SNPs that segregate in each of the 3 clonal groups considered in the Supplemental Figure S2, a unique vcf file per clonal group, containing all the segregating sites from the considered strains, were manually constructed from downloaded data (see above). Leupold’s clonal strains group were constructed by combining data from two different studies^58,88^, while NCYC132 and JB114-JB1175 clonal groups data were obtained from Jeffares et all 2015 study^58^. Shared SNPs sites between the different clonal groups were determined using the *bcftools isec* function from the BCFtools package^89^.

### Advances Inter-Cross Lines (AILs) and Advanced Self-Cross Lines (ASLs) generation

#### Advanced Inter-Cross Lines (AILs)

To generate Advanced Inter-Cross Lines in fission yeast, a diploid hybrid stock was patched in a double selection plate (see above), and used to inoculate a 100 ml of liquid YES culture that was grown over-night (ON). Next day cells were pelleted, washed in 1ml H_2_O, and resuspended in 100 ml MM-N. Cells were incubated at 25°C with shaking for 2 to 4 days following *asci* formation (azygotic *asci*) with the microscope. Next, cells were pelleted, washed in 1ml H_2_O, resuspended in 1ml 30% EtOH and incubated at room temperature for 10 minutes. This treatment efficiently kills all parental cells^87,90^, with very little effect on spore survival^91^. Treated cells were pelleted, washed with 1ml H_2_O and resuspended in 1ml liquid YES. 0.8 ml of those cells were used to inoculate a new 200 ml YES liquid culture and left ON, or longer if needed, to allow the germination of the F1 spores. The 200-ml grown culture corresponds to the haploid F1 population (AIL cycle 1 population). 2ml from this population was kept as stock in 50% glycerol at -80°C.

For all AIL cycles greater than cycle 1, cells were patched in MM-N plates to induce *asci* formation by random mating them. Mating was done in plates, rather than in liquid MM-N, as done previously, with the diploid starting culture, because from now on all matings analysed will be zygotic matings (the usual mating that normally takes place in haploid fission yeast) rather than the azygotic mating induced from the starting diploid population (see below). To patch the cells in MM-N, 0.5 ml of a 1/10th dilution of the F1 cells were plated and spread in MM-N plates. MM-N plates were incubated at 25°C for 2 to 4 days, monitoring *asci* formation as before. This plate was used both, to continue with the cycling protocol (see later), and as a source of *asci* used to monitor the spore survival by tetrad analysis (first zygotic mating spore survival determination corresponds to AIL cycle 1). To continue from cycle 1 onwards, MM-N patched cells with visible *asci* under the microscope were collected and resuspended in 1ml H_2_O, treated with 30% EtOH as before, washed with 1ml H_2_O and resuspended in 1ml liquid YES. Again 0.8 ml of treated cells were used to inoculate a new 200 ml YES culture than, once grown, corresponds to the haploid F2 population. Repeating this protocol N times allow to increase the number of AIL cycles. In each cycle a -80°C was kept, and in some of the cycles, the spore survival was monitored to test for spore survival changes associated with the AIL cycle progression. For a schematic representation of this protocol see Supplemental Figure 5A, left panel.

The different treatment carried out from the starting diploid cells (sporulated in liquid MM- N), compared with the following cycles (sporulated in MM-N plates), is mainly to avoid diploid mating in the starting diploid sporulation (diploid cells in shaking liquid cultures barely mate). The liquid MM-N applied to diploid cells produce mostly haploid spores coming from diploid azygotic meiosis, rather than diploid spores coming from diploid matings (not shown). On the contrary, to favour cell mating in the following cycles MM-N plates should be used to favour cell proximity needed for the zygotic mating.

It should be noted to the reader that, this author, started developing the AILs and ASLs protocols (see later), in fission yeast, before fully characterizing the issues affecting the differences detected in spore survival values when comparing azygotic *vs* zygotic matings (see below). This invalidated the cycle 0 spore survival determination from these experiments as it was determined in an azygotic meiosis rather than in a zygotic meiosis as all the other AIL cycles. To solve that problem, this author used as the starting spore survival values (cycle 0) in the AILs experiments (and ASLs experiments, see below), previously determined zygotic matings done from the cell stocks used to generate the starting diploid cells. This is shown in the AIL figures (and ASL figures, see below) with a vertical dotted line (to highlight the lack of continuity from the starting zygotic measurement, the cycle 0, and the first AIL zygotic measurement, the AIL cycle 1).

#### Advanced Self-Cross Lines (ASLs)

Advanced self-cross lines were performed originally as a control for the AIL experiment. The procedure to generate the starting diploid cell is the same as in AIL, but in which two differentially *ade6-tagged* clones from the same strain were used (different strains were used in an AIL). All the details explained in the above AIL section apply for the ASL experiments started from a diploid cell. For a schematic representation of this protocol see Supplemental Figure 5A, middle panel.

As explained above, when one starts the ASL (or AIL) experiments from a diploid cell, has an associated problem due to the differences detected in this study between azygotic and zygotic matings (see below). To solve that issue, latest ASL experiments performed by this author were started from a mass mating of haploid cells (which give rise to zygotic matings). For a schematic representation of this protocol see Supplemental Figure 5A, right panel. Those experiments were performed as described in AIL section for the AIL cycles greater than cycle 1 (see above).

In most of the ASLs experiments shown in this work, for statistical purposes, the later cycles were compared with the cycle 0. This is not the case for the strains JB953 and JB873 due to the high spore survival differences detected between cycle 0 and cycle 1. This phenomenon is described all along this report (see bellow). In those cases, cycle 1 data was used as reference start point to detect the effect of the ASL progression.

### JB953 ASL flocculation experiment

The JB953 flocculation phenotype it is simply observed when growing cells in 15 ml Falcon tubes (see Supplemental Figure 5E). To get a more precise visualization of this phenotype flocculation essays on plates were performed: 5 ml tubes were grown ON and poured on thin YES plates and let them incubate at 25°C with 150 rpm ON. Next day photos of the plates were taken as record.

### JB953 ASL fitness increase determination

To perform competitions experiments between original JB953 cells and ASL derived JB953 cells, 6 X 10^6^ cells of each type of cells (differentially *ade6 tagged*) were mixed and used to inoculate a 5 ml miniculture. The singly *ade6 tagged* version of ASL derived JB953 cells were obtained by platting the ASL derived haploid mixture in the needed dominant selectable marker (to select from the population only the ones having this particular marker). Growth of the mixed cells were monitored by optical density (OD) measurements, and new reinoculum were done every next day. This process was followed during 5 days accumulating approximately 20 generations of cell doubling altogether. During all this growing time, the cells freely compete all against all (competition experiment). Proportion of Nat and Hyg cells were determined by plating in triplicate around 200 cells per plate, at the beginning of the experiment, and after the 20 generations were over.

### Spore survival lost on MM-N kept cells

Separated haploid mid-stationary 50 ml YES cultures of the haploid cultures used to perform zygotic crosses were pelleted and resuspended in 50 ml MM-N (the use of heterothallic haploid cells avoids them to enter meiosis completely). At certain given timepoints 100 ml of cells was mixed with the opposite mating type cells and spread into MEA plates to generate the *asci* used to determine the spore survival. In some examples the cells used to determine the spore survival were processed as in the ASL experiment to obtain the F1 cells (ASL cycle 1). Those cells were induced meiosis to determine, as well, the spore survival values upon a self-cross.

### JB873 Freeze and Thaw experiment

JB873 haploid cells, and JB873 diploid cells (XM465 and XM591, respectively; see table S1), were grown to approx. 1 OD (2x10E-07 cells/ml). Tenfold dilutions of those cells were done. The rack of tubes containing those cells were switched repetitively from -80°C to 37°C, keeping them at the different temperatures for 30 minutes each cycle. Samples of all dilutions were plated in YES plates, and colony forming units (CFU/ml) were deduced from the counting of the plates that allow a precise counting of the number of colonies present in the plate. Colonies were manually counted with the help of a touch pressure colony counter.

## 3. Results and discussion

### Reproducible differences, of the spore survival values, are seen in inter-strain crosses involving clonally related fission yeast wild isolates, and in self-crosses, along all the described fission yeast wild population

#### Characterization of the self-cross spore survival values in Leupold’s derived fission yeast strains

The analysis of the spore survival values in inter-strain crosses of fission yeast wild isolates, to determine the degree of reproductive isolation between the different fission yeast wild isolates, that was carried out as part of the *Jeffares et al.* 2015 study, gave an unexpected observation: reproducible differences were observed in inter-strain crosses when different Leupold’s isolated strains were used. Two non-Leupold’s isolates: the strains JB759 and JB760 (showing 1119 SNPs and 2633 SNPs differences to the L972 reference strain, respectively), which are heterothallic isolates (JB759 is an *h*^+^ strain, while JB760 is an *h*^-^ strain), were crossed in both cases with L968 (*h*^90^, and therefore able to cross with either *h*^+^ or *h*^-^ strains) and with L972 (*h*^-s^) or JB32 (a Leupold’s derived *h*^+s^ strain^85^, see Materials and Methods) accordingly (see Table S1 for strain details). Significant differences were apparent when the crosses L968 x JB759 and L972 x JB759 were compared (see Figure 1A and Supplemental Figure 1A). Same was true when the crosses L968 x JB760 and JB32 x JB760 were compared (see Figure 1B and Supplemental Figure 1B). Diploid hybrid strains, obtained from those crosses, were plated on MEA plates, and the spore survival values were determined by tetrad analysis (see Materials and methods). In Both cases, with a high number of individual cells analysed (see Figure 1A and 1B), a low p-value it is observed when comparing the two crosses with Fisher’s exact tests (L968 x JB759 *vs* L972 x JB759: p-value=0.017; L968 x JB760 *vs* JB32 x JB760: p-value=4.68e-15, see Table S2).

Those results are quite striking. L968 and L972 have been fully sequenced and, as mentioned before, they only show 14 SNPs differences between them^58^ (see Table S3). The strain JB32 was not included in the sequencing pipeline applied in the *Jeffares et al. 2015* study, but sequencing data from JB32 is as well available^88^. Only 26 SNPs differences can be detected between L968 and JB32 by full genome sequencing (see Materials and methods and Table S3). This observation, together with the low spore survival values identified in self-crosses, both in the lab^58^ and in the literature^78^, and together with the non-consistent results seen in fission yeast papers regarding Leupold’s derived strains wild type spore survival^66–71^, brought my attention into systematically determine the spore survival values observed in self-crosses from individual Leupold’s derived strains. I obtained two biological independent diploid homozygotes of the following strains: L968, L972, L975 and JB32. Two independently obtained versions of L972 were used: the strain JB22, from Jürg Bähler’s laboratory, and the strain JB903, the L972 version obtained from William Brown’s laboratory^72^. L968 strain can be self-crossed straight away as it is homothallic, and therefore both phenotypically *h*^+^ and *h*^-^ cells are present in the mating mixture. To self-cross the heterothallic strains, I developed two vectors that allow me to switch the mating type of any given strain by a simple transformation protocol (vectors pMTS-h+ and pMTS- h-; see Material and methods for details). Again, although the SNPs differences between all those strains are quite low (see Table S3), highly reproducible phenotypic differences were seen in those self-crosses, as measured by determining the spore survival by tetrad analysis (see Figure 1C and Supplemental Figure 1C). Both L975 and JB32 show consistently self-cross values close to 95%, while L968 or L972 show consistently self-cross values close to 80%. A one-way ANOVA show that, while globally an effect of the strain in shaping spore survival is detected (p-value=3.67e-11), this is not the case when considering either L975 and JB32 (p-value=0.293), or L968 and L972 (p-value=0.338) separately (see Table S4). Polling all the L975 and JB32 data on one side, and the L968 and L972 data on the other side, and testing them for significant differences gives a highly significant p-value (p-value= 5.82e-15; see Figure 1C and Table S2). The inter-cross L972 (*h*^-s^) x JB32 (*h*^+s^) it is not a pure self-cross, but, this cross is added as reference cross, involving an *h*^+^ and an *h*^-^ heterothallic strains, as no background differentiation is considered by the fission yeast community while crossing Leupold’s derived strains.

Moreover, fission yeast population structure characterization carried out by *Jeffares et al. 2015* showed that, out of 161 wild strains with the whole genome fully sequenced, only 57 unique strains were found. Several clonal groups (25) of near-identical strains (<150 SNPs) were described. This was interpreted as the result of single location isolation from mitotically dividing populations, or from inadvertently repeat deposition of the same strains in stock centres. In line with this later interpretation the biggest clonal group identified was the group of clonal strains related to Leupold’s derived strains (used in most fission yeast labs), with 20 different clonal strains identified in it, and the second biggest clonal group is the NCYC132 clonal group of strains, the most used non-Leupold strain in fission yeast labs (see Table S5). I subjected some of those extra Leupold’s strains to tetrad analysis, to determine spore survival upon a self-cross by creating diploid homozygotes of them as before. Only one biological repeat was analysed in those cases and, to get an idea of the spread of the data, individually dissected dissection plates were taken as unit of repetition (see Material and Methods). Phenotypic differences became clearly apparent again (see Figure 1D and Supplemental Figure 1C). This is even more striking, as bigger phenotypic differences are seen (see Supplemental Figure 1C) than in self-crosses done with the canonical Leupold’s derived strains (see Figure 1C and Supplemental Figure 1C). It is remarkable that, with all the self-crosses carried out in Leupold’s derived strains, in 26 out of 55 (47.2%) Fisher’s exact tests comparisons, differences in spore survival values can be detected, with a global p-value <0.05 after Bonferroni correction (see Table S2). A one-way ANOVA with all Leupold’s derived strains as well detects effect of the strain in shaping the spore survival (p-value=0.0255; see Table S4).

The results obtained in all the experiments performed with Leupold’s derived strains in this study might provide a clear answer to the previously mentioned non-consistent results observed in many studies^66–71^. A clear dual behaviour is seen in canonical Leupold’s derived fission yeast strains, showing around 80% spore survival, or around 95% spore survival (see Figure 1C), which could clearly explain the previously mentioned non-consistent results, in which in some studies the wild type spore survival is stated as 80%^69–71^, while in other studies it is stated as 95%^66–68^. Although, this might be simply the case, this study uncovers many more factors affecting the spore survival. It cannot be discarded that, any of those other factors, do as well explain the non-consistent results observed in the literature^66–71^ (see later).

I carried out as well inter-strain crosses involving Leupold derived strains, which again gave significant differences despite the low SNPs differences between the parental strains involved (see Figure 1E, Supplemental Figure 1D, Table S1 and Table S3). The pattern observed in those inter-crosses suggest that multiple factors might be affecting the spore survival of the inter-cross: 1) in some cases dominant or recessive behaviours are observed.

2.1) in the inter-crosses L972 x JB32, JB32 x JB761 and JB906 x JB937, the defects that confers a reduced self-cross spore survival of the strains L972, JB761 and JB937 in respect to their partner’s self-cross spore survival value, behaves as a dominant character in those inter-strain-crosses. Defective behaviour is seen in those inter-crosses, while no defect is seen neither in JB32 nor in JB906 self-crosses; 1.2) on the contrary, a general pattern of recessiveness seems to be attached the defect(s) conferring low spore self-cross survival at the strain JB868, compared with the self-cross survival values of his partners. While in the inter-cross JB868 x JB937 a simple recessive behaviour is seen (the inter-cross show a similar behaviour as the JB937 strain), in the inter-crosses JB868 x JB906 and JB868 x JB945 a behaviour closer to JB906 or JB945 it is observed. In those later cases, there seems to be additional factors affecting the spore survival outcome (see next); 2) in the inter-cross between JB32 and JB906, although neither of the parents show a clear defect in respect to their partner, a defective spore survival is seen in the inter-cross. This most probably indicates that an interaction between JB32 and JB906 factor(s) do affect the spore survival outcome of the inter-cross. Note that the dominant and recessive patterns observed above are mostly not pure (as previously said the behaviour is closer to one strain, but not exactly like it).

In summary, major dominant or recessive defect(s) might be affecting the spore survival of clonally related fission yeast strains with SNPs differences as low as 37 in the bigger case detected (see Table S3). On top of that, additional interacting factors might be affecting the spore survival outcome as well, both in cases where no dominant or recessive defects are seen (i.e. JB32 x JB906), or in cases where a dominant or recessive defect might be as well shaping the spore survival outcome (i.e. JB868 x JB906 or JB868 x JB945). Additional experiment should be carried out to determine if the mayor dominant or recessive defects are due to a defect located in just one locus, or if more complex scenarios are should be taken into consideration (see general discussion for proposed follow-up experiments).

#### Characterization of the self-cross spore survival values in a representative sample of the fission yeast wild strains population

As mentioned above, the population structure of the available fission yeast wild strain isolates had been determined^58^. In that study, the 57 unique (non-clonal) strains identified were grouped in five clusters. I tested the self-cross spore survival values, obtained from diploid homozygotes, in a representative sample of fission yeast wild isolates spanning all the five clusters defined. In most of the cases there are examples of good-performing strains (spore survival >90%) and bad-performing strains (spore survival <90%; see Figure 2A and Table S1). This indicates that the phenomenon of low spore survival upon self-cross is not restricted to a certain group of strains of *S. pombe* with some characteristics, but rather it is observed in genetically very different individuals spanning all the fission yeast wild population.

**Figure 2:**
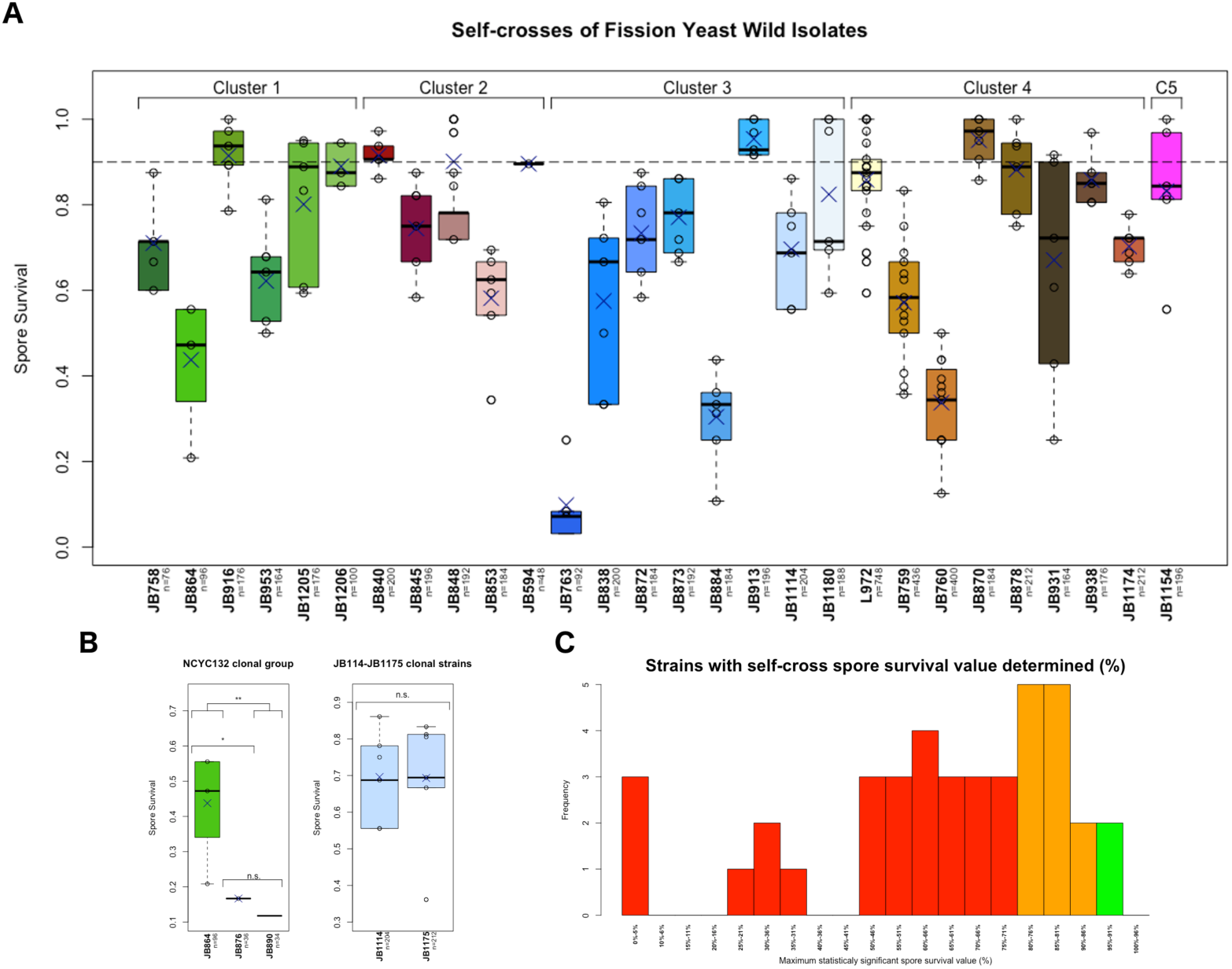
Fission yeast wild isolates self-crosses. **(A)** Self-crosses of a representative sample of the fission yeast wild isolates spanning al the five clusters defined in the Jeffares et al 2015 study{Jeffares:2015hi}. Clusters are indicated at the top of the figure. **(B)** self-crosses of strains belonging to the clonal group including the NCYC132 (JB864) strain (left), or from two clonally related strain JB1114 and JB1175 (right) (see Table S3). Each strain is coloured in a different colour. Clustered and clonally related strains boxplots are plotted in related colours (see Materials and Methods). Figure details are indicated as in Figure 1. **(C)** Strain frequencies showing the number of strains with a statistically significant spore survival value. Chi-square test to detect spore survival values higher than a certain value (from 0.95 to 0.05 in 0.05 intervals, plus 0.01) were computed for each self-cross tested strain. Strains have been binned in 5% spore survival intervals according to the maximum spore survival value detected with the available data. Histogram representing strains with spore survival significantly higher than 90% are plotted in green; strains significantly higher than 75% in are plotted in orange, while the rest of the histograms are plotted in red.

The phenotypic variability, seen between clonally related Leupold’s isolates, it is, as well, a phenomenon not only restricted to Leupold’s strains. I tested some isolates from two other clonal group of strains: the strains from the NCYC132 (JB864) clonal group of strains, and the JB1114-JB1175 clonal strains (see Table S1 and Table S5). While no phenotypic differences, in spore survival, where detected between the clonal strains JB1114 and JB1175 (see Figure 2B right panel, and Table S2), a clear difference where detected in the NCYC132 group of strains, both in self-crosses (see Figure 2B left panel, and Table S2), and in inter-strain crosses involving clonally related NCYC132 isolated strains (see Supplemental Figure 2A, Table S4 and Table S5). Although the SNPs differences between the L972 reference strain and the strains NCYC132 is quite high (66843 SNPs differences have been identified by whole genome sequencing^58^), again, a small number of SNPs differences between the clonally related strains (see Table S3) is giving huge phenotypic differences in the spore survival values both in self-crosses and in clonally related inter-strain crosses. The JB1114-JB1175 clonal group of strains also exhibit huge differences in full genome comparisons (37884 SNPs differences have been identified between the JB1114 and the L972 strain and 39690 SNPs differences have been identified between the JB1114 and the NCYC132 strain^58^).

Could it be that a common group of hyper-mutable SNPs could be the cause of the observed spore survival variability in all the different clonal groups analysed despite the overall differences between them? The analysis of the polymorphic sites in those clonal groups argues against this hypothesis (see Materials and Methods). Only four polymorphic sites are shared between the L972 clonal group and the NCYC132 clonal group, and only two polymorphic sites are shared between the NCYC132 clonal group and the JB1114 clonal group (see Supplemental Figure 2B and Table S6). This is a very small number of shared polymorphic sites and with no clear evidence of any cause-effect relationship between the shared sites and the phenotypic variability. Two main observations back this later statement: first, in the JB1114 clonal group of strains no polymorphic site seems to cause any phenotypic effect, as no significant differences exist between JB1114 and JB1175 spore survival values (see Figure 2C, Table S2 and Table S6). Giving this, no effect of this two SNPs it is expected to be the cause of the phenotypic variability in the spore survival of the NCYC132 clonal group of strains (where the same sites are polymorphic); and second, the 4 polymorphic sites shared between the L972 and the NCYC132 clonal group of strains cluster altogether in the *mal1* gene (see Table S6), with no known function in regulating the spore survival. The *mal1* gene has been related to maltose metabolism^92^, this might simply reflect that in both groups some strains have adapted differently to environments containing (or not) maltose as main sugar source.

Globally, from all self-crosses carried out in this study, only in 14 out of 40 strains analysed (35%), the spore survival value is significantly higher than 75%, and only in 2 of those strains (L975 and JB32) the data collected support spore survival values higher than 90% (see Figure 2C). The clear majority of the strains analysed (65%), have reduced spore survival values (<75%). Bigger sample sizes, and more biological replicates are needed to fine tune this dataset, but this clearly indicates that, factors that affect the outcome of the self-cross spore survival values, are pervasive in fission yeast wild populations.

### Spore survival determination showed significant differences when comparing azygotyic meiosis with zygotic meiosis in a high proportion of the strains analysed

A further surprise came when analysing spore survival data of self-crosses from fission yeast wild isolates: spore survival data obtained by mass mating experiments, rather than by selecting diploids cells (see Material and Methods for details) was significantly different in 9 out of 22 (40.9%) strains where spore survival values was determined using both methodologies (see Figure 3A and Table S2). Fission yeast is a naturally haplontic free living organism that upon poor environmental conditions (lack of nutrients) mate and immediately enters meiosis to produce spores that are released into the environment^87^. This process, where meiosis takes place just immediately after mating, is known as the zygotic meiosis. Sometimes, the diploid zygote can develop into diploid cells that, when entering meiosis, produce an azygotic ascus (azygotic meiosis)^87^. The spore survival data showed in previous figures of this report were obtained all from the analysis of azygotic meiosis, as diploid cells, either hybrids for an inter-strain cross, or diploid homozygotes, for a self-cross, were generated (see Material and Methods). On top of all the azygotic self-crosses carried out in this study, I decided to test, as well, some zygotic meiosis self-crosses, to test for the robustness of the spore survival data obtained. This analysis produced the surprising result shown in the Figure 3A of a high number of examples (40.9%) where the spore survival data obtained from a zygotic meiosis is significantly different of the spore survival obtained from an azygotic meiosis. In this report, from now on, if no specific text is shown previously, all self-crossed from azygotic meiosis, will be named in the main text, or in figures, with the strain name (i.e. L972), while all self-crossed from zygotic meiosis, will be named with the strain name followed by an M suffix (i.e. L972M).

**Figure 3:**
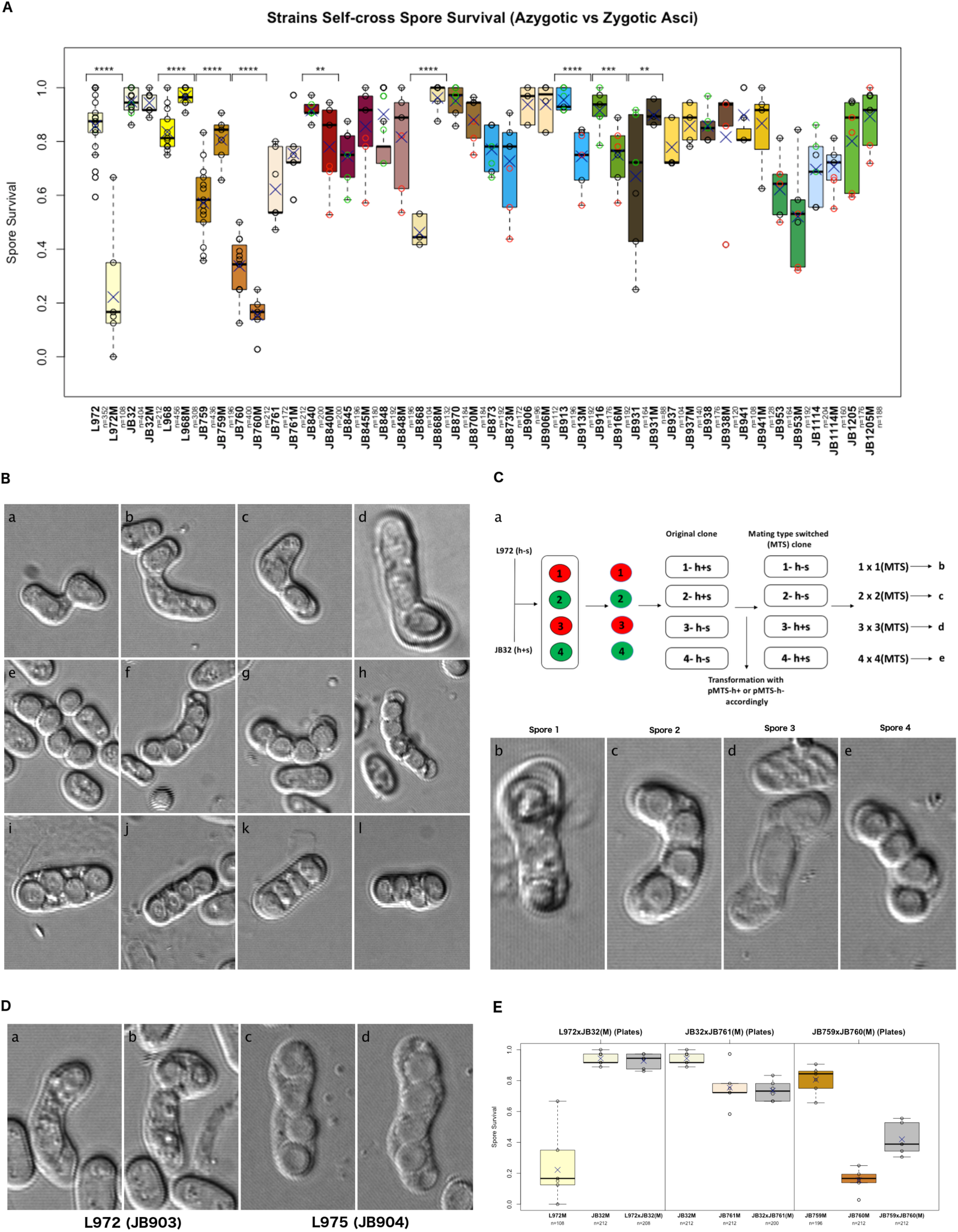
Azygotic vs zygotic asci spore survival determination. **(A)** Comparison of the spore survival determinations carried out by tetrad analysis either in diploid cells (azygotic asci) or in haploid cells by mass mating experiments (zygotic asci). Twenty-two strains were tested in both experimental conditions (zygotic asci results are suffixed with "M"). Statistically significant differences are shown (see Material and Methods and Table S2). A One-Way ANOVA of this data is available in Table S4 **(B)** Images of asci obtained in L972 self-cross experiments: panels a, b, c and d correspond to L972 zygotic self-cross asci; panels e, f, g and h correspond to asci coming from an L972 (h^-s^ naturally occurring version) inter-strain cross mass mated (zygotic mating) with JB32 Leupold related strain (h^+s^ strain made on purpose by Heim{Heim:1990vx}; panels i, j, k and l correspond to L972 azygotic self-cross asci. **(C)** Panel a: schematic experimental protocol carried out to determine that L972 zygotic mating defect it is caused by a single locus (as 2:2 segregation is seen in descendants from L972 x JB32 inter-strain cross). Spores are coloured red (bad zygotic self-cross performance) or green (good zygotic self-cross performance) and representative asci from each cross are shown in panels b, c, d and e. MTS: Mating type switched strain. **(D)** The same defect detected in Jurg Bähler’s lab version of L972 (JB22) zygotic self-cross is as well detected in William Brown’s lab L972 independently maintained stock (JB903){Brown:2011dh}: panels a and b. Strain L975 doesn’t show the zygotic mating defect: panels c and d. **(E)** Zygotic inter-strain crosses performed in this study. Figure details for boxplots are as indicated in Figure 1. In **(A)** most individually dissected plates (circles) are coloured black as in Figure 1, but some plates are coloured green or red. See main text for details. Boxplots are coloured as in previous figures.

Among the examples that showed a differential behaviour, it is especially relevant the case of L972, the fission yeast reference strain, where spore survival upon a self-cross in an azygotic meiosis gave values of 85,83%, while in a zygotic meiosis gave survival values of 22,22%, and with most of the mating figures showing aberrant meiotic shapes (see Figure 3B panels a, b, c and d). This aberrant mating figures are not seen neither in an L972 mating with another Leupold related wild type, as the JB32 strain (see Figure 3B panels e, f, g and h), nor in a L972 azygotyic self-cross (see Figure 3B panels i, j, k and l). The mating defect it is only appreciated in a zygotic L972 self-cross. Moreover, this defect is caused by just one locus, as deduced from individually phenotyping the aberrant mating figures in the four spores coming from a cross between L972 and JB32 (see Figure 3C). On top of that, this defect seems to be quite unstable. Initially, this defect, seemed to be an L972M specific defect, as it is shown by two independently maintained L972 stock clones: JB22 from Jürg Bähler’s laboratory (see Figure 3B panels a, b, c and d), and JB903 from the William Brown laboratory^72^ (see Figure 3D panels a and b). Going back to an earlier version of the source where JB22 was obtained made the defect to disappear (Jürg Bähler’s personal communication). On top of that, further data obtained in this study (see later in text and in Figure 5C, left panel), the JB22 defect seems to be eliminated in meiotic cells. The determination of the Mendelian inheritance of this defect could foster the explanation that a genetic mutation has accumulated in the defective clones, but the fact that two independently maintained L972 clones showed the defect, together with Jürg Bähler’s observation, and with the meiotic elimination of this defect, observed in this study, go against this claim. Within Leupold’s derived strains, this defect is L972M specific, as neither L975M (see Figure 3D panels c and d), nor any other Leupold derived strain show this defect in a mass mating (zygotic meiosis) spore survival determination (see Figure 3A).

I performed some inter-strain crosses as before, but using zygotic matings instead of azygotic matings as used above. Those crosses involve both, clonally related strains: L972 x JB32 and JB32 x JB761 crosses, or not clonally, but closely related strains: JB759 x JB760 cross (see Figure 3E and Table S5). In those examples, dominant or recessive behaviours are as well observed: 1) as detailed before, in a zygotic inter-strain cross between L972 and JB32, the L972M defect behaves as a recessive defect. No defect it is observed in a zygotic cross between the strains L972 and JB32, neither in the analysis of the spore survival values (see Figure 3E left panel), nor in the aberrant meiotic shapes (see Figure 3B, panels e, f, g and h); 2) a general pattern of dominance is instead observed when comparing the defect that JB761M and JB760M show compared with JB32M and JB759M respectively. In both those two zygotic inter-strain crosses, the hybrid behaves as the more defective parental partner (see Figure 1E, middle and right panels). In the not clonally related example (JB759 x JB760), not a unique relation of dominance it is observed when comparing JB759M and JB760M with the zygotic self-cross of the strains JB759 and JB760. This is what it should be expected if additional factors are as well shaping the spore survival of the hybrid (expected due the fact that they show a higher genomic divergence than clonally related strains). Strikingly, the JB759-JB760 hybrid strain has a higher zygotic spore survival self-cross value (see Figure 3E right panel). This might indicate that, additional factors might, as well as lower the spore survival value (as seen in Figure 1E example), increase it.

It is, as well noticeable that, while in the cross L972 x JB32, the azygotic L972 defect behaves as a dominant characteristic (see Figure 1E and above in the text), in the same cross but zygotic, the L972M defect behaves as a recessive character (see Figure 3B and Supplemental Figure 3A for a detailed comparison). This observation is, again, really striking. L972 and JB32 only show 29 SNPs differences on a whole genome sequencing (see Table S3). Could it be that, just 29 SNPs differences could explain functional differences in spore survival that, behave differently in zygotic and azygotic meiosis? This might imply that, between those 29 SNPs differences, at least one affects the spore survival only in azygotic meiosis, and that, the defect generated is dominant in an inter-strain cross, and that, another SNP difference, has an effect only in zygotic meiosis and is recessive in an inter-strain cross. Alternatively, the same SNP defect might behave differently in a zygotic or in an azygotic meiosis. In my opinion, both scenarios are quite unlikely. Further results shown in this report do as well make those scenarios unlikely (see bellow). On the contrary, the defect shown by JB761 when compared with the JB32 strain it behaves as a dominant character, both in azygotic and zygotic examples (see Figure 1E, 3E and Supplemental Figure 3A for a detailed comparison). See bellow in the text, and in the general discussion for a more detailed explanation about the JB761 spore survival defect (see below and General discussion).

A further, unexpected, surprise came from the analysis of this dataset. The initial purpose of this author, when performing the comparisons between azygotic *asci* and zygotic *asci*, was to ensure to have high numbers of analysed cells in all the strains analysed. For such a purpose, a target of dissecting six full dissection plates per cross was set (if all the tetrads were successfully dissected this give rise to 216 cells analysed in each condition, when using the default setup of the Singer MSM400 tetrad dissector). To speed up the global view of this phenomenon, I decided to do two batches of three plates in each of the strains considered. The first batch of plates were analysed straight away from freshly patched cells, obtained from the -80°C stocks. The second batch of plates was analysed from the same cells, once the dissection of the first three plates was finished. Patched cells were kept at 4°C in-between. This decision, rather than being anecdotic, turn out to be highly suppressive. In a highly reproducible manner, the first set of three plates analysed always show a higher spore survival value than the later set of three plates analysed. This is detailed in Figure 3A, where, in the cases analysed in this way, the first set of three plates is shown with black circles (the standard way of showing plates in the figures; see caption in Figure 1), while the set of the last three plates are shown in red coloured circles (see Figure 3A). No batch effect was initially expected by this author. Consequently, in most cases, no batch information was properly recorded. Separated from this set of 4°C kept plates, in another set of plates, batch information was, as well, recorded. In this case, no 4°C exposure occurred in-between the two determinations. This is as well detailed in Figure 3A. The set of three plates analysed in the later stage are shown in green coloured circles. A more detailed analysis of this phenomena is described further in the following section of this manuscript (see below).

Given this unexpected surprise, the prior statement where it is claimed that, in nearly 41% of the cases, the spore survival values of a self-cross are significantly different between azygotic and zygotic meiosis, may be wrong due to this unexpected effect. To check for this, a further analysis was performed, in which the values of the last three plates obtained in the 4°C kept plates (red circles in Figure 3A) were eliminated from the analysis. In this analysis, no big differences could be observed (see Supplemental Figure 3B and Table S2). In the strains JB840 and JB845, there seems to be an effect of eliminating the 4°C data (JB840 difference might be wrong, while JB845 was undetected due the 4°C effect). Nevertheless, clear differences between some zygotic and some azygotic meiosis seem to exist. Several observations support this claim, although, the data is affected by unexpected and uncontrolled experimental conditions: 1) robust differences are detected in the following strain: L972, L968, JB759, JB760, JB868, JB913, JB916 and JB931; 2) representative examples show consistent effects: 2.1) L972M has an effect that it is not only observed by spore survival values, but implies, as well, abnormal mating figures (see Figure 3B). The defect detected in L972M is recessive (only seen in a self-cross), and highly specific to L972M (it is not seen in L968, or L975, clonally related strains, neither in zygotic nor in azygotic matings); 2.2) L968, although clonally related to L972, doesn’t show any mass mating defect, but instead show a defect in the opposite direction (azygotic meiosis showed lower survival than zygotic meiosis; see Figure 3A); 2.3) JB868 is a newly identified *h*^90^ Leupold derived strain. It is probably an erroneous restocking of L968 strain^58^. Consistently with this interpretation, JB868M and L968M show the same behaviour, but JB868 seemed to have lost spore survival viability specifically in the azygotic meiosis, but no spore survival lost is seen when comparing L968M and JB868M spore survival in a zygotic meiosis (see Supplemental Figure 3C and Table S2); 2.4) as in azygotic meiosis (see above), clonally related spore survival zygotic meiosis differences, can be detected. The clonal group of strains that include the strains JB760, JB888 and JB889 (see Table S5) do show highly significant phenotypic variation in zygotic spore survival determinations (see Supplemental Figure 3D and Table S2), although, almost no SNPs differences are detected between those strains by full genome sequencing (see Table S3).

To specifically test for which factors do affect the spore survival values some ANOVA tests were carried out. A one-way ANOVA test with all the data collected (see Figure 3A) doesn’t support a clear involvement of the type of meiosis (azygotic or zygotic) in the determination of the spore survival values (p-value = 0.998; see Table S4 and Figure 3A). This might simply indicate that most of the strains analysed doesn’t show this effect, or that some other uncontrolled factors are masking this effect. Indeed, this seems to be the case as, if the same test is carried out, but without the data affected by the exposure to 4°C, a clear significant effect of the type of meiosis in the spore survival values is detected (p-value=0.00489; see Table S4 and Supplemental Figure 3B). The unexpected effect of the 4°C treatment seems to be masking the type of meiosis effect in the spore survival. Given that there is a distortion in the acquisition of the data, due to unexpected batch effects, a multifactorial ANOVA test (MANOVA) was carried out, just in the strains that include batch information. This analysis showed a clear effect of the 4°C affected batches (p-value=5.21e-19; see Table S4 and Supplemental Figure 3E), while no effect is detected for the other batches in shaping the spore survival (p-value=0.5029; see Table S4 and Supplemental Figure 3E). This MANOVA analysis as well detects an almost marginal significant effect of the type of meiosis in shaping spore survival data (p-value=0.0511; see Table S4 and Supplemental Figure 3E), which, might not be as clearly significant, as discussed earlier, simply by the fact that not all the strains analysed show this effect. Moreover, a one-way ANOVA test with the same set of strains detects an effect of the type of meiosis in the spore survival determination (p-value=0.00489; see Table S4 and Supplemental Figure 3E).

In summary, although the 4°C batch effect might have influenced some spore survival determinations, overall no elimination of the zygotic *vs* azygotic differences is detected upon elimination of the 4°C affected plates (see Figure 3A, Supplemental Figure 3B, and Supplemental Figure 3E). Altogether the data presented in this report supports for the existence, in some strains, of meiotic defects that are specific to zygotic or azygotic meiosis, or that show up differentially in both types of meiosis. The unexpected batch effects (not foreseen), requires a comprehensive study to detect and characterize those differences more accurately. In further studies, a special attention should be taken in designing the data acquisition to avoid possible experimental artifacts.

### Environmental induced variation of the spore survival values: the 4°C and -80°C examples

As mentioned before, the comparison carried out between zygotic and azygotic meiosis, accidentally gave an, unexpected, and extremely interesting observation: the spore survival values seen in fission yeast self-crosses, rather than being a precise and fix value that is a given characteristic of every particular strain, it is a changing value that it is affected by environmental factors: either the kind of meiosis in which one strain is going through, or, more importantly, simply by being exposed for a certain period of time to a harsh environmental condition, like being kept at 4°C for a certain amount of time. On the other side, most of the spore survival values determined using biological replicates obtained during this report, gave an extremely precise and reproducible spore survival values (see Supplemental Figure 4A). Moreover, an ANOVA test, couldn’t detect any effect of the replicates in shaping the spore survival values (p-value=0.232; see Table S4 and Supplemental Figure 4A). It should be noted that, the supplemental figure 4A only include azygotic spore survival data. Later in this manuscript, some zygotic biological replicates have been obtained due some extra experiments carried out. This aspect will be discussed in higher detail later in the manuscript (see below and General discussion).

The fact that, in azygotic biological replicates, even when working with diploid biological replicates obtained from different starting haploid clones (see Table S1), we can see highly reproducible values, and that in some cases, this value is close to 100% survival, argues in favour of the fact that, the variability that it is observed, is poorly affected by technical factors, but mainly reflecting some biological variability that might exist between the strains. On top of that, no difference was observed between the data acquired by this author, and the data acquired by master students who helped in some of the data acquisition (data not shown). This later observation, reinforces the above-mentioned interpretation that technical factors are almost negligible in the spore survival value determination by means of tetrad analysis.

The characterization of the effect on the spore survival by the exposure of the cells at 4°C shows that this effect is clearly majoritarian, as in most of the strains (7 out of 13; 53,8%), that where accidentally analysed in this way, statistically significant differences can be found when comparing the initial batch with the 4°C treated batch (see Figure 4A and Table S2). This effect seems to be specific to this environmental treatment, as other examples, where batch information was recorded, but in which no 4°C exposure in between happened, doesn’t clearly shown the effect (see Supplemental Figure 4B). In those strains, only in 1 out of 12 strains (8,3%), a significant effect could be detected (see Table S2). On top of that, in this significant example, an increased spore survival value in the second batch it is observed (see Supplemental Figure 4B, JB931 rows), while in the 4°C batch effect data, in all the significant examples, the spore survival always change from a higher value to a lower value. Moreover, in the 4°C batch effect data, even in the non-significant examples, this tendency is always observed, suggesting that, with a higher sample sizes, or with a stronger environmental insult, a significant change might be as well observed (see Figure 4A).

**Figure 4:**
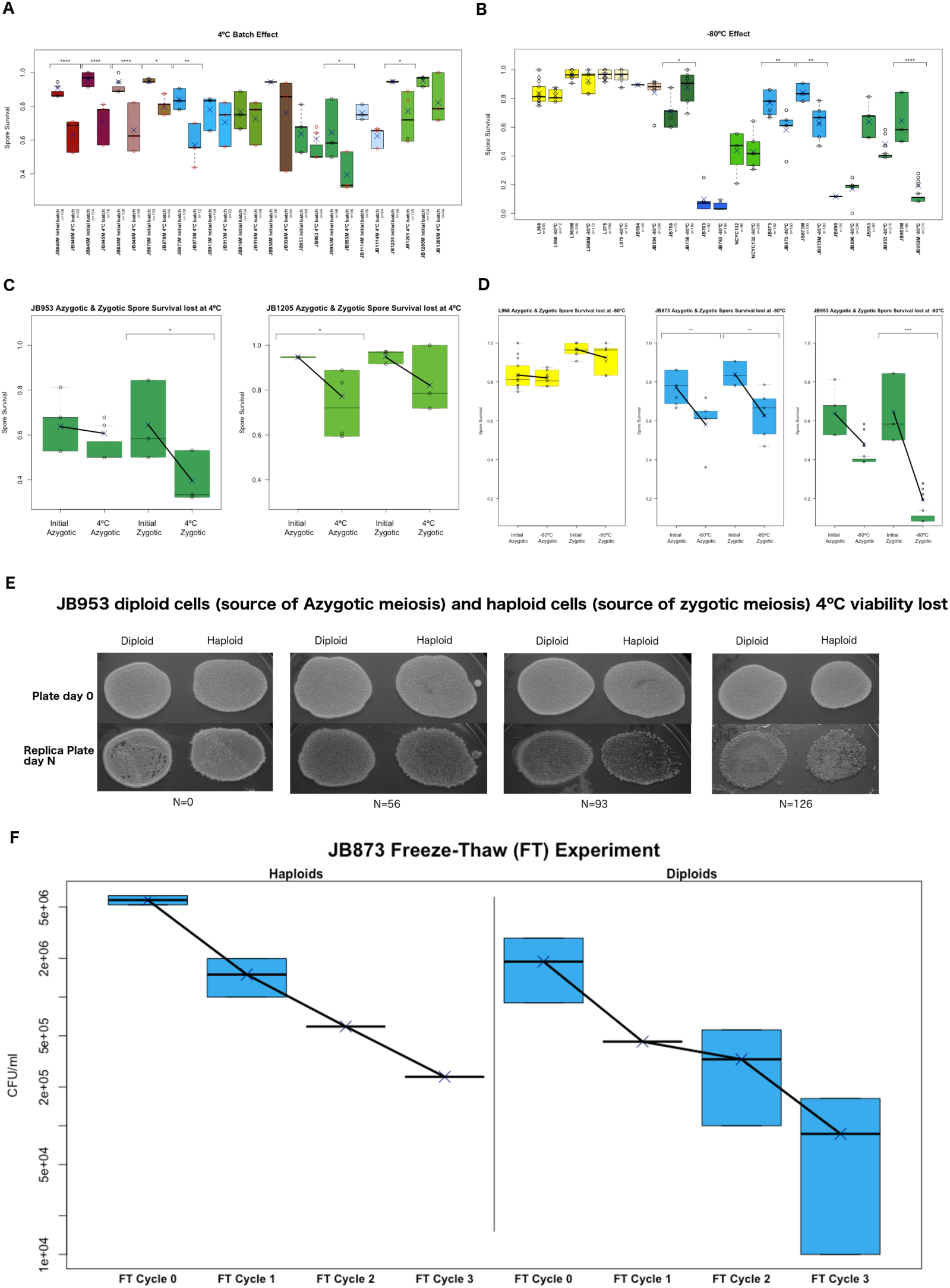
The 4°C and -80°C environmental effect on fission yeast self-cross values. **(A)** Reduction of the fission yeast self-cross spore survival values upon exposure of the vegetative cells to an uncontrolled period at 4°C inside a laboratory fridge used routinely to kept yeast plate stocks (from 1 to 4 month approx.). **(B)** Variation of the fission yeast self-cross spore survival values observed upon reanalysis of a different patch of cells obtained from the same stock tube, after the stock tube has been kept for a long period of time (longer than 6 months) in a -80°C laboratory refrigerator. This is the usual way of keeping yeast stocks. **(C)** and **(D)** Detailed view of the data shown in panels **(A)** and **(B)** for the strains JB953 and JB1205 treated at 4°C **(C)** or for the strains L968, JB873 and JB953 treated at -80°C **(D)**. Those examples correspond to the ones in which the zygotic and azygotic meiosis spore survival values were determined. **(E)** JB953 vegetative cells patched in rich media original plates (YES plates; top panels) and replica plates done in YES after N days at 4°C in a laboratory fridge (bottom panels). Number of days in which cells were maintained at 4°C are shown below. **(F)** Cell viability loss of the haploid (left panel) and diploid (right panel) JB873 cells subjected to a freeze a thaw experiment (see Material and methods). Boxplots are coloured as in previous figures. Significance level are plotted as described in Material and Methods (see Materials and Methods, Table S2 and Table S4).

**Figure 5:**
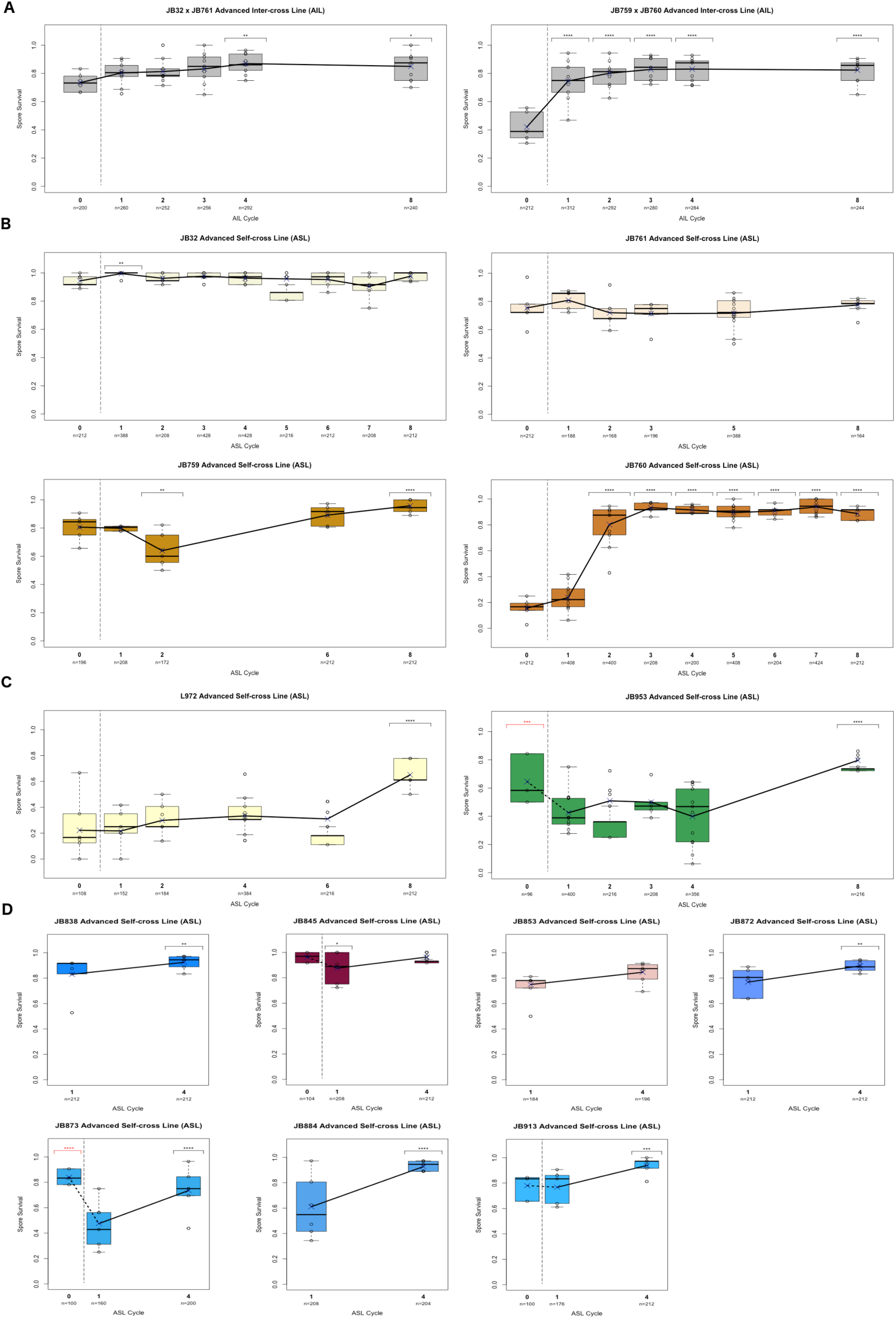
Advanced Inter-Cross Lines (AILs) and Advanced Self-Cross Lines (ASLs). **(A)** AILs of two selected fission yeast wild isolates inter-crosses showing initial spore survival values with high zygotic spore survival self-cross (JB32 x JB761=73.5%; JB759 x JB760=41.98%). **(B)** Individual ASLs of the four strains involved in the AILs shown in **(A)**. **(C)** Two additional ASLs in which the spore survival change was analysed up to the 8th ASL cycle. **(D)** Seven additional ASLs in which the spore survival change was analysed up to the 4th ASL cycle. Vertical dotted line between the cycle 0 and the cycle 1 indicate the lack of continuity between the two spore survivals measured, due that most of the AILs and ASLs were started from a single diploid clone (either a diploid hybrid in AILs or a diploid homozygote in ASLs). For details see Materials and methods and Supplemental Figure 5A. ASL cycle 0 and ASL cycle 1 are joined with a straight line when the exact same cells were used in cycle 0 zygotic spore survival determination than the cells used to produce the diploid cells used to start the ASL experiment. In the h^90^examples though, only one of the two cells different cells used to generate the diploid cells were used in the cycle 0 zygotic spore survival determination. This is shown with a dotted line joining the ASL cycle 0 with the ASL cycle 1. Figure details are as indicated in Figure 1. For statistical purposes in most cases the cycle 0 value is used as a reference value for comparison. This is not true in the strains JB953 and JB873 where ASL cycle 1 was used as a reference, this fact is indicated with red coloured statistically significant differences detected in those strains cycle 0. For details see Material and methods and Figure 4.

In summary, there seems to be an effect of the 4°C exposure, that has a clear effect of reduction of the spore survival values in a self-cross. In other words, the 4°C effect has a clear directionality effect that leads to a spore survival reduction. On the contrary, no general pattern can be detected in the other batch data available (see Supplemental Figure 4B). As discussed earlier, a MANOVA test, with only the strains where batch information was recorded, do as well support the claim that only the 4°C treatment it is significantly affecting the self-cross spore survival values (see Supplemental Figure 3E). Altogether this data argues in favour of, that, during the 4°C exposure, there are some biological changes, in most of the fission yeast strains, if not in all of them, that affect the spore survival value observed, by reducing it, when the strains are self-crossed.

A similar effect, showing a reduction of the spore survival value in a self-cross, was as well observed in cells exposed to -80°C. In some strains, when reanalysing the spore survival of a self-cross, from a new patch of cells obtained from the same stock tube, but after the stock tube has been kept for a long period of time (longer than 6 months) at -80°C, a reduction of the self-cross spore survival was observed. This is the usual way of keeping yeast stocks. Interestingly, in the Leupold’s derived strains analysed (L968 and L975) only a tiny, non-significant reduction is observed (see Figures 4B and 4D, L968, L968M, and L975 rows), while in some other strains, like JB873 and JB953, a clear effect is observed in -80°C treated cells (see Figures 4B and 4D, JB873, JB873M and JB953M rows).

The first observation of this phenomena, the 4°C effect, were done without carefully planning experiments to characterize it (as unforeseen). The unforeseen nature of those observations, lead to the fact that most meiosis, where this effect was observed, were zygotic meiosis (see Figure 4A). Fortunately, some azygotic meiosis, in some of those strains, were as well analysed in the same experiment (see Figure 4A). This fact, lead to another, unexpected, and extremely interesting observation: different strains show different patterns in the 4°C or -80°C mediated self-cross spore survival loss, depending on which kind of meiosis was analysed (zygotic or azygotic). While, in the strain JB953, a higher spore survival loss is observed in a zygotic meiosis, compared with the spore survival loss observed in the azygotic meiosis (see Figure 4C, left panel), this is not true for the JB1205 (see Figure 4C, right panel). The strain JB1205 show a similar ratio of spore survival loss in both types of meiosis (although with the current data, in zygotic meiosis this claim is not statistically significant, a clear pattern it is observed, suggesting that, with a higher sample size, the spore survival might be, indeed, significantly reduced). The same differential behaviour was as well observed in the -80°C effect. In this case, again, the JB953 strain show the same differential behaviour, as seen in the 4°C example, between zygotic and azygotic meiosis: the self-cross spore survival loss is much more pronounced in the zygotic meiosis than in the azygotic meiosis (see Figure 4D, right panel). Fortunately, again a differentially behaving strain was identified: the JB873 strain shows a clear self-cross spore survival reduction, both in zygotic and in azygotic meiosis (see Figure 4D, middle panel). In the strain JB873 exposed at -80°C, the same ratio of spore survival loss it is observed in a zygotic and in an azygotic meiosis, in a similar way to what it is observed in JB1205 when exposed at 4°C (see Figure 4C, right panel). The Leupold’s strain L968 show a slight pattern of spore survival reduction, which might indicate, that, the same phenomena might be happening but at much, much lower rate in all cells (see Figure 4D, left panel and later in General discussion). In this case, as well, the same ratio of reduction, might be happening when comparing zygotic and azygotic meiosis (see Figure 4D, left panel), although, due to the tiny effect observed, this cannot precisely be determined.

Another observation to highlight, is the fact that, when comparable examples are available, the same behaviour it is observed between the 4°C effect and the -80°C effect: in the strain JB953, the self-cross spore survival value was analysed for the two different types of meiosis, both in the 4°C effect and in the -80°C effect. In both cases, a higher spore survival reduction is seen in zygotic meiosis than in azygotic meiosis (see Figure 4C, left panel; and Figure 4D, right panel). In the strain JB873 the zygotic mating was analysed in both 4°C effect and -80°C effect (see Figure 4A and 4B). In this example, again, the behaviour observed when the cells are exposed to 4°C, is the same as the behaviour of the cells that are exposed to -80°C. Those observations suggest that, the same phenomena that it is affecting the biology of some fission yeast strains at 4°C, might be as well affecting them while they are being kept at -80°C as stocks. More examples are needed to clearly conclude that this, indeed, is a general rule (see General discussion).

Another, even more important, unexpected, and unforeseen observation was done in this experiment: the viability of the JB953 vegetative cells (previously to any mating or meiosis induction), was showing the same differential behaviour when comparing diploid cells (the cells patched in MEA plates to induce an azygotic meiosis), with haploid cells (the cells patched in MEA plates to obtain a zygotic mating), when both were kept for a while at 4°C. This was detected with a replica plate of the same cells used for the spore survival determination showed above. The original observation, as unforeseen, and, as this author didn’t had done yet the spore survival analysis, was not recorded. I could, however, successfully reproduce the same experimental observation in a newly started experiment (see Figure 4E). In this qualitative experiment, the diploid cells seemed to die at lower rates than haploid cells. This observation raises a fundamental question: could it be that, what is observed with the spore survival lost in fission yeast self-crosses, is, in fact, reflecting an underlying biological phenomenon that it is happening to all vegetative cells, regardless of the fact that they will later be induced for their entry into meiosis? As discussed later in the general discussion, precise quantitative experiments should be done to answer this question (see General discussion).

Regardless of that, I have been able to do a quick and dirty quantitative test with the JB873 strain: I subjected JB873 haploid and diploid cells to a quick -80°C environmental stress, by quickly changing them, for short periods of time, from -80°C to 37°C in repetitive cycles (freeze and thaw experiment, see Materials and Methods). In these experiments, it is clearly observed that, as seen in spore survival experiments (see Figure 4D, middle panel), the haploid and diploid vegetative cells, stressed by exposing them to a -80°C related stress, respond with a similar ratio of cell viability loss in haploids and diploids cells (see Figure 4F). This corroborates, in a semi-quantitative way, the qualitative observation done with JB953 (see Figure 4E): the loss of cell viability, seen in vegetative cells, seems to behave in the same way as the spore survival loss uncovered in this manuscript. As discussed later in the general discussion, if this is the rule rather than the exception, this might indicate that, there is a correlation between the spore survival recovery, and the vegetative cell survival recovery. This would implicate that, what is seen with the spore survival loss is, in fact, reflecting an underlying biological phenomenon that happens to all cells and affects its vegetative survival viability (see General discussion).

### A meiosis repair program corrects the spore survival defects previously shown in this report, as well as corrects some more independent defective phenotypes

As explained above, the phenomena detected by analysing fission yeast self-cross values, and the signs that, this not only affects the spore survival, but the vegetative cell survival as well, suggest an underlying biological phenomenon happening in all fission yeast cells. The data acquired so far, however, gives no direct evidence on what could this phenomenon be, and, ultimately, what it is the cause of all this huge phenotypic variability observed even when comparing clonally related fission yeast strains. An obvious suspected cause of all this phenotypic variability would be the existence of a pool of DNA mutations that might explain all these observations. Indeed, in a recent paper, the strain JB759 has been shown to be a *swc5* frameshift mutation^73^. Could it be that, this scenario is widespread in fission yeast wild isolates, and that this pool of DNA mutations explains the huge phenotypic variability seen in fission yeast self-crosses spore survival determination? Although, this might certainly be an option, it would not easily explain the variability seen in clonally related strains, nor the speed in which those changes seems to happen (see 4°C effect above, and ASL experiments shown below; for ASL definition see below or Material and methods). Nevertheless, this is a well-known scenario studied by the population genetics community: asexually dividing organism do accumulate mildly deleterious mutations due to the process known as Muller’s ratchet^93–95^. Nothing is known about the fission yeast life cycle in the wild^77^ (i.e. whether they reproduce mainly through an asexual life cycle or whether they frequently mate and reproduce mainly through a sexual life cycle), so it could well be that, by dividing mainly through asexual divisions (mitotic divisions), wild fission yeast cells have accumulated a high genetic load of mildly deleterious mutations that might explain the observed low spore viability values in fission yeast self-crosses (see Figure 2A).

To test for the Muller’s ratchet hypothesis, Advanced Inter-Cross Lines (AILs)^96^ where generated with the aim to clean the experimental cell population from the accumulated deleterious mutations, thanks to recombination and further selection of the fitter cells. One would expect that, if a spore killing allele is present in one of the two strain used in an inter-strain cross, the progression through an AIL would eliminate this variant from the population, as the cells carrying this mutation would produce less viable spores and, therefore, this allele would become underrepresented upon progression into the AIL experiment (see Supplemental Figure 5A, left panel). Two inter-strain crosses, which have a relative high inter-strain spore survival value (and therefore a low deleterious mutation load), where selected to facilitate the identification of the putative causative mutations: the JB32 x JB761, and the JB759 x JB760 crosses were chosen (showing 67.41% and 63.49% spore survival in an azygotic meiosis self-cross; and 73.5% and 41.98% in a zygotic meiosis self-cross; see Supplemental Figure 4A for azygotic matings, and Figure 5A for zygotic matings). In those experiments, a clear increase in spore survival is seen with the AIL progression (see Figure 5A and Table S2).

As a control to the AILs experiments, Advanced Self-Cross Lines (ASLs) were generated. In those experiments a similar protocol than in AILs experiments were used, but the starting clones used in those experiments were two differentially tagged clones of the same strain (see Materials and Methods). If the defects that originate the self-cross issue were genetic (Muller’s ratchet hypothesis), and, the spore survival increase observed in the AILs experiments, were due to the recombination, and further selection of the fitter alleles, one would expect that no spore survival increase should be seen in an ASL experiment. Surprisingly, in 2 out of 4 of the ASLs experiments performed (from the four strains used in the previously described AILs experiments) a clear spore survival increases were observed (see Figure 5B and Table S2). This observation was totally unexpected, and it is difficult to explain under the Muller’s ratchet hypothesis (see General discussion). I decided, then, to focus on this phenomenon as the main focus of study, to try to characterize the self-cross problems detected. The most spectacular observation is, without any doubt, the JB760 example. In this example, the spore survival values vary from 15.57% (in the original JB760 zygotic self-cross) to 88.68% (in the JB760 ASL 8th cycle), with the top value observed, in the JB760 ASL 7th cycle, being as high as 93.27% (see Figure 5B, bottom right panel). Given the exceptionality of this observation, I repeated, in triplicate, the JB760 ASL experiment, starting from three independently isolated haploid clones from the original JB760 -80°C stock (see Table S1). In the three replicate experiments, although some important differences are seen during the ASL experiment progression, in all cases, after 7 to 10 ASL cycles a high spore survival values were achieved (see Supplemental Figure 5B and Table S2).

To test whether this observation is an exception or the rule, I analysed nine more ASL from different strains. Two of them: L972 and JB953 were analysed with an ASL experiment up to 8th cycles (see Figure 5C and Table S2), while the seven remaining strains (JB838, JB845, JB853, JB872, JB873, JB884 and JB913), were analysed up to the 4th ASL cycle (see Figure 5D and Table S2). In 8 out of 9 of the extra examples analysed, a significant change in the self-cross spore survival value was observed (see Figures 5C and 5D, and Table S2). In total, in 11 out of 13 (84,6%) ASL experiments performed, a clear effect was observed (see Figure 5 and Table S2). Again, in most of the non-significant examples, a clear tendency towards a spore survival increase, is observed (see Figure 5B, 5C and 5D).

Moreover, in fact, the ASL effect on the directionality seems to counteract the previously described general directionality observed in the 4°C effect (see above). This is particularly evident in the JB953 and JB873 ASL examples. As detailed in Material and methods, ASLs experiments were initially started from a single diploid clone (ASL azygotic Start, see Supplemental Figure 5A, middle panel). This, though, was not the best choice, due the described zygotic *vs* azygotic significant changes that are observed when studying a self-cross (see above). This fact, was not yet fully known by this author at the starting of the development of the ASL experiments. This fact, together with the fact that ASL spore survival determinations, in cycles greater than one, are more easily performed in zygotic that in azygotic matings (to avoid having to select for many diploids at any given ASL cycle), lead to the use of previously obtained zygotic mating spore survival values as a starting value (cycle 0) for the ASL experiments (done mainly with the same cells). This fact is reflected by a vertical dotted line in all ASL (or AIL) representing graphs (see Figure 5A, 5B, 5C, 5D and Supplemental Figure 5B). All these experimental problems lead to the detection of the same problem as seen in the phenomenon observed with the -80°C effect (see Figure 4B). This is reflected in the ASL experiments of JB953 and JB873 (see Figures 5C and 5D, JB953 and JB873 examples). This is, as well, true for the strains JB845 and JB913, where the cycle 1 has a slight reduction in spore survival compared with the cycle 0. Indeed, if cycle 1 is used as reference to check for change when comparing with cycle 4, JB845 example cycle 4 shows as significant (p-value < 0.01), while JB913 example increases its significant level (from p-value < 0.001 to p-value < 0.0001; see Table S2). All this might indicate that the general tendency of reduction in spore survival due to the 4°C or -80°C effects are slightly masking the effects of the ASL progression (that shows an opposite general tendency to the one shown by the previously named experimental conditions).

It should be noted that in the strain JB761, where no apparent change seems to happen along the ASL (see Figure 5B, top-right panel), among its unique identified mutations^58^ it has a point mutation that generates an amino-acid change in the Cdc28 protein (R735P). Deletion of the *cdc28* gene has a strong effect in spore survival of the spores^97^, so it might well be that a single point mutation in this gene, has, as well, an effect on spore survival. In this case, this would be an example of the Muller’s ratchet affected spore survival lost, and, as discussed earlier, no change in the spore survival phenotype would be expected in an ASL experiment, as actually seen (see Figure 5B, top-right panel, and General discussion).

As well as the consistent directionality shown by the ASLs experiments, in the spore survival changes, it is remarkable the speed at with those phenotypic changes take place during a ASL experiment. As explained in Materials and Methods, during the spore germination step (see Supplemental Figure 5A), EtOH treated samples were diluted to a low OD cultures (0.8 ml cultures of the treated cells were diluted in 200 ml of fresh YES culture. These produced cultures of around 0.2 OD units). In the next day or two they grow up to saturation (approx. up to 10 OD units, see Supplemental Figure 5C). The OD measurements done in the starting of those cultures (from EtOH treated cells) is not completely comparable to the OD measurements taken to a freshly growing exponentially growing cells. The treatment, with 30% EtOH, of the mating mixture left a mixture of unmated (or unsporulated) dead parental cells, mated (or sporulated) dead parental cells with spores inside (live or dead), released free spores (live or dead), and cell debris. This does not allow a proper counting of the starting live cells (specially because of not having any efficient way to distinguish live *vs* dead spores). Anyway, at the end of this step, you end up with a saturated culture (2 to 10 OD units; arithmetic mean = 6.1 ODs units) of fission yeast cells that come from the live spores that were present in the initial EtOH treated culture. The treatment with 30% EtOH completely kill all parental cells (data not shown). Taking the initial OD measurement as an approximate measurement of the initial live cells, one can estimate the number of generations that happen during the growing of this culture step (germination step in Supplemental Figure 5A). A mean of nearly 5 generations it is observed (see Supplemental Figure 5C). Taking the standard fission yeast mutation rate described in the literature: 2.00 X 10^-10^ mutations per site per generation^98^, this gets an estimated number of 0.0125 mutations per cell (12,5 X 10^6^ sites per cell x 2.00 X 10^-10^ mutations per site per generation x 5 generations = 0.0125 mutations per cell). This is a really small number of mutation per cell to explain the observed recovery of the spore survival during an ASL experiment as the simple effect of the selection of a retromutation that corrects the genetic mutation that the wild fission yeast might have accumulated due to the genetic Muller’s ratchet. Even going through 7 to 10 meiosis cycles (which give 0.08 to 0.1 mutations per cell), still gives a really low number of accumulated mutations to explain this phenomenon. Moreover, given that those experiments were done with in a naturally haplontic free living organism, like fission yeast, that produces haploid spores, who develop in haploid organisms, one should note that, if an accumulated genetic mutation has happened, there is no room for a backup copy of the right genetic information. So, in front of a genetic mutation, and with no genetic backup information available, how would the cell know which is the right information to put back and correct the genetic mutation?

Even more importantly, during the realization of the ASL experiment, this author noted that, as one progress through the ASL experiment, the spore survival phenotype is not the only phenotype that do change. Several completely independent phenotypic changes were observed by this author in different strains. This include: 1) changes in the mating efficiency were observed in the strains JB32, JB759, JB761, JB845 and JB953 when comparing later ASL cycles with earlier ASL cycles (not shown), in most of the examples but one (JB845) this is an increase in the mating efficiency; 2) changes in the colony sizes were observed in the strains JB760, JB884 and JB913 (see Supplemental Figure 5D for JB760 example); 3) changes in cell flocculation were observed in the strains JB864 and JB953 (see Supplemental Figures 5E and 5F for JB953 example); 4) Changes in germination speed of the colonies (quicker germination speed) were observed in the strains JB760, JB838, JB872, JB884 and JB913 (not shown). This is just a glance of the unintentionally observed phenotypic changes that happen along the ASLs experiments. A comprehensive study with proper measurements should be done to characterize this phenomenon. All those observations, made even more unlikely a classic mutation- retromutation scenario. If just explaining the recovery of the spore survival is a difficult task, explaining all those phenotypic changes happening at once by retromutation and selection in such a low number of generations is a nearly impossible task.

In summary, all the observation collected during the ASLs experiments are not easily explained by a classic mutation-retromutation scenario. This fact suggests that, there it might be a non-genetic meiotic repair program that takes place in all cells during meiosis. The observations that backs this later statement are basically the following ones: 1) several phenotypic changes take place in a short number of generations during an ASL experiment; 2) most of those changes seems to have a clear directionality, i.e. the spore survival show a general pattern of spore survival recovery. This seems, as well, to be true in most of the other phenotypes that were observed to change during the ASLs experiments (they change from a worst to a best scenario); and 3) the lack of genetic backup in haploid organisms. All this makes highly unlikely that all those changes are due to a classis mutation- retromutation scenario, and this leads to the proposal of a non-genetic mechanisms, as the responsible for all those phenotypic changes.

### A vegetative cell fitness increase is seen in cells that go successively through several meiotic cycles

If the statement done in the previous section is true: the existence of a non-genetic meiotic repair program does happen during an ASL experiment, one would expect that cells from later cycles of an ASL experiment are in general fitter than they non-ASL-subjected starting cells. This is an easily testable prediction, thanks of having *ade6-tagged* strains which can easily be followed in a competition experiment.

The strain XM637, which is an JB953 *ade6-Nat tagged* version generated in this study (see Table S1), was subjected to six competition experiments with different populations of *ade6- Hyg tagged* strains: first, with the strain XM638 (a *ade6-Hyg tagged* version of the JB953 strain, see Table S1). This is a non-ASL subjected starting cell; and second, five *ade6-Hyg tagged* haploid populations obtained from the haploid mixtures generated in the cycles 1 to 5 of the JB953 ASL experiment (see Materials and Methods). A clear enrichment in *ade6- Hyg* tagged cells was observed in later ASL cycles, compared with what it was observed in earlier ASL cycles (see Figure 6A and Table S2). This clearly indicates that the ASL generated cells are fitter than the starting clone used to start the ASL experiment. *Ade6- Hyg tagged* cell enrichment it was observed after the third ASL cycle had been performed (see Figure 6A), this coincides with the main change observed in the JB953 flocculation phenotype (see Supplemental Figure 5F), but doesn’t coincide with a detectable change in the spore survival of a zygotic self-cross (see Figure 5C, right panel). This indicate that, not only more than one phenotypic change is happening as one progress in an ASL experiment, but that not all of them are happening at the same time. Altogether this experiment corroborates the predicted increase in cell fitness as one induces cells to go into repeated meiotic cycles. I would like to emphasize, that this observation agrees with the proposed existence of an overall correlation between the spore survival recovery, and the cell survival recovery.

**Figure 6:**
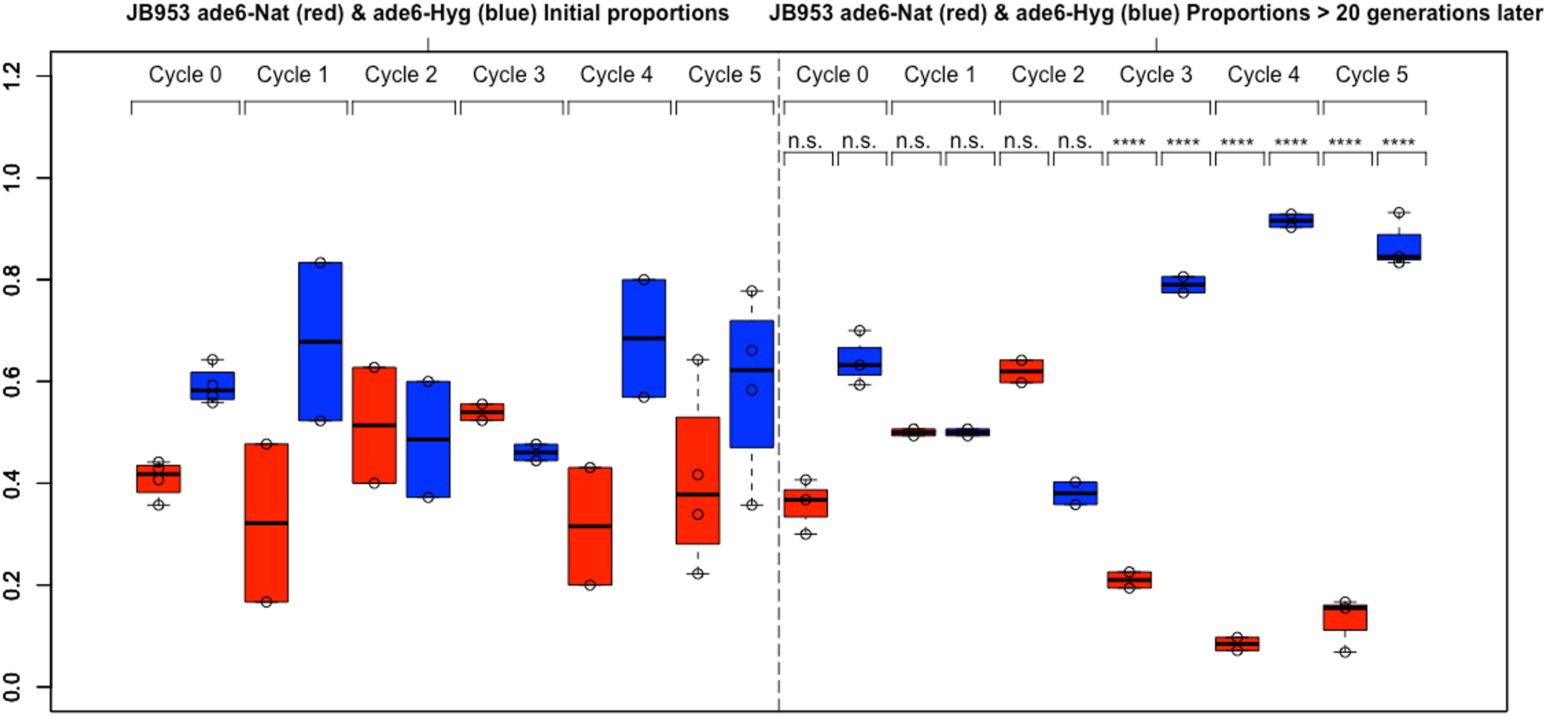
Vegetative cell fitness increase in ASL derived cells. Vegetative cell fitness increase it is observed in cells from later ASL cycles (right panels) compared with the starting cells used in the ASL experiment (left panels). Nourseothricin tagged cells (ade6-Nat) are coloured in red, while Hygromycin tagged cells (ade6-Hyg) are coloured in blue. Significance level are plotted as described in Material and Methods (see Materials and Methods and Table S2).

In summary: during an ASL experiment, among all the phenotypes that do change detected before (see above), we should include in the list an overall increase in cell fitness, or what it is the same, an increased ability of the vegetative cells to survive.

### The survival value of the spores of the F1 generation depends on the survival value of the spores of the parental generation

Yet another surprise came from the analysis of the self-cross spore survival in fission yeasts: the self-cross spore survival values were affected when cells were kept in minimal media without nitrogen (MM-N) for up to two months (longer times were not analysed). In those experiments, freshly growing haploid cells were put in 50 ml flasks with MM-N and were kept at 25°C for up to two months (see Materials and methods). Those cells were tested for spore survival in a zygotic meiosis self-cross, by merging cells from opposite mating types, at certain timepoints after the starting of the exposure to MM-N. Two different strains were analysed: JB32, which shows initially a high spore survival; and JB760, which shows initially a low spore survival (see Figure 3A, JB32M and JB760M rows). Two biological repeats in each of the strains were analysed. In both cases a clear change pattern was observed along the timecourse, although different behaviours were observed: while a slight decrease is observed in JB32M cells, an initial increase is observed in JB760M. The initial increase in JB760M it is followed by a decrease/maintenance afterwards (see Figure 7A and Table S2). Spore survival increases at initial steps of one experiment has been previously observed (see JB758 strain in Figure 4B) or in other independent experiments carried out by this author (i.e. in chronological lifespan experiments, some strains show an initial higher mortality, which leads to the survival of the fitter cells from the population. Those cells presumably might have higher spore survival; data not shown).

**Figure 7:**
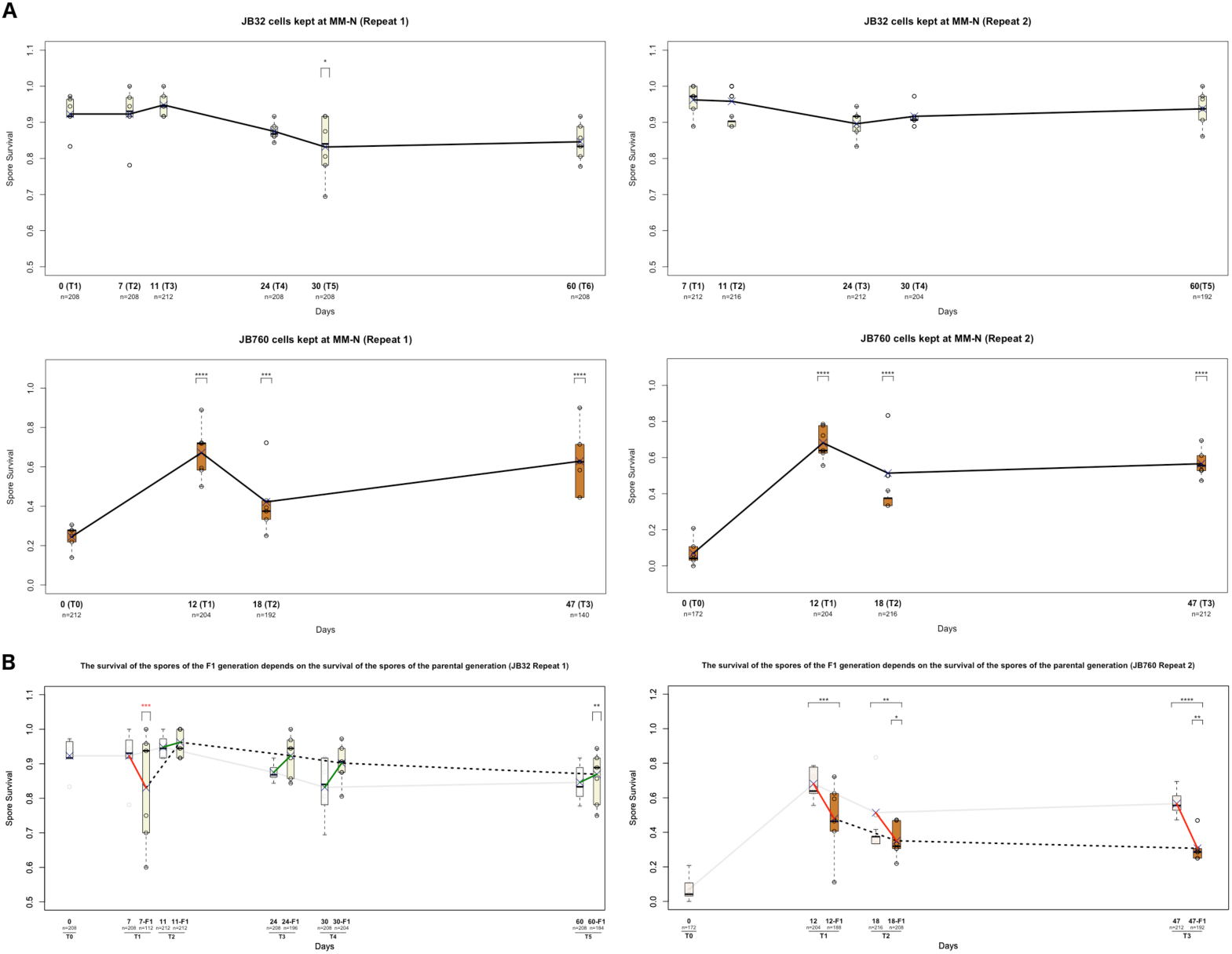
Spore survival variation seen in liquid MM-N kept cells and epigenetic degeneration phenomenon. **(A)** Haploid cells from opposite mating types kept for up to two months in MM-N at 25°C were joined at several timepoints, and the spore survival of the self-cross was determined. Two strains were analysed: JB32 and JB760. Two biological repeats of each was carried out. Statistical significances are shown on top. Significance has been calculated by comparing each timepoint with the first one available **(B)** Cells from JB32 MM-N kept cells from repeat 1 and cells from JB760 MM-N kept cells repeat 2 were tested for the spore survival after one cycle of meiosis was carried out (F1 cells; done as in the ASL experiment see Materials and methods). Original MM-N timecourse is plotted fainted. Original and F1 cells are joined with straight lines: red (when there is a spore survival reduction) or green (when there is a spore survival increase). F1 cells are joined with a black dotted line, to indicate that there is no direct continuity from the cells used in one determination from the cells used in the following determination (the continuity is through the parental cells). Statistical significances are shown on top. Double sized lines show the comparison between original and F1 cells derived from them, while single sized lines show significance from each F1 timepoint with the reference timepoint. In JB32 example T2 timepoint was used as reference, while in the JB760 example T1 timepoint was used as a reference. Significance level are plotted as described in Material and Methods (see Materials and Methods and Table S2).

This manuscript is just the first one to uncover the variation in self-cross spore survival values due to environmental factors. Examples of differently behaving repeat experiments can be seen in the replica experiments done to recheck JB760 ASL (see Supplemental Figure 5B). In particular, the JB760M crosses showed, in all the experiments that I carried out a high zygotic spore survival variability (see Supplemental Figure 6A). This is not observed in two JB32M independent zygotic spore survival determination (see Supplemental Figure 6B). As, environmental variations were not initially expected, I did not carefully control the exact procedure to prepare the cells in the different experiments carried out. It could be that, in each one of the experiments, cells treated unconsciously in a different way were used. Alternatively, clonal variation could just be due to a quickly changing nature of the causative defect leading to a low self-cross spore survival. See General discussion for a detailed discussion on the implications that a higher JB760M variability, compared with JB760 might indicate (see General discussion). Anyway, the exact mechanisms that might lead to specific changes of the self-cross spore survival changes in a given strain is beyond the scope of this manuscript.

The change in the self-cross spore survival values observed alongside those timecourses, reinforces the previously stated interpretation that, the spore survival in a self-cross, it is not a given value of a given strain, but a quickly changing phenotype affected by quite different environmental factors. So far, this manuscript describes the unintentionally detection of the temperature related environmental factors (4°C and -80°C) and the media maintenance factors (MM-N). It is well possible that many more different environmental stresses produce similar effects.

An extra test, complementary to what it was seen in Figure 5 (recovery/change of the spore survival in meiosis), was carried out in those cells. In two examples: JB32M liquid MM-N kept cells repeat 1 and in JB760M liquid MM-N kept cells repeat 2, at almost all timepoints, cells were induced to enter into meiosis and processed, as in an ASL experiment (see Material and methods), to characterize the spore survival of the F1 generation (after one meiotic cycle has taken place). In those experiments, a clear pattern it was observed: the behaviour of the F1 cells along the timecourse mimic the behaviour of the parental cells (see Figure 7B and Table S2). This suggested that the survival of the F1 generation depends on which is the survival shown by the parental generation, and that any change that it is observed in the parental generation it is as well observed in the F1 generation. This might simply be the fact of a limited capacity of the above proposed non-genetic repair/change of the spore survival value in a single meiosis cycle. This interpretation might explain why, in most of the ASL examples, shown in Figure 5 and in Supplemental Figure 5, more than one meiotic cycle is needed to achieve a high spore survival value. Remember, though, that, in JB760 ASLs repeats, it was seen that, although a reduction could be seen in initial cycles of an ASL, after many meiotic cycles, almost full recovery of the spore survival, as well as of other phenotypes, is always seen (see Supplemental Figure 5B and 5D and data not shown). MANOVA tests showed that, while in JB760M, a clear effect is seen of the generation in shaping the spore survival values (p-value=6.94E-06; see Figure 7B and Table S4), this is not the case in the JB32M example (p-value=0.09; see Figure 7B and Table S4). The lower spore survival changes observed in the JB32M examples (given that it is already in a high range of the spore survival) might require a higher sample size to detect significance.

These experiments suggest that, the non-genetic changes that the parental generation has are, somehow, reflected in the non-genetic driven fitness of the F1 generation. This might probably due to a limited capacity of change of the spore survival in just one meiotic cycle (see Figure 7B). In practical terms, this might simply mean that cells (and multicellular organisms), might need to go often through meiotic cycles, to avoid the accumulation of a high non-genetic defects load, accumulated by the parental generations.

## 4. General discussion

### Justification of the general discussion section and of the decision to share this data, and thoughts, through a bioRxiv manuscript

This manuscript is full of unexpected interesting findings that describe an intriguing phenomenon related to the values observed when one analyses the spore survival of a self-cross in a bunch of fission yeast collected from all around the world^58^. The data obtained so far, and the conclusions that could be directly obtained from that data are exposed in the previous section of this manuscript (see Results and discussion). In this section, I would like to highlight that, among the interesting observations that are described in this manuscript, there are some that, if proven to be the rule rather than the exception (something not achieved in this manuscript), would allow to describe a simple and coherent way to define the ageing process as an epigenetically-based ratchet-like mechanisms, and to conceive ageing itself as a loss of epigenetic information phenomenon. This is not a trivial statement.

In this section, I will try to do my best to provide the reader with an experimental framework to tackle this question, and with a theoretical framework to justify it.

My current situation, as an independent researcher, does not allow me to advance on this project just by myself. I am sending this manuscript to the bioRxiv repository to make these data and thoughts publicly available, with the hope of gathering the necessary supports to advance, with the proposed approach, in the study of aging.

At the end of this section I will speculate about what would imply, in the biomedical research, the confirmation of the statement made just above about the nature of the ageing process.

### The key observations

As previously said, this study uncovers a group of unexpected observations, while working with fission yeast wild isolates, and trying to characterize their spore survival by tetrad analysis after a mating has occurred. From all these unexpected observations, one of them clearly stands up: defective haploid cells are able to recover self-cross spore survival defects, just by going through meiosis. This was observed originally in self-cross spore survival data, but, while pursuing this goal, evidences of other phenotypic changes were observed (see Figure 5 and Supplemental Figure 5). These observations are simply spectacular.

The second key observation made during this manuscript was a possible correlation between the spore survival values and the vegetative cell survival data. This is not clearly addressed in this manuscript (neither was its purpose), but in my opinion is a key question to address. The data that suggests this correlation are the following observations: 1) in the JB953 strain, a different pattern of spore survival loss is seen, when comparing the 4°C and the -80°C induced loss of the spore survival values, between zygotic matings and azygotic matings (see Figure 4A, 4B, 4C and 4D). In some other strains this is not true: i.e. JB1205 4°C mediated self-cross spore survival loss, and JB873 -80°C mediated self-cross spore survival loss show the same patterns of spore survival loss when comparing zygotic with azygotic matings (see Figure 4A and 4B; detailed view of JB953 strain is shown in Figure 4C, left panel; detailed view of JB873 strain is shown in Figure 4D, middle panel; while detailed view of JB1205 strain is shown in Figure 4C, right panel); 2) In the JB953 strain, the 4°C stress show a qualitative loss of cell viability when comparing the vegetative cell survival between haploid cells (used for zygotic matings) and diploid cells (used for azygotic matings; see Figure 4E). The spore survival loss and the cell vegetative loss show the same exact behaviour; 3) the same phenomena is observed when comparing the JB873 -80°C mediated self-cross spore survival loss (see detailed view in Figure 4D, middle panel), with the -80°C vegetative cell survival loss (see Figure 4F). Again, the same behaviour between the spore survival loss and the vegetative cell survival loss is seen; 4) the JB953 ASL experiment show a spore survival increase, while when testing for fitness increase in JB953 ASL treated cells this is as well observed (this observation indirectly implies an increase in the vegetative cell survival).

The corroboration of this correlation is a must do to support the following non-trivial statement: this correlation imply that all cells go through an underlying biological phenomenon, that reduces its viability, and ultimately causes its death. In this manuscript, I propose that this underlying biological phenomenon is, indeed, the ageing process.

Those two key observations together, if proven to be the rule instead of anecdotic observations, allow to describe ageing as a non-genetic, information based loss mechanism, that happens in all non-meiotic cells, while, allows, as well, to describe the rejuvenation process, as an active specialized differentiation mechanism, which happens during meiosis, and that corrects the defects accumulated during the non-meiotic cell cycles (see Figure 8 and later in the text for a theoretical development of those ideas).

**Figure 8:**
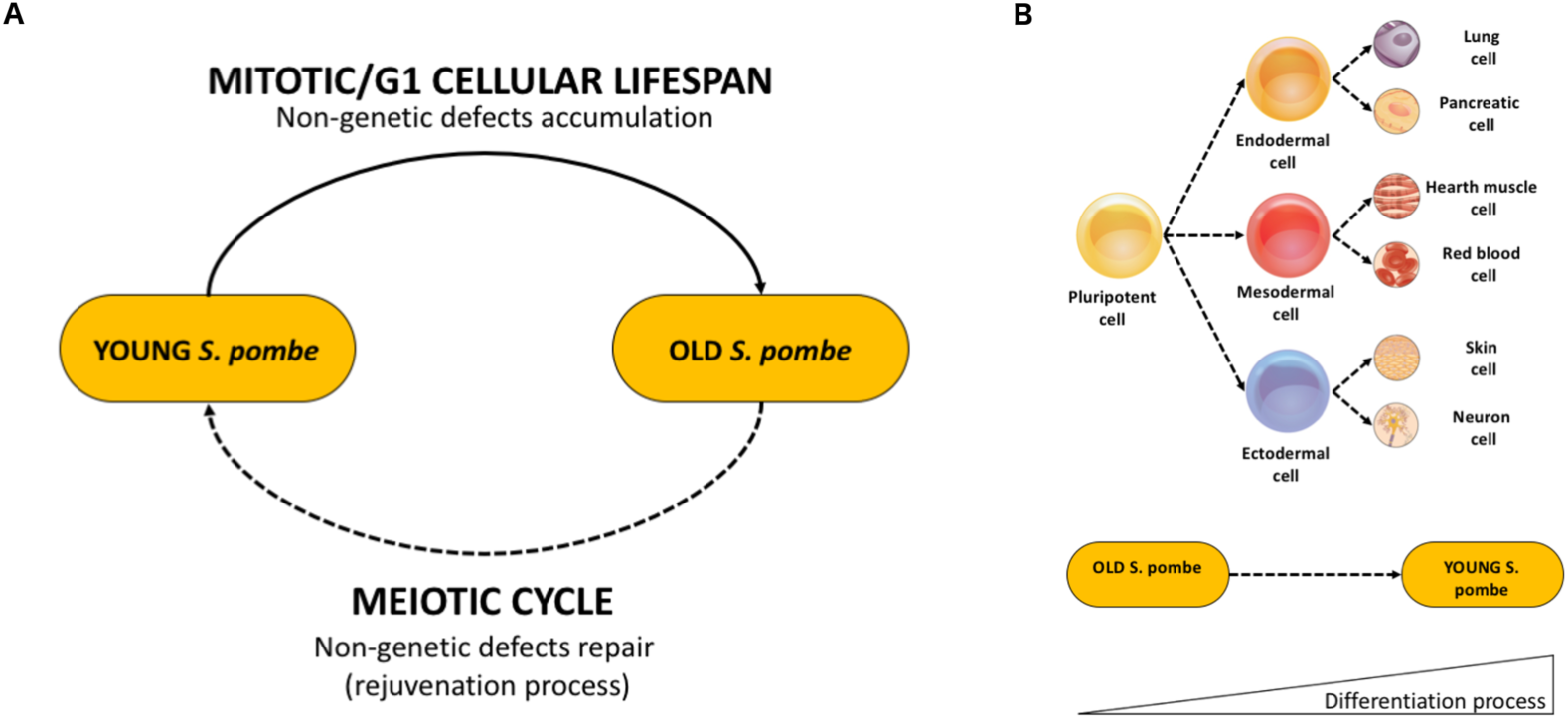
Cyclical, differentiation-like nature of the ageing process. **(A)** Schematic representation on how a simple cyclical process, with two different phases, can allow explaining the ageing process: 1) the mitotic/G1 phase (top), in which non-genetic defects are accumulated; and 2) a meiotic phase (bottom), in which non-genetic defects are specifically repaired. The non-continuous line in the meiotic process indicates that, in most cases, this process did not go back to the same individual cell, but to its next generation (see main text for details). **(B)** Schematic representation of how the rejuvenation process can be understood as a specialized differentiation process (see main text for details). This graphic, was, in some parts, adapted from the "422 Feature Stem Cell.jpg" file obtained from the Version 8.25 of the Textbook: OpenStax Anatomy and Physiology, Published May 18, 2016. By OpenStax Anatomy and PhysiologyOpenStax [CC BY 4.0 (http://creativecommons.org/licenses/by/4.0)], via Wikimedia Commons.

### An experimental framework that will allow to provide experimental corroboration of the above proposed model

#### Dominant and recessive epigenetic mutations characterization

Another of the interesting observations uncovered in this manuscript is the huge phenotypic variability that fission yeast clonally related strain show with a very little number of SNPs differences (see Figure 1C, 1D, Supplemental Figure 1C, Figure 2B, Supplemental Figure 3C and 3D, and Table S3). This difference is observed in both azygotic and zygotic matings. As mentioned before, this might simply account for the disparity of wild type spore survival values seen in the literature^6,16,21–29^. However, it could not be discarded, after seeing how easily phenotypic changes can happen, that the differences, seen in the literature, are due to this easiness in phenotypic changes. It is relevant, though, that the Leupold’s derived strains are quite resistant to, at least one of the described induced defects, the -80°C defect (see Figures 4B and 4D). This might explain why this phenomenon has not been detected before, while working just with Leupold’s derived strains, as most of the fission yeast laboratories do^61^.

Interestingly, in some of those clonal variants, signs of dominance and recessiveness behaviours can be observed in the inter-strains crosses performed (see Figure 1E and Figure 3E). In particular, in the L972 strain, the zygotic specific mating defect show a recessive behaviour, while the azygotic specific mating defect of the same strain, mated with the same clonally related partner (JB32), show a dominant defect (see Figure 3E, Table S1 and Table S5). As previously said, the number of SNPs differences between L972 and JB32 has been determined to be as low as 29 SNPs (see Table S3). How come can just 29 SNPs explain this scenario? Luckily, L972 is one of the strains that I subjected to an ASL experiment. As clearly seen, after a certain number of meiotic cycles, the zygotic specific meiotic defect starts to recover after a certain number of meiotic cycles (see Figure 5C). As discussed later, this is not easily explainable if the defect were due to a genetic defect (see below). What else could be then? Considering the observations just explained above, and easy explanation can be proposed: the defects are epigenetic defects. Why epigenetic defects? Because a non-genetic defect can only be dominant or recessive if it is caused by a factor that is closely linked to the DNA (*Epi*= “on top”). Given this, the data presented in this manuscript might suggest the existence of dominant epigenetic defects (or dominant epimutations) and recessive epigenetic defects (or recessive epimutations; see Figure 9). This is not clearly addressed in this manuscript, however, the tools provided in this manuscript, might allow to easily tackle it.

**Figure 9:**
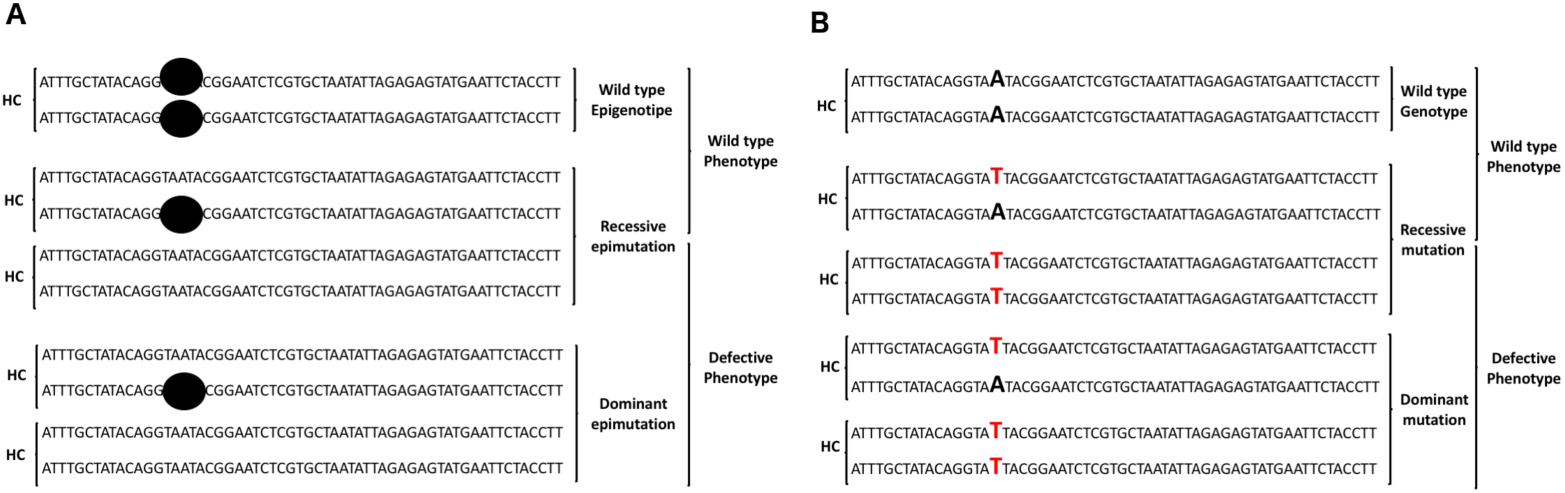
Dominant and recessive epimutations: **(A)** Schematic representation of dominant and recessive epimutations caused by the lack of one epigenetic factor (black circle) in one specific locus. **(B)** Schematic representation of a dominant and recessive mutations behaviour is shown for comparison (caused by changes in the DNA sequence). HC= Homolog Chromosomes. In **(B)** defective base pair is shown in red.

In fact, at least for the zygotic specific defect shown by the L972 strain, this seems to be the case, as genetic crosses allow to determine that it is caused by just one *locus* (see Figure 3C), crossing experiments allow to determine its recessiveness (see Supplemental Figure 3A), and ASL experiment allow to determine its non-genetic nature (see Figure 5C). This would be an example of a recessive epimutation (see Figure 9A). No azygotic ASL data have been obtained for the L972 strain (characterizing at each ASL cycle the azygotic mating spore survival value rather than the zygotic one, see Material and methods for details), neither it was tested if this defect is just due to a single locus, but if those observations were proven to be true, the azygotic specific mating defect shown by L972 (see Figure 1E and Supplemental Figure 3A) would be an example of a dominant epimutation (see Figure 9A). The combination of self-crosses, inter-strain crosses (with clonally or closely related strains), mapping for *unilocus* defects, and ASLs experiments, can help to identify more and more examples of those defects (see Table 1) in the fission yeast wild populations^58^. The examples shown in Table 1, though, are just, some of the possible test that could be done. There are many clonal groups (see Table S5) described, and cluster relations determined^58^ to easily do much more examples than the ones listed in Table 1. Even, with the strains that do not have any clonal or clustered related strains, this kind of experiments could be done thanks to the fact that an ASL experiment could allow to obtain a “clean” (not-epimutated) strain (see below).

**Table 1:** Experiments performed in this manuscript, from which conclusive date, about the epigenetic nature, and its unilocus origin could be determined by the realization of the non-determined (n.d.) gaps in this table. As in previously said, M suffixed strain numbers refers to zygotic matings (both self-crosses and inter-strain-crosses), while non-suffixed ones refer to azygotic matings. *Self-cross phenotype defective (or wild-type) phenotype is always referred as comparison with the phenotype shown by the mating partner. ** JB760 x JB759 is a cross between closely, but not clonally related strains (The SNPs difference between JB759 and JB760 is 1667). The boxplots from the cross, suggest an almost recessive behavior of the putative JB760 defect/s, but show, as well, that some other factors might be involved (see Figure 3E). It should be noted that, both JB760 and JB759 fully recover the spore survival in ASL experiments (see Figure 5B). Therefore, the interactions that affects the spore survival outcome could be due either to the 1667 SNPs differences or to the any of the undetermined number of epimutations that these two strain might have.

| Strain | Self-cross* | Inter-strain crossed with: | Outcome of the inter-strain cross | Test for unilocus defect | ASL Increase detected | Causative defect |
| --- | --- | --- | --- | --- | --- | --- |
| **L972** | defective | JB32 | dominant | n.d. | n.d. | Dominant defect/s |
| **L972M** | defective | JB32M | recessive | YES | YES | Recessive epimutation |
| **JB761** | defective | JB32 | dominant | n.d. | n.d. | Dominant defect/s |
| **JB761M** | defective | JB32M | dominant | n.d. | NO | Dominant genetic mutation/s |
| **JB868** | defective | JB906 | recessive | n.d. | n.d. | Recessive defect/s |
| **JB868** | defective | JB937 | recessive | n.d. | n.d. | Recessive defect/s |
| **JB868** | defective | JB945 | recessive | n.d. | n.d. | Recessive defect/s |
| **JB937** | defective | JB906 | recessive | n.d. | n.d. | Recessive defect/s |
| **JB760** | defective | JB759 | recessive** | n.d. | YES | Recessive** epimutation/s |

Note that JB761 it is supposed to be a genetic defect, due its inability to change spore survival in an ASL experiment (see Figure 5B). This has only been tested for zygotic meiosis, and not for azygotic meiosis. If the same behaviour was observed as well in azygotic meiosis after and ASL, this would point to the fact that in this case, the same genetic mutations is affecting both, zygotic and azygotic meiosis. As mentioned earlier, among the unique JB761 SNPs identified, there is a single base mutation that generates an amino-acid change in the Cdc28 protein (R735P). This mutation should be recreated in a strain with a good self-cross performance, both in zygotic and azygotic meiosis (i.e. JB32), to clarify if indeed, this is genetic causative defect of this self-cross spore survival loss.

Dominance and recessiveness is not only guessed from the inter-strain crosses behaviours, but as well, from environmentally induced spore survival loss, and vegetative cell survival loss seen during this manuscript. The differential behaviour of the strain JB953 spore survival/cell survival loss seen in this manuscript (see Figures 4A, 4B, 4C and 4D) might easily be due to an epigenetic recessive defect induced by the environmental factor. This would explain why diploid cells acquire the defective phenotype slower than its haploid counterparts (for a recessive epimutations, both homolog chromosomes should be affected in a diploid cell, while in haploid cells just one defect is enough to show the phenotypic defect; see Figure 9A). The behaviour seen in JB1205 and JB873, however, show the behaviour that one would expect from a dominant epimutation induced by the environment (i.e. only one defect is needed, so no difference between haploids and diploids should be expected in the speed of the acquisition of the environmentally induced defect). The same kind of experiments explained detailed above for Table 1, could be planned for those strains to test whether these defects are really caused by dominant and recessive epimutations. One clonal strain is available for the JB953 (strain JB952, see Table S5), but no clonally related strains are available for JB873 or JB1205. Anyway, as explained above, if the strain does increase its spore survival (both zygotic and azygotic) during an ASL experiment, a “clean” clonally related strain can be obtained. No test has been done in JB1205, but JB873 do show spore survival increase upon an ASL experiment (see Figure 5D). In Table 2, those experiments suggested are detailed as in Table 1.

**Table 2:** Strains in which, some signs of dominance or recessiveness could be guessed in this manuscript. As detailed in main text, conclusive data, that confirm the existence of dominant and recessive epigenetic defects, could be obtained for these strains with the realization of non-determined (n.d.) experiments detailed in the table, and with the help of clonally related strains crosses. * Defective spore survival cross it is considered if spore survival > 90%. Anyway, in all these strains regardless of what it is the original spore survival value, the defective phenotype can be induced easily (see Figure 4).

| Strain | Self-cross* | dominance/recessiveness guessed | Test for unilocus defect | ASL Increase detected | Causative defect |
| --- | --- | --- | --- | --- | --- |
| **JB953** | defective/inducible | recessive | n.d. | YES | Recessive epimutation/s |
| **JB873** | defective/inducible | dominant | n.d. | YES | Dominant epimutation/s |
| **JB1205** | defective/inducible | dominant | n.d. | n.d. | Dominant defect/s |

Again, the examples listed in Table 2, are just a few of the examples that could be obtained from within the fission yeast wild strains populations. Differential behaviour, between haploid and diploid cells can be easily determined in many more of wild fission yeast strains available^58^. This might allow to characterize genetically many more dominant or recessive epimutations.

Another observation, made during the experiments carried out in this manuscript, show that a higher phenotypic variability might exist in zygotic matings (made by platting haploids cells), compared with the phenotypic variability that azygotic matings show (made by platting diploid cells). The example observed is in the strain JB760: JB760M show a high phenotypic variability (see Supplemental Figure 6), while JB760 matings show a much more reduced variability (see Supplemental Figure 4A). This observation, if confirmed in more ASLs affected phenotypes (non-genetic ones), would support the existence of recessive epigenetic defects. One would expect that, haploids with just one copy, acquire recessive defects faster than diploid cells.

#### Characterization of the putative correlation between spore cell survival and vegetative cell survival

As mentioned above, one of the key observations has not been clearly stablished in this manuscript: the correlation between spore survival and vegetative cell survival. Two kind of experiments, that alter the spore survival have been independently identified: environmentally induced spore survival loss, and ASL linked spore survival gain. In those experiments, as detailed earlier, signs of a possible correlation between spore survival and vegetative cell survival have been observed (see above). It is, therefore, a must do experiment, to test both phenotypes in quantitative parallel experiments. This will allow to clearly corroborate if this correlation it is real and ubiquitous or not.

Carefully planned experiments should be done in which both values are measured quantitatively side by side. A special attention should be taken with spore survival determination. In my experience, although used both MEA and MM-N plates to induce meiosis and determine spore survival changes (I used mostly MEA plates), you not always obtain the same spore survival values when using one or the other kind of meiosis inducer plates (to be addressed in more detail elsewhere by this author). This is probably due, to the fact that, cells, do grow, in MEA plates, before entering meiosis. This is not the case in MM-N plates. If fitness phenotypic differences exist between clonal cells (or emerge quickly, due to quick changes), the growing step that takes place in MEA plates might alter the values by selecting for the fitter cells. Anyway, effects have been seen by this author using both kind of plates. From now on, though, every single experimental step should be done as controlled as possible. It might be more accurate to use MM-N plates on those experiments (to avoid a possible alteration due to selection for fitter cells in MEA plates).

Those experiments should be done with the strains that show a high effect to the detected environmental insults: i.e. JB953 for 4°C and -80°C effects, JB1205 for the 4°C effect, and JB873 for the -80°C effect, but it is important, as well, to determine if this is reflecting underlying mechanisms happening to all cells. This is suggested in the Figure 4D by the L968 example. In this case, no great effect is seen, but a tiny one is suggested (in spore survival). If this tiny effect is persistent, accumulative, and not only affects spore survival, but vegetative cell survival as well, this might clearly show, that, indeed, there is an underlying phenomenon happening to all cells. In fact, all yeast researchers might have experienced the observation that stocks need to be redone from time to time due to loss of viability (as apparent from the number of colonies growing from a patch of cells). This might simply be it: an underlying process is causing that cells kept in stocks are constantly losing viability.

Another easy to test correlation suggested in this manuscript is whether the 4°C effect and the -80°C effect are always correlated.

#### Sequence characterization of ASL derived cells

An obvious kind of experiments to do, in this proposed experimental framework to corroborate the proposed model of ageing, is to sequence all the strains analysed. A special attention should be put into the ASLs experiments. The behaviour than they show, strongly argues against any change in the genotype (see below for a detailed discussion), but it should not be discarded that any genotypic change can explain part of the changing behaviour. In my opinion is highly unlikely that all those phenotypic changes observed could be due only to genetic changes. If this was the case, one should consider that either: 1) any single SNP might easily influence spore survival; 2) the causative SNPs are all of them mostly located in "hidden" regions of the genome (i.e. repetitive parts of the genome that are difficult to sequence); or that 3) other non-SNPs genetic differences (i.e. structural variants (SVs): indels, rearrangements, CNVs) are responsible for all this phenotypic variability. The first two considerations are likely not true, as one should explain why this happens (not easily foreseen). A special attention, though, should be taken with other non-SNPs genetic differences.

Indeed, in a recent study it has been shown that non-SNPs genetic differences make a substantial contribution to quantitative traits in pombe, and that they do affect the phenotype even between clonally related strains^76^. This study shows, as well, that non-SNPs genetic differences have a rapid turnover in fission yeast cells. This is certainly compatible with the kind of observations obtained in this study. However, in this study, no clear estimate is available about the non-SNPs genetic change rate in fission yeast. As discussed earlier, the known mutation rate^98^ it is not able to explain the speed of the phenotypic changes observed in this study (see above). Could be that the non-SNPs genetic change speed rate might explain all this variability? In my opinion, this scenario seems unlikely, given that, even a 100 times higher non-SNPs genetic change rate, compared with the known SNPs mutation rate, would provide only 10 changes per cell in a 10 cycles ASL experiment. This is clearly not enough to be able to explain the full recovery of spore survival, and all the other changes observed in other phenotypes during the ASLs experiments performed in this study. In my opinion, what makes, this scenario really unlikely is not only any speed change rate consideration, but the consistent directionality of the changes observed during the ASLs experiments (see Figure 5 and Supplemental Figure 5). If all those phenotypic changes were due to non-SNPs genetic variants, the changes happening during meiosis (i.e. by eliminating repeated sequences during recombination), would account for the observed phenotypic changes. But, how would the cell known which exact non-SNP change configuration is the right one? should the cell always ensure that just only one CNV remains? Is all the phenotypic variability just caused for that reason? In my opinion, this seems unlikely.

Anyway, all those questions have a clear way to be answered: subject the ASL derived cells (and the original ones) to a sequence analysis. In this analysis, techniques that ensure a correct characterization of the non-SNPs genetic changes should be considered: either by applying the pipeline developed in^76^, or by using newer sequencing technologies that allow to get longer sequencing reads. This would give a more accurate answer to the actual genotypic changes that happens during an ASL experiment.

#### Characterization of the putative epigenetic changes that are causative of the phenotypic changes

If the phenotypic changes observed during the ASLs experiments, are not due to genotypic changes, and, as proposed above, are due to epigenetic changes, the next questions to ask are, obviously, the following ones: 1) which specific epigenetic changes are the responsible for those phenotypic changes? And, 2) how, the presence/absence of this specific epigenetic change/modification can explain the phenotypic change?

Those questions will not be easy to answer, and are not in the purpose of this manuscript. Once an epigenetic change has been “genetically” characterized using the tools described in this manuscript, it won’t be straightforward to locate it. If this is characterized between two clonally related strains (as most in this manuscript are), using classical genetic mappings would not give and easy answer to locate them (there won’t be genetic polymorphic markers to do a classical genetic mapping). Chip-seq experiments might help, but, one should consider that just from crossing experiments (used to “genetically” characterize them) one would have no idea of which is the specific nature of the epigenetic change/modification that might be the causative one. This information is needed *a priory* in a Chip-seq experiment. There could even be the case that, the causative defect, is due to an epigenetic factor/modification that has not yet been characterized. On top of that, as suggested by the multiple phenotypic changes happening at the same time during meiosis, characterizing, which specific epigenetic changes are responsible for a specific phenotypic change, won’t be, at all, straightforward.

Another aspect that should be taken into consideration, if one plan to use “genetic crosses” to characterize the putative epigenetic defect/s, is that, as proposed in this manuscript, the epigenetic landscape of the F1 generation it won’t be the same as the one from the parental ones. According to the proposal made in this manuscript, they would have gone through a non-genetic (epigenetic) repair program during meiosis. As discussed later, epigenetic changes are seen in mammalian cells during meiosis ^99,100^. That might simply be it. The epigenetic changes seen during meiosis, are the repair mechanisms, that restore the correct phenotypic function (see below).

All these aspects, would make that, the responses to the two above mentioned questions, would not be straightforward to obtain.

#### Experimental direct test, about the phenotypic (and lethality) recovery power of the meiotic cell cycles

The cyclical ageing model proposed above (see Figure 8A), has implicit a non-intuitive statement: ageing is a reversible process. Indeed, the model that I propose not only says that. I propose that rejuvenation is an actively process, that happens during meiosis, and that specifically reverts the loss of information occurred during ageing (see Figure 9B and below). Ageing is then, not only a reversible process, but a reversible process that happens in every single meiosis that takes place. This is, again, not a trivial statement. Ageing is characterized by an increased probability of death (both in unicellular and in pluricellular organisms). What this model imply, is that lethality due to ageing is, indeed, as well, reversed in every single meiosis, and therefore, the reason why the newer generations survive the parental generations is due to nothing but to a bunch of processes that took place during the parental meiosis, and that restore the lost functionalities accumulated by the parental generations.

One of the advantages that yeasts have in front of other model organisms is that they are quite easily malleable. This allow to a do, to yeast cells, lots of modifications much more difficult to achieve in other model organisms. Given this, one can envisage an experimental procedure to try to test if the above statements can be corroborated experimentally. What I propose here, to test this hypothesis, is to use two small tricks to uncouple meiosis from newer generation production. First, in fission yeast, *spo7* cells do enter meiosis, but do not produce the spore wall that spores normally have (Spo7 is needed to produce the spore cell wall), and so, *spo7* cells, induced to enter meiosis, end up in cells with four meiotically produced nuclei^101^. Second, meiosis can be easily, ectopically, induced in fission yeast^66,102,103^, even to haploid cells^104,105^. The experiment proposed would consist then, into ectopically induce meiosis to a group of *spo7* cells, put them back into a rich media (to allow them to re-enter a mitotic cell cycle; see below), and to compare by a Chronological Life-Span (CLS) experiment, if simply going through meiosis delays lethality. As a control, a group of isogenic cells would be treated similarly, but without inducing the entry into meiosis (or with a defective inducible system). If the proposed model is correct, one would expect that the meiosis induced cells will live longer than its non-induced isogenic cells.

A third trick, must be needed for this experiment to be successful: meiosis induced cells, do normally lyse the parental cells to free the spores. This is achieved thanks to two specific enzymes, that lyse the cell wall components: Agn2 and Eng2^106^. In consequence, all this experiment should be done in triple mutant *spo7*, *agn2* and *eng2* cells, to avoid cell wall lysis during the experiment.

I do not escape this author, that, the proposed experiment, as well as need the three tricks here discussed, might need some more tricks, or even become impossible if the four nucleated cells produced are not able no re-enter a normal mitotic cell cycle. “Return to growth” experiments have been done successfully in *S. cerevisiae*, but those experiments, are completely different from the one here proposed: cells are induced to return to growth before full commitment to meiosis is reached^107,108^. On top of that, no intention to avoid spore formation is planned on those experiments. In summary, although, if successful, the proposed experiment, would be very informative in the implication of the meiotic events into the lethality avoidance, this experiment might be tricky, or even impossible. Possible tricks to force cells to re-enter meiosis (if they do not do it spontaneously), might include to ectopically express the Tht1 protein, who is normally involved in promoting nuclear fusion (karyogamy) during zygotic meiosis^109^.

Contrary to what happens in the “return to growth” experiments done in cerevisiae^107,108^, this experiment, if successful, would not include a ploidy reduction step in any induced meiosis that has been successfully uncoupled of the spore formation. Fortunately, induced meiosis can happen both in haploids, as well as, in diploid fission yeast cells^104,105^. Given this, it would be recommendable to do this experiment using both haploid and diploid cells. This would allow to: 1) rule out any effect of ploidy in those CLS experiments; and 2) concatenate at least two induction events (starting from haploid cells), to check that any meiosis induction do really produce a consecutive lifespan increase. If ploidy increase did not produce any harm, this experiment could be tested in several reiterative cycles of meiotic induction.

If this experiment is proven impossible, alternative experiments should be thought to link meiotic epigenetic events with lethality avoidance. It should be highlighted, though, that, in agreement to this proposed experiment, in *S. cerevisiae* it has been shown that, meiosis *per se*, is able to erase the effects produced by ageing in a replicative life-span experiment^51^, this experiment aims to test the similar finding, but in a chronological lifespan experiment.

### A theoretical framework to justify ageing as an epigenetic ratchet-like based mechanisms

#### Information based nature of the biological sciences

Biology is all about information. The main characteristic of biological organisms is its ability to achieve complex tasks, and to keep the record of how to continue accomplishing those complex achievements. A proper record, of this useful achieved information, it is crucial for biological organisms. Since Mendel uncovered the laws of inheritance^110^, a clear question became apparent in the biological field: where, and how is this information stored? Watson and Crick model of the DNA gave a coherent explanation for those questions^111^. Is this all what matters? A clear answer for that question, as a negative answer, came from the observation that all the cell types, in a multicellular organism, have the same DNA information inside^112,113^. The phenotype of a specific cell (its achievements) is, therefore, not only due to the information that it is stored inside that cell, but it is as well due to which part of this information is being read in this particular cell type. In unicellular organisms, there is also a similar scenario: different environments might trigger the reading of different parts of the stored information to help the cell to cope with a particular problem that the cell is facing. Cells have, therefore, the ability to read different parts of its stored information according to their needs, and give, then, a useful answer (as phenotypically observed characteristics) to successfully overcome their given problems.

The origin of life itself, could be defined, according to what has been explained above, as the moment in which a phenotypically useful information began to be stored and transmitted to other similar structures. Since then, evolution has been maintaining and recording new information that allowed, as evolution progressed, to perform increasingly more and more complex tasks.

#### Ageing as epigenetic ratchet-like mechanisms

In this section, I plan to argue in favour of that, ageing should be considered, as well, as an information related phenomenon.

The bigger surprise in this manuscript, was, by far, the ASLs experiments. They were done initially as a control for the AILs experiments, to confirm the genetic nature of the low self-cross spore survival defects. On the contrary, though, in 11 out of 13 (84,6%) of the ASLs performed a clear change was observed (see Figure 5). The AIL experiment was done with the hypothesis of the phenomenon known as Muller’s ratchet as a causative explanation for the widely observed low self-cross spore survival seen in fission yeast wild populations (see Figure 2A). ASLs experiments, were planned as simple control to corroborate the genetic origin of the phenotypic defects. The ASLs experiments gave, however, a spectacular result: haploid derived cells (diploid homozygotes created from a haploid cell), were able to recover from the phenotypic defects observed. This observation cannot be easily explained if the causative defects were genetic. Haploid organisms do not have a backup genetic information from which the defective information could be copied, and used to repair the genetic defect. The observation that most strains do increase their spore survival upon an ASL implies that most of the observed defects that cause spore survival loss were non-genetics (not due to changes into the DNA sequence).

On top of that, the ASLs observations were not isolated. The non-genetic nature, of the self-cross spore survival defects, was highly coherent with the high phenotypic variability observed in self-cross spore survival values of clonally related strains (which show little genetic variability; see Figure 1, Supplemental Figure 1, Figure 2B, Supplemental Figure 3C and 3D, and Table S3).

The recovery seen in ASL experiments implies that the defective information was somehow being put back in its right place from somewhere else. What could all this mean, in terms of information access? The ASLs experiments suggest that, during meiosis, a repair program is taking place. This repair program is probably the reading of the genetically encoded information (where else should the information be stored?), and its translation into non-genetically encoded information. Which non-genetically coded information? As discussed earlier (see above), if the existence of dominant and recessive non-genetic information could be experimentally confirmed, and shown to be generalized, this must be epigenetic information. In that sense, what it is happening during meiosis is nothing else, but a specialized form of differentiation (see Figure 8B), where, cells with the same genetic information give rise to completely different phenotypic cells thanks to epigenetic modifications. In this case, a phenotypically old cell became a phenotypically young cell. In my opinion, any differentiation program should be understood as an information reading process, where the genetic information is translated into epigenetic information, which contribute to determine the phenotypic landscape shown by the specialized cell type. The combination of genetic and epigenetic information of any given cell is what give rise to its phenotype (see Figure 10E). Indeed, experimental findings have shown that, any cell type can be reset to form pluripotent cells able to specialize again^114^.

**Figure 10:**
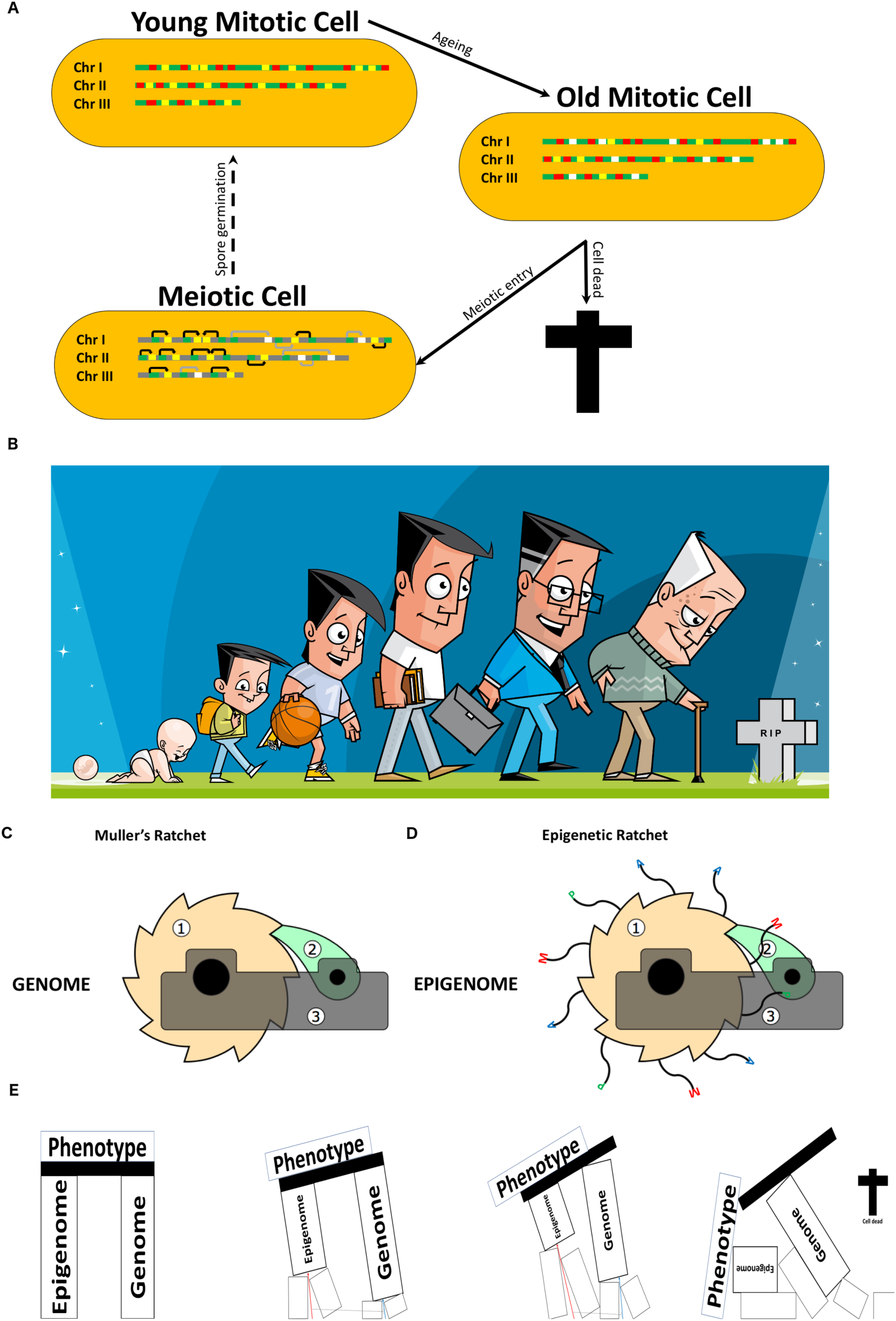
Epigenetic ratchet. **(A)** Detailed representation on how the cyclical model shown in (Figure 8A) takes place. The three fission yeast chromosomes are represented, and as a mode of simplification active regions are coloured in green while meiotic specific genes, which are silenced during normal vegetative simplified localization of functional epigenes (pieces of DNA that are necessary to achieve a phenotypic accomplishment through the epigenetic information that they contain). According to the proposed model, epigenetic defects (epimutations) are accumulate in functional epigenes during ageing. Defective epigenes are coloured in white. As suggested by the results of this study, the accumulation of those epimutations might be what drives cells to death. Meiotic genes, are activated during meiosis (they are shown in green in meiotic cells. In those cells, the rest of the chromosomes is coloured grey, as it is no relevant, for the model, whether they stay active or not). According to the proposed model, the reading of the information stored in those meiotic genes is translated into epigenetic information (black arrows), that corrects the previously accumulated defects (epimutations) in epigenes. Some defective epigenes are left white to shown that, probably, not all defects are corrected in just one single meiotic event (grey arrows symbolize, unread meiotic genes in a meiotic event). For simplification purposes, only intra-chromosomal interactions are plotted. **(B)** Cartoon symbolizing our asexual nature as organisms. From conception, to organismal death, only mitotic division takes place in the cells that form our body. **(C)** Schematic representation of the genetic Muller’s ratchet, and of the newly proposed epigenetic ratchet-like mechanism **(D)**. (1) ratchet wheel, (2) pawl, (3) ratchet bracket. In **(D)**, in the ratchet wheel, "histone tails" have been added to symbolize the ratchet wheel as a nucleosome core. A, P and M, at the end of the "histone tails", represents acetylation, phosphorylation and methylation modifications respectively. Histone tails usually have those modifications among others. For simplification purposes, only those epigenetic modifications are plotted. Note that a ratchet is a machine conceived to move just in the front direction (can’t go backwards), and that, the further it moves the worst is the damage generated. **(E)** Schematic representation on how the genomic and epigenomic information are indispensable to maintain our phenotypic achievements (functionality). The combined action of the epigenetic ratchet (red arrow) and the genetic Muller’s ratchet (blue arrow), that happens in non-meiotic cells do compromise the phenotypic functionality of the living organisms, and ultimately produces cell death. A dotted line is plotted to emphasize that the existence of the epigenetic ratchet slow down the progress of the genetic Muller’s ratchet. Licensing rights for cartoon shown in **(B)** have been bought (Dreamstime LLC, United States). Graphics from **(C)** and **(D)** have been obtained or adapted from the "Ratchet_Drawing.svg" file obtained from Dr. Schorsch (Own work) [CC BY-SA 3.0 (https://creativecommons.org/licenses/by-sa/3.0)], via Wikimedia Commons.

The proposed model to explain ageing, described in this manuscript, is based on describing ageing as a non-genetic (likely epigenetic) information-loss based mechanism. Accordingly, the rejuvenation process, is nothing else than a natural resetting mechanisms that takes place actively during meiosis (and that recovers the lost information). As described earlier, this describes a cyclical mechanism (see Figure 8A). In other words: during their non-meiotic lifetime, cells do accumulate epigenetic defects, that, ultimately, affects their viability, but, on the contrary, during meiosis, an epigenetic repair program do takes place (which might simply be the epigenetic reprograming known to happen during meiosis^99,100^). This epigenetic repair program wipe out most of the defects accumulated by the parental cells, presumably by reading the right genetic information and translating it into the right (functional) epigenetic information (see Figure 10A). The new-born individual, would have then, inherited the genetic and the epigenetic information transmitted by its parental cells, but just with one limitation, some genetic information (meiosis specific genes, including the ones that allows the reading of the right epigenetic information to restore the information lost during the ageing process), would not be read again until the next meiosis.

Indeed, most meiotic genes are known to be silenced during vegetative growth^115^. This leaves the "epigenetic repair mechanisms" inactive during this period. Allowing the cells to accumulate epigenetic defects (randomly or due to other genetic programs active during those cell cycles, or during differentiation processes in multicellular organisms). Consequently, those non-meiotic cells will have as the only source of some epigenetic information, the one that they inherited from their parents. Ageing is then, basically, a non-maintenance problem of the non-meiotic cells. In this respect, it should be noted, that, while unicellular eukaryotic organism, can divide both sexually or asexually, in multicellular organisms, this is not true for all cells. Multicellular organisms, although generally considered as sexually dividing organisms, this is only true for the species concept. The individually specialized somatic cells, that form that organism, divide only through mitotic cell divisions (Figure 10B). The only meiotically dividing cells that are present in multicellular organisms, are the specialized germline cells, who are responsible to produce the next, young, generation. In this respect, we, multicellular organisms, should be considered sexually reproducing species, but asexually reproducing organisms.

This model is reflected, though, not completely demonstrated, by the data shown in this manuscript. The environmentally induced self-cross spore survival (Figures 4A, 4B, 4C and 4D), might be, indeed, showing the phenomena of epigenetic defects accumulation, that presumably leads to cell death (this later statement, should be corroborated by the correlation between spore survival and vegetative cell survival). On the contrary, the ASLs mediated self-cross spore survival increases are showing how, epigenetics defects accumulated by its parental cells, are being corrected during meiosis (see Figures 5 and Supplemental Figure 5). The success of the *spo7* experiment (or similar) proposed above, would corroborate that meiosis, not only boost spore survival and other phenotypes (including cell fitness), but as well cell survival and death avoidance. Another key aspect suggested by this manuscript is the existence of epigenetic dominant and recessive defects. The confirmation of this as a pervasive observation, will indeed, argue in favour of an “epigenetic code” that do store phenotypic information. It will confirm that, epigenetics has, just by itself, the power to encode phenotypic information, regardless of the genetic information contained in the cell (see below).

As in the genetic Muller’s ratchet (see Figure 10C), the central point for this process is the meiosis division. Therefore, in analogy with it, an epigenetic ratchet-like mechanism, can be proposed (see Figure 10D).

According to this model, life is a continuous process to maintain the useful information (already accumulated), to achieve the necessary phenotypic accomplishments to survive. Cells, though, seemed to have a limited access to all this information. This might seem surprising. Below I will try to develop some arguments to support that, this, although contradictory, might have, indeed, more benefits than problems, from an evolutionary perspective.

This model, as well, solves one of the main mysteries of the ageing process: why germline cells are immune to ageing (amortal), while the somatic cells aren’t. The answer is clear: germline cells have access to the right information to do so.

#### Two ratchets better than one…

Having an unstoppable epigenetic ratchet-like mechanism, that drives cells (or organisms) that doesn’t divide sexually into death is, as well, indispensable to stop the progress of the genetic Muller’s ratchet. The cells that don’t go through meiosis would ultimately die due to the genetic or the epigenetic ratchet like-mechanisms (see Figure 10E). However, the cells that do enter meiosis, will be subjected to a new functional test, and to a repair mechanism. The recombination step, that happens during meiosis, helps to eliminate the mildly deleterious genetic defects. At the same time, the epigenetic repair mechanism, that happens in the meiotic process (rejuvenation step), allow to clean the cells from most of the parentally accumulated epigenetic defects.

As shown in the Figure 10E, the proposed epigenetic ratchet go faster than the already known genetic Muller’s ratchet. In other words, epigenetic defects accumulate much faster than genetic defects. This is reflected in this study: in most of the analysed ASLs experiments, a pattern compatible with an epigenetic defect is shown. Only in 1 out of 13 examples (7.6%), the expected pattern for a genetic defect is seen (see JB761 in Figure 5B). This might simply show that, many more cells, have accumulated epigenetic defects than genetic defects, due to a higher speed of the epigenetic ratchet, compared with the genetic ratchet. The higher accumulation of epigenetic defects, when compared with the accumulation of genetic defects it is a phenomenon that has been, as well, described in mammalian cells^116^.

#### The “purpose” of ageing

The proposed model of ageing detailed above has, a spectacular derived consequence: there is nothing irremediable in the ageing process. Indeed, the proper organisms know how to avoid ageing. It does it during meiosis, and it seems to avoid doing it during non-meiotic cells cycles. This leads to a simple question: why? couldn’t just be all the information always available, so that ageing doesn’t exist at all?

Short answer: No for complex organisms.

Long answer: in my opinion, questioning why ageing exist at all, it only has meaning from a teleologically based perspective, which is the predominant approach followed by us, humans. We tend to approach questions based on observations, including scientific questions, and try to understand them, by trying to answer, how we would have done it, if we had to do it, before it was done. The teleologically based thinking has been proposed to have a great impact on the interpretation of experimental results in biological science, especially in evolutionary studies^117^. Teleologically based thinking assumes that things were done according to a preconceived plan. In this author’s opinion, things just simply happen, randomly. Among all the ones that have happened randomly, the ones that work just keep going, while the ones that doesn’t work, are simply eliminated. With this reasoning, the appearance of processes like ageing and sexual reproduction increased by orders of magnitude the number of random events that could be tested, and so, ageing and sexual reproduction helped to boost the accumulation of useful information.

As we live in deeply embedded teleologically based societies, a teleologically based answer of why do we age should be useful to spread the model among the society: ageing is the price we pay for being complex. In fact, all our ancestors paid, in tiny amounts, their debts for us to be complex. We simply, just keep paying the bill.

A closer look to this aspect, will notice that among the organisms that seems not to age, are generally less complex^118,119^. As complexity increases, and organisms can do more complex tasks, this has an extra cost of increasing the chance of dying by accident. So, a complex organism, it would sooner than later die anyway. The less complex you are, the less chances you have of dying by accident, but the price you pay is a boring simple life, with less achievements. This is again a teleologically based description. In fact, if a complex amortal organisms have ever existed, it is highly likely that all of them died by accident, and therefore they didn’t proceed much in the evolutionary tree. On the contrary, the complex mortal organisms, overcome the accidentally based lethality by constantly providing young (and fit) newer organisms. The choice is then: simple and amortal or complex and mortal.

#### The success of the Eukaryotic cell

As described later, this model is coherent, with lots of other observations acquired by many more scientist (see below). Among those observations, I would like to highlight one that is really puzzling: the fact that a simple genetic base mutation in a nuclear membrane based protein could produce such a complex phenotype as a progeroid syndrome^120,121^. As discussed in the introduction, ageing is nowadays mainly perceived as a multifactorial process affected by at least nine hallmarks^2^. In this scenario, ageing is thought to be the result of a combination of the action of multiple pathways, and of multiple changes in different cell components, all over the cell. How could a simple base mutation in a protein speed up all these processes? This author can’t find an easy answer for that question. However, if we conceive ageing as a process of information loss (in this case epigenetic information), any mutation (even a point mutation) that affects a key protein needed to deal with the epigenetic information (i.e. epigenetic information storage or epigenetic information usefulness), might produce a big effect, by producing a quick, and massive information loss, that in a non-mutant scenario should take many more years to happen.

In eukaryotic cells, acquiring the nuclear membrane, could have helped to master complexity acquisition. Eukaryotic cells, with the combination, in just one process, the meiotic division, of the recombination step, and the rejuvenation step, together with the help of the nuclear membrane, in the storage, or usefulness, of the epigenetic information, might have “invented” a quick and efficient way to acquire much more useful information. The epigenetic information, would have allowed the cells to acquire an object-oriented programing language for the cell. The genetic information alone, might be analogous to the procedural programing languages, which allow to construct much less complex computer algorithms. Having a second layer of complexity acquisition, in this case the epigenetic information, with or without the help of the nuclear membrane, might have allowed, the eukaryotic cells, to store much complex “algorithms”. This fact would have boosted complexity acquisition, and allowed the cells to do really complex things like, for example, the code reuse needed, in multicellular organisms, to achieve, with one single copy of the DNA information, that the cells can have many different phenotypic outcomes (i.e. many different cell types; see Figure 8B). This epigenetically based object-oriented programing language of the cell might be, in part membrane based, as seen by the accelerated ageing in Lamin based progeroid syndromes^120,121^, but most probably not only limited to it. Increasing attention to the 3D genome organization is growing into the biological research^122^. A simple explanation for the importance of the 3D organization of the genome is that, it is what it is regulated mainly by the epigenetic information. This interpretation is compatible with recent findings that link ageing with changes in the spatial genome organization^123^. The ageing model proposed in this manuscript, allow to interpret the genome disorganization observed during ageing as a consequence of the loss of the epigenetic information needed to properly maintain it.

In summary: if genetic information alone allows to achieve levels of complexity like the complexity achieved thanks to procedural programing languages “algorithms”, having an extra layer of information storage, the epigenetic information, might have allowed the cells to acquire a higher level of complexity “algorithms”, like the boost that object-oriented programing languages allowed in comparison with the procedural programing languages. Archean and prokaryotic organisms might, probably, have similar non-genetic ways of storing information. Eukaryotic cells, though, as discussed earlier, with the meiosis process and with the help of the nuclear membrane, might have simply mastered complexity acquisition, giving rise to a whole new world of complex living forms^124^.

#### A coherent model to explain ageing

As just said, this model is coherent, with many more observations available in the scientific literature. This manuscript, gives only a series of clues, obtained from unforeseen observations, which support this model. But, although, the ageing research community, has not yet agreed a definition on what it is ageing, and therefore, not systematically seek for epigenetics as the only causative nature of ageing, several recent studies, show data that it is in complete agreement with the interpretation that this author is giving to the spore survival phenomena described in this report. Epigenetics is nowadays only considered one of the factors that affects the ageing process^2^, but, recently, the number of studies that give a prominent relevance of epigenetic factors to the ageing research are starting to become apparent. Without the pretension to be a complete review, I would like to list some of the most relevant independent observations that backs this statement: 1) a global change in DNA methylation patterns have been detected in humans all along the lifetime^125,126^, a process that has been called "epigenetic drift"; 2) aging yeast lose chromatin-associated histones during ageing and, correcting this defect increases replicative life span of yeast cells^127^; 3) meiosis eliminates aged induced cellular damage in budding yeast and resets replicative lifespan^51^; 4) DNA methylation marks show a highly elevated correlation with biological age in humans^128–130^; 5) in *C. elegans* modifications in the chromatin state of the parental cells can extend the lifespan of the progeny up to three generations^131^; 6) DNA methylations changes are associated with increased lethality^132^, this observation would clearly coincide with the observation, suggested in this paper, in which there it might be a correlation between the spore survival values (a non-genetic change), and the cell viability, which might suggest an epigenetically based mortality of the cells; 7) epigenetic dysregulation it has been shown to be a driver in ageing^123^, while epigenetic remodelling leads to a recovery of the aged defects^133^; and 8) accelerated epigenetic ageing has been associated with a progeroid syndrome^134^.

Recently acquired fission yeast data, as well, coincide with the model proposed in this report: chronological lifespan of quiescent fission yeast cells has been analysed by parallel mutant phenotyping by barcode sequencing (Bar-seq) to assay pooled haploid deletion mutants as they aged together during long-term quiescence^135^. The full list of short lived mutants obtained in this report (Jürg Bähler, personal communication) shows an enrichment in meiotic genes when analysed for specific GO term enrichments with the specifically developed tool AnGeLi^136^. Fission yeast quiescence cells show a higher chronological lifespan than freshly growing fission yeast cells in rich media^137,138^. Fission yeast quiescent state is generally considered a model for mammal quiescence cells^139^, but, in my opinion, this is not entirely true: induction of fission yeast quiescence is induced by putting fission yeast cells in nitrogen deprived medium. This environmental condition, in a wild type scenario would induce fission yeast cells to entry into meiosis. In the laboratory, due to its general use, heterothallic cells are considered as wild types, but, indeed, they are defective cells that cannot proceed normally through meiosis in nitrogen deprived media due its inability to switch mating type. This generates that, in a heterothallic haploid mixture of fission yeast cells, no meiosis is induced upon nitrogen deprivation, due the lack of a suitable partner. Those cells cannot, then, finish they entry into meiosis, but they are exposed to an environmental condition that strongly induces cells to go through it. If fission yeast cells, when going through meiosis, activate a specific program that restore cell viability (as would be confirmed by the *spo7* experiment, or similar, proposed in this manuscript), one can explain the longer lifespan of quiescent fission yeast cells, just by an, more than probably, increase in meiotic genes leakage in this condition. According to the proposed model, this might be enough to enhance the recovery of the lost information, that normally happens in a full meiosis. This might increase the quiescent cells lifespan in comparison with the non-quiescent cells by affecting the equilibrium between epigenetic defects accumulation and epigenetic defects repair (see Figure 8A and Figure 10A).

Indeed, that the lethality is produced at cellular level, and even at the organismal level, by the accumulation of epigenetic defects it is as well suggested by other independent observations like the differential mortality seen when comparing both sexes in heterogametic organisms. While in mammals females, the homogametic sex live longer than males (the heterogametic sex), this is inverse in birds, where the heterogametic sex are the males, who live longer than the heterogametic bird females^140^. Although no clear evidence for higher accumulation of mutations in the heterogametic cells have been proven to be the cause of this difference^141^, nothing has been tested about a higher epimutations accumulation in sexual chromosomes. The haploid nature of heterogametic sex, might account for more lethal consequences of the accumulation of epigenetic defects in the heterogametic sex (in mammals this would be a kind of "epigenetic unguarded X" hypothesis).

#### Meiosis dependent, Mendelian inheritance of epigenetic defects

One thing that surprises, is how frequent are those epigenetic defects in wild populations of fission yeast. This doesn’t seem to be the case of the budding yeast *Saccharomyces cerevisiae*, in which, recent studies have shown that, they even show a high spores survival values in inter-strain crosses^142^. However, in studies with natural wine related isolates of budding yeast the phenomenon of low spore viability upon a self-cross is as well observed^143–145^. From this manuscript results, one can infer that, one key factor that might affect the self-cross spore survival values, is how often do they go through sexual divisions. This aspect might simply explain the different results showed above. However, another advantage that fission yeast has in front of budding yeast is its haplontic lifestyle. If confirmed, the pervasive presence of dominant and recessive epigenetic defects among fission yeast wild populations, by the simple fact of being haploid, fission yeast should show more easily those defects (all defects are more apparent regardless of their dominance or recessiveness).

Another surprising observation is that meiotic repair of epigenetic defects not always takes place in one single meiosis. The results shown in this manuscript seems to suggest that, the repair is not as efficient when the starting defect phenotype is higher, that, when this is lower (see Figure 5 and Supplemental Figure 5). One thing to notice, though, is that, this manuscript, suggest that spore survival is not the only defect corrected, so although, the initial spore survival value might be used as an approximation, to the level of epigenetic defects accumulated (and, according to this model, to the "age" of the fission yeast), this is only an approximation and not the whole thing. The JB760 ASL replicate experiment shows that, although after several meiosis you get a full recovery (measured by the spore survival, but seen as well in colony size; not shown), you might even get a decrease of the spore survival value in-between. Altogether, these observations suggest that repair during meiosis might happen randomly, and that, with a higher level of starting defects load, no complete repair is achieved in just one meiosis.

The ability of the cell to repair in a single meiosis, seems to be limited. This is as well seen in the Figure 7B, where it is observed that, the spore survival level achieved by the F1 generation after one meiosis from their parental cells, it is dependent on which was the initial spore survival level of the parental cells. This leads to a model in which, due to limitations on the ability to repair, if the repair doesn’t happen, then the defect is inherited in a Mendelian way (i.e. in a mating between a defective cell and a non-defective cell give rise to a 2:2 inheritance of the defect; see Figure 3C), while if a full repair does happen, then deviations from Mendelian inheritance are observed (i.e. even in a mating between two defective cells a complete repair of the defect could be observed if the two defective copies are repaired). According to the proposed model, then, the inheritance of epigenetic factors might show Mendelian ratios for defects, but non-Mendelian ratios for repairs. The inheritance of epigenetic factors is, then, a meiosis repair dependent, Mendelian inheritance process (see Figure 11A and 11B). According to this model, because, repair is proposed to happen randomly, there is no way to check which is the scenario *a posteriori* (just observing repair values), without having: 1) a knowledge of the exact causative factor and nature of the defect (genetic or epigenetic, and if epigenetic, which specific epigenetic factor/modification); and 2) the complete epigenomic sequence of all the individuals involved.

**Figure 11:**
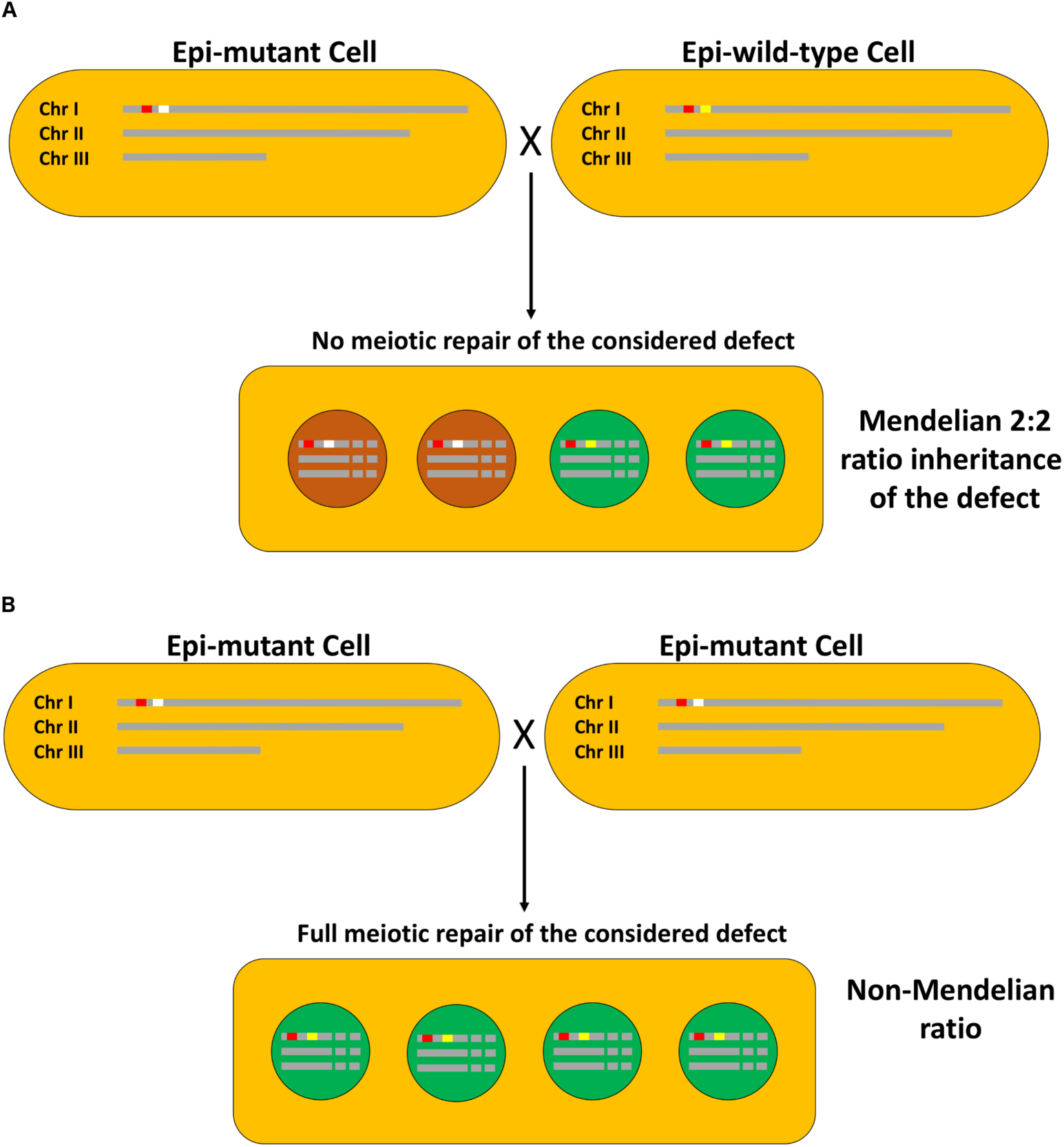
Meiosis dependent, Mendelian inheritance of epigenetic defects. Two extreme cases are shown. In **(A)** a cross between an epi-mutant and an epi- wild-type cell for a concrete phenotype created by an epigenetic defect, in a meiosis where this specific epigenetic defect has not been targeted for reparation. In **(B)** a cross between two epi-mutant cells for a concrete phenotype created by an epigenetic defect, in a meiosis in which all the defective epigenetic information has been repaired. Many more combinations between different cells (epi-mutants or epi-wild-types) and with different repair outcomes might be considered. According to the proposed model (see main text) in all cases where no specific repair has happened one could expect Mendelian ratios of inheritance, while, it is expected that, in most meiosis where a repair has happened (either in just one or both defective copies as given) non- Mendelian ratios are observed. Colours of chromosomes and genomic or epigenomic regions are plotted as in Figure 8. Spores are coloured as follow: viable spores in green; inviable or defective spores in brown-red.

It might be noticed, that, if confirmed, this model affects a commonly spread thought in biology research nowadays: "the phenotype of a given cell (or organisms) it only depends on its actual genomic sequences". GWAS studies relies on this interpretation. However, this study shows that, for example, the spore survival values of the F1 generation, is dependent on the spore survival values of their parental generation, in a situation in which not much genetic changes are expected to have happened, between the two tested parental cells (see Figure 7B). This epigenetic dependent contribution to the phenotype has already been noticed in association studies. It has been recently shown that, considering the epigenome of individual organisms along its genome, has a positive effect on the achievements accomplished by association studies^146–149^. The conception of ageing as process of loss of information, and the proposed model of inheritance of epigenetic defects (see Figure 11) has a clear impact in those approaches.

If this conception of ageing is right, then is clear that you not only need to know the genomic information to study a disease, but, you need to know, as well, the epigenomic information (which is dependent on the epigenetic information that you inherited from your parents, and to which changes has had, this information, since your conception). This might probably have a clear impact in the resolution of the missing heritability problem.

Another remarkable thing is that, if proven right, the proposed model of inheritance for the epigenetic defects, might be masking the simple nature of many so-called complex traits. Confirming that, in fission yeast wild isolates, most of the observed non-genetic defects are caused by just one locus (one epigenetic defect), might shed light into the possibility that many more than thought defects in higher organisms are as well caused by simple (one locus) epigenetic defects, rather than by the combination of many genetic defects. In this scenario, any phenotypic defect, that is thought to be genetic, but that is indeed caused by an epigenetic defect, might simply lead to a mistaken conclusion that it is complex by the fact that only the genetic information is considered when looking for causative defects.

### Implications for biomedical research of an epigenetic ratchet-like model of ageing

This author would like to emphasise, how important for biomedical research would be an ageing model as the one proposed in this report. If ageing is due to the loss of information in non-meiotic cells, but, at the same time, all cells have, indeed, the right information to correct for those defects caused by this loss of information (information that it is silenced in non-meiotic cells, but active in meiotic cells), this give an immediate direct approach to tackle all ageing related diseases: search for which is the relevant information loss causative of the defect, and try to find ways to ectopically restore the lost epigenetic information. In a scenario like the proposed in this manuscript, there is no need to search for the ways to avoid or revert ageing, it does happen in every single meiosis. In other words: there is no need to reinvent the wheel. Ageing, and all its consequences (including death) are reversed during meiosis. We just need to look at how does this happen.

But, a model like the one proposed in this report, not only could account for an explanation of the approaches to tackle ageing related diseases, but many more diseases, not apparently linked to the ageing phenomenon. As seen in Figure 7B, the spore survival of the F1 generation is dependent on the spore survival of the parental generation, probably due to a limitation in the repair capacity of the meiotic cells. If the proposed model of ageing is proven right, one could foresee, that, in organism’s lifestyle, going often through a meiotic cycle, is beneficial for their fitness. On the contrary, delaying the meiotic entry, and staying longer into non-meiotic cell types, might increase the defects accumulation in the parental germline, and therefore transmit a higher number of epigenetic defects into the next generations. We could name such a process as an epigenetic degeneration process (as it drives to a loss of useful, epigenetically achieved information, in the newer generations).

From a human related point of view, this indicate that delaying parenthood is not a good practise for the fitness of the newer generations. The emergence of culture in human populations, might have created a sudden increase (in evolutionary terms), in the delay of parenthood. This might have created and uncoupling between the biological evolutionary needs and the cultural evolutionary needs. The delaying of parenthood is, nowadays, especially relevant in contemporary western societies. According to the proposed model of ageing, and due to its presumable limitation in the repair of many epigenetic defects in just one meiosis (see Figure 7B), this might imply that delaying parenthood, drives consequently to higher epigenetic defects load in the new-borns generations. In practical terms, the more parenthood is delayed, the higher is the epigenetic defects load accumulated in the newer generations. This implies that, delaying parenthood, should cause an increase in the prevalence of any phenotypic defect that is acquired through epigenetically transmitted parental defective information.

In this regard, it should be noted that, a growing number of non-ageing related diseases, show a growing pattern of increase in prevalence. This include: food allergies and intolerances^150–153^, mental health problems^154–157^, diabetes and obesity related problems ^158–161^, impaired visual problems^162,163^, among others. Many more evidences show, as well, than a delayed parenthood is correlated with lower fitness of the offspring^164–166^. This author would like to encourage researchers working in those diseases, and many more diseases, to consider the epigenetic degeneration phenomenon described here, as a possible contributing phenomenon responsible for the observed growing prevalence of those diseases, and to check whether this phenomenon could be directly attributed to the, as well, growing phenomenon of cultural induced delaying of parenthood.

In summary, this study uncovers several unexpected observations during the study of spore survival in fission yeast, that allows the formulation of a new way to conceive ageing. According to this, ageing it is conceived as a specific loss of epigenetic information during the lifetime of non-meiotic cell types. Meiotic cells, on the contrary, might have a specific program of epigenetic reprograming that restores the lost functional epigenetic information. Although this manuscript only offers the tip of the iceberg of the possible observations to back this model, it allows to define a new simple way to conceive ageing, that is as well coherent with many more independent phenomena observed, and to design the proper experiments, in fission yeast thanks to the developed experimental framework to carefully test this hypothesis. This manuscript, in addition, provides a coherent theoretical framework to explain why a process such as aging could be the fundamental process that allowed the appearance of complex organisms such as those observed nowadays in nature.

## Supporting information

Marsellach_2017_Supplementals

## Acknowledgments and complaints

I would like to deeply thank the help of Professor Jürg Bähler into the course of this study. This study couldn’t have had place without his help. He accepted me as a postdoctoral research in his lab, and gave me all the freedom that I needed to develop my own research interests. I would like to express my deep sadness, for not finding the way we could finish this work together. I would like to specially thank Dr Alejandro Sánchez-Gracia for helpful discussions about my work. It was a three-word advice from Alejandro, on asexual populations, which allowed me to put order on all the amount of spore survival data that I had accumulated: “acumulan mutaciones deletéreas” (Spanish). I would like to thank Nada Aljassim, Emma Burton, and Josephine Hellberg for their help into some experimental work. I would like to thank Professor Fernando Azorín for his help into finishing some experimental work. I would like to thank, as well, Dr Dori Huertas, Dr Lucas Carey, Professor Ramon Trullas, Dr Alejandro Vaquero, Professor Manel Esteller, Dr Ethel Queralt, and Professor Joaquín Ariño for their willingness to help although those attempts couldn’t succeed. I would like to thank Bart Nieuwenhuis for suggestions on the presentations of the figures.

Funding: X.M acknowledges to have received funding from: Fundación Ramón A. (C.I.F.: G- 28459311), Marie Curie IEF (Project No: 235502), and funding from the Wellcome Trust through core funding in Jürg’s Bähler laboratory.

I would like to point out the most energetic complaint about the scientific policy carried out by the governments of Spain, which has condemned many “young” scientist to exile, irrelevance, or unemployment. At the same time, I would like to criticize the scientific policy chosen by the Catalan governments, which, although giving a higher priority to scientific research than the Spanish governments didn’t offer any suitable solution for “young” scientist to prosper within the Catalan research system. The Catalan governments chose to fund mainly the *primadonna*-based scientific research. This decision has allowed Catalan science to have a greater visibility in the international scientific community, but has also involved all the inconveniences linked to this conception of how scientific research should be. This policy left all the problems created by the shortage due to the incompetence of the Spanish governments, unsolved.

