## Supplementary material for "A non-genetic meiotic repair program inferred from spore survival values in fission yeast wild isolates: a clue for an epigenetic ratchet-like model of ageing?": Marsellach_2017_Supplementals

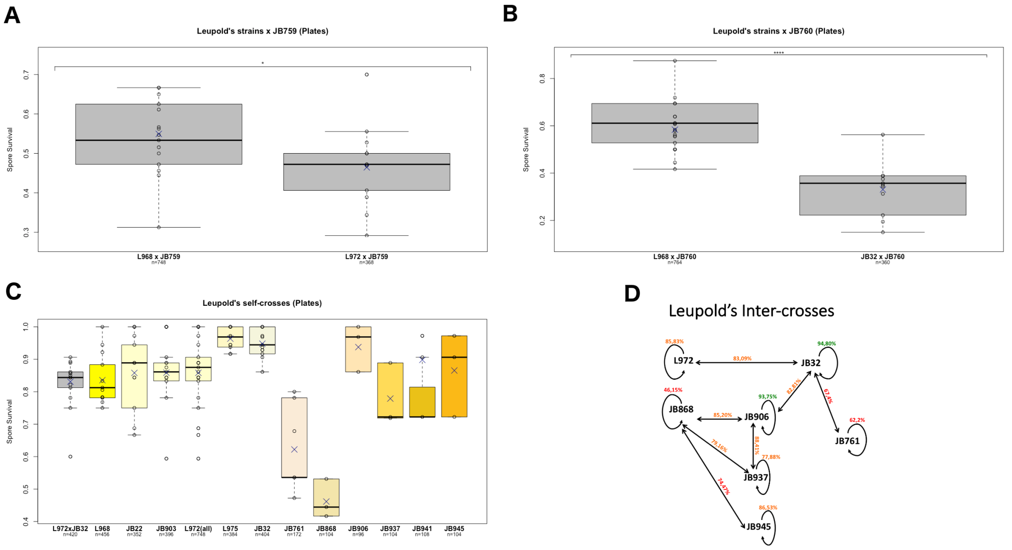

*Supplemental Figure 1:*

***Leupold's strains clonally related inter-strain crosses and Leupold's strains self-crosses.*** *Inter-strain crosses* ***(A)*** *and* ***(B)****, of Leupold's strains L968 ($h^{90}$), L972 ($h^{-s}$) and JB32 ($h^{+s}$) with non-Leopold's wild isolates JB759 ($h^{+}$)* ***(A)*** *or JB760 ($h^{-}$)* ***(B)****, accordingly. This data corresponds to the same data plotted in main Figure 1A and 1B, but with no biological replicates considered, and using the spore survival values of individually dissected plates as units of repetition (see Materials and Methods). Self-crosses* ***(C)****, of all Leupold's strains analysed{Jeffares:2015hi}. This includes canonical and newly described Leupold's derived strains. Data from canonical Leupold's isolates is the same as the data shown in main Figure 1C but plotted without considering biological replicates as in and* ***(A)*** *and* ***(B)****. The data from the newly isolated Leupold's isolates is the same as in main Figure 1D, plotted side by side with the data of the canonical Leupold's isolates, for comparison. Some one-way ANOVAs, regarding the data shown here, are available in Table S4. In* ***(D)*** *a schematic representation of all the inter-strain crosses showed in main Figure 1E is shown. Spore survival data is represented with the following colour code: crosses with spore survival significantly higher than 90% are plotted in green; crosses significantly higher than 75% are plotted in orange, while the rest of the spore survival data are plotted in red. Boxplot figure details are as indicated in the main Figure 1.*

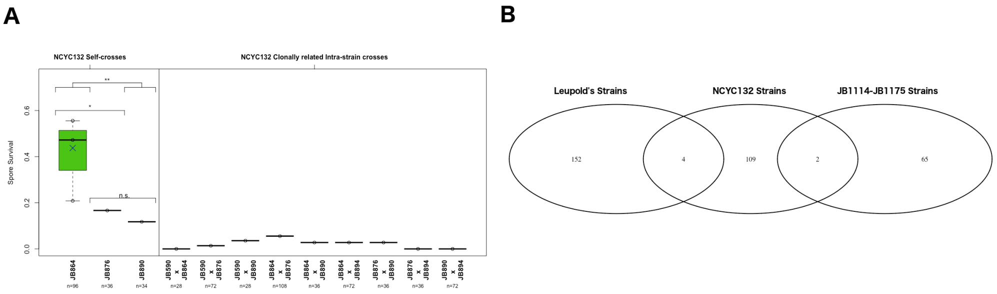

**Supplemental Figure 2:**

***Clonal group of strains: (A)*** *Self-crosses (left) and inter-strain crosses (right) of the NCYC132 clonal group of strains. Self-crosses correspond to the same data plotted in main Figure 2B, and is plotted for comparison purposes. All the data included in this graph have been analysed with a One-Way ANOVA (see Table S4).* ***(B)*** *Venn Diagram showing the polymorphic sites shared between strains analysed in the three clonal groups considered: Leupold's clonal group of strains (L972, JB32, L968, JB761, JB868, JB906, JB937, JB941 and JB945 strains), NCYC132 clonal group of strains (JB864, JB876 and JB890 strains), and the JB1114-JB1175 clonal strains.*

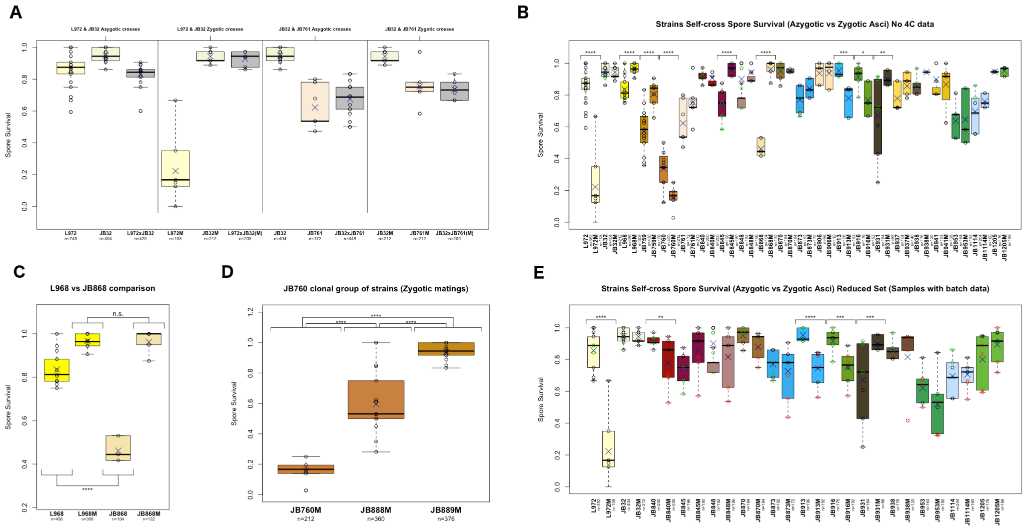

**Supplemental Figure 3:**

***Azygotic vs zygotic asci spore survival determination.*** ***(A)****Detailed comparison of the zygotic and azygotic self-crosses of the following crosses: L972 & JB32 crosses (left panels) and JB32 & JB761 crosses (right panels).* ***(B)*** *Comparison of the spore survival determinations carried out by tetrad analysis either in diploid cells (azygotic asci) or in haploid cells by mass mating experiments (zygotic asci). Same data as in main Figure 3A is shown, but 4ºC kept plates have been eliminated from the plot. A One-Way ANOVA of this data is available in Table S4.* ***(C)*** *Detailed comparison of the strains L968 and JB868. Both are $h^{90}$ Leupold's derived strains.* ***(D)*** *Comparison of zygotic self-crosses from three clonally related strain JB760, JB888 and JB889 (Table S5). In* ***(E)*** *only data from strains where batch data it was properly recorded is presented, either strains that suffered the 4ºC issue (red circles) or other strains with other batch data available (green circles). A MANOVA test with this data detects the effect of the type of meiosis (p-value=0.040635), and the 4ºC batches (p-value=5.38e-15) as significant in the spore survival determination, while no effect it is detected for the other batch information available (p-value=0.619517). Figure details for boxplots are as indicated in Figure 1. Boxplots are coloured as in previous figures. Statistically significant differences are shown (see Material and Methods and Table S2).*

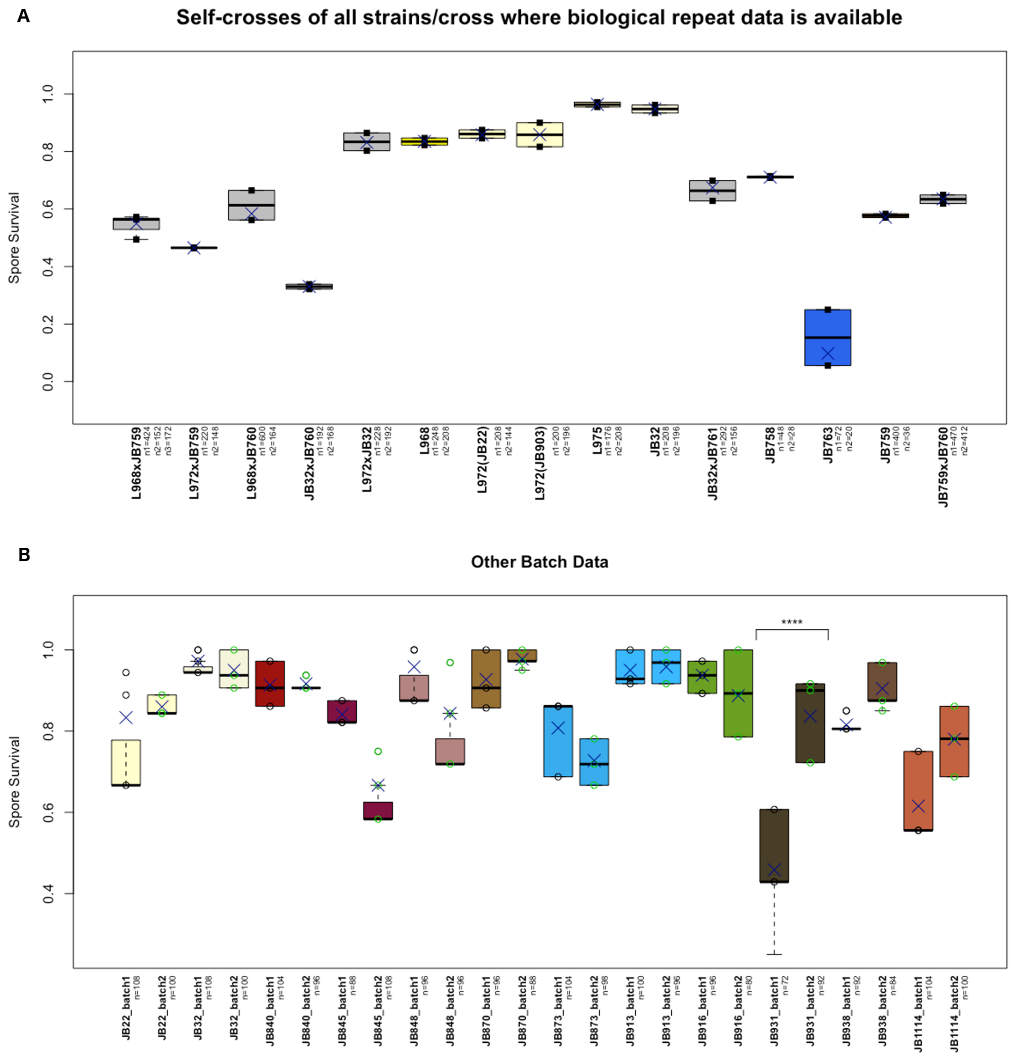

***Supplemental Figure 4:***

***Inter-strain or self-cross spore survival variation. (A)*** *Inter-strain crosses or self-crosses in which biological replicates data is available. Two or three individual diploid isolates (biological replicates) of the respective azygotic crosses were analysed by tetrad dissection. The spore survival values obtained in each of the individual isolates are represented with a closed square.* ***(B)*** *Self-cross spore survival values variability observed in strains where batch data information was recorded. Figure details are as in main Figure 3. Significance level are plotted as described in Material and Methods (see Materials and Methods, Table S2 and Table S4).*

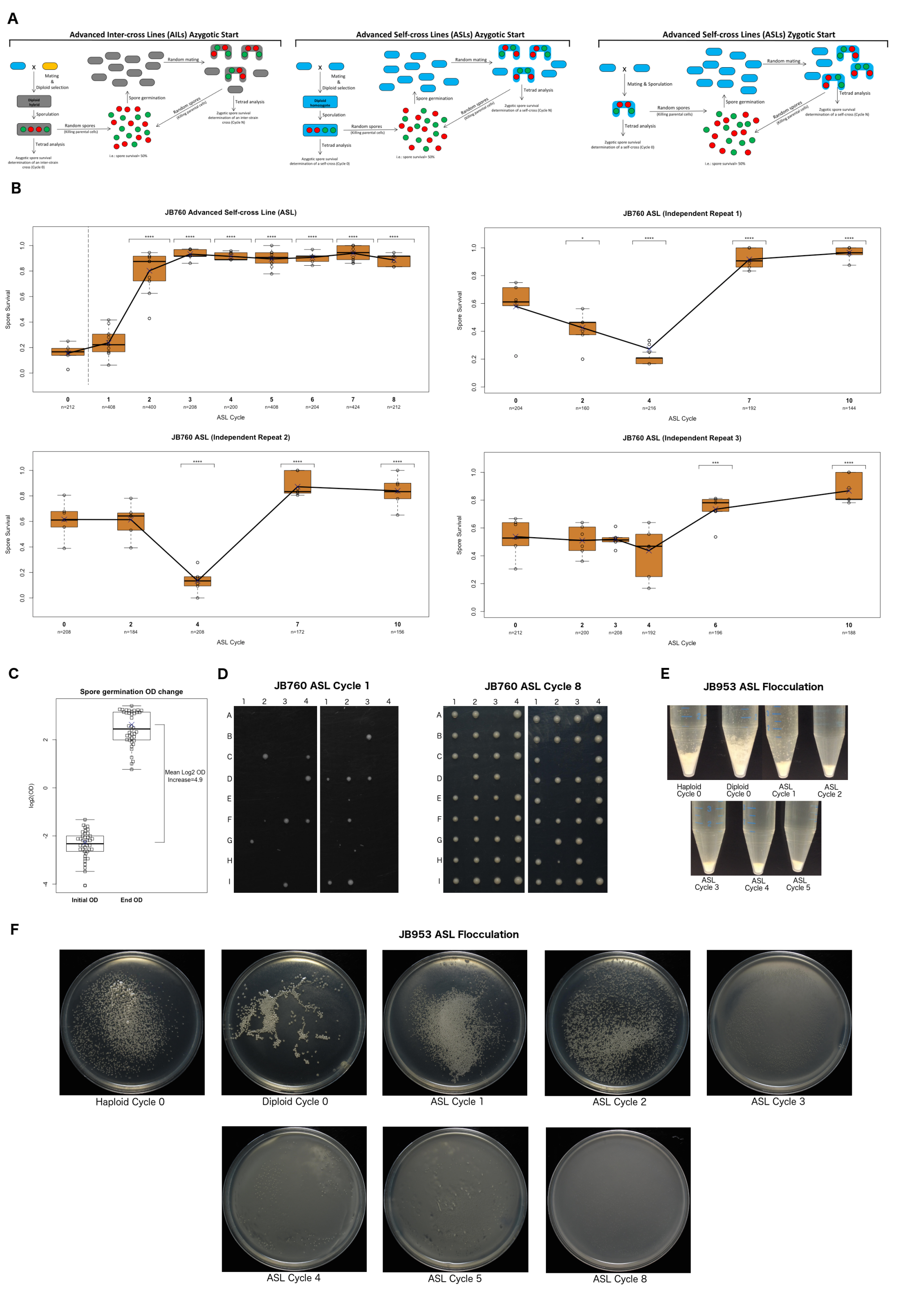

*Supplemental Figure 5:*

***Advanced Inter-Cross Lines (AILs) and Advanced Self-Cross Lines (ASLs). (A)*** *Schematic representation of the AILs azygotic start (right panel), ASLs azygotic start (middle panel) and the ASLs zygotic start (left panel). Each strain background is represented in one colour. hybrid backgrounds are represented in grey. Spores were represented in red (dead spores or non-colony forming spores) and green (live spores able to form a colony).* ***(B)*** *JB760 ASL experiment shown in main Figure 5B (top right panel), and three more independent repeats started all of them from a single isolated haploid clone (all other panels). Note than original JB760 was an azygotic started ASL (vertical dotted line is then present between cycle 0 and cycle 1, see Materials and methods and middle panel in* ***(A)****, while all other JB760 repeats are zygotic started ASLs (see right panel in* ***(A)****). Boxplot details are as in main Figure 1.* ***(C)*** *log2 representation of the Optical density (OD) variations observed in the spore germination step from the ASLs experiments performed (see materials and methods, main text and all panels in* ***(A)****). Single values are represented with open squares, while the arithmetic mean is shown with a blue X. The difference between the initial OD mean and the final OD mean is shown.* ***(D)*** *Colony size changes observed during the JB760 ASL progression. Representative dissection plates are shown.* ***(E)*** *and* ***(F)*** *phenotypic changes observed in the flocculation phenotype shown by the strain JB953 as one progress through an ASL experiment. Examples of the qualitative view of the phenotype as seen in tubes* ***(E)*** *or plates* ***(F)*** *are shown. For protocol details see Material and methods.*

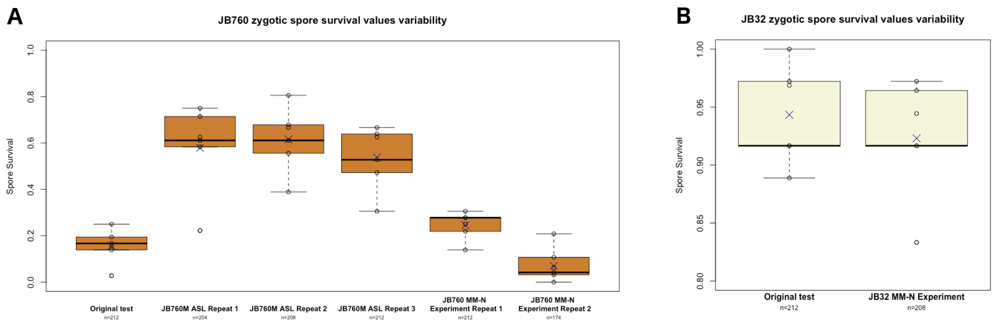

***Supplemental Figure 6:***

***Zygotic spore survival variability.*** ***(A)*** *Variability shown, during the determination of the spore survival in zygotic self-crosses, by the different haploid starting cells, used in the different JB760 related experiments. (B) Two examples of the determination of the self-cross zygotic spore survival, made from two independent clones of the haploid JB32 strain.*

#### Supplemental Table S1: Strains used in this study

| Bahler Lab strain name | Other relevant strain names | Ploidy | Genetic background | Genotype | Source | Haploid used in mass mating |
|---|---|---|---|---|---|---|
| JB759 | Y0036 | Haploid | JB759 | h+ wt | Jeffares et al 2015 | |
| JB760 | DBVPG2812 | Haploid | JB760 | h- wt | Jeffares et al 2015 | |
| JB50 | L968 | Haploid | L968 | h90 wt | Leupold 1950; Bresch et al 1968 | L968 |
| JB22 | L972 | Haploid | L972(JB22) | h-s wt | Leupold 1950; Gutz and Doe 1973; Jeffares et al | |
| JB32 | SA21 | Haploid | JB32 | h+s wt | Heim 1990 | |
| XM2 | - | Haploid | L968 | h90 Δade6::kanMX6 | This study (JB50 transformed with pΔade6-K) | |
| XM39 | - | Haploid | JB759 | h+ Δade6::natMX6 | This study (JB759 transformed with pΔAde6-N) | JB759 (XM39 x XM434) |
| XM79 | - | Haploid | L968 | h90 Δade6::hphMX6 | This study (JB50 transformed with pΔade6-H) | |
| XM191 | - | Haploid | JB759 | h90 Δade6::natMX6 | This study (JB759 spontaneous h90 revertant isolated) | |
| XM73 | - | Diploid hybrid | L968/JB759 | h+/h90 Δade6::kanMX6/Δade6::natMX6 | This study (XM2 x XM39) | |
| XM74 | - | Diploid hybrid | L968/JB759 | h+/h90 Δade6::kanMX6/Δade6::natMX6 | This study (XM2 x XM39) | |
| XM266 | - | Diploid hybrid | L968/JB759 | h90/h90 Δade6::hphMX6/Δade6::natMX6 | This study (XM79 x XM191) | |
| XM8 | - | Haploid | L972(JB22) | h-s Δade6::hphMX6 | This study (JB22 transformed with pΔade6-H) | |
| XM110 | - | Diploid hybrid | L972(JB22)/JB759 | h-s/h+ Δade6::hphMX6/Δade6::natMX6 | This study (XM8 x XM39) | |
| XM262 | - | Diploid hybrid | L972(JB22)/JB759 | h-s/h+ Δade6::hphMX6/Δade6::natMX6 | This study (XM8 x XM191) | |
| XM63 | - | Haploid | JB760 | h- Δade6::hphMX6 | This study (JB760 transformed with pΔAde6-H) | |
| XM75 | - | Diploid hybrid | L968/JB760 | h-/h90 Δade6::kanMX6/Δade6::hphMX6 | This study (XM2 x XM63) | |
| XM76 | - | Diploid hybrid | L968/JB760 | h-/h90 Δade6::kanMX6/Δade6::hphMX6 | This study (XM2 x XM63) | |
| XM1 | - | Haploid | JB32 | h+s Δade6::natMX6 | This study (JB32 transformed with pΔAde6-N) | JB32 (XM1 x XM384) |
| XM112 | - | Diploid hybrid | JB32/JB760 | h-/h+s Δade6::hphMX6/Δade6::natMX6 | This study (XM1 x XM63) | |
| XM113 | - | Diploid hybrid | JB32/JB760 | h-/h+s Δade6::hphMX6/Δade6::natMX6 | This study (XM1 x XM63) | |
| XM4 | - | Haploid | L968 | h90 Δade6::natMX6 | This study (JB50 transformed with pΔAde6-N) | |
| XM13 | - | Diploid homozygote | L968 | h90/h90 Δade6::kanMX6/Δade6::natMX6 | This study (XM2 x XM4) | |
| XM82 | - | Diploid homozygote | L968 | h90/h90 Δade6::kanMX6/Δade6::hphMX6 | This study (XM2 x XM79) | |
| XM364 | - | Haploid | L972(JB22) | h-s Δade6::natMX6 | This study (JB22 transformed with pΔade6-N) | L972-JB22 (XM383 x XM364) |
| XM383 | - | Haploid | L972(JB22) | h+(mat1P-KanMX6-ΔH1) Δade6::hphMX6 | This study (XM8 transformed with pMTS-h+) | L972-JB22 (XM383 x XM364) |
| XM381 | - | Haploid | L972(JB22) | h+(mat1P-KanMX6-ΔH1) Δade6::natMX6 | This study (XM364 transformed with pMTS-h+) | |
| XM409 | - | Diploid homozygote | L972(JB22) | h-s/h+(mat1P-KanMX6-ΔH1) Δade6::natMX6/Δade6::hphMX6 | This study (XM364 x XM383) | |
| XM410 | - | Diploid homozygote | L972(JB22) | h-s/h+(mat1P-KanMX6-ΔH1) Δade6::natMX6/Δade6::hphMX6 | This study (XM8 x XM381) | |
| JB903 | L972 | Haploid | L972(JB903) | h-s wt | Leupold 1950; Gutz and Doe 1973; Brown et al. 2011 | |
| XM641 | - | Haploid | L972(JB903) | h-s Δade6::natMX6 | This study (JB903 transformed with pΔAde6-N) | L972-JB903 (XM641 x XM647) |
| XM643 | - | Haploid | L972(JB903) | h+(mat1P-KanMX6-ΔH1) Δade6::natMX6 | This study (XM641 transformed with pMTS-h+) | |
| XM645 | - | Haploid | L972(JB903) | h-s Δade6::hphMX6 | This study (JB903 transformed with pΔAde6-H) | |
| XM647 | | Haploid | L972(JB903) | h+(mat1P-KanMX6-ΔH1) Δade6::hphMX6 | This study (XM645 transformed with pMTS-h+) | L972-JB903 (XM641 x XM647) |
| XM657 | - | Diploid homozygote | L972(JB903) | h-s/h+(mat1P-KanMX6-ΔH1) Δade6::natMX6/Δade6::hphMX6 | This study (XM641 x XM647) | |
| XM658 | - | Diploid homozygote | L972(JB903) | h-s/h+(mat1P-KanMX6-ΔH1) Δade6::natMX6/Δade6::hphMX6 | This study (XM643 x XM645) | |
| JB904 | L975 | Haploid | L975 | h+N wt | Leupold 1950; Gutz and Doe 1973; Brown et al. 2011 | |
| XM642 | - | Haploid | L975 | h+N Δade6::natMX6 | This study (JB904 transformed with pΔAde6-N) | L975 (XM642 x XM648) |
| XM644 | - | Haploid | L975 | h-(mat1M-kanMX6-ΔH1) Δade6::natMX6 | This study (XM642 transformed with pMTS-h-) | |
| XM646 | - | Haploid | L975 | h+ Δade6::hphMX6 | This study (JB904 transformed with pΔAde6-H) | |
| XM648 | - | Haploid | L975 | h-(mat1M-kanMX6-ΔH1) Δade6::hphMX6 | This study (XM646 transformed with pMTS-h-) | L975 (XM642 x XM648) |
| XM659 | - | Diploid homozygote | L975 | h-(mat1M-kanMX6-ΔH1)/h+N Δade6::natMX6/Δade6::hphMX6 | This study (XM642 x XM648) | |
| XM660 | - | Diploid homozygote | L975 | h-(mat1M-kanMX6-ΔH1)/h+N Δade6::natMX6/Δade6::hphMX6 | This study (XM644 x XM646) | |
| XM363 | - | Haploid | JB32 | h+s Δade6::hphMX6 | This study (JB32 transformed with pΔAde6-H) | |
| XM382 | - | Haploid | JB32 | h-(mat1M-kanMX6-ΔH1) Δade6::natMX6 | This study (XM1 transformed with pMTS-h-) | |
| XM384 | - | Haploid | JB32 | h-(mat1M-kanMX6-ΔH1) Δade6::hphMX6 | This study (XM363 transformed with pMTS-h-) | JB32 (XM1 x XM384) |
| XM411 | - | Diploid homozygote | JB32 | h-(mat1M-kanMX6-ΔH1)/h+N Δade6::natMX6/Δade6::hphMX6 | This study (XM1 x XM384) | |
| XM412 | - | Diploid homozygote | JB32 | h-(mat1M-kanMX6-ΔH1)/h+N Δade6::natMX6/Δade6::hphMX6 | This study (XM363 x XM382) | |
| XM126 | - | Diploid hybrid | L972(JB22)/JB32 | h-s/h+s Δade6::hphMX6/Δade6::natMX6 | This study (XM1 x XM8) | |
| XM127 | - | Diploid hybrid | L972(JB22)/JB32 | h-s/h+s Δade6::hphMX6/Δade6::natMX6 | This study (XM1 x XM8) | |
| JB761 | DBVPG6610 | Haploid | JB761 | h- wt | Brown et al. 2011 | |
| XM41 | - | Haploid | JB761 | h- Δade6::natMX6 | This Study (JB761 transformed with pΔAde6-N) | JB761 (XM41 x XM360) |
| XM64 | - | Haploid | JB761 | h- Δade6::hphMX6 | This Study (JB761 transformed with pΔAde6-H) | |
| XM360 | - | Haploid | JB761 | h+(mat1P-KanMX6-ΔH1) Δade6::hphMX6 | This study (XM64 transformed with pMTS-h+) | JB761 (XM41 x XM360) |
| XM407 | - | Diploid homozygote | JB761 | h-/h+(mat1P-KanMX6-ΔH1) Δade6::natMX6/Δade6::hphMX6 | This study (XM41 x XM360) | |
| JB868 | DBVPG6699 | Haploid | JB868 | h90 wt | Brown et al. 2011 | JB868 |
| XM437 | - | Haploid | JB868 | h90 Δade6::natMX6 | This study (JB868 transformed with pΔAde6-N) | |
| XM444 | - | Haploid | JB868 | h90 Δade6::hphMX6 | This study (JB868 transformed with pΔAde6-H) | |
| XM616 | - | Diploid homozygote | JB868 | h90/h90 Δade6::natMX6/Δade6::hphMX6 | This study (XM437 x XM444) | |
| JB906 | DBVPG6279;ATCC16979;CBS1042;CCRC21381;IFO0358 | Haploid | JB906 | h90 wt | Brown et al. 2011 | JB906 |
| XM440 | - | Haploid | JB906 | h90 Δade6::natMX6 | This study (JB906 transformed with pΔAde6-N) | |
| XM447 | - | Haploid | JB906 | h90 Δade6::hphMX6 | This study (JB906 transformed with pΔAde6-H) | |
| XM617 | - | Diploid homozygote | JB906 | h90/h90 Δade6::natMX6/Δade6::hphMX6 | This study (XM440 x XM447) | |
| JB937 | CLIB840 | Haploid | JB937 | h90 wt | Jeffares et al 2015 | JB937 |
| XM441 | - | Haploid | JB937 | h90 Δade6::natMX6 | This study (JB937 transformed with pΔAde6-N) | |
| XM448 | - | Haploid | JB937 | h90 Δade6::hphMX6 | This study (JB937 transformed with pΔAde6-H) | |
| XM618 | - | Diploid homozygote | JB937 | h90/h90 Δade6::natMX6/Δade6::hphMX6 | This study (XM441 XxXM448) | |
| JB941 | CLIB844 | Haploid | JB941 | h90 wt | Jeffares et al 2015 | JB941 |
| XM442 | - | Haploid | JB941 | h90 Δade6::natMX6 | This study (JB941 transformed with pΔAde6-N) | |
| XM449 | - | Haploid | JB941 | h90 Δade6::hphMX6 | This study (JB941 transformed with pΔAde6-H) | |
| XM619 | - | Diploid homozygote | JB941 | h90/h90 Δade6::natMX6/Δade6::hphMX6 | This study (XM442 x XM449) | |
| JB945 | CBS1042;ATCC16979;CCRC21381;DBVPG6279;IFO0358 | Haploid | JB945 | h90 wt | Brown et al. 2011 | |
| XM443 | - | Haploid | JB945 | h90 Δade6::natMX6 | This study (JB945 transformed with pΔAde6-N) | |
| XM450 | - | Haploid | JB945 | h90 Δade6::hphMX6 | This study (JB945 transformed with pΔAde6-H) | |
| XM620 | - | Diploid homozygote | JB945 | h90/h90 Δade6::natMX6/Δade6::hphMX6 | This study (XM443 x XM450) | |
| XM114 | - | Diploid hybrid | JB32/JB761 | h-/h+s Δade6::hphMX6/Δade6::natMX6 | This study (XM1 x XM64) | |
| XM115 | - | Diploid hybrid | JB32/JB761 | h-/h+s Δade6::hphMX6/Δade6::natMX6 | This study (XM1 x XM64) | |
| XM681 | - | Diploid hybrid | JB32/JB906 | h+s/h90 Δade6::natMX6/Δade6::hphMX6 | This study (XM1 X XM447) | |
| XM451 | - | Diploid hybrid | JB868/JB906 | h90/h90 Δade6::natMX6/Δade6::hphMX6 | This study (XM437 x XM447) | |
| XM452 | - | Diploid hybrid | JB868/JB937 | h90/h90 Δade6::natMX6/Δade6::hphMX6 | This study (XM437 x XM448) | |
| XM453 | - | Diploid hybrid | JB868/JB945 | h90/h90 Δade6::natMX6/Δade6::hphMX6 | This study (XM437 x XM450) | |
| XM454 | - | Diploid hybrid | JB906/JB937 | h90/h90 Δade6::natMX6/Δade6::hphMX6 | This study (XM440 x XM448) | |
| JB758 | NCYC936;CBS10394 | Haploid | JB758 | h+ wt | Brown et al. 2011 | |
| XM38 | - | Haploid | JB758 | h+ Δade6::natMX6 | This study (JB758 transformed with pΔAde6-N) | |
| XM61 | - | Haploid | JB758 | h+ Δade6::hphMX6 | This study (JB758 transformed with pΔAde6-H) | |
| XM190 | - | Haploid | JB758 | h- Δade6::hphMX6 | This study (JB758 spontaneus mating type revertant isolated) | |
| XM120 | - | Diploid homozygote | JB758 | h+/h- Δade6::hphMX6/Δade6::natMX6 | This study (XM38 x XM61) | |
| XM268 | - | Diploid homozygote | JB758 | h+/h- Δade6::hphMX6/Δade6::natMX6 | This study (XM38 x XM190) | |
| JB864 | NCYC132;ATCC24751;CBS10391 | Haploid | NCYC132(JB864) | h90 wt | Brown et al. 2011 | |
| XM323 | - | Haploid | NCYC132(JB864) | h90 Δade6::natMX6 | This study (JB864 transformed with pΔAde6-N) | |
| XM328 | - | Haploid | NCYC132(JB864) | h90 Δade6::hphMX6 | This study (JB864 transformed with pΔAde6-H) | |
| XM338 | - | Diploid homozygote | NCYC132(JB864) | h90/h90 Δade6::natMX6/Δade6::hphMX6 | This study (XM323 x XM328) | |
| JB916 | CBS352;DBVPG6373 | Haploid | JB916 | h90 wt | Brown et al. 2011 | |
| XM469 | - | Haploid | JB916 | h90 Δade6::natMX6 | This study (JB916 transformed with pΔAde6-N) | JB916 |
| XM491 | - | Haploid | JB916 | h90 Δade6::hphMX6 | This study (JB916 transformed with pΔAde6-H) | |
| XM595 | - | Diploid homozygote | JB916 | h90/h90 Δade6::natMX6/Δade6::hphMX6 | This study (XM469 x XM491) | |
| JB953 | PHAFF65-116 | Haploid | JB953 | h90 wt | Jeffares et al 2015 | |
| XM637 | - | Haploid | JB953 | h90 Δade6::natMX6 | This study (JB953 transformed with pΔAde6-N) | JB953 |
| XM638 | - | Haploid | JB953 | h90 Δade6::hphMX6 | This study (JB953 transformed with pΔAde6-H) | |
| XM603 | - | Diploid homozygote | JB953 | h90/h90 Δade6::natMX6/Δade6::hphMX6 | This study (XM637 x XM638) | |
| JB1205 | NBRC10568 | Haploid | JB1205 | h90 wt | Jeffares et al 2015 | |
| XM478 | - | Haploid | JB1205 | h90 Δade6::natMX6 | This study (JB1205 transformed with pΔAde6-N) | JB1205 |
| XM500 | - | Haploid | JB1205 | h90 Δade6::hphMX6 | This study (JB1205 transformed with pΔAde6-H) | |
| XM604 | - | Diploid homozygote | JB1205 | h90/h90 Δade6::natMX6/Δade6::hphMX6 | This study (XM478 x XM500) | |
| JB1206 | NBRC10569 | Haploid | JB1206 | h90 wt | Jeffares et al 2015 | |
| XM479 | - | Haploid | JB1206 | h90 Δade6::natMX6 | This study (JB1206 transformed with pΔAde6-N) | |
| XM501 | - | Haploid | JB1206 | h90 Δade6::hphMX6 | This study (JB1206 transformed with pΔAde6-H) | |
| XM605 | - | Diploid homozygote | JB1206 | h90/h90 Δade6::natMX6/Δade6::hphMX6 | This study (XM479 x XM501) | |
| JB840 | UFMGR435;CBS10458 | Haploid | JB840 | h90 wt | Brown et al. 2011 | |
| XM459 | - | Haploid | JB840 | h90 Δade6::natMX6 | This study (JB840 transformed with pΔAde6-N) | JB840 |
| XM481 | - | Haploid | JB840 | h90 Δade6::hphMX6 | This study (JB840 transformed with pΔAde6-H) | |
| XM585 | - | Diploid homozygote | JB840 | h90/h90 Δade6::natMX6/Δade6::hphMX6 | This study (XM459 x XM481) | |
| JB845 | UFMGR434;CBS10476 | Haploid | JB845 | h90 wt | Brown et al. 2011 | |
| XM460 | - | Haploid | JB845 | h90 Δade6::natMX6 | This study (JB845 transformed with pΔAde6-N) | JB845 |
| XM482 | - | Haploid | JB845 | h90 Δade6::hphMX6 | This study (JB845 transformed with pΔAde6-H) | |
| XM586 | - | Diploid homozygote | JB845 | h90/h90 Δade6::natMX6/Δade6::hphMX6 | This study (XM460 x XM482) | |
| JB848 | UFMGR428;CBS10475 | Haploid | JB848 | h90 wt | Brown et al. 2011 | |
| XM461 | - | Haploid | JB848 | h90 Δade6::natMX6 | This study (JB848 transformed with pΔAde6-N) | JB848 |
| XM483 | - | Haploid | JB848 | h90 Δade6::hphMX6 | This study (JB848 transformed with pΔAde6-H) | |
| XM587 | - | Diploid homozygote | JB848 | h90/h90 Δade6::natMX6/Δade6::hphMX6 | This study (XM461 x XM483) | |
| JB853 | UFMGA1000;CBS10465 | Haploid | JB853 | h90 wt | Brown et al. 2011 | |
| XM462 | - | Haploid | JB853 | h90 Δade6::natMX6 | This study (JB853 transformed with pΔAde6-N) | |
| XM484 | - | Haploid | JB853 | h90 Δade6::hphMX6 | This study (JB853 transformed with pΔAde6-H) | |
| XM588 | - | Diploid homozygote | JB853 | h90/h90 Δade6::natMX6/Δade6::hphMX6 | This study (XM462 x XM484) | |
| JB594 | CBS5682;DBVPG6376;NCYC3421 | Haploid | JB594 | h90 wt | Brown et al. 2011 | |
| XM7 | - | Haploid | JB594 | h90 Δade6::natMX6 | This study (JB594 transformed with pΔAde6-N) | |
| XM11 | - | Haploid | JB594 | h90 Δade6::hphMX6 | This study (JB594 transformed with pΔAde6-H) | |
| XM15 | - | Diploid homozygote | JB594 | h90/h90 Δade6::natMX6/Δade6::hphMX6 | This study (XM7 x XM11) | |
| JB763 | CBS357;CLIB834;DBVPG6280 | Haploid | JB763 | h- wt | Brown et al. 2011 | |
| XM59 | - | Haploid | JB763 | h- Δade6::natMX6 | This study (JB763 transformed with pΔAde6-N) | |
| XM66 | - | Haploid | JB763 | h- Δade6::hphMX6 | This study (JB763 transformed with pΔAde6-H) | |
| XM192 | - | Haploid | JB763 | h90 Δade6::natMX6 | This study (JB763 spontaneous mating type revertants isolated) | |
| XM124 | - | Diploid homozygote | JB763 | h90/h- Δade6::natMX6/Δade6::hphMX6 | This study (XM59 x XM66) | |
| XM272 | - | Diploid homozygote | JB763 | h90/h- Δade6::natMX6/Δade6::hphMX6 | This study (XM66 x XM192) | |

| | | | | | | |
|---|---|---|---|---|---|---|
| JB838 | UWOPS94.422.2 | Haploid | JB838 | h90 wt | Brown et al. 2011 | |
| XM458 | - | Haploid | JB838 | h90 Δade6::natMX6 | This study (JB838 transformed with pΔAde6-N) | |
| XM480 | - | Haploid | JB838 | h90 Δade6::hphMX6 | This study (JB838 transformed with pΔAde6-H) | |
| XM584 | - | Diploid homozygote | JB838 | h90/h90 Δade6::natMX6/Δade6::hphMX6 | This study (XM458 XxXM480) | |
| JB872 | CBS2775;IFO0347;NBRC0347;NCYC3418 | Haploid | JB872 | h90 wt | Brown et al. 2011 | |
| XM464 | - | Haploid | JB872 | h90 Δade6::natMX6 | This study (JB872 transformed with pΔAde6-N) | |
| XM872 | - | Haploid | JB872 | h90 Δade6::hphMX6 | This study (JB872 transformed with pΔAde6-H) | |
| XM590 | - | Diploid homozygote | JB872 | h90/h90 Δade6::natMX6/Δade6::hphMX6 | This study (XM464 x XM872) | |
| JB873 | CBS5680;DBVPG6448;NCYC3422;EF5 | Haploid | JB873 | h90 wt | Brown et al. 2011 | |
| XM465 | - | Haploid | JB873 | h90 Δade6::natMX6 | This study (JB873 transformed with pΔAde6-N) | JB873 |
| XM487 | - | Haploid | JB873 | h90 Δade6::hphMX6 | This study (JB873 transformed with pΔAde6-H) | |
| XM591 | - | Diploid homozygote | JB873 | h90/h90 Δade6::natMX6/Δade6::hphMX6 | This study (XM465 x XM487) | |
| JB884 | DBVPG2810 | Haploid | JB884 | h90 wt | Brown et al. 2011 | |
| XM467 | - | Haploid | JB884 | h90 Δade6::natMX6 | This study (JB884 transformed with pΔAde6-N) | |
| XM489 | - | Haploid | JB884 | h90 Δade6::hphMX6 | This study (JB884 transformed with pΔAde6-H) | |
| XM593 | - | Diploid homozygote | JB884 | h90/h90 Δade6::natMX6/Δade6::hphMX6 | This study (XM467 x XM489) | |
| JB913 | CBS1058 | Haploid | JB913 | h90 wt | Brown et al. 2011 | |
| XM468 | - | Haploid | JB913 | h90 Δade6::natMX6 | This study (JB913 transformed with pΔAde6-N) | JB913 |
| XM490 | - | Haploid | JB913 | h90 Δade6::hphMX6 | This study (JB913 transformed with pΔAde6-H) | |
| XM594 | - | Diploid homozygote | JB913 | h90/h90 Δade6::natMX6/Δade6::hphMX6 | This study (XM468 x XM490) | |
| JB1114 | YHL278;CSIRY-830 | Haploid | JB1114 | h90 wt | Jeffares et al 2015 | |
| XM472 | - | Haploid | JB1114 | h90 Δade6::natMX6 | This study (JB1114 transformed with pΔAde6-N) | JB1114 |
| XM494 | - | Haploid | JB1114 | h90 Δade6::hphMX6 | This study (JB1114 transformed with pΔAde6-H) | |
| XM598 | - | Diploid homozygote | JB1114 | h90/h90 Δade6::natMX6/Δade6::hphMX6 | This study (XM472 x XM494) | |
| JB1180 | Kambucha; YFS276; SPK1820 | Haploid | Kambucha(JB1180) | h90 wt | Jeffares et al 2015 | |
| XM476 | - | Haploid | Kambucha(JB1180) | h90 Δade6::natMX6 | This study (JB1180 transformed with pΔAde6-N) | |
| XM498 | - | Haploid | Kambucha(JB1180) | h90 Δade6::hphMX6 | This study (JB1180 transformed with pΔAde6-H) | |
| XM602 | - | Diploid homozygote | Kambucha(JB1180) | h90/h90 Δade6::natMX6/Δade6::hphMX6 | This study (XM476 x XM498) | |
| XM62 | - | Haploid | JB759 | h+ Δade6::hphMX6 | This study (JB759 transformed with pΔAde6-H) | |
| XM122 | - | Diploid homozygote | JB759 | h+/h90 Δade6::hphMX6/Δade6::natMX6 | This study (XM39 x XM62) | |
| XM270 | - | Diploid homozygote | JB759 | h+/h90 Δade6::hphMX6/Δade6::natMX6 | This study (XM62 x XM191) | |
| XM40 | - | Haploid | JB760 | h- Δade6::natMX6 | This study (JB760 transformed with pΔAde6-N) | JB760 (XM40 x XM359) |
| XM359 | - | Haploid | JB760 | h+(mat1P-KanMX6-ΔH1) Δade6::hphMX6 | This study (XM63 transformed with pMTS-h+) | |
| XM405 | - | Diploid homozygote | JB760 | h-/h+(mat1P-KanMX6-ΔH1) Δade6::natMX6/Δade6::hphMX6 | This study (XM40 x XM359) | |
| JB870 | Y0037;CBS10499 | Haploid | JB870 | h90 wt | Brown et al. 2011 | |
| XM463 | - | Haploid | JB870 | h90 Δade6::natMX6 | This study (JB870 transformed with pΔAde6-N) | JB870 |
| XM485 | - | Haploid | JB870 | h90 Δade6::hphMX6 | This study (JB870 transformed with pΔAde6-H) | |
| XM589 | - | Diploid homozygote | JB870 | h90/h90 Δade6::natMX6/Δade6::hphMX6 | This study (XM463 x XM485) | |
| JB878 | DBVPG2804 | Haploid | JB878 | h90 wt | Brown et al. 2011 | |
| XM466 | - | Haploid | JB878 | h90 Δade6::natMX6 | This study (JB878 transformed with pΔAde6-N) | |
| XM488 | - | Haploid | JB878 | h90 Δade6::hphMX6 | This study (JB878 transformed with pΔAde6-H) | |
| XM592 | - | Diploid homozygote | JB878 | h90/h90 Δade6::natMX6/Δade6::hphMX6 | This study (XM466 x XM488) | |
| JB931 | FRR2535 | Haploid | JB931 | h90 wt | Jeffares et al 2015 | |
| XM470 | - | Haploid | JB931 | h90 Δade6::natMX6 | This study (JB931 transformed with pΔAde6-N) | JB931 |
| XM492 | - | Haploid | JB931 | h90 Δade6::hphMX6 | This study (JB931 transformed with pΔAde6-H) | |
| XM596 | - | Diploid homozygote | JB931 | h90/h90 Δade6::natMX6/Δade6::hphMX6 | This study (XM470 x XM492) | |
| JB938 | CLIB841 | Haploid | JB938 | h90 wt | Jeffares et al 2015 | |
| XM471 | - | Haploid | JB938 | h90 Δade6::natMX6 | This study (JB938 transformed with pΔAde6-N) | JB938 |
| XM493 | - | Haploid | JB938 | h90 Δade6::hphMX6 | This study (JB938 transformed with pΔAde6-H) | |
| XM597 | - | Diploid homozygote | JB938 | h90/h90 Δade6::natMX6/Δade6::hphMX6 | This study (XM471 x XM493) | |
| JB1174 | CECT12918;IFI2139 | Haploid | JB1174 | h90 wt | Jeffares et al 2015 | |
| XM474 | - | Haploid | JB1174 | h90 Δade6::natMX6 | This study (JB1174 transformed with pΔAde6-N) | |
| XM496 | - | Haploid | JB1174 | h90 Δade6::hphMX6 | This study (JB1174 transformed with pΔAde6-H) | |
| XM600 | - | Diploid homozygote | JB1174 | h90/h90 Δade6::natMX6/Δade6::hphMX6 | This study (XM474 x XM496) | |
| JB1154 | AWRI1875 | Haploid | JB1154 | h90 wt | Jeffares et al 2015 | |
| XM473 | - | Haploid | JB1154 | h90 Δade6::natMX6 | This study (JB1154 transformed with pΔAde6-N) | |
| XM495 | - | Haploid | JB1154 | h90 Δade6::hphMX6 | This study (JB1154 transformed with pΔAde6-H) | |
| XM599 | - | Diploid homozygote | JB1154 | h90/h90 Δade6::natMX6/Δade6::hphMX6 | This study (XM473 x XM495) | |
| JB876 | NCYC535;ATCC16491;CBS4100 | Haploid | JB876 | h90 wt | Brown et al. 2011 | |
| XM324 | - | Haploid | JB876 | h90 Δade6::natMX6 | This study (JB876 transformed with pΔAde6-N) | |
| XM329 | - | Haploid | JB876 | h90 Δade6::hphMX6 | This study (JB876 transformed with pΔAde6-H) | |
| XM344 | - | Diploid homozygote | JB876 | h90/h90 Δade6::natMX6/Δade6::hphMX6 | This study (XM324 x XM329) | |
| JB890 | DBVPG2817 | Haploid | JB890 | h90 wt | Brown et al. 2011 | |
| XM325 | - | Haploid | JB890 | h90 Δade6::natMX6 | This study (JB890 transformed with pΔAde6-N) | |
| XM330 | - | Haploid | JB890 | h90 Δade6::hphMX6 | This study (JB890 transformed with pΔAde6-H) | |
| XM350 | - | Diploid homozygote | JB890 | h90/h90 Δade6::natMX6/Δade6::hphMX6 | This study (XM325 x XM330) | |
| JB590 | CBS356;DBVPG6277;JCM8274;JB861 | Haploid | JB590 | h90 wt | Brown et al. 2011 | |
| XM322 | - | Haploid | JB590 | h90 Δade6::natMX6 | This study (JB590 transformed with pΔAde6-N) | |
| XM327 | - | Haploid | JB590 | h90 Δade6::hphMX6 | This study (JB590 transformed with pΔAde6-H) | |
| JB890 | DBVPG2817 | Haploid | JB890 | h90 wt | Brown et al. 2011 | |
| XM325 | - | Haploid | JB890 | h90 Δade6::natMX6 | This study (JB890 transformed with pΔAde6-N) | |
| XM330 | - | Haploid | JB890 | h90 Δade6::hphMX6 | This study (JB890 transformed with pΔAde6-H) | |
| JB894 | DBVPG4437 | Haploid | JB894 | h90 wt | Brown et al. 2011 | |
| XM326 | - | Haploid | JB894 | h90 Δade6::natMX6 | This study (JB894 transformed with pΔAde6-N) | |
| XM331 | - | Haploid | JB894 | h90 Δade6::hphMX6 | This study (JB894 transformed with pΔAde6-H) | |
| XM333 | - | Diploid hybrid | JB590/JB864 | h90/h90 Δade6::natMX6/Δade6::hphMX6 | This study (XM322 x XM328) | |
| XM334 | - | Diploid hybrid | JB590/JB876 | h90/h90 Δade6::natMX6/Δade6::hphMX6 | This study (XM322 x XM329) | |
| XM335 | - | Diploid hybrid | JB590/JB890 | h90/h90 Δade6::natMX6/Δade6::hphMX6 | This study (XM322 x XM330) | |
| XM339 | - | Diploid hybrid | JB864/JB876 | h90/h90 Δade6::natMX6/Δade6::hphMX6 | This study (XM323 x XM329) | |
| XM340 | - | Diploid hybrid | JB864/JB890 | h90/h90 Δade6::natMX6/Δade6::hphMX6 | This study (XM323 x XM330) | |
| XM341 | - | Diploid hybrid | JB864/JB894 | h90/h90 Δade6::natMX6/Δade6::hphMX6 | This study (XM323 x XM331) | |
| XM345 | - | Diploid hybrid | JB876/JB890 | h90/h90 Δade6::natMX6/Δade6::hphMX6 | This study (XM324 x XM330) | |
| XM346 | - | Diploid hybrid | JB876/JB894 | h90/h90 Δade6::natMX6/Δade6::hphMX6 | This study (XM324 x XM331) | |
| XM351 | - | Diploid hybrid | JB890/JB894 | h90/h90 Δade6::natMX6/Δade6::hphMX6 | This study (XM325 x XM331) | |
| JB1175 | CECT12926 | Haploid | JB1175 | h90 wt | Jeffares et al 2015 | |
| XM475 | - | Haploid | JB1175 | h90 Δade6::natMX6 | This study (JB1154 transformed with pΔAde6-N) | |
| XM497 | - | Haploid | JB1175 | h90 Δade6::hphMX6 | This study (JB1154 transformed with pΔAde6-H) | |
| XM601 | - | Diploid homozygote | JB1175 | h90/h90 Δade6::natMX6/Δade6::hphMX6 | This study (XM475 x XM497) | |
| XM434 | - | Haploid | JB759 | h-(mat1M-kanMX6-ΔH1) Δade6::hphMX6 | This study (XM62 transformed with pMTS-h-) | JB759 (XM39 x XM434) |
| XM773 | Xavi Marsellach T14-1; Figure 3C Spore 1 | Haploid | Recombinant (JB22/JB32) | h+N Δade6::hphMX6 | This study (From tetrad disection of XM126 diploid hybrid) | XM773 (XM773 x XM699) |
| XM774 | Xavi Marsellach T14-2; Figure 3C Spore 2 | Haploid | Recombinant (JB22/JB32) | h+N Δade6::hphMX6 | This study (From tetrad disection of XM126 diploid hybrid) | XM774 (XM774 x XM700) |
| XM775 | Xavi Marsellach T14-3; Figure 3C Spore 3 | Haploid | Recombinant (JB22/JB32) | h-s Δade6::natMX6 | This study (From tetrad disection of XM126 diploid hybrid) | XM775 (XM775 x XM701) |
| XM776 | Xavi Marsellach T14-4; Figure 3C Spore 4 | Haploid | Recombinant (JB22/JB32) | h-s Δade6::natMX6 | This study (From tetrad disection of XM126 diploid hybrid) | XM776 (XM776 x XM702) |
| XM695 | - | Haploid | Recombinant (JB22/JB32) | h-(mat1M-kanMX6-ΔH1) Δade6::hphMX6 | This study (XM773 transformed with pMTSh-) | |
| XM696 | - | Haploid | Recombinant (JB22/JB32) | h-(mat1M-kanMX6-ΔH1) Δade6::hphMX6 | This study (XM774 transformed with pMTSh-) | |
| XM697 | - | Haploid | Recombinant (JB22/JB32) | h+(mat1M-kanMX6-ΔH1) Δade6::natMX6 | This study (XM775 transformed with pMTSh+) | |
| XM698 | - | Haploid | Recombinant (JB22/JB32) | h+(mat1M-kanMX6-ΔH1) Δade6::natMX6 | This study (XM776 transformed with pMTSh+) | |
| XM699 | - | Haploid | Recombinant (JB22/JB32) | h-(mat1M-kanMX6-ΔH1) Δade6::natMX6 | This study (XM695 transformed with pΔAde6-N) | XM773 (XM773 x XM699) |
| XM700 | - | Haploid | Recombinant (JB22/JB32) | h-(mat1M-kanMX6-ΔH1) Δade6::natMX6 | This study (XM696 transformed with pΔAde6-N) | XM774 (XM774 x XM700) |
| XM701 | - | Haploid | Recombinant (JB22/JB32) | h+(mat1M-kanMX6-ΔH1) Δade6::hphMX6 | This study (XM697 transformed with pΔAde6-H) | XM775 (XM775 x XM701) |
| XM702 | - | Haploid | Recombinant (JB22/JB32) | h+(mat1M-kanMX6-ΔH1) Δade6::hphMX6 | This study (XM698 transformed with pΔAde6-H) | XM776 (XM776 x XM702) |
| JB888 | DBVPG2815 | Haploid | JB888 | h90 wt | Brown et al. 2011 | |
| XM438 | - | Haploid | JB888 | h90 Δade6::natMX6 | This study (JB888 transformed with pΔAde6-N) | JB888 |
| XM445 | - | Haploid | JB888 | h90 Δade6::hphMX6 | This study (JB888 transformed with pΔAde6-H) | |
| JB889 | DBVPG2816 | Haploid | JB889 | h90 wt | Brown et al. 2011 | JB889 |
| XM128 | - | Diploid | JB759/JB760 | h-/h+ Δade6::hphMX6/Δade6::natMX6 | This study (XM39 x XM63) | |
| XM114 AIL Cycle 1 | - | Haploid population | JB32/JB761 | h- or h+s Δade6::hphMX6 or Δade6::natMX6 | This study (XM114 azygotic started AIL) | |
| XM114 AIL Cycle 2 | - | Haploid population | JB32/JB761 | h- or h+s Δade6::hphMX6 or Δade6::natMX6 | This study (XM114 azygotic started AIL) | |
| XM114 AIL Cycle 3 | - | Haploid population | JB32/JB761 | h- or h+s Δade6::hphMX6 or Δade6::natMX6 | This study (XM114 azygotic started AIL) | |
| XM114 AIL Cycle 4 | - | Haploid population | JB32/JB761 | h- or h+s Δade6::hphMX6 or Δade6::natMX6 | This study (XM114 azygotic started AIL) | |
| XM114 AIL Cycle 5 | - | Haploid population | JB32/JB761 | h- or h+s Δade6::hphMX6 or Δade6::natMX6 | This study (XM114 azygotic started AIL) | |
| XM114 AIL Cycle 6 | - | Haploid population | JB32/JB761 | h- or h+s Δade6::hphMX6 or Δade6::natMX6 | This study (XM114 azygotic started AIL) | |
| XM114 AIL Cycle 7 | - | Haploid population | JB32/JB761 | h- or h+s Δade6::hphMX6 or Δade6::natMX6 | This study (XM114 azygotic started AIL) | |
| XM114 AIL Cycle 8 | - | Haploid population | JB32/JB761 | h- or h+s Δade6::hphMX6 or Δade6::natMX6 | This study (XM114 azygotic started AIL) | |
| XM128 AIL Cycle 1 | - | Haploid population | JB759/JB760 | h- or h+ Δade6::hphMX6 or Δade6::natMX6 | This study (XM128 azygotic started AIL) | |
| XM128 AIL Cycle 2 | - | Haploid population | JB759/JB760 | h- or h+ Δade6::hphMX6 or Δade6::natMX6 | This study (XM128 azygotic started AIL) | |
| XM128 AIL Cycle 3 | - | Haploid population | JB759/JB760 | h- or h+ Δade6::hphMX6 or Δade6::natMX6 | This study (XM128 azygotic started AIL) | |
| XM128 AIL Cycle 4 | - | Haploid population | JB759/JB760 | h- or h+ Δade6::hphMX6 or Δade6::natMX6 | This study (XM128 azygotic started AIL) | |
| XM128 AIL Cycle 5 | - | Haploid population | JB759/JB760 | h- or h+ Δade6::hphMX6 or Δade6::natMX6 | This study (XM128 azygotic started AIL) | |
| XM128 AIL Cycle 6 | - | Haploid population | JB759/JB760 | h- or h+ Δade6::hphMX6 or Δade6::natMX6 | This study (XM128 azygotic started AIL) | |
| XM128 AIL Cycle 7 | - | Haploid population | JB759/JB760 | h- or h+ Δade6::hphMX6 or Δade6::natMX6 | This study (XM128 azygotic started AIL) | |
| XM128 AIL Cycle 8 | - | Haploid population | JB759/JB760 | h- or h+ Δade6::hphMX6 or Δade6::natMX6 | This study (XM128 azygotic started AIL) | |
| XM411 ASL Cycle 1 | - | Haploid population | JB32 | h-(mat1M-kanMX6-ΔH1) or h+N Δade6::natMX6 or Δade6::hphMX6 | This study (XM411 azygotic started ASL) | |
| XM411 ASL Cycle 2 | - | Haploid population | JB32 | h-(mat1M-kanMX6-ΔH1) or h+N Δade6::natMX6 or Δade6::hphMX6 | This study (XM411 azygotic started ASL) | |
| XM411 ASL Cycle 3 | - | Haploid population | JB32 | h-(mat1M-kanMX6-ΔH1) or h+N Δade6::natMX6 or Δade6::hphMX6 | This study (XM411 azygotic started ASL) | |
| XM411 ASL Cycle 4 | - | Haploid population | JB32 | h-(mat1M-kanMX6-ΔH1) or h+N Δade6::natMX6 or Δade6::hphMX6 | This study (XM411 azygotic started ASL) | |
| XM411 ASL Cycle 5 | - | Haploid population | JB32 | h-(mat1M-kanMX6-ΔH1) or h+N Δade6::natMX6 or Δade6::hphMX6 | This study (XM411 azygotic started ASL) | |
| XM411 ASL Cycle 6 | - | Haploid population | JB32 | h-(mat1M-kanMX6-ΔH1) or h+N Δade6::natMX6 or Δade6::hphMX6 | This study (XM411 azygotic started ASL) | |
| XM411 ASL Cycle 7 | - | Haploid population | JB32 | h-(mat1M-kanMX6-ΔH1) or h+N Δade6::natMX6 or Δade6::hphMX6 | This study (XM411 azygotic started ASL) | |
| XM411 ASL Cycle 8 | - | Haploid population | JB32 | h-(mat1M-kanMX6-ΔH1) or h+N Δade6::natMX6 or Δade6::hphMX6 | This study (XM411 azygotic started ASL) | |
| XM407 ASL Cycle 1 | - | Haploid population | JB761 | h- or h+(mat1P-KanMX6-ΔH1) Δade6::natMX6 or Δade6::hphMX6 | This study (XM407 azygotic started ASL) | |
| XM407 ASL Cycle 2 | - | Haploid population | JB761 | h- or h+(mat1P-KanMX6-ΔH1) Δade6::natMX6 or Δade6::hphMX6 | This study (XM407 azygotic started ASL) | |
| XM407 ASL Cycle 3 | - | Haploid population | JB761 | h- or h+(mat1P-KanMX6-ΔH1) Δade6::natMX6 or Δade6::hphMX6 | This study (XM407 azygotic started ASL) | |
| XM407 ASL Cycle 4 | - | Haploid population | JB761 | h- or h+(mat1P-KanMX6-ΔH1) Δade6::natMX6 or Δade6::hphMX6 | This study (XM407 azygotic started ASL) | |
| XM407 ASL Cycle 5 | - | Haploid population | JB761 | h- or h+(mat1P-KanMX6-ΔH1) Δade6::natMX6 or Δade6::hphMX6 | This study (XM407 azygotic started ASL) | |
| XM407 ASL Cycle 6 | - | Haploid population | JB761 | h- or h+(mat1P-KanMX6-ΔH1) Δade6::natMX6 or Δade6::hphMX6 | This study (XM407 azygotic started ASL) | |
| XM407 ASL Cycle 7 | - | Haploid population | JB761 | h- or h+(mat1P-KanMX6-ΔH1) Δade6::natMX6 or Δade6::hphMX6 | This study (XM407 azygotic started ASL) | |
| XM407 ASL Cycle 8 | - | Haploid population | JB761 | h- or h+(mat1P-KanMX6-ΔH1) Δade6::natMX6 or Δade6::hphMX6 | This study (XM407 azygotic started ASL) | |

| | | | | | |
|---|---|---|---|---|---|
| XM427 | - | Haploid | JB759 | h-(mat1M-kanMX6-ΔH1) Δade6::natMX6 | This study (XM39 transformed with pMTSh-) |
| XM635 | - | Diploid homozygote | JB759 | h-(mat1M-kanMX6-ΔH1)/h+ Δade6::natMX6/Δade6::hphMX6 | This study (XM62 x XM427) |
| XM635 ASL Cycle 1 | - | Haploid population | JB759 | h-(mat1M-kanMX6-ΔH1) or h+ Δade6::natMX6 or Δade6::hphMX6 | This study (XM635 azygotic started ASL) |
| XM635 ASL Cycle 2 | - | Haploid population | JB759 | h-(mat1M-kanMX6-ΔH1) or h+ Δade6::natMX6 or Δade6::hphMX6 | This study (XM635 azygotic started ASL) |
| XM635 ASL Cycle 3 | - | Haploid population | JB759 | h-(mat1M-kanMX6-ΔH1) or h+ Δade6::natMX6 or Δade6::hphMX6 | This study (XM635 azygotic started ASL) |
| XM635 ASL Cycle 4 | - | Haploid population | JB759 | h-(mat1M-kanMX6-ΔH1) or h+ Δade6::natMX6 or Δade6::hphMX6 | This study (XM635 azygotic started ASL) |
| XM635 ASL Cycle 5 | - | Haploid population | JB759 | h-(mat1M-kanMX6-ΔH1) or h+ Δade6::natMX6 or Δade6::hphMX6 | This study (XM635 azygotic started ASL) |
| XM635 ASL Cycle 6 | - | Haploid population | JB759 | h-(mat1M-kanMX6-ΔH1) or h+ Δade6::natMX6 or Δade6::hphMX6 | This study (XM635 azygotic started ASL) |
| XM635 ASL Cycle 7 | - | Haploid population | JB759 | h-(mat1M-kanMX6-ΔH1) or h+ Δade6::natMX6 or Δade6::hphMX6 | This study (XM635 azygotic started ASL) |
| XM635 ASL Cycle 8 | - | Haploid population | JB759 | h-(mat1M-kanMX6-ΔH1) or h+ Δade6::natMX6 or Δade6::hphMX6 | This study (XM635 azygotic started ASL) |
| XM405 ASL Cycle 1 | - | Haploid population | JB760 | h- or h+(mat1P-KanMX6-ΔH1) Δade6::natMX6 or Δade6::hphMX6 | This study (XM405 azygotic started ASL) |
| XM405 ASL Cycle 2 | - | Haploid population | JB760 | h- or h+(mat1P-KanMX6-ΔH1) Δade6::natMX6 or Δade6::hphMX6 | This study (XM405 azygotic started ASL) |
| XM405 ASL Cycle 3 | - | Haploid population | JB760 | h- or h+(mat1P-KanMX6-ΔH1) Δade6::natMX6 or Δade6::hphMX6 | This study (XM405 azygotic started ASL) |
| XM405 ASL Cycle 4 | - | Haploid population | JB760 | h- or h+(mat1P-KanMX6-ΔH1) Δade6::natMX6 or Δade6::hphMX6 | This study (XM405 azygotic started ASL) |
| XM405 ASL Cycle 5 | - | Haploid population | JB760 | h- or h+(mat1P-KanMX6-ΔH1) Δade6::natMX6 or Δade6::hphMX6 | This study (XM405 azygotic started ASL) |
| XM405 ASL Cycle 6 | - | Haploid population | JB760 | h- or h+(mat1P-KanMX6-ΔH1) Δade6::natMX6 or Δade6::hphMX6 | This study (XM405 azygotic started ASL) |
| XM405 ASL Cycle 7 | - | Haploid population | JB760 | h- or h+(mat1P-KanMX6-ΔH1) Δade6::natMX6 or Δade6::hphMX6 | This study (XM405 azygotic started ASL) |
| XM405 ASL Cycle 8 | - | Haploid population | JB760 | h- or h+(mat1P-KanMX6-ΔH1) Δade6::natMX6 or Δade6::hphMX6 | This study (XM405 azygotic started ASL) |
| XM711 | - | Haploid | JB760 | h- wt | This study (JB760 individualy picked clone) |
| XM712 | - | Haploid | JB760 | h- wt | This study (JB760 individualy picked clone) |
| XM713 | - | Haploid | JB760 | h- wt | This study (JB760 individualy picked clone) |
| XM731 | - | Haploid | JB760 | h+(mat1P-KanMX6-ΔH1) | This study (XM711 transformed with pMTS-h+) |
| XM732 | - | Haploid | JB760 | h+(mat1P-KanMX6-ΔH1) | This study (XM712 transformed with pMTS-h+) |
| XM733 | - | Haploid | JB760 | h+(mat1P-KanMX6-ΔH1) | This study (XM713 transformed with pMTS-h+) |
| JB760 ASL Repeat 1 Cycle 1 | - | Haploid population | JB760 | h- or h+(mat1P-KanMX6-ΔH1) | This study (XM711 x XM731 zygotic started ASL) |
| JB760 ASL Repeat 1 Cycle 2 | - | Haploid population | JB760 | h- or h+(mat1P-KanMX6-ΔH1) | This study (XM711 x XM731 zygotic started ASL) |
| JB760 ASL Repeat 1 Cycle 3 | - | Haploid population | JB760 | h- or h+(mat1P-KanMX6-ΔH1) | This study (XM711 x XM731 zygotic started ASL) |
| JB760 ASL Repeat 1 Cycle 4 | - | Haploid population | JB760 | h- or h+(mat1P-KanMX6-ΔH1) | This study (XM711 x XM731 zygotic started ASL) |
| JB760 ASL Repeat 1 Cycle 5 | - | Haploid population | JB760 | h- or h+(mat1P-KanMX6-ΔH1) | This study (XM711 x XM731 zygotic started ASL) |
| JB760 ASL Repeat 1 Cycle 6 | - | Haploid population | JB760 | h- or h+(mat1P-KanMX6-ΔH1) | This study (XM711 x XM731 zygotic started ASL) |
| JB760 ASL Repeat 1 Cycle 7 | - | Haploid population | JB760 | h- or h+(mat1P-KanMX6-ΔH1) | This study (XM711 x XM731 zygotic started ASL) |
| JB760 ASL Repeat 1 Cycle 8 | - | Haploid population | JB760 | h- or h+(mat1P-KanMX6-ΔH1) | This study (XM711 x XM731 zygotic started ASL) |
| JB760 ASL Repeat 1 Cycle 9 | - | Haploid population | JB760 | h- or h+(mat1P-KanMX6-ΔH1) | This study (XM711 x XM731 zygotic started ASL) |
| JB760 ASL Repeat 1 Cycle 10 | - | Haploid population | JB760 | h- or h+(mat1P-KanMX6-ΔH1) | This study (XM711 x XM731 zygotic started ASL) |
| JB760 ASL Repeat 2 Cycle 1 | - | Haploid population | JB760 | h- or h+(mat1P-KanMX6-ΔH1) | This study (XM712 x XM732 zygotic started ASL) |
| JB760 ASL Repeat 2 Cycle 2 | - | Haploid population | JB760 | h- or h+(mat1P-KanMX6-ΔH1) | This study (XM712 x XM732 zygotic started ASL) |
| JB760 ASL Repeat 2 Cycle 3 | - | Haploid population | JB760 | h- or h+(mat1P-KanMX6-ΔH1) | This study (XM712 x XM732 zygotic started ASL) |
| JB760 ASL Repeat 2 Cycle 4 | - | Haploid population | JB760 | h- or h+(mat1P-KanMX6-ΔH1) | This study (XM712 x XM732 zygotic started ASL) |
| JB760 ASL Repeat 2 Cycle 5 | - | Haploid population | JB760 | h- or h+(mat1P-KanMX6-ΔH1) | This study (XM712 x XM732 zygotic started ASL) |
| JB760 ASL Repeat 2 Cycle 6 | - | Haploid population | JB760 | h- or h+(mat1P-KanMX6-ΔH1) | This study (XM712 x XM732 zygotic started ASL) |
| JB760 ASL Repeat 2 Cycle 7 | - | Haploid population | JB760 | h- or h+(mat1P-KanMX6-ΔH1) | This study (XM712 x XM732 zygotic started ASL) |
| JB760 ASL Repeat 2 Cycle 8 | - | Haploid population | JB760 | h- or h+(mat1P-KanMX6-ΔH1) | This study (XM712 x XM732 zygotic started ASL) |
| JB760 ASL Repeat 2 Cycle 9 | - | Haploid population | JB760 | h- or h+(mat1P-KanMX6-ΔH1) | This study (XM712 x XM732 zygotic started ASL) |
| JB760 ASL Repeat 2 Cycle 10 | - | Haploid population | JB760 | h- or h+(mat1P-KanMX6-ΔH1) | This study (XM712 x XM732 zygotic started ASL) |
| JB760 ASL Repeat 3 Cycle 1 | - | Haploid population | JB760 | h- or h+(mat1P-KanMX6-ΔH1) | This study (XM713 x XM733 zygotic started ASL) |
| JB760 ASL Repeat 3 Cycle 2 | - | Haploid population | JB760 | h- or h+(mat1P-KanMX6-ΔH1) | This study (XM713 x XM733 zygotic started ASL) |
| JB760 ASL Repeat 3 Cycle 3 | - | Haploid population | JB760 | h- or h+(mat1P-KanMX6-ΔH1) | This study (XM713 x XM733 zygotic started ASL) |
| JB760 ASL Repeat 3 Cycle 4 | - | Haploid population | JB760 | h- or h+(mat1P-KanMX6-ΔH1) | This study (XM713 x XM733 zygotic started ASL) |
| JB760 ASL Repeat 3 Cycle 5 | - | Haploid population | JB760 | h- or h+(mat1P-KanMX6-ΔH1) | This study (XM713 x XM733 zygotic started ASL) |
| JB760 ASL Repeat 3 Cycle 6 | - | Haploid population | JB760 | h- or h+(mat1P-KanMX6-ΔH1) | This study (XM713 x XM733 zygotic started ASL) |
| JB760 ASL Repeat 3 Cycle 7 | - | Haploid population | JB760 | h- or h+(mat1P-KanMX6-ΔH1) | This study (XM713 x XM733 zygotic started ASL) |
| JB760 ASL Repeat 3 Cycle 8 | - | Haploid population | JB760 | h- or h+(mat1P-KanMX6-ΔH1) | This study (XM713 x XM733 zygotic started ASL) |
| JB760 ASL Repeat 3 Cycle 9 | - | Haploid population | JB760 | h- or h+(mat1P-KanMX6-ΔH1) | This study (XM713 x XM733 zygotic started ASL) |
| JB760 ASL Repeat 3 Cycle 10 | - | Haploid population | JB760 | h- or h+(mat1P-KanMX6-ΔH1) | This study (XM713 x XM733 zygotic started ASL) |
| XM409 ASL Cycle 1 | - | Haploid population | L972(JB22) | h-s or h+(mat1P-KanMX6-ΔH1) Δade6::natMX6 or Δade6::hphMX6 | This study (XM409 azygotic started ASL) |
| XM409 ASL Cycle 2 | - | Haploid population | L972(JB22) | h-s or h+(mat1P-KanMX6-ΔH1) Δade6::natMX6 or Δade6::hphMX6 | This study (XM409 azygotic started ASL) |
| XM409 ASL Cycle 3 | - | Haploid population | L972(JB22) | h-s or h+(mat1P-KanMX6-ΔH1) Δade6::natMX6 or Δade6::hphMX6 | This study (XM409 azygotic started ASL) |
| XM409 ASL Cycle 4 | - | Haploid population | L972(JB22) | h-s or h+(mat1P-KanMX6-ΔH1) Δade6::natMX6 or Δade6::hphMX6 | This study (XM409 azygotic started ASL) |
| XM409 ASL Cycle 5 | - | Haploid population | L972(JB22) | h-s or h+(mat1P-KanMX6-ΔH1) Δade6::natMX6 or Δade6::hphMX6 | This study (XM409 azygotic started ASL) |
| XM409 ASL Cycle 6 | - | Haploid population | L972(JB22) | h-s or h+(mat1P-KanMX6-ΔH1) Δade6::natMX6 or Δade6::hphMX6 | This study (XM409 azygotic started ASL) |
| XM409 ASL Cycle 7 | - | Haploid population | L972(JB22) | h-s or h+(mat1P-KanMX6-ΔH1) Δade6::natMX6 or Δade6::hphMX6 | This study (XM409 azygotic started ASL) |
| XM409 ASL Cycle 8 | - | Haploid population | L972(JB22) | h-s or h+(mat1P-KanMX6-ΔH1) Δade6::natMX6 or Δade6::hphMX6 | This study (XM409 azygotic started ASL) |
| XM603 ASL Cycle 1 | - | Haploid population | JB953 | h90 Δade6::natMX6 or Δade6::hphMX6 | This study (XM603 azygotic started ASL) |
| XM603 ASL Cycle 2 | - | Haploid population | JB953 | h90 Δade6::natMX6 or Δade6::hphMX6 | This study (XM603 azygotic started ASL) |
| XM603 ASL Cycle 3 | - | Haploid population | JB953 | h90 Δade6::natMX6 or Δade6::hphMX6 | This study (XM603 azygotic started ASL) |
| XM603 ASL Cycle 4 | - | Haploid population | JB953 | h90 Δade6::natMX6 or Δade6::hphMX6 | This study (XM603 azygotic started ASL) |
| XM603 ASL Cycle 5 | - | Haploid population | JB953 | h90 Δade6::natMX6 or Δade6::hphMX6 | This study (XM603 azygotic started ASL) |
| XM603 ASL Cycle 6 | - | Haploid population | JB953 | h90 Δade6::natMX6 or Δade6::hphMX6 | This study (XM603 azygotic started ASL) |
| XM603 ASL Cycle 7 | - | Haploid population | JB953 | h90 Δade6::natMX6 or Δade6::hphMX6 | This study (XM603 azygotic started ASL) |
| XM603 ASL Cycle 8 | - | Haploid population | JB953 | h90 Δade6::natMX6 or Δade6::hphMX6 | This study (XM603 azygotic started ASL) |
| XM584 ASL Cycle 1 | - | Haploid population | JB838 | h90/h90 Δade6::natMX6/Δade6::hphMX6 | This study (XM584 azygotic started ASL) |
| XM584 ASL Cycle 2 | - | Haploid population | JB838 | h90/h90 Δade6::natMX6/Δade6::hphMX6 | This study (XM584 azygotic started ASL) |
| XM584 ASL Cycle 3 | - | Haploid population | JB838 | h90/h90 Δade6::natMX6/Δade6::hphMX6 | This study (XM584 azygotic started ASL) |
| XM584 ASL Cycle 4 | - | Haploid population | JB838 | h90/h90 Δade6::natMX6/Δade6::hphMX6 | This study (XM584 azygotic started ASL) |
| XM586 ASL Cycle 1 | - | Haploid population | JB845 | h90 Δade6::natMX6 or Δade6::hphMX6 | This study (XM586 azygotic strated ASL) |
| XM586 ASL Cycle 2 | - | Haploid population | JB845 | h90 Δade6::natMX6 or Δade6::hphMX6 | This study (XM586 azygotic strated ASL) |
| XM586 ASL Cycle 3 | - | Haploid population | JB845 | h90 Δade6::natMX6 or Δade6::hphMX6 | This study (XM586 azygotic strated ASL) |
| XM586 ASL Cycle 4 | - | Haploid population | JB845 | h90 Δade6::natMX6 or Δade6::hphMX6 | This study (XM586 azygotic strated ASL) |
| XM588 ASL Cycle 1 | - | Haploid population | JB853 | h90 Δade6::natMX6 or Δade6::hphMX6 | This study (XM588 azygotic started ASL) |
| XM588 ASL Cycle 2 | - | Haploid population | JB853 | h90 Δade6::natMX6 or Δade6::hphMX6 | This study (XM588 azygotic started ASL) |
| XM588 ASL Cycle 3 | - | Haploid population | JB853 | h90 Δade6::natMX6 or Δade6::hphMX6 | This study (XM588 azygotic started ASL) |
| XM588 ASL Cycle 4 | - | Haploid population | JB853 | h90 Δade6::natMX6 or Δade6::hphMX6 | This study (XM588 azygotic started ASL) |
| XM590 ASL Cycle 1 | - | Haploid population | JB872 | h90 Δade6::natMX6 or Δade6::hphMX6 | This study (XM590 azygotic started ASL) |
| XM590 ASL Cycle 2 | - | Haploid population | JB872 | h90 Δade6::natMX6 or Δade6::hphMX6 | This study (XM590 azygotic started ASL) |
| XM590 ASL Cycle 3 | - | Haploid population | JB872 | h90 Δade6::natMX6 or Δade6::hphMX6 | This study (XM590 azygotic started ASL) |
| XM590 ASL Cycle 4 | - | Haploid population | JB872 | h90 Δade6::natMX6 or Δade6::hphMX6 | This study (XM590 azygotic started ASL) |
| XM591 ASL Cycle 1 | - | Haploid population | JB873 | h90 Δade6::natMX6 or Δade6::hphMX6 | This study (XM591 azygotic started ASL) |
| XM591 ASL Cycle 2 | - | Haploid population | JB873 | h90 Δade6::natMX6 or Δade6::hphMX6 | This study (XM591 azygotic started ASL) |
| XM591 ASL Cycle 3 | - | Haploid population | JB873 | h90 Δade6::natMX6 or Δade6::hphMX6 | This study (XM591 azygotic started ASL) |
| XM591 ASL Cycle 4 | - | Haploid population | JB873 | h90 Δade6::natMX6 or Δade6::hphMX6 | This study (XM591 azygotic started ASL) |
| XM593 ASL Cycle 1 | - | Haploid population | JB884 | h90 Δade6::natMX6 or Δade6::hphMX6 | This study (XM593 azygotic started ASL) |
| XM593 ASL Cycle 2 | - | Haploid population | JB884 | h90 Δade6::natMX6 or Δade6::hphMX6 | This study (XM593 azygotic started ASL) |
| XM593 ASL Cycle 3 | - | Haploid population | JB884 | h90 Δade6::natMX6 or Δade6::hphMX6 | This study (XM593 azygotic started ASL) |
| XM593 ASL Cycle 4 | - | Haploid population | JB884 | h90 Δade6::natMX6 or Δade6::hphMX6 | This study (XM593 azygotic started ASL) |
| XM594 ASL Cycle 1 | - | Haploid population | JB913 | h90 Δade6::natMX6 or Δade6::hphMX6 | This study (XM594 azygotic started ASL) |
| XM594 ASL Cycle 2 | - | Haploid population | JB913 | h90 Δade6::natMX6 or Δade6::hphMX6 | This study (XM594 azygotic started ASL) |
| XM594 ASL Cycle 3 | - | Haploid population | JB913 | h90 Δade6::natMX6 or Δade6::hphMX6 | This study (XM594 azygotic started ASL) |
| XM594 ASL Cycle 4 | - | Haploid population | JB913 | h90 Δade6::natMX6 or Δade6::hphMX6 | This study (XM594 azygotic started ASL) |

### Supplemental Table S2: Fisher's exact tests

| Experiment | Strain/cross 1 | Viable | Total | Viability | Strain/cross 2 | Viable | Total | Viability | Number of tests | p-value(alt="two.sided") | Bonferroni Adjusted P-value | * (<0.05) | ** (<0.01) | *** (<0.001) | **** (<0.0001) |
|---|---|---|---|---|---|---|---|---|---|---|---|---|---|---|---|
| Leupold's x JB759 or JB760 | L968xJB759 | 411 | 748 | 0,55 | L972xJB759 | 171 | 368 | 0,46 | 2 | 0,008933753 | 0,01786751 | TRUE | FALSE | FALSE | FALSE |
| (Figure 1A & 1B) | L968xJB760 | 446 | 764 | 0,58 | JB32xJB760 | 119 | 360 | 0,33 | 2 | 2,34079E-15 | 4,68E-15 | TRUE | TRUE | TRUE | TRUE |
| L975+JB32 vs L968+L972 (Figure 1C) | L975+JB32 | 753 | 788 | 0,96 | L968+L972 | 1023 | 1204 | 0,85 | 1 | 5,82822E-15 | 5,83E-15 | TRUE | TRUE | TRUE | TRUE |
| All Leupold's comparisons | L972 | 302 | 352 | 0,857954545 | JB32 | 383 | 404 | 0,948019802 | 55 | 2,81E-05 | 1,55E-03 | TRUE | TRUE | FALSE | FALSE |
| | L972 | 302 | 352 | 0,857954545 | L968 | 381 | 456 | 0,835526316 | 55 | 0,4327051 | 1 | FALSE | FALSE | FALSE | FALSE |
| | L972 | 302 | 352 | 0,857954545 | JB761 | 107 | 172 | 0,622093023 | 55 | 2,73E-09 | 1,50E-07 | TRUE | TRUE | TRUE | TRUE |
| | L972 | 302 | 352 | 0,857954545 | JB868 | 48 | 104 | 0,461538462 | 55 | 2,29E-15 | 1,26E-13 | TRUE | TRUE | TRUE | TRUE |
| | L972 | 302 | 352 | 0,857954545 | JB906 | 90 | 96 | 0,9375 | 55 | 0,0369876 | 1 | FALSE | FALSE | FALSE | FALSE |
| | L972 | 302 | 352 | 0,857954545 | JB937 | 81 | 104 | 0,778846154 | 55 | 0,06699841 | 1 | FALSE | FALSE | FALSE | FALSE |
| | L972 | 302 | 352 | 0,857954545 | JB941 | 97 | 108 | 0,898148148 | 55 | 0,3323575 | 1 | FALSE | FALSE | FALSE | FALSE |
| | L972 | 302 | 352 | 0,857954545 | JB945 | 90 | 104 | 0,865384615 | 55 | 1 | 1 | FALSE | FALSE | FALSE | FALSE |
| | L972 | 302 | 352 | 0,857954545 | JB903 | 340 | 396 | 0,858585859 | 55 | 1 | 1 | FALSE | FALSE | FALSE | FALSE |
| | L972 | 302 | 352 | 0,857954545 | JB904 | 370 | 384 | 0,963541667 | 55 | 3,20E-07 | 1,76E-05 | TRUE | TRUE | TRUE | TRUE |
| | JB32 | 383 | 404 | 0,948019802 | L968 | 381 | 456 | 0,835526316 | 55 | 1,12E-07 | 6,18E-06 | TRUE | TRUE | TRUE | TRUE |
| | JB32 | 383 | 404 | 0,948019802 | JB761 | 107 | 172 | 0,622093023 | 55 | 1,13E-21 | 6,19E-20 | TRUE | TRUE | TRUE | TRUE |
| | JB32 | 383 | 404 | 0,948019802 | JB868 | 48 | 104 | 0,461538462 | 55 | 2,14E-28 | 1,18E-26 | TRUE | TRUE | TRUE | TRUE |
| | JB32 | 383 | 404 | 0,948019802 | JB906 | 90 | 96 | 0,9375 | 55 | 0,6219522 | 1 | FALSE | FALSE | FALSE | FALSE |
| | JB32 | 383 | 404 | 0,948019802 | JB937 | 81 | 104 | 0,778846154 | 55 | 8,30E-07 | 4,57E-05 | TRUE | TRUE | TRUE | TRUE |
| | JB32 | 383 | 404 | 0,948019802 | JB941 | 97 | 108 | 0,898148148 | 55 | 0,0718251 | 1 | FALSE | FALSE | FALSE | FALSE |
| | JB32 | 383 | 404 | 0,948019802 | JB945 | 90 | 104 | 0,865384615 | 55 | 0,007481824 | 0,4115003 | FALSE | FALSE | FALSE | FALSE |
| | JB32 | 383 | 404 | 0,948019802 | JB903 | 340 | 396 | 0,858585859 | 55 | 2,04E-05 | 1,12E-03 | TRUE | TRUE | FALSE | FALSE |
| | JB32 | 383 | 404 | 0,948019802 | JB904 | 370 | 384 | 0,963541667 | 55 | 0,3051677 | 1 | FALSE | FALSE | FALSE | FALSE |
| | L968 | 381 | 456 | 0,835526316 | JB761 | 107 | 172 | 0,622093023 | 55 | 4,58E-08 | 2,52E-06 | TRUE | TRUE | TRUE | TRUE |
| | L968 | 381 | 456 | 0,835526316 | JB868 | 48 | 104 | 0,461538462 | 55 | 2,98E-14 | 1,64E-12 | TRUE | TRUE | TRUE | TRUE |
| | L968 | 381 | 456 | 0,835526316 | JB906 | 90 | 96 | 0,9375 | 55 | 0,01034199 | 0,5688093 | FALSE | FALSE | FALSE | FALSE |
| | L968 | 381 | 456 | 0,835526316 | JB937 | 81 | 104 | 0,778846154 | 55 | 0,1973827 | 1 | FALSE | FALSE | FALSE | FALSE |
| | L968 | 381 | 456 | 0,835526316 | JB941 | 97 | 108 | 0,898148148 | 55 | 0,1355268 | 1 | FALSE | FALSE | FALSE | FALSE |
| | L968 | 381 | 456 | 0,835526316 | JB945 | 90 | 104 | 0,865384615 | 55 | 0,5524013 | 1 | FALSE | FALSE | FALSE | FALSE |
| | L968 | 381 | 456 | 0,835526316 | JB903 | 340 | 396 | 0,858585859 | 55 | 0,3916956 | 1 | FALSE | FALSE | FALSE | FALSE |
| | L968 | 381 | 456 | 0,835526316 | JB904 | 370 | 384 | 0,963541667 | 55 | 4,29E-10 | 2,36E-08 | TRUE | TRUE | TRUE | TRUE |
| | JB761 | 107 | 172 | 0,622093023 | JB868 | 48 | 104 | 0,461538462 | 55 | 0,01215153 | 0,6683342 | FALSE | FALSE | FALSE | FALSE |
| | JB761 | 107 | 172 | 0,622093023 | JB906 | 90 | 96 | 0,9375 | 55 | 2,97E-09 | 1,63E-07 | TRUE | TRUE | TRUE | TRUE |
| | JB761 | 107 | 172 | 0,622093023 | JB937 | 81 | 104 | 0,778846154 | 55 | 0,007640364 | 0,42022 | FALSE | FALSE | FALSE | FALSE |
| | JB761 | 107 | 172 | 0,622093023 | JB941 | 97 | 108 | 0,898148148 | 55 | 2,04E-07 | 1,12E-05 | TRUE | TRUE | TRUE | TRUE |
| | JB761 | 107 | 172 | 0,622093023 | JB945 | 90 | 104 | 0,865384615 | 55 | 9,68E-06 | 5,33E-04 | TRUE | TRUE | TRUE | FALSE |
| | JB761 | 107 | 172 | 0,622093023 | JB903 | 340 | 396 | 0,858585859 | 55 | 1,50E-09 | 8,23E-08 | TRUE | TRUE | TRUE | TRUE |
| | JB761 | 107 | 172 | 0,622093023 | JB904 | 370 | 384 | 0,963541667 | 55 | 1,28E-24 | 7,04E-23 | TRUE | TRUE | TRUE | TRUE |
| | JB868 | 48 | 104 | 0,461538462 | JB906 | 90 | 96 | 0,9375 | 55 | 5,48E-14 | 3,01E-12 | TRUE | TRUE | TRUE | TRUE |
| | JB868 | 48 | 104 | 0,461538462 | JB937 | 81 | 104 | 0,778846154 | 55 | 3,84E-06 | 2,11E-04 | TRUE | TRUE | TRUE | FALSE |
| | JB868 | 48 | 104 | 0,461538462 | JB941 | 97 | 108 | 0,898148148 | 55 | 3,10E-12 | 1,70E-10 | TRUE | TRUE | TRUE | TRUE |
| | JB868 | 48 | 104 | 0,461538462 | JB945 | 90 | 104 | 0,865384615 | 55 | 7,31E-10 | 4,02E-08 | TRUE | TRUE | TRUE | TRUE |
| | JB868 | 48 | 104 | 0,461538462 | JB903 | 340 | 396 | 0,858585859 | 55 | 6,91E-16 | 3,80E-14 | TRUE | TRUE | TRUE | TRUE |
| | JB868 | 48 | 104 | 0,461538462 | JB904 | 370 | 384 | 0,963541667 | 55 | 2,32E-31 | 1,28E-29 | TRUE | TRUE | TRUE | TRUE |
| | JB906 | 90 | 96 | 0,9375 | JB937 | 81 | 104 | 0,778846154 | 55 | 0,002131106 | 0,1172108 | FALSE | FALSE | FALSE | FALSE |
| | JB906 | 90 | 96 | 0,9375 | JB941 | 97 | 108 | 0,898148148 | 55 | 0,4474602 | 1 | FALSE | FALSE | FALSE | FALSE |
| | JB906 | 90 | 96 | 0,9375 | JB945 | 90 | 104 | 0,865384615 | 55 | 0,1028891 | 1 | FALSE | FALSE | FALSE | FALSE |
| | JB906 | 90 | 96 | 0,9375 | JB903 | 340 | 396 | 0,858585859 | 55 | 0,03917015 | 1 | FALSE | FALSE | FALSE | FALSE |
| | JB906 | 90 | 96 | 0,9375 | JB904 | 370 | 384 | 0,963541667 | 55 | 0,2567423 | 1 | FALSE | FALSE | FALSE | FALSE |
| | JB937 | 81 | 104 | 0,778846154 | JB941 | 97 | 108 | 0,898148148 | 55 | 0,02399485 | 1 | FALSE | FALSE | FALSE | FALSE |
| | JB937 | 81 | 104 | 0,778846154 | JB945 | 90 | 104 | 0,865384615 | 55 | 0,1462474 | 1 | FALSE | FALSE | FALSE | FALSE |
| | JB937 | 81 | 104 | 0,778846154 | JB903 | 340 | 396 | 0,858585859 | 55 | 0,05089598 | 1 | FALSE | FALSE | FALSE | FALSE |
| | JB937 | 81 | 104 | 0,778846154 | JB904 | 370 | 384 | 0,963541667 | 55 | 2,03E-08 | 1,11E-06 | TRUE | TRUE | TRUE | TRUE |
| | JB941 | 97 | 108 | 0,898148148 | JB945 | 90 | 104 | 0,865384615 | 55 | 0,5260202 | 1 | FALSE | FALSE | FALSE | FALSE |
| | JB941 | 97 | 108 | 0,898148148 | JB903 | 340 | 396 | 0,858585859 | 55 | 0,3387561 | 1 | FALSE | FALSE | FALSE | FALSE |
| | JB941 | 97 | 108 | 0,898148148 | JB904 | 370 | 384 | 0,963541667 | 55 | 0,01130554 | 0,6218047 | FALSE | FALSE | FALSE | FALSE |
| | JB945 | 90 | 104 | 0,865384615 | JB903 | 340 | 396 | 0,858585859 | 55 | 1 | 1 | FALSE | FALSE | FALSE | FALSE |
| | JB945 | 90 | 104 | 0,865384615 | JB904 | 370 | 384 | 0,963541667 | 55 | 0,000495077 | 0,02722925 | TRUE | FALSE | FALSE | FALSE |
| | JB903 | 340 | 396 | 0,86 | JB904 | 370 | 384 | 0,96 | 55 | 1,58381E-07 | 8,71093E-06 | TRUE | TRUE | TRUE | TRUE |
| JB1114-JB1175 Clonal strain (Figure 2B) | JB1114 | 142 | 204 | 0,70 | JB1175 | 147 | 212 | 0,69 | 1 | 1 | 1 | FALSE | FALSE | FALSE | FALSE |
| NCYC132 Clonal group | JB864 | 42 | 96 | 0,44 | JB876 | 6 | 36 | 0,17 | 3 | 0,004294546 | 0,01288364 | TRUE | FALSE | FALSE | FALSE |
| (Figure 2B) | JB864 | 42 | 96 | 0,44 | JB890 | 4 | 34 | 0,12 | 3 | 0,000717649 | 0,002152948 | TRUE | TRUE | FALSE | FALSE |
| | JB876 | 6 | 36 | 0,17 | JB890 | 4 | 34 | 0,12 | 3 | 0,7355707 | 1 | FALSE | FALSE | FALSE | FALSE |
| Zygotic vs Azygotic meiosis | L972 | 302 | 352 | 0,857954545 | L972M | 24 | 108 | 0,222222222 | 22 | 4,93E-41 | 1,08E-39 | TRUE | TRUE | TRUE | TRUE |
| (Figure 3A) | JB32 | 383 | 404 | 0,948019802 | JB32M | 200 | 212 | 0,943396226 | 22 | 0,8513258 | 1 | FALSE | FALSE | FALSE | FALSE |
| | L968 | 381 | 456 | 0,835526316 | L968M | 298 | 308 | 0,967532468 | 22 | 1,64E-09 | 3,61E-08 | TRUE | TRUE | TRUE | TRUE |
| | JB759 | 249 | 436 | 0,571100917 | JB759M | 158 | 196 | 0,806122449 | 22 | 6,68E-09 | 1,47E-07 | TRUE | TRUE | TRUE | TRUE |
| | JB760 | 135 | 400 | 0,3375 | JB760M | 33 | 212 | 0,155660377 | 22 | 1,03E-06 | 2,27E-05 | TRUE | TRUE | TRUE | TRUE |
| | JB761 | 107 | 172 | 0,622093023 | JB761M | 160 | 212 | 0,754716981 | 22 | 0,005420559 | 0,1192523 | FALSE | FALSE | FALSE | FALSE |
| | JB840 | 183 | 200 | 0,915 | JB840M | 156 | 200 | 0,78 | 22 | 0,000249993 | 0,005499845 | TRUE | TRUE | FALSE | FALSE |
| | JB845 | 146 | 196 | 0,744897959 | JB845M | 154 | 180 | 0,855555556 | 22 | 0,009871593 | 0,217175 | FALSE | FALSE | FALSE | FALSE |
| | JB848 | 173 | 192 | 0,901041667 | JB848M | 160 | 196 | 0,816326531 | 22 | 0,01965771 | 0,4324697 | FALSE | FALSE | FALSE | FALSE |
| | JB868 | 48 | 104 | 0,461538462 | JB868M | 127 | 132 | 0,962121212 | 22 | 2,87E-19 | 6,31E-18 | TRUE | TRUE | TRUE | TRUE |
| | JB870 | 175 | 184 | 0,951086957 | JB870M | 162 | 184 | 0,880434783 | 22 | 0,02297154 | 0,505374 | FALSE | FALSE | FALSE | FALSE |
| | JB873 | 148 | 192 | 0,770833333 | JB873M | 125 | 172 | 0,726744186 | 22 | 0,3355576 | 1 | FALSE | FALSE | FALSE | FALSE |
| | JB906 | 90 | 96 | 0,9375 | JB906M | 105 | 112 | 0,9375 | 22 | 1 | 1 | FALSE | FALSE | FALSE | FALSE |
| | JB913 | 187 | 196 | 0,954081633 | JB913M | 143 | 192 | 0,744791667 | 22 | 3,45E-09 | 7,60E-08 | TRUE | TRUE | TRUE | TRUE |
| | JB916 | 161 | 176 | 0,914772727 | JB916M | 144 | 192 | 0,75 | 22 | 2,57E-05 | 0,000566085 | TRUE | TRUE | TRUE | FALSE |
| | JB931 | 110 | 164 | 0,670731707 | JB931M | 79 | 88 | 0,897727273 | 22 | 5,73784E-05 | 0,001262324 | TRUE | TRUE | FALSE | FALSE |
| | JB937 | 81 | 104 | 0,778846154 | JB937M | 120 | 140 | 0,857142857 | 22 | 0,1277644 | 1 | FALSE | FALSE | FALSE | FALSE |
| | JB938 | 151 | 176 | 0,857954545 | JB938M | 98 | 120 | 0,816666667 | 22 | 0,4182095 | 1 | FALSE | FALSE | FALSE | FALSE |
| | JB941 | 97 | 108 | 0,898148148 | JB941M | 111 | 128 | 0,8671875 | 22 | 0,5465866 | 1 | FALSE | FALSE | FALSE | FALSE |
| | JB953 | 102 | 164 | 0,62195122 | JB953M | 100 | 192 | 0,520833333 | 22 | 0,06796897 | 1 | FALSE | FALSE | FALSE | FALSE |
| | JB1114 | 142 | 204 | 0,696078431 | JB1114M | 113 | 160 | 0,70625 | 22 | 0,9082892 | 1 | FALSE | FALSE | FALSE | FALSE |
| | JB1205 | 141 | 176 | 0,80 | JB1205M | 168 | 188 | 0,89 | 22 | 0,0185908 | 0,4089976 | FALSE | FALSE | FALSE | FALSE |
| Zygotic vs Azygotic meiosis | L972 | 642 | 748 | 0,85828877 | L972M | 24 | 108 | 0,222222222 | 22 | 4,93E-41 | 1,08E-39 | TRUE | TRUE | TRUE | TRUE |
| No 4C data | JB32 | 383 | 404 | 0,948019802 | JB32M | 200 | 212 | 0,943396226 | 22 | 0,8513258 | 1 | FALSE | FALSE | FALSE | FALSE |
| (Supplemental Figure 3B) | L968 | 381 | 456 | 0,835526316 | L968M | 298 | 308 | 0,967532468 | 22 | 1,64E-09 | 3,61E-08 | TRUE | TRUE | TRUE | TRUE |
| | JB759 | 249 | 436 | 0,571100917 | JB759M | 158 | 196 | 0,806122449 | 22 | 6,68E-09 | 1,47E-07 | TRUE | TRUE | TRUE | TRUE |
| | JB760 | 135 | 400 | 0,3375 | JB760M | 33 | 212 | 0,155660377 | 22 | 1,03E-06 | 2,27E-05 | TRUE | TRUE | TRUE | TRUE |
| | JB761 | 107 | 172 | 0,622093023 | JB761M | 160 | 212 | 0,754716981 | 22 | 0,005420559 | 0,1192523 | FALSE | FALSE | FALSE | FALSE |
| | JB840 | 183 | 200 | 0,915 | JB840M_no4C | 98 | 108 | 0,907407407 | 22 | 0,8347901 | 1 | FALSE | FALSE | FALSE | FALSE |
| | JB845 | 146 | 196 | 0,744897959 | JB845M_no4C | 100 | 104 | 0,961538462 | 22 | 5,80E-07 | 1,27683E-05 | TRUE | TRUE | TRUE | TRUE |
| | JB848 | 173 | 192 | 0,901041667 | JB848M_no4C | 102 | 108 | 0,944444444 | 22 | 0,2761778 | 1 | FALSE | FALSE | FALSE | FALSE |
| | JB868 | 48 | 104 | 0,461538462 | JB868M | 127 | 132 | 0,962121212 | 22 | 2,87E-19 | 6,31E-18 | TRUE | TRUE | TRUE | TRUE |
| | JB870 | 175 | 184 | 0,951086957 | JB870M_no4C | 95 | 100 | 0,95 | 22 | 1 | 1 | FALSE | FALSE | FALSE | FALSE |
| | JB873 | 148 | 192 | 0,770833333 | JB873M_no4C | 84 | 100 | 0,84 | 22 | 0,1738167 | 1 | FALSE | FALSE | FALSE | FALSE |
| | JB906 | 90 | 96 | 0,9375 | JB906M | 105 | 112 | 0,9375 | 22 | 1 | 1 | FALSE | FALSE | FALSE | FALSE |
| | JB913 | 187 | 196 | 0,954081633 | JB913M_no4C | 78 | 100 | 0,78 | 22 | 9,91E-06 | 0,000217928 | TRUE | TRUE | TRUE | FALSE |
| | JB916 | 161 | 176 | 0,914772727 | JB916M_no4C | 80 | 104 | 0,769230769 | 22 | 0,001107037 | 0,02435482 | TRUE | FALSE | FALSE | FALSE |
| | JB931 | 110 | 164 | 0,670731707 | JB931M_no4C | 79 | 88 | 0,897727273 | 22 | 5,73784E-05 | 0,001262324 | TRUE | TRUE | FALSE | FALSE |
| | JB937 | 81 | 104 | 0,778846154 | JB937M | 120 | 140 | 0,857142857 | 22 | 0,1277644 | 1 | FALSE | FALSE | FALSE | FALSE |
| | JB938 | 151 | 176 | 0,857954545 | JB938M | 34 | 36 | 0,944444444 | 22 | 0,2690584 | 1 | FALSE | FALSE | FALSE | FALSE |
| | JB941 | 97 | 108 | 0,898148148 | JB941M | 111 | 128 | 0,8671875 | 22 | 0,5465866 | 1 | FALSE | FALSE | FALSE | FALSE |
| | JB953_no4C | 51 | 80 | 0,6375 | JB953M_no4C | 62 | 96 | 0,645833333 | 22 | 1 | 1 | FALSE | FALSE | FALSE | FALSE |
| | JB1114 | 142 | 204 | 0,696078431 | JB1114M_no4C | 73 | 96 | 0,760416667 | 22 | 0,2739464 | 1 | FALSE | FALSE | FALSE | FALSE |
| | JB1205_no4C | 53 | 56 | 0,95 | JB1205M_no4C | 99 | 104 | 0,95 | 22 | 1 | 1 | FALSE | FALSE | FALSE | FALSE |
| Zygotic vs Azygotic meiosis | L968 | 381 | 456 | 0,835526316 | JB868 | 48 | 104 | 0,461538462 | 2 | 2,98E-14 | 5,96E-14 | TRUE | TRUE | TRUE | TRUE |
| L968 vs JB868 (Supplemental Figure 3C) | L968M | 298 | 308 | 0,97 | JB868M | 127 | 132 | 0,96 | 2 | 0,7781162 | 1 | FALSE | FALSE | FALSE | FALSE |
| JB760 Clonal Group (Zygotic) | JB760M | 33 | 212 | 0,155660377 | JB888M | 214 | 360 | 0,594444444 | 3 | 5,10E-26 | 1,53E-25 | TRUE | TRUE | TRUE | TRUE |
| (Supplemental Figure 3D) | JB760M | 33 | 212 | 0,155660377 | JB889M | 353 | 376 | 0,938829787 | 3 | 2,42E-88 | 7,25E-88 | TRUE | TRUE | TRUE | TRUE |
| | JB888M | 214 | 360 | 0,59 | JB889M | 353 | 376 | 0,94 | 3 | 1,0681E-30 | 3,20431E-30 | TRUE | TRUE | TRUE | TRUE |
| Zygotic vs Azygotic meiosis | L972 | 642 | 748 | 0,86 | L972M | 24 | 108 | 0,22 | 14 | 8,84E-35 | 1,24E-33 | TRUE | TRUE | TRUE | TRUE |
| Batch data | JB32 | 383 | 404 | 0,95 | JB32M | 200 | 212 | 0,94 | 14 | 0,8513258 | 1 | FALSE | FALSE | FALSE | FALSE |
| (Supplemental Figure 3E) | JB840 | 183 | 200 | 0,92 | JB840M | 156 | 200 | 0,78 | 14 | 0,000249993 | 0,003499901 | TRUE | TRUE | FALSE | FALSE |
| | JB845 | 146 | 196 | 0,74 | JB845M | 154 | 180 | 0,86 | 14 | 0,009871593 | 0,1382023 | FALSE | FALSE | FALSE | FALSE |
| | JB848 | 173 | 192 | 0,90 | JB848M | 160 | 196 | 0,82 | 14 | 0,01965771 | 0,275208 | FALSE | FALSE | FALSE | FALSE |
| | JB870 | 175 | 184 | 0,95 | JB870M | 162 | 184 | 0,88 | 14 | 0,02297154 | 0,3216016 | FALSE | FALSE | FALSE | FALSE |
| | JB873 | 148 | 192 | 0,77 | JB873M | 125 | 172 | 0,73 | 14 | 0,3355576 | 1 | FALSE | FALSE | FALSE | FALSE |
| | JB913 | 187 | 196 | 0,95 | JB913M | 143 | 192 | 0,74 | 14 | 3,45E-09 | 4,84E-08 | TRUE | TRUE | TRUE | TRUE |
| | JB916 | 161 | 176 | 0,91 | JB916M | 144 | 192 | 0,75 | 14 | 2,57E-05 | 0,000360236 | TRUE | TRUE | TRUE | FALSE |
| | JB931 | 110 | 164 | 0,67 | JB931M | 79 | 88 | 0,90 | 14 | 5,73784E-05 | 0,000803297 | TRUE | TRUE | TRUE | FALSE |
| | JB938 | 151 | 176 | 0,86 | JB938M | 98 | 120 | 0,82 | 14 | 0,4182095 | 1 | FALSE | FALSE | FALSE | FALSE |
| | JB953 | 102 | 164 | 0,62 | JB953M | 100 | 192 | 0,52 | 14 | 0,06796897 | 0,9515655 | FALSE | FALSE | FALSE | FALSE |
| | JB1114 | 142 | 204 | 0,70 | JB1114M | 113 | 160 | 0,71 | 14 | 0,9082892 | 1 | FALSE | FALSE | FALSE | FALSE |
| | JB1205 | 141 | 176 | 0,80 | JB1205M | 168 | 188 | 0,89 | 14 | 0,0185908 | 0,2602712 | FALSE | FALSE | FALSE | FALSE |
| Biological Repeats (Azygotic) | L968xJB759 N1 | 239 | 424 | 0,563679245 | L968xJB759 N2 | 87 | 152 | 0,572368421 | 33 | 0,9240478 | 1 | FALSE | FALSE | FALSE | FALSE |
| (Supplemental Figure 4A) | L968xJB759 N1 | 239 | 424 | 0,563679245 | L968xJB759N3 | 85 | 172 | 0,494186047 | 33 | 0,1242613 | 1 | FALSE | FALSE | FALSE | FALSE |
| | L968xJB759 N2 | 87 | 152 | 0,572368421 | L968xJB759N3 | 85 | 172 | 0,494186047 | 33 | 0,1810668 | 1 | FALSE | FALSE | FALSE | FALSE |
| | L972xJB759 N1 | 102 | 220 | 0,463636364 | L972xJB759 N2 | 69 | 148 | 0,466216216 | 33 | 1 | 1 | FALSE | FALSE | FALSE | FALSE |
| | L968xJB760 N1 | 337 | 600 | 0,561666667 | L968xJB760 N2 | 109 | 164 | 0,664634146 | 33 | 0,02002529 | 0,6608346 | FALSE | FALSE | FALSE | FALSE |
| | JB32xJB760 N1 | 65 | 192 | 0,338541667 | JB32xJB760 N2 | 54 | 168 | 0,321428571 | 33 | 0,7377512 | 1 | FALSE | FALSE | FALSE | FALSE |
| | L972xJB32 N1 | 183 | 228 | 0,802631579 | L972xJB32 N2 | 166 | 192 | 0,864583333 | 33 | 0,1164482 | 1 | FALSE | FALSE | FALSE | FALSE |
| | L968 N1 | 210 | 248 | 0,846774194 | L968 N2 | 171 | 208 | 0,822115385 | 33 | 0,5266829 | 1 | FALSE | FALSE | FALSE | FALSE |
| | L972(L972) N1 | 176 | 208 | 0,846153846 | L972(L972) N2 | 126 | 144 | 0,875 | 33 | 0,5350368 | 1 | FALSE | FALSE | FALSE | FALSE |
| | L972(JB903) N1 | 180 | 200 | 0,9 | L972(JB903) N2 | 160 | 196 | 0,816326531 | 33 | 0,02066354 | 0,6818967 | FALSE | FALSE | FALSE | FALSE |
| | L975 N1 | 168 | 176 | 0,954545455 | L975 N2 | 202 | 208 | 0,971153846 | 33 | 0,4234789 | 1 | FALSE | FALSE | FALSE | FALSE |
| | JB32 N1 | 200 | 208 | 0,961538462 | JB32 N2 | 183 | 196 | 0,933673469 | 33 | 0,2634588 | 1 | FALSE | FALSE | FALSE | FALSE |
| | JB32xJB761 N1 | 204 | 292 | 0,698630137 | JB32xJB761 N2 | 98 | 156 | 0,628205128 | 33 | 0,1393231 | 1 | FALSE | FALSE | FALSE | FALSE |

| | | | | | | | | | | | | | | | |
|---|---|---|---|---|---|---|---|---|---|---|---|---|---|---|---|
| | JB758 N1 | 34 | 48 | 0,708333333 | JB758 N2 | 20 | 28 | 0,714285714 | 33 | 1 | 1 | FALSE | FALSE | FALSE | FALSE |
| | JB763 N1 | 4 | 72 | 0,055555556 | JB763 N2 | 5 | 20 | 0,25 | 33 | 0,0212596 | 0,7015669 | FALSE | FALSE | FALSE | FALSE |
| | JB759 N1 | 228 | 400 | 0,57 | JB759 N2 | 21 | 36 | 0,583333333 | 33 | 1 | 1 | FALSE | FALSE | FALSE | FALSE |
| | JB759xJB760 N1 | 305 | 470 | 0,65 | JB759xJB760 N2 | 255 | 412 | 0,62 | 33 | 0,3629969 | 1 | FALSE | FALSE | FALSE | FALSE |
| 4C Batch Effect | JB840M_no4C | 98 | 108 | 0,907407407 | JB840M_4C | 58 | 92 | 0,630434783 | 13 | 2,52E-06 | 3,27E-05 | TRUE | TRUE | TRUE | TRUE |
| (Figure 4A) | JB845M_no4C | 100 | 104 | 0,961538462 | JB845M_4C | 54 | 76 | 0,710526316 | 13 | 2,42E-06 | 3,14E-05 | TRUE | TRUE | TRUE | TRUE |
| | JB848M_no4C | 102 | 108 | 0,944444444 | JB848M_4C | 58 | 88 | 0,659090909 | 13 | 2,39E-07 | 3,10E-06 | TRUE | TRUE | TRUE | TRUE |
| | JB870M_no4C | 95 | 100 | 0,95 | JB870M_4C | 67 | 84 | 0,797619048 | 13 | 0,002370126 | 0,03081164 | TRUE | FALSE | FALSE | FALSE |
| | JB873M_no4C | 84 | 100 | 0,84 | JB873M_4C | 41 | 72 | 0,569444444 | 13 | 0,000119116 | 0,001548512 | TRUE | TRUE | FALSE | FALSE |
| | JB913M_no4C | 78 | 100 | 0,78 | JB913M_4C | 65 | 92 | 0,706521739 | 13 | 0,2518025 | 1 | FALSE | FALSE | FALSE | FALSE |
| | JB916M_no4C | 80 | 104 | 0,769230769 | JB916M_4C | 64 | 88 | 0,727272727 | 13 | 0,5092421 | 1 | FALSE | FALSE | FALSE | FALSE |
| | JB838M_no4C | 34 | 36 | 0,944444444 | JB838M_4C | 64 | 84 | 0,761904762 | 13 | 0,01985614 | 0,2581298 | FALSE | FALSE | FALSE | FALSE |
| | JB953_no4C | 51 | 80 | 0,6375 | JB953_4C | 51 | 84 | 0,607142857 | 13 | 0,748314 | 1 | FALSE | FALSE | FALSE | FALSE |
| | JB953M_no4C | 62 | 96 | 0,645833333 | JB953M_4C | 38 | 96 | 0,395833333 | 13 | 0,000845372 | 0,01098983 | TRUE | FALSE | FALSE | FALSE |
| | JB1114M_no4C | 73 | 96 | 0,760416667 | JB1114M_4C | 40 | 64 | 0,625 | 13 | 0,07749061 | 1 | FALSE | FALSE | FALSE | FALSE |
| | JB1205_no4C | 53 | 56 | 0,946428571 | JB1205_4C | 88 | 120 | 0,733333333 | 13 | 0,000917905 | 0,01193277 | TRUE | FALSE | FALSE | FALSE |
| | JB1205M_no4C | 99 | 104 | 0,95 | JB1205M_4C | 69 | 84 | 0,82 | 13 | 0,007465526 | 0,09705183 | FALSE | FALSE | FALSE | FALSE |
| Other batches | L972_batch1 | 90 | 108 | 0,833333333 | L972_batch2 | 86 | 100 | 0,86 | 12 | 0,7013312 | 1 | FALSE | FALSE | FALSE | FALSE |
| (Supplemental Figure 4B) | JB32_batch1 | 105 | 108 | 0,972222222 | JB32_batch2 | 95 | 100 | 0,95 | 12 | 0,4851409 | 1 | FALSE | FALSE | FALSE | FALSE |
| | JB840_batch1 | 95 | 104 | 0,913461538 | JB840_batch2 | 88 | 96 | 0,916666667 | 12 | 1 | 1 | FALSE | FALSE | FALSE | FALSE |
| | JB845_batch1 | 74 | 88 | 0,840909091 | JB845_batch2 | 72 | 108 | 0,666666667 | 12 | 0,008014827 | 0,09617792 | FALSE | FALSE | FALSE | FALSE |
| | JB848_batch1 | 92 | 96 | 0,958333333 | JB848_batch2 | 81 | 96 | 0,84375 | 12 | 0,01378678 | 0,1654413 | FALSE | FALSE | FALSE | FALSE |
| | JB870_batch1 | 89 | 96 | 0,927083333 | JB870_batch2 | 86 | 88 | 0,977272727 | 12 | 0,1726408 | 1 | FALSE | FALSE | FALSE | FALSE |
| | JB873_batch1 | 84 | 104 | 0,807692308 | JB873_batch2 | 64 | 88 | 0,727272727 | 12 | 0,228181 | 1 | FALSE | FALSE | FALSE | FALSE |
| | JB913_batch1 | 95 | 100 | 0,95 | JB913_batch2 | 92 | 96 | 0,958333333 | 12 | 1 | 1 | FALSE | FALSE | FALSE | FALSE |
| | JB916_batch1 | 90 | 96 | 0,9375 | JB916_batch2 | 71 | 80 | 0,8875 | 12 | 0,2840003 | 1 | FALSE | FALSE | FALSE | FALSE |
| | JB931_batch1 | 33 | 72 | 0,458333333 | JB931_batch2 | 77 | 92 | 0,836956522 | 12 | 3,61E-07 | 4,33E-06 | TRUE | TRUE | TRUE | TRUE |
| | JB938_batch1 | 75 | 92 | 0,815217391 | JB938_batch2 | 76 | 84 | 0,904761905 | 12 | 0,1292634 | 1 | FALSE | FALSE | FALSE | FALSE |
| | JB1114_batch1 | 64 | 104 | 0,62 | JB1114_batch2 | 78 | 100 | 0,78 | 12 | 0,01455006 | 0,1746007 | FALSE | FALSE | FALSE | FALSE |
| minus80C Batch Effect | L968 | 381 | 456 | 0,835526316 | L968_80C | 174 | 212 | 0,820754717 | 12 | 0,6579926 | 1 | FALSE | FALSE | FALSE | FALSE |
| (Figure 4B) | L968M | 298 | 308 | 0,967532468 | L968M_80C | 159 | 172 | 0,924418605 | 12 | 0,04405114 | 0,5286137 | FALSE | FALSE | FALSE | FALSE |
| | JB904 | 370 | 384 | 0,963541667 | JB904_80C | 172 | 180 | 0,955555556 | 12 | 0,6457989 | 1 | FALSE | FALSE | FALSE | FALSE |
| | JB594 | 43 | 48 | 0,895833333 | JB594_80C | 175 | 208 | 0,841346154 | 12 | 0,498654 | 1 | FALSE | FALSE | FALSE | FALSE |
| | JB758 | 54 | 76 | 0,710526316 | JB758_80C | 172 | 196 | 0,87755102 | 12 | 0,001878882 | 0,02254658 | TRUE | FALSE | FALSE | FALSE |
| | JB763 | 9 | 92 | 0,097826087 | JB763_80C | 9 | 188 | 0,04787234 | 12 | 0,1237061 | 1 | FALSE | FALSE | FALSE | FALSE |
| | JB864 | 42 | 96 | 0,4375 | JB864_80C. | 89 | 208 | 0,427884615 | 12 | 0,9012582 | 1 | FALSE | FALSE | FALSE | FALSE |
| | JB873 | 148 | 192 | 0,770833333 | JB873_80C | 93 | 160 | 0,58125 | 12 | 0,000211517 | 0,002538203 | TRUE | TRUE | FALSE | FALSE |
| | JB873M | 84 | 100 | 0,84 | JB873M_80C | 113 | 180 | 0,627777778 | 12 | 0,000202015 | 0,00242418 | TRUE | TRUE | FALSE | FALSE |
| | JB890 | 4 | 34 | 0,117647059 | JB890_80C | 35 | 204 | 0,171568627 | 12 | 0,6169769 | 1 | FALSE | FALSE | FALSE | FALSE |
| | JB953 | 51 | 80 | 0,6375 | JB953_80C | 104 | 216 | 0,481481481 | 12 | 0,01862493 | 0,2234991 | FALSE | FALSE | FALSE | FALSE |
| | JB953M | 62 | 96 | 0,65 | JB953M_80C | 42 | 216 | 0,19 | 12 | 1,73585E-14 | 2,08302E-13 | TRUE | TRUE | TRUE | TRUE |
| AIL JB32 x JB761 | JB32 x JB761 M | 147 | 200 | 0,735 | JB32 x JB761 M | 147 | 200 | 0,735 | 6 | 1 | 1 | FALSE | FALSE | FALSE | FALSE |
| (Figure 5A) | XM114 Cycle 1 | 209 | 260 | 0,803846154 | JB32 x JB761 M | 147 | 200 | 0,735 | 6 | 0,0917935 | 0,550761 | FALSE | FALSE | FALSE | FALSE |
| | XM114 Cycle 2 | 205 | 252 | 0,813492063 | JB32 x JB761 M | 147 | 200 | 0,735 | 6 | 0,05252033 | 0,315122 | FALSE | FALSE | FALSE | FALSE |
| | XM114 Cycle 3 | 213 | 256 | 0,83203125 | JB32 x JB761 M | 147 | 200 | 0,735 | 6 | 0,01484131 | 0,08904784 | FALSE | FALSE | FALSE | FALSE |
| | XM114 Cycle 4 | 254 | 292 | 0,869863014 | JB32 x JB761 M | 147 | 200 | 0,735 | 6 | 0,000223343 | 0,00134006 | TRUE | TRUE | FALSE | FALSE |
| | XM114 Cycle 8 | 204 | 240 | 0,85 | JB32 x JB761 M | 147 | 200 | 0,74 | 6 | 0,004063965 | 0,02438379 | TRUE | FALSE | FALSE | FALSE |
| AIL JB759 x JB760 | JB759 x JB760 M | 89 | 212 | 0,419811321 | JB759 x JB760 M | 89 | 212 | 0,419811321 | 6 | 1,00E+00 | 1,00E+00 | FALSE | FALSE | FALSE | FALSE |
| (Figure 5A) | XM128 Cycle 1 | 233 | 312 | 0,746794872 | JB759 x JB760 M | 89 | 212 | 0,419811321 | 6 | 6,52E-14 | 3,91E-13 | TRUE | TRUE | TRUE | TRUE |
| | XM128 Cycle 2 | 234 | 292 | 0,801369863 | JB759 x JB760 M | 89 | 212 | 0,419811321 | 6 | 1,24E-18 | 7,43E-18 | TRUE | TRUE | TRUE | TRUE |
| | XM128 Cycle 3 | 232 | 280 | 0,828571429 | JB759 x JB760 M | 89 | 212 | 0,419811321 | 6 | 2,74E-21 | 1,64E-20 | TRUE | TRUE | TRUE | TRUE |
| | XM128 Cycle 4 | 236 | 284 | 0,830985915 | JB759 x JB760 M | 89 | 212 | 0,419811321 | 6 | 1,07E-21 | 6,40E-21 | TRUE | TRUE | TRUE | TRUE |
| | XM128 Cycle 8 | 201 | 244 | 0,82 | JB759 x JB760 M | 89 | 212 | 0,42 | 6 | 1,71E-19 | 1,03E-18 | TRUE | TRUE | TRUE | TRUE |
| ASL JB32 | JB32M | 200 | 212 | 0,943396226 | JB32M | 200 | 212 | 0,943396226 | 9 | 1 | 1 | FALSE | FALSE | FALSE | FALSE |
| (Figure 5B) | XM411 Cycle 1 | 386 | 388 | 0,994845361 | JB32M | 200 | 212 | 0,943396226 | 9 | 0,00013199 | 0,001187913 | TRUE | TRUE | FALSE | FALSE |
| | XM411 Cycle 2 | 200 | 208 | 0,961538462 | JB32M | 200 | 212 | 0,943396226 | 9 | 0,4932462 | 1 | FALSE | FALSE | FALSE | FALSE |
| | XM411 Cycle 3 | 418 | 428 | 0,976635514 | JB32M | 200 | 212 | 0,943396226 | 9 | 0,03774081 | 0,3396672 | FALSE | FALSE | FALSE | FALSE |
| | XM411 Cycle 4 | 412 | 428 | 0,962616822 | JB32M | 200 | 212 | 0,943396226 | 9 | 0,3050734 | 1 | FALSE | FALSE | FALSE | FALSE |
| | XM411 Cycle 5 | 203 | 216 | 0,939814815 | JB32M | 200 | 212 | 0,943396226 | 9 | 1,00E+00 | 1 | FALSE | FALSE | FALSE | FALSE |
| | XM411 Cycle 6 | 202 | 212 | 0,952830189 | JB32M | 200 | 212 | 0,943396226 | 9 | 0,8272766 | 1 | FALSE | FALSE | FALSE | FALSE |
| | XM411 Cycle 7 | 188 | 208 | 0,903846154 | JB32M | 200 | 212 | 0,943396226 | 9 | 1,43E-01 | 1,00E+00 | FALSE | FALSE | FALSE | FALSE |
| | XM411 Cycle 8 | 207 | 212 | 0,98 | JB32M | 200 | 212 | 0,94 | 9 | 0,1353809 | 1 | FALSE | FALSE | FALSE | FALSE |
| ASL JB761 | JB761M | 160 | 212 | 0,754716981 | JB761M | 160 | 212 | 0,754716981 | 6 | 1 | 1 | FALSE | FALSE | FALSE | FALSE |
| (Figure 5B) | XM407 Cycle 1 | 152 | 188 | 0,808510638 | JB761M | 160 | 212 | 0,754716981 | 6 | 0,2267656 | 1 | FALSE | FALSE | FALSE | FALSE |
| | XM407 Cycle 2 | 121 | 168 | 0,720238095 | JB761M | 160 | 212 | 0,754716981 | 6 | 0,4810246 | 1 | FALSE | FALSE | FALSE | FALSE |
| | XM407 Cycle 3 | 140 | 196 | 0,714285714 | JB761M | 160 | 212 | 0,754716981 | 6 | 0,3707019 | 1 | FALSE | FALSE | FALSE | FALSE |
| | XM407 Cycle 5 | 278 | 388 | 0,716494845 | JB761M | 160 | 212 | 0,754716981 | 6 | 0,3369371 | 1 | FALSE | FALSE | FALSE | FALSE |
| | XM407 Cycle 8 | 127 | 164 | 0,77 | JB761M | 160 | 212 | 0,75 | 9 | 0,7141351 | 1 | FALSE | FALSE | FALSE | FALSE |
| ASL JB759 | JB759M | 158 | 196 | 0,806122449 | JB759M | 158 | 196 | 0,806122449 | 5 | 1,00E+00 | 1,00E+00 | FALSE | FALSE | FALSE | FALSE |
| (Figure 5B) | XM635 Cycle 1 | 166 | 208 | 0,798076923 | JB759M | 158 | 196 | 0,806122449 | 5 | 9,01E-01 | 1,00E+00 | FALSE | FALSE | FALSE | FALSE |
| | XM635 Cycle 2 | 110 | 172 | 0,639534884 | JB759M | 158 | 196 | 0,806122449 | 5 | 4,09E-04 | 2,05E-03 | TRUE | TRUE | FALSE | FALSE |
| | XM635 Cycle 6 | 189 | 212 | 0,891509434 | JB759M | 158 | 196 | 0,806122449 | 5 | 0,01809021 | 0,09045103 | FALSE | FALSE | FALSE | FALSE |
| | XM635 Cycle 8 | 203 | 212 | 0,96 | JB759M | 158 | 196 | 0,81 | 5 | 1,54E-06 | 7,72E-06 | TRUE | TRUE | TRUE | TRUE |
| ASL JB760 | JB760M | 33 | 212 | 0,155660377 | JB760M | 33 | 212 | 0,155660377 | 9 | 1,00E+00 | 1,00E+00 | FALSE | FALSE | FALSE | FALSE |
| (Figure 5B) | XM405 Cycle 1 | 97 | 408 | 0,237745098 | JB760M | 33 | 212 | 0,155660377 | 9 | 1,70E-02 | 1,53E-01 | FALSE | FALSE | FALSE | FALSE |
| | XM405 Cycle 2 | 321 | 400 | 0,8025 | JB760M | 33 | 212 | 0,155660377 | 9 | 1,95E-56 | 1,76E-55 | TRUE | TRUE | TRUE | TRUE |
| | XM405 Cycle 3 | 194 | 208 | 0,932692308 | JB760M | 33 | 212 | 0,155660377 | 9 | 4,06E-65 | 3,66E-64 | TRUE | TRUE | TRUE | TRUE |
| | XM405 Cycle 4 | 183 | 200 | 0,915 | JB760M | 33 | 212 | 0,155660377 | 9 | 4,08E-60 | 3,67E-59 | TRUE | TRUE | TRUE | TRUE |
| | XM405 Cycle 5 | 365 | 408 | 0,894607843 | JB760M | 33 | 212 | 0,155660377 | 9 | 1,04E-77 | 9,34E-77 | TRUE | TRUE | TRUE | TRUE |
| | XM405 Cycle 6 | 185 | 204 | 0,906862745 | JB760M | 33 | 212 | 0,155660377 | 9 | 3,45E-59 | 3,11E-58 | TRUE | TRUE | TRUE | TRUE |
| | XM405 Cycle 7 | 398 | 424 | 0,938679245 | JB760M | 33 | 212 | 0,155660377 | 9 | 7,97E-93 | 7,17E-92 | TRUE | TRUE | TRUE | TRUE |
| | XM405 Cycle 8 | 188 | 212 | 0,89 | JB760M | 33 | 212 | 0,16 | 9 | 2,47E-56 | 2,22E-55 | TRUE | TRUE | TRUE | TRUE |
| ASL JB60M Repeat 1 | JB760M Repeat 1 | 118 | 204 | 0,578431373 | JB760M Repeat 1 | 118 | 204 | 0,578431373 | 5 | 1 | 1 | FALSE | FALSE | FALSE | FALSE |
| (Supplemental Figure 5B) | JB760M Repeat 1 Cycle 2 | 68 | 160 | 0,425 | JB760M Repeat 1 | 118 | 204 | 0,578431373 | 5 | 0,004329193 | 0,02164596 | TRUE | FALSE | FALSE | FALSE |
| | JB760M Repeat 1 Cycle 4 | 59 | 216 | 0,273148148 | JB760M Repeat 1 | 118 | 204 | 0,578431373 | 5 | 3,35E-10 | 1,67E-09 | TRUE | TRUE | TRUE | TRUE |
| | JB760M Repeat 1 Cycle 7 | 176 | 192 | 0,916666667 | JB760M Repeat 1 | 118 | 204 | 0,578431373 | 5 | 2,32E-15 | 1,16E-14 | TRUE | TRUE | TRUE | TRUE |
| | JB760M Repeat 1 Cycle 10 | 139 | 144 | 0,97 | JB760M Repeat 1 | 118 | 204 | 0,58 | 5 | 3,02E-18 | 1,51E-17 | TRUE | TRUE | TRUE | TRUE |
| ASL JB60M Repeat 2 | JB760M Repeat 2 | 128 | 208 | 0,615384615 | JB760M Repeat 2 | 128 | 208 | 0,615384615 | 5 | 1,00E+00 | 1,00E+00 | FALSE | FALSE | FALSE | FALSE |
| (Supplemental Figure 5B) | JB760M Repeat 2 Cycle 2 | 113 | 184 | 0,614130435 | JB760M Repeat 2 | 128 | 208 | 0,615384615 | 5 | 1 | 1 | FALSE | FALSE | FALSE | FALSE |
| | JB760M Repeat 2 Cycle 4 | 28 | 208 | 0,134615385 | JB760M Repeat 2 | 128 | 208 | 0,615384615 | 5 | 5,66E-25 | 2,83E-24 | TRUE | TRUE | TRUE | TRUE |
| | JB760M Repeat 2 Cycle 7 | 150 | 172 | 0,872093023 | JB760M Repeat 2 | 128 | 208 | 0,615384615 | 5 | 1,46E-08 | 7,31E-08 | TRUE | TRUE | TRUE | TRUE |
| | JB760M Repeat 2 Cycle 10 | 131 | 156 | 0,84 | JB760M Repeat 2 | 128 | 208 | 0,62 | 5 | 2,47E-06 | 1,24E-05 | TRUE | TRUE | TRUE | TRUE |
| ASL JB60M Repeat 3 | JB760M Repeat 3 | 114 | 212 | 0,537735849 | JB760M Repeat 3 | 114 | 212 | 0,537735849 | 6 | 1 | 1 | FALSE | FALSE | FALSE | FALSE |
| (Supplemental Figure 5B) | JB760M Repeat 3 Cycle 2 | 102 | 200 | 0,51 | JB760M Repeat 3 | 114 | 212 | 0,537735849 | 6 | 0,6218886 | 1 | FALSE | FALSE | FALSE | FALSE |
| | JB760M Repeat 3 Cycle 3 | 108 | 208 | 0,519230769 | JB760M Repeat 3 | 114 | 212 | 0,537735849 | 6 | 0,7693774 | 1 | FALSE | FALSE | FALSE | FALSE |
| | JB760M Repeat 3 Cycle 4 | 84 | 192 | 0,4375 | JB760M Repeat 3 | 114 | 212 | 0,537735849 | 6 | 0,04683443 | 0,2810066 | FALSE | FALSE | FALSE | FALSE |
| | JB760M Repeat 3 Cycle 6 | 144 | 196 | 0,734693878 | JB760M Repeat 3 | 114 | 212 | 0,537735849 | 6 | 3,91E-05 | 0,000234372 | TRUE | TRUE | TRUE | FALSE |
| | JB760M Repeat 3 Cycle 10 | 163 | 188 | 0,87 | JB760M Repeat 3 | 114 | 212 | 0,54 | 6 | 3,65E-13 | 2,19E-12 | TRUE | TRUE | TRUE | TRUE |
| ASL L972 | L972M | 24 | 108 | 0,222222222 | L972M | 24 | 108 | 0,222222222 | 6 | 1,00E+00 | 1,00E+00 | FALSE | FALSE | FALSE | FALSE |
| (Figure 5C) | XM409 Cycle 1 | 33 | 152 | 0,217105263 | L972M | 24 | 108 | 0,222222222 | 6 | 1,00E+00 | 1,00E+00 | FALSE | FALSE | FALSE | FALSE |
| | XM409 Cycle 2 | 55 | 184 | 0,298913043 | L972M | 24 | 108 | 0,222222222 | 6 | 1,74E-01 | 1,00E+00 | FALSE | FALSE | FALSE | FALSE |
| | XM409 Cycle 4 | 128 | 384 | 0,333333333 | L972M | 24 | 108 | 0,222222222 | 6 | 3,35E-02 | 2,01E-01 | FALSE | FALSE | FALSE | FALSE |
| | XM409 Cycle 6 | 67 | 216 | 0,310185185 | L972M | 24 | 108 | 0,222222222 | 6 | 1,16E-01 | 6,94E-01 | FALSE | FALSE | FALSE | FALSE |
| | XM409 Cycle 8 | 138 | 212 | 0,65 | L972M | 24 | 108 | 0,22 | 6 | 2,10E-13 | 1,26E-12 | TRUE | TRUE | TRUE | TRUE |
| ASL JB953 | JB953M_no4C | 62 | 96 | 0,645833333 | XM603 Cycle 1 | 170 | 400 | 0,425 | 6 | 0,000104132 | 0,000624792 | TRUE | TRUE | TRUE | FALSE |
| (Figure 5C) | XM603 Cycle 1 | 170 | 400 | 0,425 | XM603 Cycle 1 | 170 | 400 | 0,425 | 6 | 1,00E+00 | 1,00E+00 | FALSE | FALSE | FALSE | FALSE |
| | XM603 Cycle 2 | 110 | 216 | 0,509259259 | XM603 Cycle 1 | 170 | 400 | 0,425 | 6 | 5,11E-02 | 3,07E-01 | FALSE | FALSE | FALSE | FALSE |
| | XM603 Cycle 3 | 104 | 208 | 0,5 | XM603 Cycle 1 | 170 | 400 | 0,425 | 6 | 8,59E-02 | 5,15E-01 | FALSE | FALSE | FALSE | FALSE |
| | XM603 Cycle 4 | 142 | 356 | 0,398876404 | XM603 Cycle 1 | 170 | 400 | 0,425 | 6 | 5,05E-01 | 1,00E+00 | FALSE | FALSE | FALSE | FALSE |
| | XM603 Cycle 8 | 172 | 216 | 0,80 | XM603 Cycle 1 | 170 | 400 | 0,43 | 6 | 1,50E-19 | 9,02E-19 | TRUE | TRUE | TRUE | TRUE |
| ASL JB838 | XM584 Cycle 1 | 176 | 212 | 0,830188679 | XM584 Cycle 1 | 176 | 212 | 0,830188679 | 2 | 1 | 1 | FALSE | FALSE | FALSE | FALSE |
| (Figure 5D) | XM584 Cycle 4 | 196 | 212 | 0,92 | XM584 Cycle 1 | 176 | 212 | 0,83 | 2 | 4,54E-03 | 9,07E-03 | TRUE | TRUE | FALSE | FALSE |
| ASL JB845 | JB845M_no4C | 100 | 104 | 0,961538462 | JB845M_no4C | 100 | 104 | 0,961538462 | 3 | 1 | 1 | FALSE | FALSE | FALSE | FALSE |
| (Figure 5D) | XM586 Cycle 1 | 182 | 208 | 0,875 | JB845M_no4C | 100 | 104 | 0,961538462 | 3 | 0,01416378 | 0,04249134 | TRUE | FALSE | FALSE | FALSE |
| | XM586 Cycle 4 | 208 | 216 | 0,96 | JB845M_no4C | 100 | 104 | 0,96 | 3 | 1 | 1 | FALSE | FALSE | FALSE | FALSE |
| ASL 853 | XM588 Cycle 1 | 138 | 184 | 0,75 | XM588 Cycle 1 | 138 | 184 | 0,75 | 2 | 1 | 1 | FALSE | FALSE | FALSE | FALSE |
| (Figure 5D) | XM588 Cycle 4 | 165 | 196 | 0,84 | XM588 Cycle 1 | 138 | 184 | 0,75 | 2 | 0,02997884 | 0,05995767 | FALSE | FALSE | FALSE | FALSE |
| ASL 872 | XM590 Cycle 1 | 163 | 212 | 0,768867925 | XM590 Cycle 1 | 163 | 212 | 0,768867925 | 2 | 1 | 1 | FALSE | FALSE | FALSE | FALSE |
| (Figure 5D) | XM590 Cycle 4 | 190 | 212 | 0,90 | XM590 Cycle 1 | 163 | 212 | 0,77 | 2 | 0,000644522 | 0,001289045 | TRUE | TRUE | FALSE | FALSE |
| ASL 873 | JB873M_no4C | 84 | 100 | 0,84 | XM591 Cycle 1 | 76 | 160 | 0,475 | 3 | 2,13E-09 | 6,38E-09 | TRUE | TRUE | TRUE | TRUE |
| (Figure 5D) | XM591 Cycle 1 | 76 | 160 | 0,475 | XM591 Cycle 1 | 76 | 160 | 0,475 | 3 | 1,00E+00 | 1,00E+00 | FALSE | FALSE | FALSE | FALSE |
| | XM591 Cycle 4 | 147 | 200 | 0,74 | XM591 Cycle 1 | 76 | 160 | 0,48 | 3 | 4,82017E-07 | 1,44605E-06 | TRUE | TRUE | TRUE | TRUE |
| ASL 884 | XM593 Cycle 1 | 127 | 208 | 0,610576923 | XM593 Cycle 1 | 127 | 208 | 0,610576923 | 2 | 1 | 1 | FALSE | FALSE | FALSE | FALSE |
| (Figure 5D) | XM593 Cycle 4 | 189 | 204 | 0,93 | XM593 Cycle 1 | 127 | 208 | 0,61 | 2 | 5,78709E-15 | 1,15742E-14 | TRUE | TRUE | TRUE | TRUE |
| ASL 913 | JB913M_no4C | 78 | 100 | 0,78 | JB913M_no4C | 78 | 100 | 0,78 | 3 | 1 | 1 | FALSE | FALSE | FALSE | FALSE |
| (Figure 5D) | XM594 Cycle 1 | 135 | 176 | 0,767045455 | JB913M_no4C | 78 | 100 | 0,78 | 3 | 0,8818196 | 1 | FALSE | FALSE | FALSE | FALSE |
| | XM594 Cycle 4 | 199 | 212 | 0,94 | JB913M_no4C | 78 | 100 | 0,78 | 3 | 7,84516E-05 | 0,000235355 | TRUE | TRUE | TRUE | FALSE |
| ASL JB845 & JB913 F1 refered | JB845M_no4C | 100 | 104 | 0,961538462 | XM586 Cycle 1 | 182 | 208 | 0,875 | 6 | 0,01416378 | 0,08498269 | FALSE | FALSE | FALSE | FALSE |
| (Figure 5D and text) | XM586 Cycle 1 | 182 | 208 | 0,875 | XM586 Cycle 1 | 182 | 208 | 0,875 | 6 | 1 | 1 | FALSE | FALSE | FALSE | FALSE |
| | XM586 Cycle 4 | 208 | 216 | 0,962962963 | XM586 Cycle 1 | 182 | 208 | 0,875 | 6 | 0,001039411 | 0,006236468 | TRUE | TRUE | FALSE | FALSE |
| | JB913M_no4C | 78 | 100 | 0,78 | XM594 Cycle 1 | 135 | 176 | 0,767045455 | 6 | 0,8818196 | 1 | FALSE | FALSE | FALSE | FALSE |
| | XM594 Cycle 1 | 135 | 176 | 0,767045455 | XM594 Cycle 1 | 135 | 176 | 0,767045455 | 6 | 1 | 1 | FALSE | FALSE | FALSE | FALSE |
| | XM594 Cycle 4 | 199 | 212 | 0,94 | XM594 Cycle 1 | 135 | 176 | 0,77 | 6 | 1,33207E-06 | 7,99243E-06 | TRUE | TRUE | TRUE | TRUE |
| **Experiment** | **Sample** | **Counted Ade6-Nat** | **Total** | **% Nat** | **Sample** | **Counted Ade6-Nat** | **Total** | **% Nat** | **Number of tests** | **p-value(alt="two.sided")** | **Bonferroni Adjusted P-value** | *** (<0.05)** | **** (<0.01)** | ***** (<0.001)** | ****** (<0.0001)** |
| JB953 ASL Fitness | Cycle 0 Initial | 88 | 206 | 0,427184466 | Cycle 0 Final | 83 | 246 | 0,337398374 | 6 | 0,05217668 | 0,3130601 | FALSE | FALSE | FALSE | FALSE |
| Nat cels | Cycle 1 Initial | 52 | 109 | 0,47706422 | Cycle 1 Final | 37 | 75 | 0,493333333 | 6 | 0,8811427 | 1 | FALSE | FALSE | FALSE | FALSE |
| | Cycle 2 Initial | 59 | 94 | 0,627659574 | Cycle 2 Final | 43 | 67 | 0,641791045 | 6 | 0,8698848 | 1 | FALSE | FALSE | FALSE | FALSE |
| | Cycle 3 Initial | 56 | 107 | 0,523364486 | Cycle 3 Final | 20 | 103 | 0,194174757 | 6 | 7,91E-07 | 4,75E-06 | TRUE | TRUE | TRUE | TRUE |
| | Cycle 4 Initial | 53 | 123 | 0,430894309 | Cycle 4 Final | 4 | 56 | 0,071428571 | 6 | 6,28E-07 | 3,77E-06 | TRUE | TRUE | TRUE | TRUE |
| | Cycle 4 Initial | 85 | 226 | 0,38 | Cycle 4 Final | 25 | 156 | 0,16 | 6 | 3,70179E-06 | 2,22108E-05 | TRUE | TRUE | TRUE | TRUE |
| **Experiment** | **Sample** | **Counted Ade6-Hyg** | **Total** | **% Hyg** | **Sample** | **Counted Ade6-Hyg** | **Total** | **% Hyg** | **Number of tests** | **p-value(alt="two.sided")** | **Bonferroni Adjusted P-value** | *** (<0.05)** | **** (<0.01)** | ***** (<0.001)** | ****** (<0.0001)** |

| | | | | | | | | | | | | | | | |
|---|---|---|---|---|---|---|---|---|---|---|---|---|---|---|---|
| JB953 ASL Fitness | Cycle 0 Initial | 118 | 206 | 0,572815534 | Cycle 0 Final | 163 | 246 | 0,662601626 | 6 | 0,05217668 | 0,3130601 | FALSE | FALSE | FALSE | FALSE |
| Hyg cels | Cycle 1 Initial | 57 | 109 | 0,52293578 | Cycle 1 Final | 38 | 75 | 0,506666667 | 6 | 0,8811427 | 1 | FALSE | FALSE | FALSE | FALSE |
| | Cycle 2 Initial | 35 | 94 | 0,372340426 | Cycle 2 Final | 24 | 67 | 0,358208955 | 6 | 0,8698848 | 1 | FALSE | FALSE | FALSE | FALSE |
| | Cycle 3 Initial | 51 | 107 | 0,476635514 | Cycle 3 Final | 83 | 103 | 0,805825243 | 6 | 7,91E-07 | 4,75E-06 | TRUE | TRUE | TRUE | TRUE |
| | Cycle 4 Initial | 70 | 123 | 0,569105691 | Cycle 4 Final | 52 | 56 | 0,928571429 | 6 | 6,28E-07 | 3,77E-06 | TRUE | TRUE | TRUE | TRUE |
| | Cycle 4 Initial | 141 | 226 | 0,62 | Cycle 4 Final | 131 | 156 | 0,84 | 6 | 3,70179E-06 | 2,22108E-05 | TRUE | TRUE | TRUE | TRUE |

| **Experiment** | **Strain/cross 1** | **Viable** | **Total** | **Viability** | **Strain/cross 2** | **Viable** | **Total** | **Viability** | **Number of tests** | **p-value(alt="two.sided")** | **Bonferroni Adjusted P-value** | *** (<0.05)** | **** (<0.01)** | ***** (<0.001)** | ****** (<0.0001)** |
|---|---|---|---|---|---|---|---|---|---|---|---|---|---|---|---|
| JB32 MM-N | MMN_JB32_R1_T0 | 192 | 208 | 0,923076923 | MMN_JB32_R1_T0 | 192 | 208 | 0,923076923 | 6 | 1 | 1 | FALSE | FALSE | FALSE | FALSE |
| Repeat 1 | MMN-JB32_R1_T1 | 192 | 208 | 0,923076923 | MMN_JB32_R1_T0 | 192 | 208 | 0,923076923 | 6 | 1 | 1 | FALSE | FALSE | FALSE | FALSE |
| | MMN-JB32_R1_T2 | 201 | 212 | 0,948113208 | MMN_JB32_R1_T0 | 192 | 208 | 0,923076923 | 6 | 0,3250716 | 1 | FALSE | FALSE | FALSE | FALSE |
| | MMN-JB32_R1_T3 | 182 | 208 | 0,875 | MMN_JB32_R1_T0 | 192 | 208 | 0,923076923 | 6 | 0,1422976 | 0,8537859 | FALSE | FALSE | FALSE | FALSE |
| | MMN-JB32_R1_T4 | 173 | 208 | 0,831730769 | MMN_JB32_R1_T0 | 192 | 208 | 0,923076923 | 6 | 0,006668476 | 0,04001086 | TRUE | FALSE | FALSE | FALSE |
| | MMN-JB32_R1_T5 | 176 | 208 | 0,85 | MMN_JB32_R1_T0 | 192 | 208 | 0,92 | 6 | 0,02056773 | 0,1234064 | FALSE | FALSE | FALSE | FALSE |
| JB32 MM-N | MMN_JB32_R2_T1 | 204 | 212 | 0,962264151 | MMN_JB32_R2_T1 | 204 | 212 | 0,962264151 | 5 | 1 | 1 | FALSE | FALSE | FALSE | FALSE |
| Repeat 2 | MMN-JB32_R2_T2 | 207 | 216 | 0,958333333 | MMN_JB32_R2_T1 | 204 | 212 | 0,962264151 | 5 | 1 | 1 | FALSE | FALSE | FALSE | FALSE |
| | MMN-JB32_R2_T3 | 190 | 212 | 0,896226415 | MMN_JB32_R2_T1 | 204 | 212 | 0,962264151 | 5 | 0,01262809 | 0,06314047 | FALSE | FALSE | FALSE | FALSE |
| | MMN-JB32_R2_T4 | 187 | 204 | 0,916666667 | MMN_JB32_R2_T1 | 204 | 212 | 0,962264151 | 5 | 0,06293814 | 0,3146907 | FALSE | FALSE | FALSE | FALSE |
| | MMN-JB32_R2_T5 | 180 | 192 | 0,94 | MMN_JB32_R2_T1 | 204 | 212 | 0,96 | 5 | 0,2623984 | 1 | FALSE | FALSE | FALSE | FALSE |
| JB760 MM-N | MMN_JB760_R1_T0 | 52 | 212 | 0,245283019 | MMN_JB760_R1_T0 | 52 | 212 | 0,245283019 | 4 | 1 | 1 | FALSE | FALSE | FALSE | FALSE |
| Repeat 1 | MMN_JB760_R1_T1 | 137 | 204 | 0,671568627 | MMN_JB760_R1_T0 | 52 | 212 | 0,245283019 | 4 | 1,34E-18 | 5,35E-18 | TRUE | TRUE | TRUE | TRUE |
| | MMN_JB760_R1_T2 | 81 | 192 | 0,421875 | MMN_JB760_R1_T0 | 52 | 212 | 0,245283019 | 4 | 0,000198769 | 0,000795075 | TRUE | TRUE | TRUE | FALSE |
| | MMN_JB760_R1_T3 | 88 | 140 | 0,63 | MMN_JB760_R1_T0 | 52 | 212 | 0,25 | 4 | 1,02476E-12 | 4,09903E-12 | TRUE | TRUE | TRUE | TRUE |
| JB760 MM-N | MMN_JB760_R2_T0 | 12 | 172 | 0,069767442 | MMN_JB760_R2_T0 | 12 | 172 | 0,069767442 | 4 | 1 | 1 | FALSE | FALSE | FALSE | FALSE |
| Repeat 1 | MMN_JB760_R2_T1 | 139 | 204 | 0,681372549 | MMN_JB760_R2_T0 | 12 | 172 | 0,069767442 | 4 | 4,59E-37 | 1,84E-36 | TRUE | TRUE | TRUE | TRUE |
| | MMN_JB760_R2_T2 | 111 | 216 | 0,513888889 | MMN_JB760_R2_T0 | 12 | 172 | 0,069767442 | 4 | 1,03E-22 | 4,10E-22 | TRUE | TRUE | TRUE | TRUE |
| | MMN_JB760_R2_T3 | 120 | 212 | 0,57 | MMN_JB760_R2_T0 | 12 | 172 | 0,07 | 4 | 1,08421E-26 | 4,33685E-26 | TRUE | TRUE | TRUE | TRUE |
| JB32 MM-N | MMN-JB32_R1_T1 | 192 | 208 | 0,923076923 | MMN-JB32_R1_T1_F1 | 93 | 112 | 0,830357143 | 5 | 0,01447794 | 0,07238968 | FALSE | FALSE | FALSE | FALSE |
| Epigenetic degeneration | MMN-JB32_R1_T2 | 201 | 212 | 0,948113208 | MMN-JB32_R1_T2_F1 | 204 | 212 | 0,962264151 | 5 | 0,6398883 | 1 | FALSE | FALSE | FALSE | FALSE |
| F1 effect | MMN-JB32_R1_T3 | 182 | 208 | 0,875 | MMN-JB32_R1_T3_F1 | 181 | 196 | 0,923469388 | 5 | 0,1374511 | 0,6872554 | FALSE | FALSE | FALSE | FALSE |
| | MMN-JB32_R1_T4 | 173 | 208 | 0,831730769 | MMN-JB32_R1_T4_F1 | 184 | 204 | 0,901960784 | 5 | 0,04243027 | 0,2121513 | FALSE | FALSE | FALSE | FALSE |
| | MMN-JB32_R1_T5 | 176 | 208 | 0,85 | MMN-JB32_R1_T5_F1 | 160 | 184 | 0,87 | 5 | 0,5641509 | 1 | FALSE | FALSE | FALSE | FALSE |
| JB32 MM-N | MMN-JB32_R1_T1_F1 | 93 | 112 | 0,830357143 | MMN-JB32_R1_T2_F1 | 204 | 212 | 0,962264151 | 5 | 8,78E-05 | 0,000438785 | TRUE | TRUE | TRUE | FALSE |
| Epigenetic degeneration | MMN-JB32_R1_T2_F1 | 204 | 212 | 0,962264151 | MMN-JB32_R1_T2_F1 | 204 | 212 | 0,962264151 | 5 | 1 | 1 | FALSE | FALSE | FALSE | FALSE |
| Black dotted line test | MMN-JB32_R1_T3_F1 | 181 | 196 | 0,923469388 | MMN-JB32_R1_T2_F1 | 204 | 212 | 0,962264151 | 5 | 0,1311647 | 0,6558237 | FALSE | FALSE | FALSE | FALSE |
| | MMN-JB32_R1_T4_F1 | 184 | 204 | 0,901960784 | MMN-JB32_R1_T2_F1 | 204 | 212 | 0,962264151 | 5 | 0,01793738 | 0,0896869 | FALSE | FALSE | FALSE | FALSE |
| | MMN-JB32_R1_T5_F1 | 160 | 184 | 0,87 | MMN-JB32_R1_T2_F1 | 204 | 212 | 0,96 | 5 | 0,000782119 | 0,003910593 | TRUE | TRUE | FALSE | FALSE |
| JB760 MM-N | MMN_JB760_R2_T1 | 139 | 204 | 0,681372549 | MMN_JB760_R2_T1_F1 | 90 | 188 | 0,478723404 | 3 | 6,03E-05 | 0,000180999 | TRUE | TRUE | TRUE | FALSE |
| Epigenetic degeneration | MMN_JB760_R2_T2 | 111 | 216 | 0,513888889 | MMN_JB760_R2_T2_F1 | 73 | 208 | 0,350961538 | 3 | 0,000850479 | 0,002551437 | TRUE | TRUE | FALSE | FALSE |
| F1 effect | MMN_JB760_R2_T3 | 120 | 212 | 0,57 | MMN_JB760_R2_T3_F1 | 59 | 192 | 0,31 | 3 | 1,7161E-07 | 5,14831E-07 | TRUE | TRUE | TRUE | TRUE |
| JB760 MM-N | MMN_JB760_R2_T1_F1 | 90 | 188 | 0,478723404 | MMN_JB760_R2_T1_F1 | 90 | 188 | 0,478723404 | 3 | 1 | 1 | FALSE | FALSE | FALSE | FALSE |
| Epigenetic degeneration | MMN_JB760_R2_T2_F1 | 73 | 208 | 0,350961538 | MMN_JB760_R2_T1_F1 | 90 | 188 | 0,478723404 | 3 | 0,01073004 | 0,03219011 | TRUE | FALSE | FALSE | FALSE |
| Black dotted line test | MMN_JB760_R2_T3_F1 | 59 | 192 | 0,31 | MMN_JB760_R2_T1_F1 | 90 | 188 | 0,48 | 3 | 0,000756791 | 0,002270372 | TRUE | TRUE | FALSE | FALSE |

### Supplemental Table S3: SNPs differences

| Strain Clonal Group | Strain 1 | Strain 2 | SNPs differences |
|---|---|---|---|
| Leupold's strains | L972 | JB32 | 29 |
| | L972 | L968 | 14 |
| | L972 | JB761 | 21 |
| | L972 | JB868 | 22 |
| | L972 | JB906 | 12 |
| | L972 | JB937 | 13 |
| | L972 | JB941 | 17 |
| | L972 | JB945 | 14 |
| | JB32 | L968 | 26 |
| | JB32 | JB761 | 36 |
| | JB32 | JB868 | 37 |
| | JB32 | JB906 | 24 |
| | JB32 | JB937 | 28 |
| | JB32 | JB941 | 31 |
| | JB32 | JB945 | 29 |
| | L968 | JB761 | 21 |
| | L968 | JB868 | 22 |
| | L968 | JB906 | 10 |
| | L968 | JB937 | 12 |
| | L968 | JB941 | 16 |
| | L968 | JB945 | 13 |
| | JB761 | JB868 | 31 |
| | JB761 | JB906 | 19 |
| | JB761 | JB937 | 21 |
| | JB761 | JB941 | 25 |
| | JB761 | JB945 | 22 |
| | JB868 | JB906 | 20 |
| | JB868 | JB937 | 22 |
| | JB868 | JB941 | 26 |
| | JB868 | JB945 | 23 |
| | JB906 | JB937 | 3 |
| | JB906 | JB941 | 5 |
| | JB906 | JB945 | 0 |
| | JB937 | JB941 | 6 |
| | JB937 | JB945 | 5 |
| | JB941 | JB945 | 7 |
| NCYC132 clonal group | JB864 | JB876 | 41 |
| | JB864 | JB890 | 77 |
| | JB876 | JB890 | 86 |
| JB1114-JB1175 clonal strains | JB1114 | JB1175 | 12 |
| JB760 clonal group | JB760 | JB888 | 23 |
| | JB760 | JB889 | 24 |
| | JB888 | JB889 | 13 |

### Supplemental Table S4: ANOVA tests

| Experiment | Strain name | Group | Viable | No_viable | Total |
|---|---|---|---|---|---|
| One way ANOVA | L968 | 1 | 8 | 0 | 8 |
| Canonical Leupold's strains | L968 | 1 | 47 | 13 | 60 |
| All data | L968 | 1 | 53 | 7 | 60 |
| (Suplemental Figure 1C) | L968 | 1 | 43 | 13 | 56 |
| | L968 | 1 | 59 | 5 | 64 |
| | L968 | 1 | 30 | 6 | 36 |
| | L968 | 1 | 25 | 7 | 32 |
| | L968 | 1 | 27 | 9 | 36 |
| | L968 | 1 | 26 | 6 | 32 |
| | L968 | 1 | 29 | 7 | 36 |
| | L968 | 1 | 34 | 2 | 36 |
| | L972(JB22) | 2 | 24 | 12 | 36 |
| | L972(JB22) | 2 | 32 | 4 | 36 |
| | L972(JB22) | 2 | 34 | 2 | 36 |
| | L972(JB22) | 2 | 27 | 5 | 32 |
| | L972(JB22) | 2 | 32 | 4 | 36 |
| | L972(JB22) | 2 | 27 | 5 | 32 |
| | L972(JB22) | 2 | 12 | 0 | 12 |
| | L972(JB22) | 2 | 8 | 0 | 8 |
| | L972(JB22) | 2 | 22 | 10 | 32 |
| | L972(JB22) | 2 | 21 | 7 | 28 |
| | L972(JB22) | 2 | 35 | 1 | 36 |
| | L972(JB22) | 2 | 28 | 0 | 28 |
| | L972(JB903) | 3 | 32 | 4 | 36 |
| | L972(JB903) | 3 | 30 | 6 | 36 |
| | L972(JB903) | 3 | 31 | 5 | 36 |
| | L972(JB903) | 3 | 28 | 0 | 28 |
| | L972(JB903) | 3 | 32 | 0 | 32 |
| | L972(JB903) | 3 | 27 | 5 | 32 |
| | L972(JB903) | 3 | 32 | 4 | 36 |
| | L972(JB903) | 3 | 28 | 4 | 32 |
| | L972(JB903) | 3 | 19 | 13 | 32 |
| | L972(JB903) | 3 | 29 | 3 | 32 |
| | L972(JB903) | 3 | 25 | 7 | 32 |
| | L972(JB903) | 3 | 27 | 5 | 32 |
| | L975 | 4 | 31 | 1 | 32 |
| | L975 | 4 | 36 | 0 | 36 |
| | L975 | 4 | 33 | 3 | 36 |
| | L975 | 4 | 33 | 3 | 36 |
| | L975 | 4 | 35 | 1 | 36 |
| | L975 | 4 | 30 | 2 | 32 |
| | L975 | 4 | 34 | 2 | 36 |
| | L975 | 4 | 32 | 0 | 32 |
| | L975 | 4 | 36 | 0 | 36 |
| | L975 | 4 | 34 | 2 | 36 |
| | L975 | 4 | 36 | 0 | 36 |
| | JB32 | 5 | 36 | 0 | 36 |
| | JB32 | 5 | 35 | 1 | 36 |
| | JB32 | 5 | 34 | 2 | 36 |
| | JB32 | 5 | 29 | 3 | 32 |
| | JB32 | 5 | 36 | 0 | 36 |
| | JB32 | 5 | 30 | 2 | 32 |
| | JB32 | 5 | 26 | 2 | 28 |
| | JB32 | 5 | 31 | 5 | 36 |
| | JB32 | 5 | 33 | 3 | 36 |
| | JB32 | 5 | 31 | 1 | 32 |
| | JB32 | 5 | 26 | 2 | 28 |
| | JB32 | 5 | 36 | 0 | 36 |

| term | estimate | std.error | statistic | p.value |
|---|---|---|---|---|
| **(Intercept)** | -1,155860781 | 0,148219602 | -7,79829905 | 6,27E-15 |
| **Group** | -0,355011328 | 0,053651489 | -6,616989295 | 3,67E-11 |

| Experiment | Strain name | Group | Viable | No_viable | Total |
|---|---|---|---|---|---|
| One way ANOVA | L975 | 1 | 31 | 1 | 32 |
| Canonical Leupold's strains | L975 | 1 | 36 | 0 | 36 |
| L975 and JB32 | L975 | 1 | 33 | 3 | 36 |
| (Suplemental Figure 1C) | L975 | 1 | 33 | 3 | 36 |
| | L975 | 1 | 35 | 1 | 36 |
| | L975 | 1 | 30 | 2 | 32 |
| | L975 | 1 | 34 | 2 | 36 |
| | L975 | 1 | 32 | 0 | 32 |
| | L975 | 1 | 36 | 0 | 36 |
| | L975 | 1 | 34 | 2 | 36 |
| | L975 | 1 | 36 | 0 | 36 |
| | JB32 | 2 | 36 | 0 | 36 |
| | JB32 | 2 | 35 | 1 | 36 |
| | JB32 | 2 | 34 | 2 | 36 |
| | JB32 | 2 | 29 | 3 | 32 |

| | | | | | |
|---|---|---|---|---|---|
| | JB32 | 2 | 36 | 0 | 36 |
| | JB32 | 2 | 30 | 2 | 32 |
| | JB32 | 2 | 26 | 2 | 28 |
| | JB32 | 2 | 31 | 5 | 36 |
| | JB32 | 2 | 33 | 3 | 36 |
| | JB32 | 2 | 31 | 1 | 32 |
| | JB32 | 2 | 26 | 2 | 28 |
| | JB32 | 2 | 36 | 0 | 36 |

| term | estimate | std.error | statistic | p.value |
|---|---|---|---|---|
| **(Intercept)** | -3,645378801 | 0,588859144 | -6,190578573 | 5,99E-10 |
| **Group** | 0,370933125 | 0,35264895 | 1,051848089 | 0,292869252 |

| Experiment | Strain name | Group | Viable | No_viable | Total |
|---|---|---|---|---|---|
| One way ANOVA | L968 | 1 | 8 | 0 | 8 |
| Canonical Leupold's strains | L968 | 1 | 47 | 13 | 60 |
| L968 and L972 | L968 | 1 | 53 | 7 | 60 |
| (Suplemental Figure 1C) | L968 | 1 | 43 | 13 | 56 |
| | L968 | 1 | 59 | 5 | 64 |
| | L968 | 1 | 30 | 6 | 36 |
| | L968 | 1 | 25 | 7 | 32 |
| | L968 | 1 | 27 | 9 | 36 |
| | L968 | 1 | 26 | 6 | 32 |
| | L968 | 1 | 29 | 7 | 36 |
| | L968 | 1 | 34 | 2 | 36 |
| | L972(JB22) | 2 | 24 | 12 | 36 |
| | L972(JB22) | 2 | 32 | 4 | 36 |
| | L972(JB22) | 2 | 34 | 2 | 36 |
| | L972(JB22) | 2 | 27 | 5 | 32 |
| | L972(JB22) | 2 | 32 | 4 | 36 |
| | L972(JB22) | 2 | 27 | 5 | 32 |
| | L972(JB22) | 2 | 12 | 0 | 12 |
| | L972(JB22) | 2 | 8 | 0 | 8 |
| | L972(JB22) | 2 | 22 | 10 | 32 |
| | L972(JB22) | 2 | 21 | 7 | 28 |
| | L972(JB22) | 2 | 35 | 1 | 36 |
| | L972(JB22) | 2 | 28 | 0 | 28 |
| | L972(JB903) | 3 | 32 | 4 | 36 |
| | L972(JB903) | 3 | 30 | 6 | 36 |
| | L972(JB903) | 3 | 31 | 5 | 36 |
| | L972(JB903) | 3 | 28 | 0 | 28 |
| | L972(JB903) | 3 | 32 | 0 | 32 |
| | L972(JB903) | 3 | 27 | 5 | 32 |
| | L972(JB903) | 3 | 32 | 4 | 36 |
| | L972(JB903) | 3 | 28 | 4 | 32 |
| | L972(JB903) | 3 | 19 | 13 | 32 |
| | L972(JB903) | 3 | 29 | 3 | 32 |
| | L972(JB903) | 3 | 25 | 7 | 32 |
| | L972(JB903) | 3 | 27 | 5 | 32 |

| term | estimate | std.error | statistic | p.value |
|---|---|---|---|---|
| **(Intercept)** | -1,553966002 | 0,20061554 | -7,745990168 | 9,48E-15 |
| **Group** | -0,092368028 | 0,096435466 | -0,957822179 | 0,338152433 |

| Experiment | Strain name | Group | Viable | No_viable | Total |
|---|---|---|---|---|---|
| One Way Anova | L968 | 1 | 8 | 0 | 8 |
| All Leupold's strains | L968 | 1 | 47 | 13 | 60 |
| All data | L968 | 1 | 53 | 7 | 60 |
| (Suplemental Figure 1C) | L968 | 1 | 43 | 13 | 56 |
| | L968 | 1 | 59 | 5 | 64 |
| | L968 | 1 | 30 | 6 | 36 |
| | L968 | 1 | 25 | 7 | 32 |
| | L968 | 1 | 27 | 9 | 36 |
| | L968 | 1 | 26 | 6 | 32 |
| | L968 | 1 | 29 | 7 | 36 |
| | L968 | 1 | 34 | 2 | 36 |
| | L972(JB22) | 2 | 24 | 12 | 36 |
| | L972(JB22) | 2 | 32 | 4 | 36 |
| | L972(JB22) | 2 | 34 | 2 | 36 |
| | L972(JB22) | 2 | 27 | 5 | 32 |
| | L972(JB22) | 2 | 32 | 4 | 36 |
| | L972(JB22) | 2 | 27 | 5 | 32 |
| | L972(JB22) | 2 | 12 | 0 | 12 |
| | L972(JB22) | 2 | 8 | 0 | 8 |
| | L972(JB22) | 2 | 22 | 10 | 32 |
| | L972(JB22) | 2 | 21 | 7 | 28 |
| | L972(JB22) | 2 | 35 | 1 | 36 |
| | L972(JB22) | 2 | 28 | 0 | 28 |
| | L972(JB903) | 3 | 32 | 4 | 36 |
| | L972(JB903) | 3 | 30 | 6 | 36 |
| | L972(JB903) | 3 | 31 | 5 | 36 |
| | L972(JB903) | 3 | 28 | 0 | 28 |
| | L972(JB903) | 3 | 32 | 0 | 32 |

| | | | | |
|---|---|---|---|---|
| L972(JB903) | 3 | 27 | 5 | 32 |
| L972(JB903) | 3 | 32 | 4 | 36 |
| L972(JB903) | 3 | 28 | 4 | 32 |
| L972(JB903) | 3 | 19 | 13 | 32 |
| L972(JB903) | 3 | 29 | 3 | 32 |
| L972(JB903) | 3 | 25 | 7 | 32 |
| L972(JB903) | 3 | 27 | 5 | 32 |
| L975 | 4 | 31 | 1 | 32 |
| L975 | 4 | 36 | 0 | 36 |
| L975 | 4 | 33 | 3 | 36 |
| L975 | 4 | 33 | 3 | 36 |
| L975 | 4 | 35 | 1 | 36 |
| L975 | 4 | 30 | 2 | 32 |
| L975 | 4 | 34 | 2 | 36 |
| L975 | 4 | 32 | 0 | 32 |
| L975 | 4 | 36 | 0 | 36 |
| L975 | 4 | 34 | 2 | 36 |
| L975 | 4 | 36 | 0 | 36 |
| JB32 | 5 | 36 | 0 | 36 |
| JB32 | 5 | 35 | 1 | 36 |
| JB32 | 5 | 34 | 2 | 36 |
| JB32 | 5 | 29 | 3 | 32 |
| JB32 | 5 | 36 | 0 | 36 |
| JB32 | 5 | 30 | 2 | 32 |
| JB32 | 5 | 26 | 2 | 28 |
| JB32 | 5 | 31 | 5 | 36 |
| JB32 | 5 | 33 | 3 | 36 |
| JB32 | 5 | 31 | 1 | 32 |
| JB32 | 5 | 26 | 2 | 28 |
| JB32 | 5 | 36 | 0 | 36 |
| JB761 | 6 | 19 | 9 | 28 |
| JB761 | 6 | 16 | 4 | 20 |
| JB761 | 6 | 17 | 19 | 36 |
| JB761 | 6 | 15 | 13 | 28 |
| JB761 | 6 | 25 | 7 | 32 |
| JB761 | 6 | 15 | 13 | 28 |
| JB868 | 7 | 15 | 21 | 36 |
| JB868 | 7 | 16 | 20 | 36 |
| JB868 | 7 | 17 | 15 | 32 |
| JB906 | 8 | 31 | 1 | 32 |
| JB906 | 8 | 31 | 5 | 36 |
| JB906 | 8 | 28 | 0 | 28 |
| JB937 | 9 | 23 | 9 | 32 |
| JB937 | 9 | 32 | 4 | 36 |
| JB937 | 9 | 26 | 10 | 36 |
| JB941 | 10 | 36 | 0 | 36 |
| JB941 | 10 | 32 | 4 | 36 |
| JB941 | 10 | 29 | 7 | 36 |
| JB945 | 11 | 29 | 3 | 32 |
| JB945 | 11 | 26 | 10 | 36 |
| JB945 | 11 | 35 | 1 | 36 |

| term | estimate | std.error | statistic | p.value |
|---|---|---|---|---|
| **(Intercept)** | -1,956431429 | 0,102832781 | -19,02536733 | 1,05E-80 |
| **Group** | 0,042595339 | 0,019076003 | 2,23292783 | 0,025553704 |

| Experiment | Strain name | Group | Viable | No_viable | Total |
|---|---|---|---|---|---|
| One Way ANOVA | JB864 | 1 | 5 | 19 | 24 |
| All NCYC132 self-crosses | JB864 | 1 | 17 | 19 | 36 |
| and Inter-strain crosses | JB864 | 1 | 20 | 16 | 36 |
| (Suplemental Figure 2A) | JB876 | 2 | 6 | 30 | 36 |
| | JB890 | 3 | 4 | 30 | 34 |
| | JB590 x JB864 | 4 | 0 | 28 | 28 |
| | JB590 x JB876 | 5 | 1 | 71 | 72 |
| | JB590 x JB890 | 6 | 1 | 27 | 28 |
| | JB864 x JB876 | 7 | 6 | 102 | 108 |
| | JB864 xJB890 | 8 | 1 | 35 | 36 |
| | JB864 x JB894 | 9 | 2 | 70 | 72 |
| | JB876 x JB890 | 10 | 1 | 35 | 36 |
| | JB876 x JB894 | 11 | 0 | 36 | 36 |
| | JB890 x JB894 | 12 | 0 | 72 | 72 |

| term | estimate | std.error | statistic | p.value |
|---|---|---|---|---|
| **(Intercept)** | 0,082237967 | 0,227394419 | 0,361653409 | 0,717611047 |
| **Group** | 0,49596466 | 0,063486693 | 7,812104175 | 5,62E-15 |

| Experiment | Strain name | Group | Viable | No_viable | Total | Zygotic-azygotic |
|---|---|---|---|---|---|---|
| One Way ANOVA | L972(JB22) | 1 | 24 | 12 | 36 | 1 |
| Zygotic vs azygotic | L972(JB22) | 1 | 32 | 4 | 36 | 1 |
| Alldata | L972(JB22) | 1 | 34 | 2 | 36 | 1 |
| (Figure 3A) | L972(JB22) | 1 | 27 | 5 | 32 | 1 |
| | L972(JB22) | 1 | 32 | 4 | 36 | 1 |
| | L972(JB22) | 1 | 27 | 5 | 32 | 1 |

| | | | | | |
|---|---|---|---|---|---|
| L972(JB22) | 1 | 12 | 0 | 12 | 1 |
| L972(JB22) | 1 | 8 | 0 | 8 | 1 |
| L972(JB22) | 1 | 22 | 10 | 32 | 1 |
| L972(JB22) | 1 | 21 | 7 | 28 | 1 |
| L972(JB22) | 1 | 35 | 1 | 36 | 1 |
| L972(JB22) | 1 | 28 | 0 | 28 | 1 |
| L972(JB903) | 1 | 32 | 4 | 36 | 1 |
| L972(JB903) | 1 | 30 | 6 | 36 | 1 |
| L972(JB903) | 1 | 31 | 5 | 36 | 1 |
| L972(JB903) | 1 | 28 | 0 | 28 | 1 |
| L972(JB903) | 1 | 32 | 0 | 32 | 1 |
| L972(JB903) | 1 | 27 | 5 | 32 | 1 |
| L972(JB903) | 1 | 32 | 4 | 36 | 1 |
| L972(JB903) | 1 | 28 | 4 | 32 | 1 |
| L972(JB903) | 1 | 19 | 13 | 32 | 1 |
| L972(JB903) | 1 | 29 | 3 | 32 | 1 |
| L972(JB903) | 1 | 25 | 7 | 32 | 1 |
| L972(JB903) | 1 | 27 | 5 | 32 | 1 |
| L972M(JB22M) | 1 | 8 | 4 | 12 | 2 |
| L972M(JB22M) | 1 | 7 | 13 | 20 | 2 |
| L972M(JB22M) | 1 | 4 | 20 | 24 | 2 |
| L972M(JB22M) | 1 | 2 | 14 | 16 | 2 |
| L972M(JB22M) | 1 | 3 | 17 | 20 | 2 |
| L972M(JB22M) | 1 | 0 | 16 | 16 | 2 |
| JB32 | 2 | 36 | 0 | 36 | 1 |
| JB32 | 2 | 35 | 1 | 36 | 1 |
| JB32 | 2 | 34 | 2 | 36 | 1 |
| JB32 | 2 | 29 | 3 | 32 | 1 |
| JB32 | 2 | 36 | 0 | 36 | 1 |
| JB32 | 2 | 30 | 2 | 32 | 1 |
| JB32 | 2 | 26 | 2 | 28 | 1 |
| JB32 | 2 | 31 | 5 | 36 | 1 |
| JB32 | 2 | 33 | 3 | 36 | 1 |
| JB32 | 2 | 31 | 1 | 32 | 1 |
| JB32 | 2 | 26 | 2 | 28 | 1 |
| JB32 | 2 | 36 | 0 | 36 | 1 |
| J32M | 2 | 32 | 4 | 36 | 2 |
| J32M | 2 | 33 | 3 | 36 | 2 |
| J32M | 2 | 35 | 1 | 36 | 2 |
| J32M | 2 | 36 | 0 | 36 | 2 |
| J32M | 2 | 31 | 1 | 32 | 2 |
| J32M | 2 | 33 | 3 | 36 | 2 |
| L968 | 3 | 8 | 0 | 8 | 1 |
| L968 | 3 | 47 | 13 | 60 | 1 |
| L968 | 3 | 53 | 7 | 60 | 1 |
| L968 | 3 | 43 | 13 | 56 | 1 |
| L968 | 3 | 59 | 5 | 64 | 1 |
| L968 | 3 | 30 | 6 | 36 | 1 |
| L968 | 3 | 25 | 7 | 32 | 1 |
| L968 | 3 | 27 | 9 | 36 | 1 |
| L968 | 3 | 26 | 6 | 32 | 1 |
| L968 | 3 | 29 | 7 | 36 | 1 |
| L968 | 3 | 34 | 2 | 36 | 1 |
| L968M | 3 | 32 | 0 | 32 | 2 |
| L968M | 3 | 28 | 0 | 28 | 2 |
| L968M | 3 | 34 | 2 | 36 | 2 |
| L968M | 3 | 29 | 3 | 32 | 2 |
| L968M | 3 | 27 | 1 | 28 | 2 |
| L968M | 3 | 31 | 1 | 32 | 2 |
| L968M | 3 | 20 | 0 | 20 | 2 |
| L968M | 3 | 34 | 2 | 36 | 2 |
| L968M | 3 | 27 | 1 | 28 | 2 |
| L968M | 3 | 36 | 0 | 36 | 2 |
| JB759 | 4 | 20 | 4 | 24 | 1 |
| JB759 | 4 | 18 | 6 | 24 | 1 |
| JB759 | 4 | 13 | 19 | 32 | 1 |
| JB759 | 4 | 20 | 12 | 32 | 1 |
| JB759 | 4 | 24 | 12 | 36 | 1 |
| JB759 | 4 | 16 | 16 | 32 | 1 |
| JB759 | 4 | 19 | 17 | 36 | 1 |
| JB759 | 4 | 12 | 20 | 32 | 1 |
| JB759 | 4 | 22 | 10 | 32 | 1 |
| JB759 | 4 | 10 | 18 | 28 | 1 |
| JB759 | 4 | 23 | 13 | 36 | 1 |
| JB759 | 4 | 18 | 14 | 32 | 1 |
| JB759 | 4 | 13 | 11 | 24 | 1 |
| JB759 | 4 | 21 | 15 | 36 | 1 |
| JB759M | 4 | 31 | 5 | 36 | 2 |
| JB759M | 4 | 29 | 7 | 36 | 2 |
| JB759M | 4 | 21 | 7 | 28 | 2 |
| JB759M | 4 | 29 | 3 | 32 | 2 |
| JB759M | 4 | 27 | 5 | 32 | 2 |
| JB759M | 4 | 21 | 11 | 32 | 2 |

| | | | | | |
|---|---|---|---|---|---|
| JB760 | 5 | 14 | 18 | 32 | 1 |
| JB760 | 5 | 14 | 18 | 32 | 1 |
| JB760 | 5 | 9 | 27 | 36 | 1 |
| JB760 | 5 | 9 | 27 | 36 | 1 |
| JB760 | 5 | 11 | 17 | 28 | 1 |
| JB760 | 5 | 11 | 21 | 32 | 1 |
| JB760 | 5 | 4 | 28 | 32 | 1 |
| JB760 | 5 | 9 | 27 | 36 | 1 |
| JB760 | 5 | 11 | 21 | 32 | 1 |
| JB760 | 5 | 18 | 18 | 36 | 1 |
| JB760 | 5 | 12 | 20 | 32 | 1 |
| JB760 | 5 | 13 | 23 | 36 | 1 |
| JB760M | 5 | 7 | 29 | 36 | 2 |
| JB760M | 5 | 5 | 31 | 36 | 2 |
| JB760M | 5 | 1 | 35 | 36 | 2 |
| JB760M | 5 | 6 | 30 | 36 | 2 |
| JB760M | 5 | 9 | 27 | 36 | 2 |
| JB760M | 5 | 5 | 27 | 32 | 2 |
| JB761 | 6 | 19 | 9 | 28 | 1 |
| JB761 | 6 | 16 | 4 | 20 | 1 |
| JB761 | 6 | 17 | 19 | 36 | 1 |
| JB761 | 6 | 15 | 13 | 28 | 1 |
| JB761 | 6 | 25 | 7 | 32 | 1 |
| JB761 | 6 | 15 | 13 | 28 | 1 |
| JB761M | 6 | 25 | 7 | 32 | 2 |
| JB761M | 6 | 27 | 9 | 36 | 2 |
| JB761M | 6 | 35 | 1 | 36 | 2 |
| JB761M | 6 | 21 | 15 | 36 | 2 |
| JB761M | 6 | 26 | 10 | 36 | 2 |
| JB761M | 6 | 26 | 10 | 36 | 2 |
| JB840 | 7 | 31 | 5 | 36 | 1 |
| JB840 | 7 | 29 | 3 | 32 | 1 |
| JB840 | 7 | 35 | 1 | 36 | 1 |
| JB840 | 7 | 30 | 2 | 32 | 1 |
| JB840 | 7 | 29 | 3 | 32 | 1 |
| JB840 | 7 | 29 | 3 | 32 | 1 |
| JB840M | 7 | 33 | 3 | 36 | 2 |
| JB840M | 7 | 34 | 2 | 36 | 2 |
| JB840M | 7 | 31 | 5 | 36 | 2 |
| JB840M | 7 | 17 | 7 | 24 | 2 |
| JB840M | 7 | 22 | 10 | 32 | 2 |
| JB840M | 7 | 19 | 17 | 36 | 2 |
| JB845 | 8 | 23 | 5 | 28 | 1 |
| JB845 | 8 | 23 | 5 | 28 | 1 |
| JB845 | 8 | 28 | 4 | 32 | 1 |
| JB845 | 8 | 24 | 12 | 36 | 1 |
| JB845 | 8 | 27 | 9 | 36 | 1 |
| JB845 | 8 | 21 | 15 | 36 | 1 |
| JB845M | 8 | 33 | 3 | 36 | 2 |
| JB845M | 8 | 31 | 1 | 32 | 2 |
| JB845M | 8 | 36 | 0 | 36 | 2 |
| JB845M | 8 | 13 | 3 | 16 | 2 |
| JB845M | 8 | 16 | 12 | 28 | 2 |
| JB845M | 8 | 25 | 7 | 32 | 2 |
| JB848 | 9 | 32 | 0 | 32 | 1 |
| JB848 | 9 | 32 | 0 | 32 | 1 |
| JB848 | 9 | 28 | 4 | 32 | 1 |
| JB848 | 9 | 23 | 9 | 32 | 1 |
| JB848 | 9 | 31 | 1 | 32 | 1 |
| JB848 | 9 | 27 | 5 | 32 | 1 |
| JB848M | 9 | 36 | 0 | 36 | 2 |
| JB848M | 9 | 32 | 4 | 36 | 2 |
| JB848M | 9 | 34 | 2 | 36 | 2 |
| JB848M | 9 | 15 | 13 | 28 | 2 |
| JB848M | 9 | 20 | 12 | 32 | 2 |
| JB848M | 9 | 23 | 5 | 28 | 2 |
| JB868 | 10 | 15 | 21 | 36 | 1 |
| JB868 | 10 | 16 | 20 | 36 | 1 |
| JB868 | 10 | 17 | 15 | 32 | 1 |
| JB868M | 10 | 19 | 1 | 20 | 2 |
| JB868M | 10 | 20 | 0 | 20 | 2 |
| JB868M | 10 | 16 | 0 | 16 | 2 |
| JB868M | 10 | 28 | 4 | 32 | 2 |
| JB868M | 10 | 32 | 0 | 32 | 2 |
| JB868M | 10 | 12 | 0 | 12 | 2 |
| JB870 | 11 | 29 | 3 | 32 | 1 |
| JB870 | 11 | 24 | 4 | 28 | 1 |
| JB870 | 11 | 36 | 0 | 36 | 1 |
| JB870 | 11 | 32 | 0 | 32 | 1 |
| JB870 | 11 | 35 | 1 | 36 | 1 |
| JB870 | 11 | 19 | 1 | 20 | 1 |
| JB870M | 11 | 34 | 2 | 36 | 2 |
| JB870M | 11 | 34 | 2 | 36 | 2 |

| | | | | | |
|---|---|---|---|---|---|
| JB870M | 11 | 27 | 1 | 28 | 2 |
| JB870M | 11 | 14 | 2 | 16 | 2 |
| JB870M | 11 | 26 | 6 | 32 | 2 |
| JB870M | 11 | 27 | 9 | 36 | 2 |
| JB873 | 12 | 31 | 5 | 36 | 1 |
| JB873 | 12 | 31 | 5 | 36 | 1 |
| JB873 | 12 | 22 | 10 | 32 | 1 |
| JB873 | 12 | 23 | 9 | 32 | 1 |
| JB873 | 12 | 16 | 8 | 24 | 1 |
| JB873 | 12 | 25 | 7 | 32 | 1 |
| JB873M | 12 | 29 | 3 | 32 | 2 |
| JB873M | 12 | 30 | 6 | 36 | 2 |
| JB873M | 12 | 25 | 7 | 32 | 2 |
| JB873M | 12 | 7 | 9 | 16 | 2 |
| JB873M | 12 | 14 | 6 | 20 | 2 |
| JB873M | 12 | 20 | 16 | 36 | 2 |
| JB906 | 13 | 31 | 1 | 32 | 1 |
| JB906 | 13 | 31 | 5 | 36 | 1 |
| JB906 | 13 | 28 | 0 | 28 | 1 |
| JB906M | 13 | 19 | 1 | 20 | 2 |
| JB906M | 13 | 28 | 0 | 28 | 2 |
| JB906M | 13 | 30 | 6 | 36 | 2 |
| JB906M | 13 | 28 | 0 | 28 | 2 |
| JB913 | 14 | 33 | 3 | 36 | 1 |
| JB913 | 14 | 36 | 0 | 36 | 1 |
| JB913 | 14 | 26 | 2 | 28 | 1 |
| JB913 | 14 | 31 | 1 | 32 | 1 |
| JB913 | 14 | 28 | 0 | 28 | 1 |
| JB913 | 14 | 33 | 3 | 36 | 1 |
| JB913M | 14 | 30 | 6 | 36 | 2 |
| JB913M | 14 | 27 | 5 | 32 | 2 |
| JB913M | 14 | 21 | 11 | 32 | 2 |
| JB913M | 14 | 24 | 8 | 32 | 2 |
| JB913M | 14 | 23 | 5 | 28 | 2 |
| JB913M | 14 | 18 | 14 | 32 | 2 |
| JB916 | 15 | 25 | 3 | 28 | 1 |
| JB916 | 15 | 30 | 2 | 32 | 1 |
| JB916 | 15 | 35 | 1 | 36 | 1 |
| JB916 | 15 | 22 | 6 | 28 | 1 |
| JB916 | 15 | 24 | 0 | 24 | 1 |
| JB916 | 15 | 25 | 3 | 28 | 1 |
| JB916M | 15 | 32 | 4 | 36 | 2 |
| JB916M | 15 | 24 | 12 | 36 | 2 |
| JB916M | 15 | 24 | 8 | 32 | 2 |
| JB916M | 15 | 16 | 12 | 28 | 2 |
| JB916M | 15 | 25 | 7 | 32 | 2 |
| JB916M | 15 | 23 | 5 | 28 | 2 |
| JB931 | 16 | 4 | 12 | 16 | 1 |
| JB931 | 16 | 17 | 11 | 28 | 1 |
| JB931 | 16 | 12 | 16 | 28 | 1 |
| JB931 | 16 | 33 | 3 | 36 | 1 |
| JB931 | 16 | 26 | 10 | 36 | 1 |
| JB931 | 16 | 18 | 2 | 20 | 1 |
| JB931M | 16 | 25 | 3 | 28 | 2 |
| JB931M | 16 | 23 | 1 | 24 | 2 |
| JB931M | 16 | 31 | 5 | 36 | 2 |
| JB937 | 17 | 23 | 9 | 32 | 1 |
| JB937 | 17 | 32 | 4 | 36 | 1 |
| JB937 | 17 | 26 | 10 | 36 | 1 |
| JB937M | 17 | 25 | 7 | 32 | 2 |
| JB937M | 17 | 29 | 7 | 36 | 2 |
| JB937M | 17 | 34 | 2 | 36 | 2 |
| JB937M | 17 | 32 | 4 | 36 | 2 |
| JB938 | 18 | 29 | 7 | 36 | 1 |
| JB938 | 18 | 29 | 7 | 36 | 1 |
| JB938 | 18 | 17 | 3 | 20 | 1 |
| JB938 | 18 | 28 | 4 | 32 | 1 |
| JB938 | 18 | 17 | 3 | 20 | 1 |
| JB938 | 18 | 31 | 1 | 32 | 1 |
| JB938M | 18 | 34 | 2 | 36 | 2 |
| JB938M | 18 | 10 | 14 | 24 | 2 |
| JB938M | 18 | 30 | 2 | 32 | 2 |
| JB938M | 18 | 24 | 4 | 28 | 2 |
| JB941 | 19 | 36 | 0 | 36 | 1 |
| JB941 | 19 | 32 | 4 | 36 | 1 |
| JB941 | 19 | 29 | 7 | 36 | 1 |
| JB941M | 19 | 33 | 3 | 36 | 2 |
| JB941M | 19 | 30 | 2 | 32 | 2 |
| JB941M | 19 | 28 | 0 | 28 | 2 |
| JB941M | 19 | 20 | 12 | 32 | 2 |
| JB953 | 20 | 13 | 3 | 16 | 1 |
| JB953 | 20 | 19 | 9 | 28 | 1 |
| JB953 | 20 | 19 | 17 | 36 | 1 |

| | | | | | |
|---|---|---|---|---|---|
| JB953 | 20 | 14 | 14 | 28 | 1 |
| JB953 | 20 | 18 | 10 | 28 | 1 |
| JB953 | 20 | 19 | 9 | 28 | 1 |
| JB953M | 20 | 21 | 15 | 36 | 2 |
| JB953M | 20 | 14 | 14 | 28 | 2 |
| JB953M | 20 | 27 | 5 | 32 | 2 |
| JB953M | 20 | 12 | 24 | 36 | 2 |
| JB953M | 20 | 9 | 19 | 28 | 2 |
| JB953M | 20 | 17 | 15 | 32 | 2 |
| JB1114 | 21 | 20 | 16 | 36 | 1 |
| JB1114 | 21 | 24 | 8 | 32 | 1 |
| JB1114 | 21 | 20 | 16 | 36 | 1 |
| JB1114 | 21 | 25 | 7 | 32 | 1 |
| JB1114 | 21 | 31 | 5 | 36 | 1 |
| JB1114 | 21 | 22 | 10 | 32 | 1 |
| JB1114M | 21 | 21 | 7 | 28 | 2 |
| JB1114M | 21 | 26 | 10 | 36 | 2 |
| JB1114M | 21 | 26 | 6 | 32 | 2 |
| JB1114M | 21 | 8 | 4 | 12 | 2 |
| JB1114M | 21 | 11 | 9 | 20 | 2 |
| JB1114M | 21 | 21 | 11 | 32 | 2 |
| JB1205 | 22 | 34 | 2 | 36 | 1 |
| JB1205 | 22 | 19 | 1 | 20 | 1 |
| JB1205 | 22 | 17 | 11 | 28 | 1 |
| JB1205 | 22 | 32 | 4 | 36 | 1 |
| JB1205 | 22 | 19 | 13 | 32 | 1 |
| JB1205 | 22 | 20 | 4 | 24 | 1 |
| JB1205M | 22 | 33 | 3 | 36 | 2 |
| JB1205M | 22 | 35 | 1 | 36 | 2 |
| JB1205M | 22 | 31 | 1 | 32 | 2 |
| JB1205M | 22 | 23 | 9 | 32 | 2 |
| JB1205M | 22 | 22 | 6 | 28 | 2 |
| JB1205M | 22 | 24 | 0 | 24 | 2 |

| term | estimate | std.error | statistic | p.value |
|---|---|---|---|---|
| **(Intercept)** | -1,198624087 | 0,075831523 | -15,80640915 | 2,81E-56 |
| **Zygotic-Azygotic** | -7,16597E-05 | 0,050392157 | -0,001422041 | 0,998865376 |

| Experiment | Strain name | Group | Viable | No_viable | Total | Zygotic-azygotic |
|---|---|---|---|---|---|---|
| One Way ANOVA | L972(JB22) | 1 | 24 | 12 | 36 | 1 |
| Zygotic vs azygotic | L972(JB22) | 1 | 32 | 4 | 36 | 1 |
| No 4C data | L972(JB22) | 1 | 34 | 2 | 36 | 1 |
| (Suplemental Figure 3B) | L972(JB22) | 1 | 27 | 5 | 32 | 1 |
| | L972(JB22) | 1 | 32 | 4 | 36 | 1 |
| | L972(JB22) | 1 | 27 | 5 | 32 | 1 |
| | L972(JB22) | 1 | 12 | 0 | 12 | 1 |
| | L972(JB22) | 1 | 8 | 0 | 8 | 1 |
| | L972(JB22) | 1 | 22 | 10 | 32 | 1 |
| | L972(JB22) | 1 | 21 | 7 | 28 | 1 |
| | L972(JB22) | 1 | 35 | 1 | 36 | 1 |
| | L972(JB22) | 1 | 28 | 0 | 28 | 1 |
| | L972(JB903) | 1 | 32 | 4 | 36 | 1 |
| | L972(JB903) | 1 | 30 | 6 | 36 | 1 |
| | L972(JB903) | 1 | 31 | 5 | 36 | 1 |
| | L972(JB903) | 1 | 28 | 0 | 28 | 1 |
| | L972(JB903) | 1 | 32 | 0 | 32 | 1 |
| | L972(JB903) | 1 | 27 | 5 | 32 | 1 |
| | L972(JB903) | 1 | 32 | 4 | 36 | 1 |
| | L972(JB903) | 1 | 28 | 4 | 32 | 1 |
| | L972(JB903) | 1 | 19 | 13 | 32 | 1 |
| | L972(JB903) | 1 | 29 | 3 | 32 | 1 |
| | L972(JB903) | 1 | 25 | 7 | 32 | 1 |
| | L972(JB903) | 1 | 27 | 5 | 32 | 1 |
| | L972M(JB22M) | 1 | 8 | 4 | 12 | 2 |
| | L972M(JB22M) | 1 | 7 | 13 | 20 | 2 |
| | L972M(JB22M) | 1 | 4 | 20 | 24 | 2 |
| | L972M(JB22M) | 1 | 2 | 14 | 16 | 2 |
| | L972M(JB22M) | 1 | 3 | 17 | 20 | 2 |
| | L972M(JB22M) | 1 | 0 | 16 | 16 | 2 |
| | JB32 | 2 | 36 | 0 | 36 | 1 |
| | JB32 | 2 | 35 | 1 | 36 | 1 |
| | JB32 | 2 | 34 | 2 | 36 | 1 |
| | JB32 | 2 | 29 | 3 | 32 | 1 |
| | JB32 | 2 | 36 | 0 | 36 | 1 |
| | JB32 | 2 | 30 | 2 | 32 | 1 |
| | JB32 | 2 | 26 | 2 | 28 | 1 |
| | JB32 | 2 | 31 | 5 | 36 | 1 |
| | JB32 | 2 | 33 | 3 | 36 | 1 |
| | JB32 | 2 | 31 | 1 | 32 | 1 |
| | JB32 | 2 | 26 | 2 | 28 | 1 |
| | JB32 | 2 | 36 | 0 | 36 | 1 |
| | J32M | 2 | 32 | 4 | 36 | 2 |
| | J32M | 2 | 33 | 3 | 36 | 2 |

| | | | | | |
|---|---|---|---|---|---|
| J32M | 2 | 35 | 1 | 36 | 2 |
| J32M | 2 | 36 | 0 | 36 | 2 |
| J32M | 2 | 31 | 1 | 32 | 2 |
| J32M | 2 | 33 | 3 | 36 | 2 |
| L968 | 3 | 8 | 0 | 8 | 1 |
| L968 | 3 | 47 | 13 | 60 | 1 |
| L968 | 3 | 53 | 7 | 60 | 1 |
| L968 | 3 | 43 | 13 | 56 | 1 |
| L968 | 3 | 59 | 5 | 64 | 1 |
| L968 | 3 | 30 | 6 | 36 | 1 |
| L968 | 3 | 25 | 7 | 32 | 1 |
| L968 | 3 | 27 | 9 | 36 | 1 |
| L968 | 3 | 26 | 6 | 32 | 1 |
| L968 | 3 | 29 | 7 | 36 | 1 |
| L968 | 3 | 34 | 2 | 36 | 1 |
| L968M | 3 | 32 | 0 | 32 | 2 |
| L968M | 3 | 28 | 0 | 28 | 2 |
| L968M | 3 | 34 | 2 | 36 | 2 |
| L968M | 3 | 29 | 3 | 32 | 2 |
| L968M | 3 | 27 | 1 | 28 | 2 |
| L968M | 3 | 31 | 1 | 32 | 2 |
| L968M | 3 | 20 | 0 | 20 | 2 |
| L968M | 3 | 34 | 2 | 36 | 2 |
| L968M | 3 | 27 | 1 | 28 | 2 |
| L968M | 3 | 36 | 0 | 36 | 2 |
| JB759 | 4 | 20 | 4 | 24 | 1 |
| JB759 | 4 | 18 | 6 | 24 | 1 |
| JB759 | 4 | 13 | 19 | 32 | 1 |
| JB759 | 4 | 20 | 12 | 32 | 1 |
| JB759 | 4 | 24 | 12 | 36 | 1 |
| JB759 | 4 | 16 | 16 | 32 | 1 |
| JB759 | 4 | 19 | 17 | 36 | 1 |
| JB759 | 4 | 12 | 20 | 32 | 1 |
| JB759 | 4 | 22 | 10 | 32 | 1 |
| JB759 | 4 | 10 | 18 | 28 | 1 |
| JB759 | 4 | 23 | 13 | 36 | 1 |
| JB759 | 4 | 18 | 14 | 32 | 1 |
| JB759 | 4 | 13 | 11 | 24 | 1 |
| JB759 | 4 | 21 | 15 | 36 | 1 |
| JB759M | 4 | 31 | 5 | 36 | 2 |
| JB759M | 4 | 29 | 7 | 36 | 2 |
| JB759M | 4 | 21 | 7 | 28 | 2 |
| JB759M | 4 | 29 | 3 | 32 | 2 |
| JB759M | 4 | 27 | 5 | 32 | 2 |
| JB759M | 4 | 21 | 11 | 32 | 2 |
| JB760 | 5 | 14 | 18 | 32 | 1 |
| JB760 | 5 | 14 | 18 | 32 | 1 |
| JB760 | 5 | 9 | 27 | 36 | 1 |
| JB760 | 5 | 9 | 27 | 36 | 1 |
| JB760 | 5 | 11 | 17 | 28 | 1 |
| JB760 | 5 | 11 | 21 | 32 | 1 |
| JB760 | 5 | 4 | 28 | 32 | 1 |
| JB760 | 5 | 9 | 27 | 36 | 1 |
| JB760 | 5 | 11 | 21 | 32 | 1 |
| JB760 | 5 | 18 | 18 | 36 | 1 |
| JB760 | 5 | 12 | 20 | 32 | 1 |
| JB760 | 5 | 13 | 23 | 36 | 1 |
| JB760M | 5 | 7 | 29 | 36 | 2 |
| JB760M | 5 | 5 | 31 | 36 | 2 |
| JB760M | 5 | 1 | 35 | 36 | 2 |
| JB760M | 5 | 6 | 30 | 36 | 2 |
| JB760M | 5 | 9 | 27 | 36 | 2 |
| JB760M | 5 | 5 | 27 | 32 | 2 |
| JB761 | 6 | 19 | 9 | 28 | 1 |
| JB761 | 6 | 16 | 4 | 20 | 1 |
| JB761 | 6 | 17 | 19 | 36 | 1 |
| JB761 | 6 | 15 | 13 | 28 | 1 |
| JB761 | 6 | 25 | 7 | 32 | 1 |
| JB761 | 6 | 15 | 13 | 28 | 1 |
| JB761M | 6 | 25 | 7 | 32 | 2 |
| JB761M | 6 | 27 | 9 | 36 | 2 |
| JB761M | 6 | 35 | 1 | 36 | 2 |
| JB761M | 6 | 21 | 15 | 36 | 2 |
| JB761M | 6 | 26 | 10 | 36 | 2 |
| JB761M | 6 | 26 | 10 | 36 | 2 |
| JB840 | 7 | 31 | 5 | 36 | 1 |
| JB840 | 7 | 29 | 3 | 32 | 1 |
| JB840 | 7 | 35 | 1 | 36 | 1 |
| JB840 | 7 | 30 | 2 | 32 | 1 |
| JB840 | 7 | 29 | 3 | 32 | 1 |
| JB840 | 7 | 29 | 3 | 32 | 1 |
| JB840M | 7 | 33 | 3 | 36 | 2 |
| JB840M | 7 | 34 | 2 | 36 | 2 |

| | | | | | |
|---|---|---|---|---|---|
| JB840M | 7 | 31 | 5 | 36 | 2 |
| JB845 | 8 | 23 | 5 | 28 | 1 |
| JB845 | 8 | 23 | 5 | 28 | 1 |
| JB845 | 8 | 28 | 4 | 32 | 1 |
| JB845 | 8 | 24 | 12 | 36 | 1 |
| JB845 | 8 | 27 | 9 | 36 | 1 |
| JB845 | 8 | 21 | 15 | 36 | 1 |
| JB845M | 8 | 33 | 3 | 36 | 2 |
| JB845M | 8 | 31 | 1 | 32 | 2 |
| JB845M | 8 | 36 | 0 | 36 | 2 |
| JB848 | 9 | 32 | 0 | 32 | 1 |
| JB848 | 9 | 32 | 0 | 32 | 1 |
| JB848 | 9 | 28 | 4 | 32 | 1 |
| JB848 | 9 | 23 | 9 | 32 | 1 |
| JB848 | 9 | 31 | 1 | 32 | 1 |
| JB848 | 9 | 27 | 5 | 32 | 1 |
| JB848M | 9 | 36 | 0 | 36 | 2 |
| JB848M | 9 | 32 | 4 | 36 | 2 |
| JB848M | 9 | 34 | 2 | 36 | 2 |
| JB868 | 10 | 15 | 21 | 36 | 1 |
| JB868 | 10 | 16 | 20 | 36 | 1 |
| JB868 | 10 | 17 | 15 | 32 | 1 |
| JB868M | 10 | 19 | 1 | 20 | 2 |
| JB868M | 10 | 20 | 0 | 20 | 2 |
| JB868M | 10 | 16 | 0 | 16 | 2 |
| JB868M | 10 | 28 | 4 | 32 | 2 |
| JB868M | 10 | 32 | 0 | 32 | 2 |
| JB868M | 10 | 12 | 0 | 12 | 2 |
| JB870 | 11 | 29 | 3 | 32 | 1 |
| JB870 | 11 | 24 | 4 | 28 | 1 |
| JB870 | 11 | 36 | 0 | 36 | 1 |
| JB870 | 11 | 32 | 0 | 32 | 1 |
| JB870 | 11 | 35 | 1 | 36 | 1 |
| JB870 | 11 | 19 | 1 | 20 | 1 |
| JB870M | 11 | 34 | 2 | 36 | 2 |
| JB870M | 11 | 34 | 2 | 36 | 2 |
| JB870M | 11 | 27 | 1 | 28 | 2 |
| JB873 | 12 | 31 | 5 | 36 | 1 |
| JB873 | 12 | 31 | 5 | 36 | 1 |
| JB873 | 12 | 22 | 10 | 32 | 1 |
| JB873 | 12 | 23 | 9 | 32 | 1 |
| JB873 | 12 | 16 | 8 | 24 | 1 |
| JB873 | 12 | 25 | 7 | 32 | 1 |
| JB873M | 12 | 29 | 3 | 32 | 2 |
| JB873M | 12 | 30 | 6 | 36 | 2 |
| JB873M | 12 | 25 | 7 | 32 | 2 |
| JB906 | 13 | 31 | 1 | 32 | 1 |
| JB906 | 13 | 31 | 5 | 36 | 1 |
| JB906 | 13 | 28 | 0 | 28 | 1 |
| JB906M | 13 | 19 | 1 | 20 | 2 |
| JB906M | 13 | 28 | 0 | 28 | 2 |
| JB906M | 13 | 30 | 6 | 36 | 2 |
| JB906M | 13 | 28 | 0 | 28 | 2 |
| JB913 | 14 | 33 | 3 | 36 | 1 |
| JB913 | 14 | 36 | 0 | 36 | 1 |
| JB913 | 14 | 26 | 2 | 28 | 1 |
| JB913 | 14 | 31 | 1 | 32 | 1 |
| JB913 | 14 | 28 | 0 | 28 | 1 |
| JB913 | 14 | 33 | 3 | 36 | 1 |
| JB913M | 14 | 30 | 6 | 36 | 2 |
| JB913M | 14 | 27 | 5 | 32 | 2 |
| JB913M | 14 | 21 | 11 | 32 | 2 |
| JB916 | 15 | 25 | 3 | 28 | 1 |
| JB916 | 15 | 30 | 2 | 32 | 1 |
| JB916 | 15 | 35 | 1 | 36 | 1 |
| JB916 | 15 | 22 | 6 | 28 | 1 |
| JB916 | 15 | 24 | 0 | 24 | 1 |
| JB916 | 15 | 25 | 3 | 28 | 1 |
| JB916M | 15 | 32 | 4 | 36 | 2 |
| JB916M | 15 | 24 | 12 | 36 | 2 |
| JB916M | 15 | 24 | 8 | 32 | 2 |
| JB931 | 16 | 4 | 12 | 16 | 1 |
| JB931 | 16 | 17 | 11 | 28 | 1 |
| JB931 | 16 | 12 | 16 | 28 | 1 |
| JB931 | 16 | 33 | 3 | 36 | 1 |
| JB931 | 16 | 26 | 10 | 36 | 1 |
| JB931 | 16 | 18 | 2 | 20 | 1 |
| JB931M | 16 | 25 | 3 | 28 | 2 |
| JB931M | 16 | 23 | 1 | 24 | 2 |
| JB931M | 16 | 31 | 5 | 36 | 2 |
| JB937 | 17 | 23 | 9 | 32 | 1 |
| JB937 | 17 | 32 | 4 | 36 | 1 |
| JB937 | 17 | 26 | 10 | 36 | 1 |

| | | | | | |
|---|---|---|---|---|---|
| JB937M | 17 | 25 | 7 | 32 | 2 |
| JB937M | 17 | 29 | 7 | 36 | 2 |
| JB937M | 17 | 34 | 2 | 36 | 2 |
| JB937M | 17 | 32 | 4 | 36 | 2 |
| JB938 | 18 | 29 | 7 | 36 | 1 |
| JB938 | 18 | 29 | 7 | 36 | 1 |
| JB938 | 18 | 17 | 3 | 20 | 1 |
| JB938 | 18 | 28 | 4 | 32 | 1 |
| JB938 | 18 | 17 | 3 | 20 | 1 |
| JB938 | 18 | 31 | 1 | 32 | 1 |
| JB938M | 18 | 34 | 2 | 36 | 2 |
| JB941 | 19 | 36 | 0 | 36 | 1 |
| JB941 | 19 | 32 | 4 | 36 | 1 |
| JB941 | 19 | 29 | 7 | 36 | 1 |
| JB941M | 19 | 33 | 3 | 36 | 2 |
| JB941M | 19 | 30 | 2 | 32 | 2 |
| JB941M | 19 | 28 | 0 | 28 | 2 |
| JB941M | 19 | 20 | 12 | 32 | 2 |
| JB953 | 20 | 13 | 3 | 16 | 1 |
| JB953 | 20 | 19 | 9 | 28 | 1 |
| JB953 | 20 | 19 | 17 | 36 | 1 |
| JB953M | 20 | 21 | 15 | 36 | 2 |
| JB953M | 20 | 14 | 14 | 28 | 2 |
| JB953M | 20 | 27 | 5 | 32 | 2 |
| JB1114 | 21 | 20 | 16 | 36 | 1 |
| JB1114 | 21 | 24 | 8 | 32 | 1 |
| JB1114 | 21 | 20 | 16 | 36 | 1 |
| JB1114 | 21 | 25 | 7 | 32 | 1 |
| JB1114 | 21 | 31 | 5 | 36 | 1 |
| JB1114 | 21 | 22 | 10 | 32 | 1 |
| JB1114M | 21 | 21 | 7 | 28 | 2 |
| JB1114M | 21 | 26 | 10 | 36 | 2 |
| JB1114M | 21 | 26 | 6 | 32 | 2 |
| JB1205 | 22 | 34 | 2 | 36 | 1 |
| JB1205 | 22 | 19 | 1 | 20 | 1 |
| JB1205M | 22 | 33 | 3 | 36 | 2 |
| JB1205M | 22 | 35 | 1 | 36 | 2 |
| JB1205M | 22 | 31 | 1 | 32 | 2 |

| term | estimate | std.error | statistic | p.value |
|---|---|---|---|---|
| **(Intercept)** | -1,057676394 | 0,081522449 | -12,97405074 | 1,72E-38 |
| **Zygotic-Azygotic** | -0,160872544 | 0,057175062 | -2,81368379 | 0,004897738 |

| Experiment | Strain name | Group | Viable | No_viable | Total | Zygotic-azigotic | 4C Batch | Other Batches |
|---|---|---|---|---|---|---|---|---|
| Multifactorial ANOVA | L972(JB22) | 1 | 12 | 24 | 36 | 1 | 1 | 1 |
| Zygotic vs azygotic | L972(JB22) | 1 | 4 | 32 | 36 | 1 | 1 | 1 |
| Batched samples | L972(JB22) | 1 | 2 | 34 | 36 | 1 | 1 | 1 |
| (Suplemental Figure 3E) | L972(JB22) | 1 | 5 | 27 | 32 | 1 | 1 | 3 |
| | L972(JB22) | 1 | 4 | 32 | 36 | 1 | 1 | 3 |
| | L972(JB22) | 1 | 5 | 27 | 32 | 1 | 1 | 3 |
| | L972(JB22) | 1 | 0 | 12 | 12 | 1 | 1 | 1 |
| | L972(JB22) | 1 | 0 | 8 | 8 | 1 | 1 | 1 |
| | L972(JB22) | 1 | 10 | 22 | 32 | 1 | 1 | 1 |
| | L972(JB22) | 1 | 7 | 21 | 28 | 1 | 1 | 1 |
| | L972(JB22) | 1 | 1 | 35 | 36 | 1 | 1 | 1 |
| | L972(JB22) | 1 | 0 | 28 | 28 | 1 | 1 | 1 |
| | L972(JB903) | 1 | 32 | 4 | 36 | 1 | 1 | 1 |
| | L972(JB903) | 1 | 30 | 6 | 36 | 1 | 1 | 1 |
| | L972(JB903) | 1 | 31 | 5 | 36 | 1 | 1 | 1 |
| | L972(JB903) | 1 | 28 | 0 | 28 | 1 | 1 | 1 |
| | L972(JB903) | 1 | 32 | 0 | 32 | 1 | 1 | 1 |
| | L972(JB903) | 1 | 27 | 5 | 32 | 1 | 1 | 1 |
| | L972(JB903) | 1 | 32 | 4 | 36 | 1 | 1 | 1 |
| | L972(JB903) | 1 | 28 | 4 | 32 | 1 | 1 | 1 |
| | L972(JB903) | 1 | 19 | 13 | 32 | 1 | 1 | 1 |
| | L972(JB903) | 1 | 29 | 3 | 32 | 1 | 1 | 1 |
| | L972(JB903) | 1 | 25 | 7 | 32 | 1 | 1 | 1 |
| | L972(JB903) | 1 | 27 | 5 | 32 | 1 | 1 | 1 |
| | L972M(JB22M) | 1 | 4 | 8 | 12 | 2 | 1 | 1 |
| | L972M(JB22M) | 1 | 13 | 7 | 20 | 2 | 1 | 1 |
| | L972M(JB22M) | 1 | 20 | 4 | 24 | 2 | 1 | 1 |
| | L972M(JB22M) | 1 | 14 | 2 | 16 | 2 | 1 | 1 |
| | L972M(JB22M) | 1 | 17 | 3 | 20 | 2 | 1 | 1 |
| | L972M(JB22M) | 1 | 16 | 0 | 16 | 2 | 1 | 1 |
| | JB32 | 2 | 0 | 36 | 36 | 1 | 1 | 1 |
| | JB32 | 2 | 1 | 35 | 36 | 1 | 1 | 1 |
| | JB32 | 2 | 2 | 34 | 36 | 1 | 1 | 1 |
| | JB32 | 2 | 3 | 29 | 32 | 1 | 1 | 3 |
| | JB32 | 2 | 0 | 36 | 36 | 1 | 1 | 3 |
| | JB32 | 2 | 2 | 30 | 32 | 1 | 1 | 3 |
| | JB32 | 2 | 2 | 26 | 28 | 1 | 1 | 1 |
| | JB32 | 2 | 5 | 31 | 36 | 1 | 1 | 1 |
| | JB32 | 2 | 3 | 33 | 36 | 1 | 1 | 1 |

| | | | | | | | |
|---|---|---|---|---|---|---|---|
| JB32 | 2 | 1 | 31 | 32 | 1 | 1 | 1 |
| JB32 | 2 | 2 | 26 | 28 | 1 | 1 | 1 |
| JB32 | 2 | 0 | 36 | 36 | 1 | 1 | 1 |
| J32M | 2 | 4 | 32 | 36 | 2 | 1 | 1 |
| J32M | 2 | 3 | 33 | 36 | 2 | 1 | 1 |
| J32M | 2 | 1 | 35 | 36 | 2 | 1 | 1 |
| J32M | 2 | 0 | 36 | 36 | 2 | 1 | 1 |
| J32M | 2 | 1 | 31 | 32 | 2 | 1 | 1 |
| J32M | 2 | 3 | 33 | 36 | 2 | 1 | 1 |
| JB840 | 3 | 5 | 31 | 36 | 1 | 1 | 1 |
| JB840 | 3 | 3 | 29 | 32 | 1 | 1 | 1 |
| JB840 | 3 | 1 | 35 | 36 | 1 | 1 | 1 |
| JB840 | 3 | 2 | 30 | 32 | 1 | 1 | 3 |
| JB840 | 3 | 3 | 29 | 32 | 1 | 1 | 3 |
| JB840 | 3 | 3 | 29 | 32 | 1 | 1 | 3 |
| JB840M | 3 | 3 | 33 | 36 | 2 | 1 | 1 |
| JB840M | 3 | 2 | 34 | 36 | 2 | 1 | 1 |
| JB840M | 3 | 5 | 31 | 36 | 2 | 1 | 1 |
| JB840M | 3 | 7 | 17 | 24 | 2 | 2 | 1 |
| JB840M | 3 | 10 | 22 | 32 | 2 | 2 | 1 |
| JB840M | 3 | 17 | 19 | 36 | 2 | 2 | 1 |
| JB845 | 4 | 5 | 23 | 28 | 1 | 1 | 1 |
| JB845 | 4 | 5 | 23 | 28 | 1 | 1 | 1 |
| JB845 | 4 | 4 | 28 | 32 | 1 | 1 | 1 |
| JB845 | 4 | 12 | 24 | 36 | 1 | 1 | 3 |
| JB845 | 4 | 9 | 27 | 36 | 1 | 1 | 3 |
| JB845 | 4 | 15 | 21 | 36 | 1 | 1 | 3 |
| JB845M | 4 | 3 | 33 | 36 | 2 | 1 | 1 |
| JB845M | 4 | 1 | 31 | 32 | 2 | 1 | 1 |
| JB845M | 4 | 0 | 36 | 36 | 2 | 1 | 1 |
| JB845M | 4 | 3 | 13 | 16 | 2 | 2 | 1 |
| JB845M | 4 | 12 | 16 | 28 | 2 | 2 | 1 |
| JB845M | 4 | 7 | 25 | 32 | 2 | 2 | 1 |
| JB848 | 5 | 0 | 32 | 32 | 1 | 1 | 1 |
| JB848 | 5 | 0 | 32 | 32 | 1 | 1 | 1 |
| JB848 | 5 | 4 | 28 | 32 | 1 | 1 | 1 |
| JB848 | 5 | 9 | 23 | 32 | 1 | 1 | 3 |
| JB848 | 5 | 1 | 31 | 32 | 1 | 1 | 3 |
| JB848 | 5 | 5 | 27 | 32 | 1 | 1 | 3 |
| JB848M | 5 | 0 | 36 | 36 | 2 | 1 | 1 |
| JB848M | 5 | 4 | 32 | 36 | 2 | 1 | 1 |
| JB848M | 5 | 2 | 34 | 36 | 2 | 1 | 1 |
| JB848M | 5 | 13 | 15 | 28 | 2 | 2 | 1 |
| JB848M | 5 | 12 | 20 | 32 | 2 | 2 | 1 |
| JB848M | 5 | 5 | 23 | 28 | 2 | 2 | 1 |
| JB870 | 6 | 3 | 29 | 32 | 1 | 1 | 1 |
| JB870 | 6 | 4 | 24 | 28 | 1 | 1 | 1 |
| JB870 | 6 | 0 | 36 | 36 | 1 | 1 | 1 |
| JB870 | 6 | 0 | 32 | 32 | 1 | 1 | 3 |
| JB870 | 6 | 1 | 35 | 36 | 1 | 1 | 3 |
| JB870 | 6 | 1 | 19 | 20 | 1 | 1 | 3 |
| JB870M | 6 | 2 | 34 | 36 | 2 | 1 | 1 |
| JB870M | 6 | 2 | 34 | 36 | 2 | 1 | 1 |
| JB870M | 6 | 1 | 27 | 28 | 2 | 1 | 1 |
| JB870M | 6 | 2 | 14 | 16 | 2 | 2 | 1 |
| JB870M | 6 | 6 | 26 | 32 | 2 | 2 | 1 |
| JB870M | 6 | 9 | 27 | 36 | 2 | 2 | 1 |
| JB873 | 7 | 5 | 31 | 36 | 1 | 1 | 1 |
| JB873 | 7 | 5 | 31 | 36 | 1 | 1 | 1 |
| JB873 | 7 | 10 | 22 | 32 | 1 | 1 | 1 |
| JB873 | 7 | 9 | 23 | 32 | 1 | 1 | 3 |
| JB873 | 7 | 8 | 16 | 24 | 1 | 1 | 3 |
| JB873 | 7 | 7 | 25 | 32 | 1 | 1 | 3 |
| JB873M | 7 | 3 | 29 | 32 | 2 | 1 | 1 |
| JB873M | 7 | 6 | 30 | 36 | 2 | 1 | 1 |
| JB873M | 7 | 7 | 25 | 32 | 2 | 1 | 1 |
| JB873M | 7 | 9 | 7 | 16 | 2 | 2 | 1 |
| JB873M | 7 | 6 | 14 | 20 | 2 | 2 | 1 |
| JB873M | 7 | 16 | 20 | 36 | 2 | 2 | 1 |
| JB913 | 8 | 3 | 33 | 36 | 1 | 1 | 1 |
| JB913 | 8 | 0 | 36 | 36 | 1 | 1 | 1 |
| JB913 | 8 | 2 | 26 | 28 | 1 | 1 | 1 |
| JB913 | 8 | 1 | 31 | 32 | 1 | 1 | 3 |
| JB913 | 8 | 0 | 28 | 28 | 1 | 1 | 3 |
| JB913 | 8 | 3 | 33 | 36 | 1 | 1 | 3 |
| JB913M | 8 | 6 | 30 | 36 | 2 | 1 | 1 |
| JB913M | 8 | 5 | 27 | 32 | 2 | 1 | 1 |
| JB913M | 8 | 11 | 21 | 32 | 2 | 1 | 1 |
| JB913M | 8 | 8 | 24 | 32 | 2 | 2 | 1 |
| JB913M | 8 | 5 | 23 | 28 | 2 | 2 | 1 |
| JB913M | 8 | 14 | 18 | 32 | 2 | 2 | 1 |
| JB916 | 9 | 3 | 25 | 28 | 1 | 1 | 1 |
| JB916 | 9 | 2 | 30 | 32 | 1 | 1 | 1 |

| | | | | | | | |
|---|---|---|---|---|---|---|---|
| JB916 | 9 | 1 | 35 | 36 | 1 | 1 | 1 |
| JB916 | 9 | 6 | 22 | 28 | 1 | 1 | 3 |
| JB916 | 9 | 0 | 24 | 24 | 1 | 1 | 3 |
| JB916 | 9 | 3 | 25 | 28 | 1 | 1 | 3 |
| JB916M | 9 | 4 | 32 | 36 | 2 | 1 | 1 |
| JB916M | 9 | 12 | 24 | 36 | 2 | 1 | 1 |
| JB916M | 9 | 8 | 24 | 32 | 2 | 1 | 1 |
| JB916M | 9 | 12 | 16 | 28 | 2 | 2 | 1 |
| JB916M | 9 | 7 | 25 | 32 | 2 | 2 | 1 |
| JB916M | 9 | 5 | 23 | 28 | 2 | 2 | 1 |
| JB931 | 10 | 12 | 4 | 16 | 1 | 1 | 1 |
| JB931 | 10 | 11 | 17 | 28 | 1 | 1 | 1 |
| JB931 | 10 | 16 | 12 | 28 | 1 | 1 | 1 |
| JB931 | 10 | 3 | 33 | 36 | 1 | 1 | 3 |
| JB931 | 10 | 10 | 26 | 36 | 1 | 1 | 3 |
| JB931 | 10 | 2 | 18 | 20 | 1 | 1 | 3 |
| JB931M | 10 | 3 | 25 | 28 | 2 | 1 | 1 |
| JB931M | 10 | 1 | 23 | 24 | 2 | 1 | 1 |
| JB931M | 10 | 5 | 31 | 36 | 2 | 1 | 1 |
| JB938 | 11 | 7 | 29 | 36 | 1 | 1 | 1 |
| JB938 | 11 | 7 | 29 | 36 | 1 | 1 | 1 |
| JB938 | 11 | 3 | 17 | 20 | 1 | 1 | 1 |
| JB938 | 11 | 4 | 28 | 32 | 1 | 1 | 3 |
| JB938 | 11 | 3 | 17 | 20 | 1 | 1 | 3 |
| JB938 | 11 | 1 | 31 | 32 | 1 | 1 | 3 |
| JB938M | 11 | 2 | 34 | 36 | 2 | 1 | 1 |
| JB938M | 11 | 14 | 10 | 24 | 2 | 2 | 1 |
| JB938M | 11 | 2 | 30 | 32 | 2 | 2 | 1 |
| JB938M | 11 | 4 | 24 | 28 | 2 | 2 | 1 |
| JB953 | 12 | 3 | 13 | 16 | 1 | 1 | 1 |
| JB953 | 12 | 9 | 19 | 28 | 1 | 1 | 1 |
| JB953 | 12 | 17 | 19 | 36 | 1 | 1 | 1 |
| JB953 | 12 | 14 | 14 | 28 | 1 | 2 | 1 |
| JB953 | 12 | 10 | 18 | 28 | 1 | 2 | 1 |
| JB953 | 12 | 9 | 19 | 28 | 1 | 2 | 1 |
| JB953M | 12 | 15 | 21 | 36 | 2 | 1 | 1 |
| JB953M | 12 | 14 | 14 | 28 | 2 | 1 | 1 |
| JB953M | 12 | 5 | 27 | 32 | 2 | 1 | 1 |
| JB953M | 12 | 24 | 12 | 36 | 2 | 2 | 1 |
| JB953M | 12 | 19 | 9 | 28 | 2 | 2 | 1 |
| JB953M | 12 | 15 | 17 | 32 | 2 | 2 | 1 |
| JB1114 | 13 | 16 | 20 | 36 | 1 | 1 | 1 |
| JB1114 | 13 | 8 | 24 | 32 | 1 | 1 | 1 |
| JB1114 | 13 | 16 | 20 | 36 | 1 | 1 | 1 |
| JB1114 | 13 | 7 | 25 | 32 | 1 | 1 | 3 |
| JB1114 | 13 | 5 | 31 | 36 | 1 | 1 | 3 |
| JB1114 | 13 | 10 | 22 | 32 | 1 | 1 | 3 |
| JB1114M | 13 | 7 | 21 | 28 | 2 | 1 | 1 |
| JB1114M | 13 | 10 | 26 | 36 | 2 | 1 | 1 |
| JB1114M | 13 | 6 | 26 | 32 | 2 | 1 | 1 |
| JB1114M | 13 | 4 | 8 | 12 | 2 | 2 | 1 |
| JB1114M | 13 | 9 | 11 | 20 | 2 | 2 | 1 |
| JB1114M | 13 | 11 | 21 | 32 | 2 | 2 | 1 |
| JB1205 | 14 | 2 | 34 | 36 | 1 | 1 | 1 |
| JB1205 | 14 | 1 | 19 | 20 | 1 | 1 | 1 |
| JB1205 | 14 | 11 | 17 | 28 | 1 | 2 | 1 |
| JB1205 | 14 | 4 | 32 | 36 | 1 | 2 | 1 |
| JB1205 | 14 | 13 | 19 | 32 | 1 | 2 | 1 |
| JB1205 | 14 | 4 | 20 | 24 | 1 | 2 | 1 |
| JB1205M | 14 | 3 | 33 | 36 | 2 | 1 | 1 |
| JB1205M | 14 | 1 | 35 | 36 | 2 | 1 | 1 |
| JB1205M | 14 | 1 | 31 | 32 | 2 | 1 | 1 |
| JB1205M | 14 | 9 | 23 | 32 | 2 | 2 | 1 |
| JB1205M | 14 | 6 | 22 | 28 | 2 | 2 | 1 |
| JB1205M | 14 | 0 | 24 | 24 | 2 | 2 | 1 |

| **term** | **estimate** | **std.error** | **statistic** | **p.value** |
|---|---|---|---|---|
| **(Intercept)** | -2,820995212 | 0,176192738 | -16,0108484 | 1,07E-57 |
| **Group** | 0,030117088 | 0,008394247 | 3,587824872 | 0,000333448 |
| **Zygotic-Azygotic** | 0,158100622 | 0,081062135 | 1,950363399 | 0,051132821 |
| **4C Batch** | 0,782906515 | 0,087891002 | 8,90769814 | 5,21E-19 |
| **Other Batches** | -0,034710154 | 0,051812042 | -0,669924453 | 0,502905951 |

| **Experiment** | **Strain name** | **Group** | **Viable** | **No_viable** | **Total** | **Zygotic-azigotic** |
|---|---|---|---|---|---|---|
| One Way ANOVA | L972(JB22) | 1 | 12 | 24 | 36 | 1 |
| Zygotic vs azygotic | L972(JB22) | 1 | 4 | 32 | 36 | 1 |
| Batched samples | L972(JB22) | 1 | 2 | 34 | 36 | 1 |
| (Suplemental Figure 3E) | L972(JB22) | 1 | 5 | 27 | 32 | 1 |
| | L972(JB22) | 1 | 4 | 32 | 36 | 1 |
| | L972(JB22) | 1 | 5 | 27 | 32 | 1 |
| | L972(JB22) | 1 | 0 | 12 | 12 | 1 |
| | L972(JB22) | 1 | 0 | 8 | 8 | 1 |
| | L972(JB22) | 1 | 10 | 22 | 32 | 1 |

| | | | | | |
|---|---|---|---|---|---|
| L972(JB22) | 1 | 7 | 21 | 28 | 1 |
| L972(JB22) | 1 | 1 | 35 | 36 | 1 |
| L972(JB22) | 1 | 0 | 28 | 28 | 1 |
| L972(JB903) | 1 | 32 | 4 | 36 | 1 |
| L972(JB903) | 1 | 30 | 6 | 36 | 1 |
| L972(JB903) | 1 | 31 | 5 | 36 | 1 |
| L972(JB903) | 1 | 28 | 0 | 28 | 1 |
| L972(JB903) | 1 | 32 | 0 | 32 | 1 |
| L972(JB903) | 1 | 27 | 5 | 32 | 1 |
| L972(JB903) | 1 | 32 | 4 | 36 | 1 |
| L972(JB903) | 1 | 28 | 4 | 32 | 1 |
| L972(JB903) | 1 | 19 | 13 | 32 | 1 |
| L972(JB903) | 1 | 29 | 3 | 32 | 1 |
| L972(JB903) | 1 | 25 | 7 | 32 | 1 |
| L972(JB903) | 1 | 27 | 5 | 32 | 1 |
| L972M(JB22M) | 1 | 4 | 8 | 12 | 2 |
| L972M(JB22M) | 1 | 13 | 7 | 20 | 2 |
| L972M(JB22M) | 1 | 20 | 4 | 24 | 2 |
| L972M(JB22M) | 1 | 14 | 2 | 16 | 2 |
| L972M(JB22M) | 1 | 17 | 3 | 20 | 2 |
| L972M(JB22M) | 1 | 16 | 0 | 16 | 2 |
| JB32 | 2 | 0 | 36 | 36 | 1 |
| JB32 | 2 | 1 | 35 | 36 | 1 |
| JB32 | 2 | 2 | 34 | 36 | 1 |
| JB32 | 2 | 3 | 29 | 32 | 1 |
| JB32 | 2 | 0 | 36 | 36 | 1 |
| JB32 | 2 | 2 | 30 | 32 | 1 |
| JB32 | 2 | 2 | 26 | 28 | 1 |
| JB32 | 2 | 5 | 31 | 36 | 1 |
| JB32 | 2 | 3 | 33 | 36 | 1 |
| JB32 | 2 | 1 | 31 | 32 | 1 |
| JB32 | 2 | 2 | 26 | 28 | 1 |
| JB32 | 2 | 0 | 36 | 36 | 1 |
| J32M | 2 | 4 | 32 | 36 | 2 |
| J32M | 2 | 3 | 33 | 36 | 2 |
| J32M | 2 | 1 | 35 | 36 | 2 |
| J32M | 2 | 0 | 36 | 36 | 2 |
| J32M | 2 | 1 | 31 | 32 | 2 |
| J32M | 2 | 3 | 33 | 36 | 2 |
| JB840 | 3 | 5 | 31 | 36 | 1 |
| JB840 | 3 | 3 | 29 | 32 | 1 |
| JB840 | 3 | 1 | 35 | 36 | 1 |
| JB840 | 3 | 2 | 30 | 32 | 1 |
| JB840 | 3 | 3 | 29 | 32 | 1 |
| JB840 | 3 | 3 | 29 | 32 | 1 |
| JB840M | 3 | 3 | 33 | 36 | 2 |
| JB840M | 3 | 2 | 34 | 36 | 2 |
| JB840M | 3 | 5 | 31 | 36 | 2 |
| JB840M | 3 | 7 | 17 | 24 | 2 |
| JB840M | 3 | 10 | 22 | 32 | 2 |
| JB840M | 3 | 17 | 19 | 36 | 2 |
| JB845 | 4 | 5 | 23 | 28 | 1 |
| JB845 | 4 | 5 | 23 | 28 | 1 |
| JB845 | 4 | 4 | 28 | 32 | 1 |
| JB845 | 4 | 12 | 24 | 36 | 1 |
| JB845 | 4 | 9 | 27 | 36 | 1 |
| JB845 | 4 | 15 | 21 | 36 | 1 |
| JB845M | 4 | 3 | 33 | 36 | 2 |
| JB845M | 4 | 1 | 31 | 32 | 2 |
| JB845M | 4 | 0 | 36 | 36 | 2 |
| JB845M | 4 | 3 | 13 | 16 | 2 |
| JB845M | 4 | 12 | 16 | 28 | 2 |
| JB845M | 4 | 7 | 25 | 32 | 2 |
| JB848 | 5 | 0 | 32 | 32 | 1 |
| JB848 | 5 | 0 | 32 | 32 | 1 |
| JB848 | 5 | 4 | 28 | 32 | 1 |
| JB848 | 5 | 9 | 23 | 32 | 1 |
| JB848 | 5 | 1 | 31 | 32 | 1 |
| JB848 | 5 | 5 | 27 | 32 | 1 |
| JB848M | 5 | 0 | 36 | 36 | 2 |
| JB848M | 5 | 4 | 32 | 36 | 2 |
| JB848M | 5 | 2 | 34 | 36 | 2 |
| JB848M | 5 | 13 | 15 | 28 | 2 |
| JB848M | 5 | 12 | 20 | 32 | 2 |
| JB848M | 5 | 5 | 23 | 28 | 2 |
| JB870 | 6 | 3 | 29 | 32 | 1 |
| JB870 | 6 | 4 | 24 | 28 | 1 |
| JB870 | 6 | 0 | 36 | 36 | 1 |
| JB870 | 6 | 0 | 32 | 32 | 1 |
| JB870 | 6 | 1 | 35 | 36 | 1 |
| JB870 | 6 | 1 | 19 | 20 | 1 |
| JB870M | 6 | 2 | 34 | 36 | 2 |
| JB870M | 6 | 2 | 34 | 36 | 2 |

| | | | | | |
|---|---|---|---|---|---|
| JB870M | 6 | 1 | 27 | 28 | 2 |
| JB870M | 6 | 2 | 14 | 16 | 2 |
| JB870M | 6 | 6 | 26 | 32 | 2 |
| JB870M | 6 | 9 | 27 | 36 | 2 |
| JB873 | 7 | 5 | 31 | 36 | 1 |
| JB873 | 7 | 5 | 31 | 36 | 1 |
| JB873 | 7 | 10 | 22 | 32 | 1 |
| JB873 | 7 | 9 | 23 | 32 | 1 |
| JB873 | 7 | 8 | 16 | 24 | 1 |
| JB873 | 7 | 7 | 25 | 32 | 1 |
| JB873M | 7 | 3 | 29 | 32 | 2 |
| JB873M | 7 | 6 | 30 | 36 | 2 |
| JB873M | 7 | 7 | 25 | 32 | 2 |
| JB873M | 7 | 9 | 7 | 16 | 2 |
| JB873M | 7 | 6 | 14 | 20 | 2 |
| JB873M | 7 | 16 | 20 | 36 | 2 |
| JB913 | 8 | 3 | 33 | 36 | 1 |
| JB913 | 8 | 0 | 36 | 36 | 1 |
| JB913 | 8 | 2 | 26 | 28 | 1 |
| JB913 | 8 | 1 | 31 | 32 | 1 |
| JB913 | 8 | 0 | 28 | 28 | 1 |
| JB913 | 8 | 3 | 33 | 36 | 1 |
| JB913M | 8 | 6 | 30 | 36 | 2 |
| JB913M | 8 | 5 | 27 | 32 | 2 |
| JB913M | 8 | 11 | 21 | 32 | 2 |
| JB913M | 8 | 8 | 24 | 32 | 2 |
| JB913M | 8 | 5 | 23 | 28 | 2 |
| JB913M | 8 | 14 | 18 | 32 | 2 |
| JB916 | 9 | 3 | 25 | 28 | 1 |
| JB916 | 9 | 2 | 30 | 32 | 1 |
| JB916 | 9 | 1 | 35 | 36 | 1 |
| JB916 | 9 | 6 | 22 | 28 | 1 |
| JB916 | 9 | 0 | 24 | 24 | 1 |
| JB916 | 9 | 3 | 25 | 28 | 1 |
| JB916M | 9 | 4 | 32 | 36 | 2 |
| JB916M | 9 | 12 | 24 | 36 | 2 |
| JB916M | 9 | 8 | 24 | 32 | 2 |
| JB916M | 9 | 12 | 16 | 28 | 2 |
| JB916M | 9 | 7 | 25 | 32 | 2 |
| JB916M | 9 | 5 | 23 | 28 | 2 |
| JB931 | 10 | 12 | 4 | 16 | 1 |
| JB931 | 10 | 11 | 17 | 28 | 1 |
| JB931 | 10 | 16 | 12 | 28 | 1 |
| JB931 | 10 | 3 | 33 | 36 | 1 |
| JB931 | 10 | 10 | 26 | 36 | 1 |
| JB931 | 10 | 2 | 18 | 20 | 1 |
| JB931M | 10 | 3 | 25 | 28 | 2 |
| JB931M | 10 | 1 | 23 | 24 | 2 |
| JB931M | 10 | 5 | 31 | 36 | 2 |
| JB938 | 11 | 7 | 29 | 36 | 1 |
| JB938 | 11 | 7 | 29 | 36 | 1 |
| JB938 | 11 | 3 | 17 | 20 | 1 |
| JB938 | 11 | 4 | 28 | 32 | 1 |
| JB938 | 11 | 3 | 17 | 20 | 1 |
| JB938 | 11 | 1 | 31 | 32 | 1 |
| JB938M | 11 | 2 | 34 | 36 | 2 |
| JB938M | 11 | 14 | 10 | 24 | 2 |
| JB938M | 11 | 2 | 30 | 32 | 2 |
| JB938M | 11 | 4 | 24 | 28 | 2 |
| JB953 | 12 | 3 | 13 | 16 | 1 |
| JB953 | 12 | 9 | 19 | 28 | 1 |
| JB953 | 12 | 17 | 19 | 36 | 1 |
| JB953 | 12 | 14 | 14 | 28 | 1 |
| JB953 | 12 | 10 | 18 | 28 | 1 |
| JB953 | 12 | 9 | 19 | 28 | 1 |
| JB953M | 12 | 15 | 21 | 36 | 2 |
| JB953M | 12 | 14 | 14 | 28 | 2 |
| JB953M | 12 | 5 | 27 | 32 | 2 |
| JB953M | 12 | 24 | 12 | 36 | 2 |
| JB953M | 12 | 19 | 9 | 28 | 2 |
| JB953M | 12 | 15 | 17 | 32 | 2 |
| JB1114 | 13 | 16 | 20 | 36 | 1 |
| JB1114 | 13 | 8 | 24 | 32 | 1 |
| JB1114 | 13 | 16 | 20 | 36 | 1 |
| JB1114 | 13 | 7 | 25 | 32 | 1 |
| JB1114 | 13 | 5 | 31 | 36 | 1 |
| JB1114 | 13 | 10 | 22 | 32 | 1 |
| JB1114M | 13 | 7 | 21 | 28 | 2 |
| JB1114M | 13 | 10 | 26 | 36 | 2 |
| JB1114M | 13 | 6 | 26 | 32 | 2 |
| JB1114M | 13 | 4 | 8 | 12 | 2 |
| JB1114M | 13 | 9 | 11 | 20 | 2 |
| JB1114M | 13 | 11 | 21 | 32 | 2 |

| | | | | | | |
|---|---|---|---|---|---|---|
| | JB1205 | 14 | 2 | 34 | 36 | 1 |
| | JB1205 | 14 | 1 | 19 | 20 | 1 |
| | JB1205 | 14 | 11 | 17 | 28 | 1 |
| | JB1205 | 14 | 4 | 32 | 36 | 1 |
| | JB1205 | 14 | 13 | 19 | 32 | 1 |
| | JB1205 | 14 | 4 | 20 | 24 | 1 |
| | JB1205M | 14 | 3 | 33 | 36 | 2 |
| | JB1205M | 14 | 1 | 35 | 36 | 2 |
| | JB1205M | 14 | 1 | 31 | 32 | 2 |
| | JB1205M | 14 | 9 | 23 | 32 | 2 |
| | JB1205M | 14 | 6 | 22 | 28 | 2 |
| | JB1205M | 14 | 0 | 24 | 24 | 2 |

| term | estimate | std.error | statistic | p.value |
|---|---|---|---|---|
| **(Intercept)** | -1,057676394 | 0,081522449 | -12,97405074 | 1,72E-38 |
| **Zygotic-Azygotic** | -0,160872544 | 0,057175062 | -2,81368379 | 0,004897738 |

| Experiment | Strain/Cross name | Group | Viable | No_viable | Total | Repeat |
|---|---|---|---|---|---|---|
| One Way ANOVA | L968xJB759 | 1 | 239 | 185 | 424 | 1 |
| Biological repeats | L968xJB759 | 1 | 87 | 65 | 152 | 2 |
| Azygotic data | L968xJB759 | 1 | 85 | 87 | 172 | 3 |
| (Supplemental Figure 4A) | L972xJB759 | 2 | 102 | 118 | 220 | 1 |
| | L972xJB759 | 2 | 69 | 79 | 148 | 2 |
| | L968xJB760 | 3 | 337 | 263 | 600 | 1 |
| | L968xJB760 | 3 | 109 | 55 | 164 | 2 |
| | JB32xJB760 | 4 | 65 | 127 | 192 | 1 |
| | JB32xJB760 | 4 | 54 | 114 | 168 | 2 |
| | L972xJB32 | 5 | 183 | 45 | 228 | 1 |
| | L972xJB32 | 5 | 166 | 26 | 192 | 2 |
| | L968 | 6 | 210 | 38 | 248 | 1 |
| | L968 | 6 | 171 | 37 | 208 | 2 |
| | L972(JB22) | 7 | 176 | 32 | 208 | 1 |
| | L972(JB22) | 7 | 126 | 18 | 144 | 2 |
| | L972(JB903) | 8 | 180 | 20 | 200 | 1 |
| | L972(JB903) | 8 | 160 | 36 | 196 | 2 |
| | L975 | 9 | 168 | 8 | 176 | 1 |
| | L975 | 9 | 202 | 6 | 208 | 2 |
| | JB32 | 10 | 200 | 8 | 208 | 1 |
| | JB32 | 10 | 183 | 13 | 196 | 2 |
| | JB32xJB761 | 11 | 204 | 88 | 292 | 1 |
| | JB32xJB761 | 11 | 98 | 58 | 156 | 2 |
| | JB758 | 12 | 34 | 14 | 48 | 1 |
| | JB758 | 12 | 20 | 8 | 28 | 2 |
| | JB763 | 13 | 4 | 68 | 72 | 1 |
| | JB763 | 13 | 5 | 15 | 20 | 2 |
| | JB759 | 14 | 228 | 172 | 400 | 1 |
| | JB759 | 14 | 21 | 15 | 36 | 2 |
| | JB759xJB760 | 15 | 305 | 165 | 470 | 1 |
| | JB759xJB760 | 15 | 255 | 157 | 412 | 2 |

| term | estimate | std.error | statistic | p.value |
|---|---|---|---|---|
| **(Intercept)** | -0,648876484 | 0,073610348 | -8,815017199 | 1,20E-18 |
| **Repeat** | -0,058061018 | 0,048590858 | -1,19489593 | 0,232127693 |

| Experiment | Strain/Cross name | Group (Timepoint) | Viable | No_viable | Total | F1 |
|---|---|---|---|---|---|---|
| Multifactorial ANOVA | MMN_JB32_R1_T0 | 1 | 35 | 1 | 36 | 0 |
| JB32 MM-N | MMN_JB32_R1_T0 | 1 | 33 | 3 | 36 | 0 |
| Epigenetic degeneration | MMN_JB32_R1_T0 | 1 | 33 | 3 | 36 | 0 |
| F1 effect | MMN_JB32_R1_T0 | 1 | 30 | 6 | 36 | 0 |
| (Figure 7B) | MMN_JB32_R1_T0 | 1 | 34 | 2 | 36 | 0 |
| | MMN_JB32_R1_T0 | 1 | 27 | 1 | 28 | 0 |
| | MMN_JB32_R1_T1 | 2 | 25 | 7 | 32 | 0 |
| | MMN_JB32_R1_T1 | 2 | 33 | 3 | 36 | 0 |
| | MMN_JB32_R1_T1 | 2 | 33 | 3 | 36 | 0 |
| | MMN_JB32_R1_T1 | 2 | 34 | 2 | 36 | 0 |
| | MMN_JB32_R1_T1 | 2 | 31 | 1 | 32 | 0 |
| | MMN_JB32_R1_T1 | 2 | 36 | 0 | 36 | 0 |
| | MMN_JB32_R1_T1_F1 | 2 | 12 | 8 | 20 | 1 |
| | MMN_JB32_R1_T1_F1 | 2 | 20 | 0 | 20 | 1 |
| | MMN_JB32_R1_T1_F1 | 2 | 15 | 1 | 16 | 1 |
| | MMN_JB32_R1_T1_F1 | 2 | 14 | 6 | 20 | 1 |
| | MMN_JB32_R1_T1_F1 | 2 | 23 | 1 | 24 | 1 |
| | MMN_JB32_R1_T1_F1 | 2 | 9 | 3 | 12 | 1 |
| | MMN_JB32_R1_T2 | 3 | 34 | 2 | 36 | 0 |
| | MMN_JB32_R1_T2 | 3 | 33 | 3 | 36 | 0 |
| | MMN_JB32_R1_T2 | 3 | 34 | 2 | 36 | 0 |
| | MMN_JB32_R1_T2 | 3 | 33 | 3 | 36 | 0 |
| | MMN_JB32_R1_T2 | 3 | 32 | 0 | 32 | 0 |
| | MMN_JB32_R1_T2 | 3 | 35 | 1 | 36 | 0 |
| | MMN_JB32_R1_T2_F1 | 3 | 32 | 0 | 32 | 1 |
| | MMN_JB32_R1_T2_F1 | 3 | 33 | 3 | 36 | 1 |
| | MMN_JB32_R1_T2_F1 | 3 | 36 | 0 | 36 | 1 |
| | MMN_JB32_R1_T2_F1 | 3 | 36 | 0 | 36 | 1 |

| | | | | | |
|---|---|---|---|---|---|
| MMN_JB32_R1_T2_F1 | 3 | 34 | 2 | 36 | 1 |
| MMN_JB32_R1_T2_F1 | 3 | 33 | 3 | 36 | 1 |
| MMN_JB32_R1_T3 | 4 | 31 | 5 | 36 | 0 |
| MMN_JB32_R1_T3 | 4 | 31 | 5 | 36 | 0 |
| MMN_JB32_R1_T3 | 4 | 28 | 4 | 32 | 0 |
| MMN_JB32_R1_T3 | 4 | 33 | 3 | 36 | 0 |
| MMN_JB32_R1_T3 | 4 | 32 | 4 | 36 | 0 |
| MMN_JB32_R1_T3 | 4 | 27 | 5 | 32 | 0 |
| MMN_JB32_R1_T3_F1 | 4 | 33 | 3 | 36 | 1 |
| MMN_JB32_R1_T3_F1 | 4 | 27 | 5 | 32 | 1 |
| MMN_JB32_R1_T3_F1 | 4 | 32 | 0 | 32 | 1 |
| MMN_JB32_R1_T3_F1 | 4 | 24 | 4 | 28 | 1 |
| MMN_JB32_R1_T3_F1 | 4 | 31 | 1 | 32 | 1 |
| MMN_JB32_R1_T3_F1 | 4 | 34 | 2 | 36 | 1 |
| MMN_JB32_R1_T4 | 5 | 29 | 7 | 36 | 0 |
| MMN_JB32_R1_T4 | 5 | 28 | 4 | 32 | 0 |
| MMN_JB32_R1_T4 | 5 | 25 | 7 | 32 | 0 |
| MMN_JB32_R1_T4 | 5 | 33 | 3 | 36 | 0 |
| MMN_JB32_R1_T4 | 5 | 33 | 3 | 36 | 0 |
| MMN_JB32_R1_T4 | 5 | 25 | 11 | 36 | 0 |
| MMN_JB32_R1_T4_F1 | 5 | 34 | 2 | 36 | 1 |
| MMN_JB32_R1_T4_F1 | 5 | 35 | 1 | 36 | 1 |
| MMN_JB32_R1_T4_F1 | 5 | 28 | 4 | 32 | 1 |
| MMN_JB32_R1_T4_F1 | 5 | 28 | 4 | 32 | 1 |
| MMN_JB32_R1_T4_F1 | 5 | 29 | 3 | 32 | 1 |
| MMN_JB32_R1_T4_F1 | 5 | 29 | 7 | 36 | 1 |
| MMN_JB32_R1_T5 | 6 | 32 | 4 | 36 | 0 |
| MMN_JB32_R1_T5 | 6 | 24 | 4 | 28 | 0 |
| MMN_JB32_R1_T5 | 6 | 33 | 3 | 36 | 0 |
| MMN_JB32_R1_T5 | 6 | 30 | 6 | 36 | 0 |
| MMN_JB32_R1_T5 | 6 | 28 | 8 | 36 | 0 |
| MMN_JB32_R1_T5 | 6 | 29 | 7 | 36 | 0 |
| MMN_JB32_R1_T5_F1 | 6 | 24 | 4 | 28 | 1 |
| MMN_JB32_R1_T5_F1 | 6 | 25 | 7 | 32 | 1 |
| MMN_JB32_R1_T5_F1 | 6 | 34 | 2 | 36 | 1 |
| MMN_JB32_R1_T5_F1 | 6 | 12 | 4 | 16 | 1 |
| MMN_JB32_R1_T5_F1 | 6 | 33 | 3 | 36 | 1 |
| MMN_JB32_R1_T5_F1 | 6 | 32 | 4 | 36 | 1 |

| term | estimate | std.error | statistic | p.value |
|---|---|---|---|---|
| **(Intercept)** | 2,767512753 | 0,202344253 | 13,67724909 | 1,39E-42 |
| **Timepoint** | -0,179061865 | 0,046308942 | -3,866680085 | 0,000110327 |
| **Generation (Parental or F1)** | 0,242539641 | 0,147059875 | 1,649257767 | 0,099094838 |

| Experiment | Strain/Cross name | Group (Timepoint) | Viable | No_viable | Total | F1 |
|---|---|---|---|---|---|---|
| Multifactorial ANOVA | MMN_JB760_R2_T0 | 1 | 1 | 23 | 24 | 0 |
| JB760 MM-N | MMN_JB760_R2_T0 | 1 | 5 | 19 | 24 | 0 |
| Epigenetic degeneration | MMN_JB760_R2_T0 | 1 | 2 | 30 | 32 | 0 |
| F1 effect | MMN_JB760_R2_T0 | 1 | 1 | 31 | 32 | 0 |
| (Figure 7B) | MMN_JB760_R2_T0 | 1 | 0 | 32 | 32 | 0 |
| | MMN_JB760_R2_T0 | 1 | 3 | 25 | 28 | 0 |
| | MMN_JB760_R2_T1 | 2 | 20 | 12 | 32 | 0 |
| | MMN_JB760_R2_T1 | 2 | 28 | 8 | 36 | 0 |
| | MMN_JB760_R2_T1 | 2 | 20 | 16 | 36 | 0 |
| | MMN_JB760_R2_T1 | 2 | 22 | 6 | 28 | 0 |
| | MMN_JB760_R2_T1 | 2 | 26 | 10 | 36 | 0 |
| | MMN_JB760_R2_T1 | 2 | 23 | 13 | 36 | 0 |
| | MMN_JB760_R2_T1_F1 | 2 | 19 | 13 | 32 | 1 |
| | MMN_JB760_R2_T1_F1 | 2 | 13 | 19 | 32 | 1 |
| | MMN_JB760_R2_T1_F1 | 2 | 15 | 9 | 24 | 1 |
| | MMN_JB760_R2_T1_F1 | 2 | 26 | 10 | 36 | 1 |
| | MMN_JB760_R2_T1_F1 | 2 | 13 | 15 | 28 | 1 |
| | MMN_JB760_R2_T1_F1 | 2 | 4 | 32 | 36 | 1 |
| | MMN_JB760_R2_T2 | 3 | 18 | 18 | 36 | 0 |
| | MMN_JB760_R2_T2 | 3 | 30 | 6 | 36 | 0 |
| | MMN_JB760_R2_T2 | 3 | 18 | 18 | 36 | 0 |
| | MMN_JB760_R2_T2 | 3 | 12 | 24 | 36 | 0 |
| | MMN_JB760_R2_T2 | 3 | 15 | 21 | 36 | 0 |
| | MMN_JB760_R2_T2 | 3 | 18 | 18 | 36 | 0 |
| | MMN_JB760_R2_T2_F1 | 3 | 12 | 24 | 36 | 1 |
| | MMN_JB760_R2_T2_F1 | 3 | 7 | 25 | 32 | 1 |
| | MMN_JB760_R2_T2_F1 | 3 | 11 | 25 | 36 | 1 |
| | MMN_JB760_R2_T2_F1 | 3 | 17 | 19 | 36 | 1 |
| | MMN_JB760_R2_T2_F1 | 3 | 11 | 25 | 36 | 1 |
| | MMN_JB760_R2_T2_F1 | 3 | 15 | 17 | 32 | 1 |
| | MMN_JB760_R2_T3 | 4 | 19 | 17 | 36 | 0 |
| | MMN_JB760_R2_T3 | 4 | 17 | 15 | 32 | 0 |
| | MMN_JB760_R2_T3 | 4 | 20 | 16 | 36 | 0 |
| | MMN_JB760_R2_T3 | 4 | 22 | 14 | 36 | 0 |
| | MMN_JB760_R2_T3 | 4 | 25 | 11 | 36 | 0 |
| | MMN_JB760_R2_T3 | 4 | 17 | 19 | 36 | 0 |
| | MMN_JB760_R2_T3_F1 | 4 | 15 | 17 | 32 | 1 |
| | MMN_JB760_R2_T3_F1 | 4 | 8 | 20 | 28 | 1 |

| | | | | | |
|---|---|---|---|---|---|
| MMN_JB760_R2_T3_F1 | 4 | 8 | 20 | 28 | 1 |
| MMN_JB760_R2_T3_F1 | 4 | 11 | 25 | 36 | 1 |
| MMN_JB760_R2_T3_F1 | 4 | 8 | 24 | 32 | 1 |
| MMN_JB760_R2_T3_F1 | 4 | 9 | 27 | 36 | 1 |

| term | estimate | std.error | statistic | p.value |
|---|---|---|---|---|
| **(Intercept)** | -0,767669968 | 0,161925008 | -4,740898129 | 2,13E-06 |
| **Timepoint** | 0,257904019 | 0,055925483 | 4,611565353 | 3,99648E-06 |
| **Generation (Parental or F1)** | -0,513019678 | 0,11412405 | -4,495281041 | 6,94781E-06 |

### Supplemental Table S5: Clonal groups

| Reference strain (Jeffares et al 2015) | Number of members | Clonal Group members (see Jeffares et al 2015 for strain details) | Cluster group (Jeffares et al 2015) |
|---|---|---|---|
| JB22(L972) | 20 | JB1168;JB1169;JB1170;JB1179;JB1204;JB22(L972);JB374;JB50;JB761;JB868;JB906;JB936;JB937;JB940;JB941;JB945;JB892;JB32;JB903(L972);JB904(L975) | 4 |
| JB864 (NCYC132) | 14 | JB1108;JB1112;JB1113;JB1176;JB1177;JB1198;JB1201;JB588;JB861;JB864;JB876;JB890;JB894;JB590 | 1 |
| JB900 | 10 | JB1114;JB1175;JB1189;JB1190;JB1191;JB1192;JB1193;JB1194;JB900;JB901 | 3 |
| JB869 | 8 | JB1181;JB1182;JB1183;JB1184;JB1195;JB759;JB869;JB897 | 4 |
| JB760 | 7 | JB760;JB885;JB886;JB887;JB888;JB889;JB891 | 4 |
| JB914 | 6 | JB1116;JB1153;JB1178;JB1199;JB763;JB914 | 3 |
| JB840 | 6 | JB840;JB843;JB847;JB850;JB855;JB856 | 2 |
| JB4 | 5 | JB4;JB593;JB859;JB893;JB905 | 5 |
| JB878 | 5 | JB878;JB880;JB881;JB882;JB883 | 4 |
| JB929 | 5 | JB907;JB908;JB929;JB944;JB947 | 3 |
| JB910 | 5 | JB1202;JB909;JB910;JB946;JB948 | 3 |
| JB872 | 5 | JB1203;JB592;JB872;JB912;JB915 | 3 |
| JB899 | 5 | JB1185;JB1186;JB1187;JB1188;JB899 | 5 |
| JB939 | 4 | JB1166;JB1172;JB877;JB939 | 2 |
| JB762 | 4 | JB1109;JB1115;JB762;JB911 | 3 |
| JB934 | 3 | JB932;JB933;JB934 | 5 |
| JB870 | 3 | JB1111;JB870;JB898 | 4 |
| JB842 | 3 | JB842;JB851;JB857 | 2 |
| JB853 | 3 | JB844;JB849;JB853 | 2 |
| JB1171 | 2 | JB1167;JB1171 | 5 |
| JB902 | 2 | JB1196;JB902 | 3 |
| JB841 | 2 | JB839;JB841 | 2 |
| JB758 | 2 | JB758;JB866 | 1 |
| JB874 | 2 | JB874;JB594 | 2 |
| JB953 | 2 | JB952;JB953 | 1 |

### Supplemental Table S6: Polymorphic sites

**GREEN= SHARED POLYMORPHIC SITES**

| CHROM | POS | REF | ALT | L972 CLONAL GROUP | NCYC132 CLONAL GROUP | JB1114-JB1175 CLONAL STRAINS |
|---|---|---|---|---|---|---|
| I | 65141 | A | G | 0 | 0 | 1 |
| I | 65144 | A | C,G | 0 | 0 | 1 |
| I | 65147 | C | A,T | 0 | 0 | 1 |
| I | 65150 | T | A | 0 | 0 | 1 |
| I | 65168 | T | C | 0 | 0 | 1 |
| I | 65922 | T | G | 0 | 1 | 0 |
| I | 66223 | T | G | 0 | 0 | 1 |
| I | 67769 | A | G | 0 | 1 | 0 |
| I | 67781 | A | T,G | 0 | 1 | 0 |
| I | 67784 | C | T | 0 | 1 | 0 |
| I | 68555 | A | G | 0 | 0 | 1 |
| I | 68576 | T | G | 0 | 0 | 1 |
| I | 68615 | T | A | 0 | 0 | 1 |
| I | 121590 | G | A | 1 | 0 | 0 |
| I | 207457 | A | C | 0 | 1 | 0 |
| I | 252458 | G | A | 1 | 0 | 0 |
| I | 365062 | G | A | 0 | 1 | 0 |
| I | 365072 | T | A | 1 | 0 | 0 |
| I | 405322 | G | A | 1 | 0 | 0 |
| I | 410610 | T | C | 0 | 1 | 0 |
| I | 461925 | T | G | 1 | 0 | 0 |
| I | 477739 | T | C | 1 | 0 | 0 |
| I | 488673 | G | A | 0 | 1 | 0 |
| I | 523603 | G | A | 1 | 0 | 0 |
| I | 523883 | G | A | 1 | 0 | 0 |
| I | 527156 | C | A | 1 | 0 | 0 |
| I | 633906 | T | C | 0 | 1 | 1 |
| I | 567033 | G | A | 0 | 1 | 0 |
| I | 641341 | C | A | 0 | 1 | 0 |
| I | 759334 | G | T | 1 | 0 | 0 |
| I | 862964 | G | T | 1 | 0 | 0 |
| I | 878336 | C | T | 0 | 1 | 0 |
| I | 879269 | A | G | 0 | 0 | 1 |
| I | 921071 | A | T | 1 | 0 | 0 |
| I | 927629 | C | G | 0 | 1 | 0 |
| I | 1129447 | G | A | 0 | 0 | 1 |
| I | 1149519 | A | C | 0 | 0 | 1 |
| I | 1155495 | C | T | 0 | 1 | 0 |
| I | 1234445 | T | A | 0 | 0 | 1 |
| I | 1240632 | C | T | 0 | 1 | 0 |
| I | 1369596 | A | T | 0 | 1 | 0 |
| I | 1434718 | T | C | 0 | 1 | 0 |
| I | 1474835 | A | G | 1 | 0 | 0 |
| I | 1528455 | T | A | 0 | 1 | 0 |
| I | 1562817 | G | A | 0 | 1 | 0 |
| I | 1562818 | T | A | 0 | 1 | 0 |
| I | 1572455 | T | A | 0 | 1 | 0 |
| I | 1664907 | T | A | 1 | 0 | 0 |
| I | 1779005 | C | A | 0 | 1 | 0 |
| I | 1783458 | C | T | 1 | 0 | 0 |
| I | 1816034 | C | T | 1 | 0 | 0 |
| I | 1906124 | C | T | 1 | 0 | 0 |
| I | 1921486 | G | A | 0 | 1 | 0 |
| I | 1966097 | G | T | 0 | 1 | 0 |
| I | 1999192 | G | C | 1 | 0 | 0 |
| I | 2025398 | C | T | 1 | 0 | 0 |
| I | 2323403 | C | T | 1 | 0 | 0 |
| I | 2603703 | G | A | 1 | 0 | 0 |
| I | 2605764 | T | A | 1 | 0 | 0 |

| | | | | | | |
|---|---|---|---|---|---|---|
| I | 2642318 | G | A | 1 | 0 | 0 |
| I | 2643596 | C | A | 1 | 0 | 0 |
| I | 2655865 | G | A | 1 | 0 | 0 |
| I | 2874735 | C | T | 0 | 1 | 0 |
| I | 2874777 | G | C | 0 | 1 | 0 |
| I | 2992238 | T | C | 0 | 1 | 0 |
| I | 3020428 | G | A | 0 | 1 | 0 |
| I | 3043452 | T | C | 0 | 1 | 0 |
| I | 3055184 | C | T | 1 | 0 | 0 |
| I | 3126991 | T | A | 0 | 1 | 1 |
| I | 3060076 | T | C | 0 | 1 | 0 |
| I | 3111508 | A | G | 1 | 0 | 0 |
| I | 3115841 | C | A | 1 | 0 | 0 |
| I | 3119299 | G | A | 1 | 0 | 0 |
| I | 3179238 | G | C | 1 | 0 | 0 |
| I | 3204845 | G | A | 1 | 0 | 0 |
| I | 3350687 | C | T | 0 | 1 | 0 |
| I | 3416193 | C | T | 1 | 0 | 0 |
| I | 3451522 | T | A | 0 | 1 | 0 |
| I | 3479252 | C | T | 1 | 0 | 0 |
| I | 3545238 | G | T | 0 | 1 | 0 |
| I | 3559922 | G | T | 0 | 1 | 0 |
| I | 3571024 | G | A | 0 | 1 | 0 |
| I | 3583106 | G | A | 1 | 0 | 0 |
| I | 3638413 | A | T | 0 | 0 | 1 |
| I | 3742761 | G | A | 1 | 0 | 0 |
| I | 3805756 | T | C | 1 | 0 | 0 |
| I | 3920563 | C | A,T | 0 | 1 | 0 |
| I | 3978510 | T | C | 0 | 1 | 0 |
| I | 3987405 | G | T,A | 0 | 1 | 0 |
| I | 4139300 | A | T | 1 | 0 | 0 |
| I | 4141559 | A | G | 1 | 0 | 0 |
| I | 4141560 | A | G | 1 | 0 | 0 |
| I | 4160606 | C | T | 0 | 1 | 0 |
| I | 4237712 | T | A | 1 | 0 | 0 |
| I | 4276701 | T | A | 0 | 1 | 0 |
| I | 4280513 | C | T | 0 | 1 | 0 |
| I | 4281979 | G | A | 0 | 1 | 0 |
| I | 4299616 | T | G | 0 | 0 | 1 |
| I | 4299617 | T | G | 0 | 0 | 1 |
| I | 4375421 | A | T | 1 | 0 | 0 |
| I | 4427125 | G | T | 0 | 1 | 0 |
| I | 4442714 | G | A | 1 | 0 | 0 |
| I | 4462986 | T | A,C | 0 | 1 | 0 |
| I | 4565906 | G | A | 0 | 1 | 0 |
| I | 4573959 | C | A | 0 | 1 | 0 |
| I | 4599222 | C | A | 1 | 0 | 0 |
| I | 4614261 | C | A | 0 | 1 | 0 |
| I | 4673274 | A | G | 0 | 0 | 1 |
| I | 4764033 | C | A | 1 | 0 | 0 |
| I | 4809907 | C | A | 0 | 1 | 0 |
| I | 4919603 | A | G | 1 | 0 | 0 |
| I | 4936568 | G | T | 0 | 1 | 0 |
| I | 4993828 | G | T | 0 | 1 | 0 |
| I | 5310098 | A | T | 0 | 1 | 0 |
| I | 5310099 | C | T | 0 | 1 | 0 |
| I | 5314711 | G | A | 0 | 1 | 0 |
| I | 5400080 | C | T | 1 | 0 | 0 |
| I | 5456432 | G | A | 1 | 0 | 0 |
| I | 5466209 | T | G | 0 | 1 | 0 |
| I | 5467854 | G | T | 0 | 1 | 0 |
| I | 5482903 | A | T | 0 | 1 | 0 |
| I | 5506394 | C | T | 0 | 0 | 1 |
| I | 5506403 | A | T | 0 | 0 | 1 |
| I | 5506406 | C | T,A | 0 | 0 | 1 |

| | | | | | | |
|---|---|---|---|---|---|---|
| I | 5506409 | A | C | 0 | 0 | 1 |
| I | 5506415 | C | T | 0 | 0 | 1 |
| I | 5506418 | T | A,G | 0 | 0 | 1 |
| I | 5506433 | C | T | 0 | 0 | 1 |
| I | 5506448 | A | G | 0 | 0 | 1 |
| I | 5511129 | G | A | 0 | 1 | 0 |
| I | 5536460 | A | G | 1 | 0 | 0 |
| I | 5546377 | G | A | 0 | 0 | 1 |
| I | 5547005 | G | A | 0 | 0 | 1 |
| I | 5547288 | A | G | 0 | 0 | 1 |
| I | 5547298 | C | G | 0 | 0 | 1 |
| I | 5547849 | C | T | 0 | 0 | 1 |
| I | 5548012 | T | C | 0 | 0 | 1 |
| I | 5548675 | A | G | 0 | 0 | 1 |
| I | 5549040 | T | A,G | 0 | 0 | 1 |
| I | 5549595 | T | C | 0 | 0 | 1 |
| I | 5549827 | A | G | 0 | 0 | 1 |
| I | 5549847 | C | T | 0 | 0 | 1 |
| I | 5549877 | C | A | 0 | 0 | 1 |
| I | 5550571 | T | C | 0 | 0 | 1 |
| I | 5550582 | T | A | 0 | 0 | 1 |
| I | 5551505 | A | G | 0 | 0 | 1 |
| I | 5551524 | A | G | 0 | 0 | 1 |
| II | 72910 | T | A | 0 | 1 | 0 |
| II | 82108 | A | C | 1 | 0 | 0 |
| II | 153847 | C | T | 1 | 0 | 0 |
| II | 154533 | A | G | 1 | 0 | 0 |
| II | 154545 | G | A | 1 | 1 | 0 |
| II | 154550 | C | T | 1 | 1 | 0 |
| II | 154551 | A | G | 1 | 1 | 0 |
| II | 154560 | C | T | 1 | 1 | 0 |
| II | 154603 | A | C | 1 | 0 | 0 |
| II | 155034 | G | A | 0 | 0 | 1 |
| II | 155115 | A | G | 0 | 1 | 0 |
| II | 174367 | C | T | 0 | 1 | 0 |
| II | 213679 | G | A | 1 | 0 | 0 |
| II | 250906 | G | A | 1 | 0 | 0 |
| II | 266718 | A | C | 1 | 0 | 0 |
| II | 266780 | T | A | 1 | 0 | 0 |
| II | 276539 | A | T | 1 | 0 | 0 |
| II | 320145 | C | T | 1 | 0 | 0 |
| II | 333869 | A | T | 0 | 1 | 0 |
| II | 339643 | A | G | 0 | 1 | 0 |
| II | 474356 | T | A | 1 | 0 | 0 |
| II | 495754 | G | A | 1 | 0 | 0 |
| II | 526743 | C | T | 0 | 0 | 1 |
| II | 538980 | A | T,G | 0 | 0 | 1 |
| II | 685728 | G | A | 1 | 0 | 0 |
| II | 699951 | T | G | 1 | 0 | 0 |
| II | 699961 | A | T | 1 | 0 | 0 |
| II | 700028 | A | T | 1 | 0 | 0 |
| II | 700113 | A | T | 1 | 0 | 0 |
| II | 752407 | C | A | 0 | 1 | 0 |
| II | 778181 | A | G | 0 | 1 | 0 |
| II | 819796 | C | T | 0 | 1 | 0 |
| II | 828279 | G | A | 0 | 1 | 0 |
| II | 862314 | A | T | 1 | 0 | 0 |
| II | 863347 | C | T | 1 | 0 | 0 |
| II | 869806 | G | C | 1 | 0 | 0 |
| II | 879033 | T | A | 1 | 0 | 0 |
| II | 905938 | C | T | 1 | 0 | 0 |
| II | 923543 | C | A | 1 | 0 | 0 |
| II | 943554 | C | A | 1 | 0 | 0 |
| II | 953201 | A | G | 1 | 0 | 0 |
| II | 967094 | A | G | 1 | 0 | 0 |

| | | | | | | |
|---|---|---|---|---|---|---|
| II | 1040271 | G | A | 0 | 1 | 0 |
| II | 1049434 | G | A | 0 | 1 | 0 |
| II | 1120634 | A | G | 0 | 0 | 1 |
| II | 1155903 | T | A | 1 | 0 | 0 |
| II | 1159678 | G | C | 1 | 0 | 0 |
| II | 1159694 | A | T | 1 | 0 | 0 |
| II | 1209990 | G | A | 0 | 1 | 0 |
| II | 1233738 | G | T | 0 | 1 | 0 |
| II | 1297891 | C | T | 0 | 1 | 0 |
| II | 1300085 | A | T,G | 0 | 0 | 1 |
| II | 1302841 | G | A | 1 | 0 | 0 |
| II | 1416994 | G | T | 0 | 1 | 0 |
| II | 1422317 | T | A | 1 | 0 | 0 |
| II | 1505255 | T | A | 0 | 1 | 0 |
| II | 1553976 | A | T | 1 | 0 | 0 |
| II | 1672785 | G | C | 1 | 0 | 0 |
| II | 1672859 | C | T | 1 | 0 | 0 |
| II | 1673002 | G | C | 1 | 0 | 0 |
| II | 1678693 | T | A | 1 | 0 | 0 |
| II | 1683324 | T | G | 1 | 0 | 0 |
| II | 1697124 | C | T | 1 | 0 | 0 |
| II | 1784826 | G | A | 1 | 0 | 0 |
| II | 1787400 | C | T | 0 | 1 | 0 |
| II | 1849777 | G | A | 1 | 0 | 0 |
| II | 1890505 | C | A | 1 | 0 | 0 |
| II | 1962419 | T | A | 1 | 0 | 0 |
| II | 1962439 | A | G | 1 | 0 | 0 |
| II | 1962458 | T | A | 1 | 0 | 0 |
| II | 1976644 | G | A | 1 | 0 | 0 |
| II | 1987098 | C | A | 1 | 0 | 0 |
| II | 2007715 | A | G | 1 | 0 | 0 |
| II | 2024227 | A | T | 1 | 0 | 0 |
| II | 2044929 | A | T | 1 | 0 | 0 |
| II | 2160074 | C | T | 1 | 0 | 0 |
| II | 2200004 | A | G | 0 | 0 | 1 |
| II | 2228775 | T | G | 1 | 0 | 0 |
| II | 2228777 | G | T | 1 | 0 | 0 |
| II | 2327066 | T | G | 0 | 1 | 0 |
| II | 2394638 | C | T | 1 | 0 | 0 |
| II | 2402384 | C | T | 1 | 0 | 0 |
| II | 2410250 | A | C | 0 | 1 | 0 |
| II | 2436580 | C | G | 1 | 0 | 0 |
| II | 2444543 | T | C | 1 | 0 | 0 |
| II | 2578154 | C | T | 1 | 0 | 0 |
| II | 2676336 | C | T | 1 | 0 | 0 |
| II | 2693982 | T | A | 0 | 1 | 0 |
| II | 2693983 | G | A | 0 | 1 | 0 |
| II | 2756603 | C | T | 1 | 0 | 0 |
| II | 2806053 | G | A | 1 | 0 | 0 |
| II | 2806808 | C | T | 1 | 0 | 0 |
| II | 2817130 | A | C | 0 | 1 | 0 |
| II | 2824897 | T | C | 0 | 1 | 0 |
| II | 2871636 | C | T | 0 | 1 | 0 |
| II | 2888520 | G | T | 0 | 1 | 0 |
| II | 2901509 | G | T | 1 | 0 | 0 |
| II | 2906868 | G | T | 0 | 1 | 0 |
| II | 2942156 | G | T | 0 | 1 | 0 |
| II | 2969812 | T | A | 0 | 1 | 0 |
| II | 2984089 | G | A | 1 | 0 | 0 |
| II | 2988458 | C | T | 0 | 1 | 0 |
| II | 3006137 | A | T | 1 | 0 | 0 |
| II | 3056228 | C | A | 0 | 1 | 0 |
| II | 3091462 | C | T | 1 | 0 | 0 |
| II | 3141396 | C | T | 0 | 1 | 0 |
| II | 3194729 | G | A | 1 | 0 | 0 |

| | | | | | | |
|---|---|---|---|---|---|---|
| II | 3207303 | G | T | 0 | 1 | 0 |
| II | 3323243 | C | A | 0 | 1 | 0 |
| II | 3323274 | T | G | 0 | 1 | 0 |
| II | 3349288 | G | T | 1 | 0 | 0 |
| II | 3357187 | G | A | 1 | 0 | 0 |
| II | 3414754 | G | T | 1 | 0 | 0 |
| II | 3608749 | T | C | 1 | 0 | 0 |
| II | 3653326 | C | G | 1 | 0 | 0 |
| II | 3855013 | G | A | 1 | 0 | 0 |
| II | 3903996 | G | A | 1 | 0 | 0 |
| II | 3904114 | G | C | 1 | 0 | 0 |
| II | 3911202 | C | T,A | 0 | 0 | 1 |
| II | 3986676 | C | T | 0 | 1 | 0 |
| II | 4122326 | T | C | 1 | 0 | 0 |
| II | 4132718 | C | T | 0 | 1 | 0 |
| II | 4180262 | A | G | 0 | 1 | 0 |
| II | 4203147 | T | C | 0 | 1 | 0 |
| II | 4225787 | A | C | 0 | 1 | 0 |
| II | 4336867 | G | A | 1 | 0 | 0 |
| II | 4400666 | A | C | 1 | 0 | 0 |
| II | 4534970 | C | T | 0 | 0 | 1 |
| II | 4534990 | A | G | 0 | 0 | 1 |
| II | 4535001 | C | T | 0 | 0 | 1 |
| II | 4535023 | T | C | 0 | 0 | 1 |
| III | 38079 | C | G | 0 | 1 | 0 |
| III | 38385 | T | A | 0 | 0 | 1 |
| III | 41574 | C | A,T | 0 | 1 | 0 |
| III | 82090 | C | T | 1 | 0 | 0 |
| III | 114743 | G | T,A | 0 | 0 | 1 |
| III | 227730 | A | T | 1 | 0 | 0 |
| III | 284657 | C | A | 1 | 0 | 0 |
| III | 346648 | T | A,C | 0 | 0 | 1 |
| III | 346719 | T | A | 0 | 1 | 0 |
| III | 418867 | A | C | 0 | 1 | 0 |
| III | 510083 | G | T | 0 | 1 | 0 |
| III | 531936 | G | A | 0 | 1 | 0 |
| III | 564266 | A | G,T | 0 | 1 | 0 |
| III | 679772 | T | C | 1 | 0 | 0 |
| III | 867244 | T | G | 0 | 1 | 0 |
| III | 947336 | G | C | 1 | 0 | 0 |
| III | 954518 | A | G | 0 | 0 | 1 |
| III | 954527 | C | T | 0 | 0 | 1 |
| III | 1174401 | C | T | 0 | 0 | 1 |
| III | 1176072 | C | G | 0 | 0 | 1 |
| III | 1176576 | G | A | 0 | 0 | 1 |
| III | 1250922 | C | T | 1 | 0 | 0 |
| III | 1346913 | G | A | 0 | 1 | 0 |
| III | 1374938 | A | T | 1 | 0 | 0 |
| III | 1377097 | G | A | 1 | 0 | 0 |
| III | 1492509 | T | C | 0 | 0 | 1 |
| III | 1793354 | T | A | 1 | 0 | 0 |
| III | 1797171 | C | A | 1 | 0 | 0 |
| III | 1820329 | G | T | 1 | 0 | 0 |
| III | 1862803 | A | G | 1 | 0 | 0 |
| III | 1927053 | C | T | 1 | 0 | 0 |
| III | 1947259 | G | A | 0 | 1 | 0 |
| III | 2055592 | A | T | 0 | 1 | 0 |
| III | 2128341 | C | T | 1 | 0 | 0 |
| III | 2131559 | C | T | 1 | 0 | 0 |
| III | 2153109 | C | T | 0 | 1 | 0 |
| III | 2221426 | G | A | 0 | 0 | 1 |
| III | 2316266 | T | C | 0 | 0 | 1 |
| III | 2362468 | C | T | 1 | 0 | 0 |
| III | 2426557 | T | A | 0 | 0 | 1 |
| III | 2428702 | G | T | 0 | 1 | 0 |

| | | | | | | |
|---|---|---|---|---|---|---|
| III | 2428915 | A | G | 0 | 1 | 0 |
| III | 2429197 | T | C | 0 | 0 | 1 |
| MT | 1788 | C | T | 1 | 0 | 0 |
| MT | 2091 | A | G | 1 | 0 | 0 |
| MT | 4102 | C | G | 1 | 0 | 0 |
| MT | 6906 | A | T | 1 | 0 | 0 |
| MT | 8529 | G | T | 1 | 0 | 0 |
| MT | 10309 | T | C | 1 | 0 | 0 |
| MT | 10310 | C | T | 1 | 0 | 0 |
| MT | 13751 | C | T | 1 | 0 | 0 |
| MT | 15200 | G | T | 1 | 0 | 0 |
| MT | 18648 | G | A | 1 | 0 | 0 |
| MT | 19152 | A | G | 1 | 0 | 0 |
